# A developmentally programmed splicing failure attenuates the DNA damage response during mammalian zygotic genome activation

**DOI:** 10.1101/2020.11.25.397794

**Authors:** Christopher D. R. Wyatt, Barbara Pernaute, André Gohr, Marta Miret-Cuesta, Lucia Goyeneche, Quirze Rovira, Ozren Bogdanovic, Sophie Bonnal, Manuel Irimia

## Abstract

The transition from maternal to embryonic transcriptional control is a crucial step in embryogenesis. However, how alternative splicing is regulated during this process and how it contributes to early development is unknown. Using transcriptomic data from pre-implantation stages of human, mouse and cow, we show that the stage of zygotic genome activation (ZGA) exhibits the highest levels of exon skipping diversity reported for any cell or tissue type. Interestingly, much of this exon skipping is temporary, leads to disruptive non-canonical isoforms, and occurs in genes enriched for DNA damage response in the three species. We identified two core spliceosomal components, *Snrpb* and *Snrpd2*, as regulators of these patterns. These genes have low maternal expression at the time of ZGA and increase sharply thereafter. Consistently, microinjection of *Snrpb/d2* mRNA into mouse zygotes reduces the levels of temporary exon skipping at ZGA, and leads to an increase in etoposide-induced DNA damage response. Altogether, our results suggest that mammalian embryos undergo an evolutionarily conserved and developmentally programmed specific splicing failure at the time of genome activation that attenuates cellular responses to DNA damage at these early stages.

## INTRODUCTION

Pre-implantation embryonic development follows a morphogenetically similar path in all placental mammals. It progresses from an unfertilized oocyte to a fertilized zygote through fusion with sperm, followed by symmetric cell divisions, morula compaction, establishment of the first cell lineages, and formation of a blastocyst that implants into the uterine wall. A distinctive control over the cell cycle and DNA damage response (DDR) has been observed during the first cell divisions ^1,2^, a moment in which the embryo needs to ensure cell cycle progression whilst preserving genome integrity, as any unrepaired damage will be inherited by all embryo lineages. These pre-implantation events are mirrored by large epigenetic and transcriptomic changes. Arguably, the most studied aspect is the transition from maternal to embryonic transcriptional control, at the stage of zygotic genome activation (ZGA). Here, the maternal mRNA contribution is cleared both actively and passively ^3^, and the zygotic genome activates in different waves ^4^. The relative timing of the major wave differs between species, disconnecting morphogenetic and transcriptomic events. For example, while morula compaction occurs roughly at the 8-cell (8C) stage in both human and mouse embryos, the major wave of ZGA takes place also at the 8C stage in human, but at the 2-cell (2C) stage in mouse. These differences, added to the difficulty of obtaining reliable transcriptomic data from pre-implantation embryos from multiple mammalian species, complicate transcriptomic evolutionary comparisons between species. Thus, despite some initial reports based on microarrays (e.g. ^5,6^), in-depth investigation of transcriptome-wide remodeling had remained elusive until the advent of single-cell and low-input RNA sequencing (RNA-Seq). Using these techniques, several studies have undertaken large transcriptomic analyses of pre-implantation development in human ^7–9^, mouse ^3,7,10–13^ and cow ^14–16^. These studies have confirmed some previous findings (e.g. the different timing of the ZGA in each species) and provided novel comparative insights. However, they focused nearly exclusively on the variations of steady-state levels of protein-coding genes.

Alternative splicing (AS) is the process by which different pairs of splice sites are selected in precursor mRNAs leading to different combinations of exons in the final mature mRNA. It is responsible for greatly expanding the functional and regulatory capacity of metazoan genomes ^17^, potentially generating numerous transcript and protein products from a single gene. Over half of human protein-coding genes produce multiple transcript isoforms that are widely regulated across cell and tissue types ^18^, with particularly high prevalence in brain and testis ^19,20^. These AS events may generate distinct functional protein isoforms or lead to unproductive mRNA products that are degraded by non-sense mediated decay (NMD), thereby contributing to modulate gene expression ^21^. Although proper AS regulation is crucial for post-implantation mammalian embryo development ^22^, only a handful of studies have descriptively assessed isoform diversity during pre-implantation stages, concluding that hundreds of genes dynamically express different transcripts in human and mouse, particularly at the maternal-to-zygotic transition ^8,12,23–25^. However, the regulatory mechanisms, evolutionary conservation and physiological implications of these patterns are unknown.

Here, we generated a comprehensive dataset of AS quantifications for pre-implantation development of human, mouse and cow. We found that the blastomeres that undergo the major wave of ZGA show the highest levels of exon skipping reported so far for any cell and tissue type. In most cases, this exon skipping was temporary and restored soon after ZGA. Remarkably, such AS events often disrupt the open reading frame (ORF) and are enriched for genes involved in DDR in the three studied species. We identified Sm ring components *Snrpb* and *Snrpd2* as major regulators of these patterns, and showed that induced expression of these factors prior to ZGA leads to reduced splicing disruption and increased DDR upon etoposide treatment at this developmental point.

## RESULTS

### AS profiles in early embryo development of human, mouse and cow

To investigate AS during pre-implantation development, we took advantage of the abundant publicly available RNA-Seq datasets (Fig. 1a and Supplementary Table 1). These data comprised samples from oocyte to blastocyst stage embryos from multiple studies, obtained from either single blastomere or bulk embryo RNA-Seq. To confidently estimate AS levels across the time courses, we performed the following steps (Fig. 1b; see Methods for details). We first measured gene steady-state mRNA levels (hereafter referred to as gene expression, GE) genome-wide for each individual sample, and clustered these samples using hierarchical clustering (Supplementary Figs. 1-3). Samples largely grouped by stage and not by experiment, and outlier cells were removed from further study (Supplementary Table 1 for details). Based on this information, we merged single cells/samples into pools of ~160 million reads on average (Supplementary Table 1) to acquire high coverage on exon-exon junctions and improve quantifications of AS. Principal Component Analyses (PCA) of GE measurements for the pooled groups showed a V-shaped temporal profile for PCs 1 and 2, with the largest difference occurring at the time of ZGA in the three species (Fig. 1c; between the 4-cell (4C) and 8C stage in human, zygote and 2C stage in mouse, and 4C and 8C/morula in cow).

**Figure 1.**
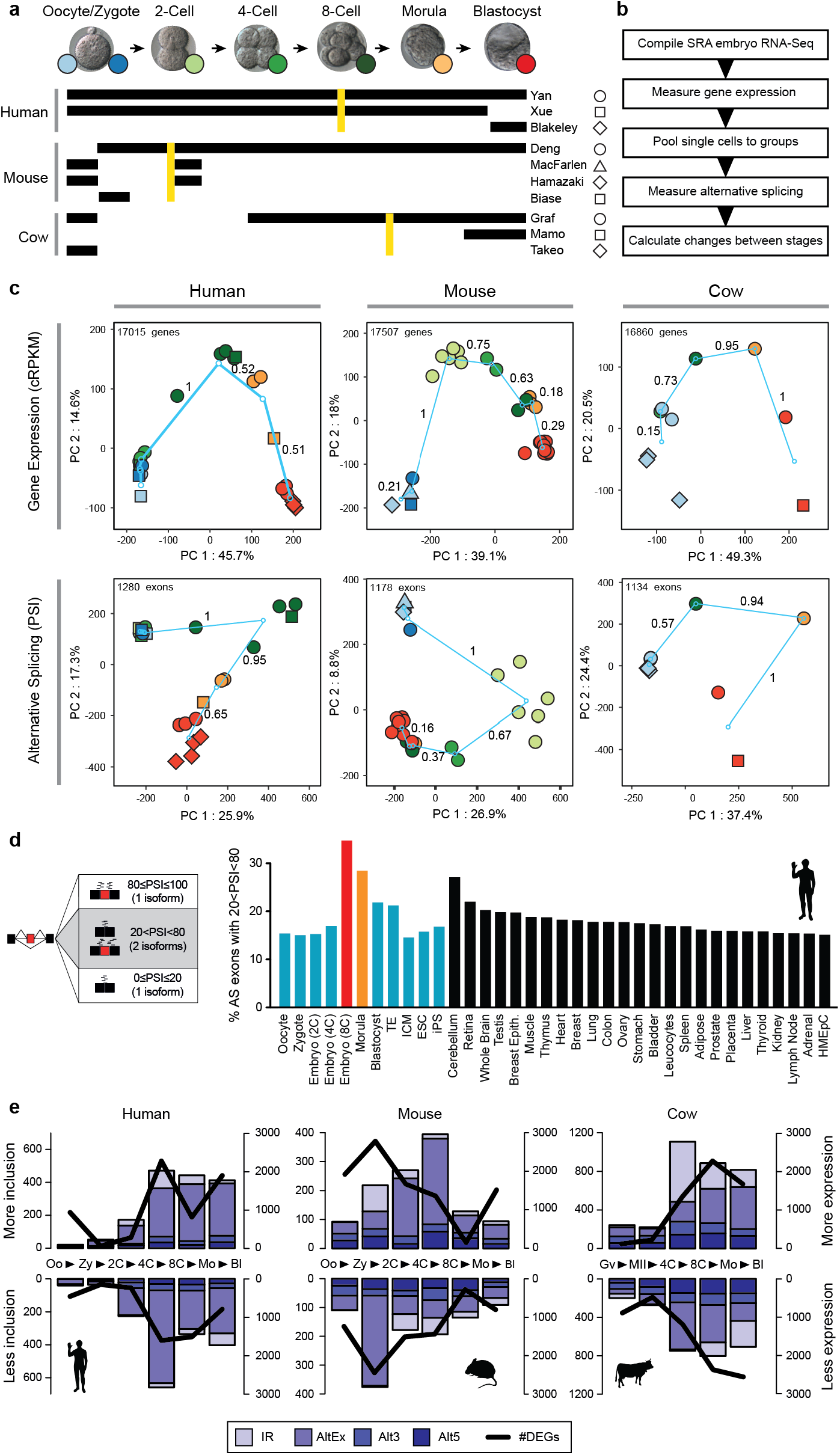
AS profiles during early development reveal the highest exon skipping diversity at ZGA. **(a)** Summary of datasets used in this study for each species. Golden lines indicate the ZGA for each species. **(b)** Schematic view of the methodological steps taken to process the RNA-Seq data and obtain AS quantifications with *vast-tools*. **(c)** Principle component analysis (PCA) for the merged groups of samples per stage for GE (top; cRPKMs) and AS (bottom; exons with sufficient read coverage in at least 80% of the samples). Color codes and shapes for each stage as depicted in (a). Turquoise circles represent the centroids of each stage, and lines show the change between consecutive stages, with values representing the relative change respect to the largest change (=1). **(d)** Percentage of exons with 20<PSI<80 (i.e. generating two substantial isoforms) in each sample was calculated for every stage and differentiated tissue, showing the highest relative exon skipping levels for the ZGA stage. Equivalent plots for mouse and cow are shown in Extended Data Fig. 2a. **(e)** Numbers of alternative exons (AltEx), alternative 3/5′ splice sites (Alt3/5) and retained introns (IR) with increased/decreased inclusion levels in consecutive pairwise transitions (bars, left Y-axes) or genes with higher/lower expression (DEGs) (line, right Y-axes). Abbreviations: Oo, oocyte; Zy, zygote; Mo, morula; Bl, blastocyst; Gv, oocyte GV; MII, oocyte MII.

We then used *vast-tools* ^18,26^ to quantify alternative sequence inclusion levels (using the percent-spliced-in metric, PSI) for alternative exons, alternative donor/acceptor sites and retained introns (Extended Data Fig. 1a). In addition, to ensure that early embryo-specific exons were not missed from *vast-tools* annotations, we conducted a *de novo* search for cassette exons using a custom pipeline (see Methods for details; Supplementary Table 2). Similar to GE, PCAs of exon PSIs separated embryos by cell stage and not experiment, showed V-shaped temporal dynamics, and had the largest difference between consecutive stages at ZGA for the three species (Fig. 1c). To facilitate the access of this large resource to the research community, AS and GE plots are provided as special datasets in VastDB (http://vastdb.crg.eu).

### ZGA stage embryos show the highest levels of exon skipping diversity of any cell and tissue type

To have a first assessment of the contribution that AS has to diversify transcriptomes at each developmental stage, we used a simple measure of diversity where alternative exons with sufficient read coverage were classified as either producing one main isoform (PSI≤20 or PSI≥80) or two (20<PSI<80) (Fig. 1d). Remarkably, the stage undergoing the ZGA showed the highest levels of exon skipping diversity in the three studied species (8C in human, 2C in mouse and 8C in cow), returning to the preceding lower levels in the subsequent stages (Fig. 1d and Extended Data Fig. 2a). Furthermore, the level of AS of cassette exons upon ZGA was not matched by any differentiated cell or tissue type, including neural, muscle and testis (Fig. 1d and Extended Data Fig. 2a). The increased level of AS among alternative exons was robust to different cut-offs of PSI range and read coverage (Extended Data Fig. 2b), also observed at the single blastomere level (Extended Data Fig. 2c), and was not found for intron retention or alternative 3′/5′ splice site choices (Extended Data Fig. 2b). Altogether, these results reveal that transcriptome diversification driven by exon skipping reaches its maximum in mammals only for a brief moment in life during ZGA.

**Figure 2.**
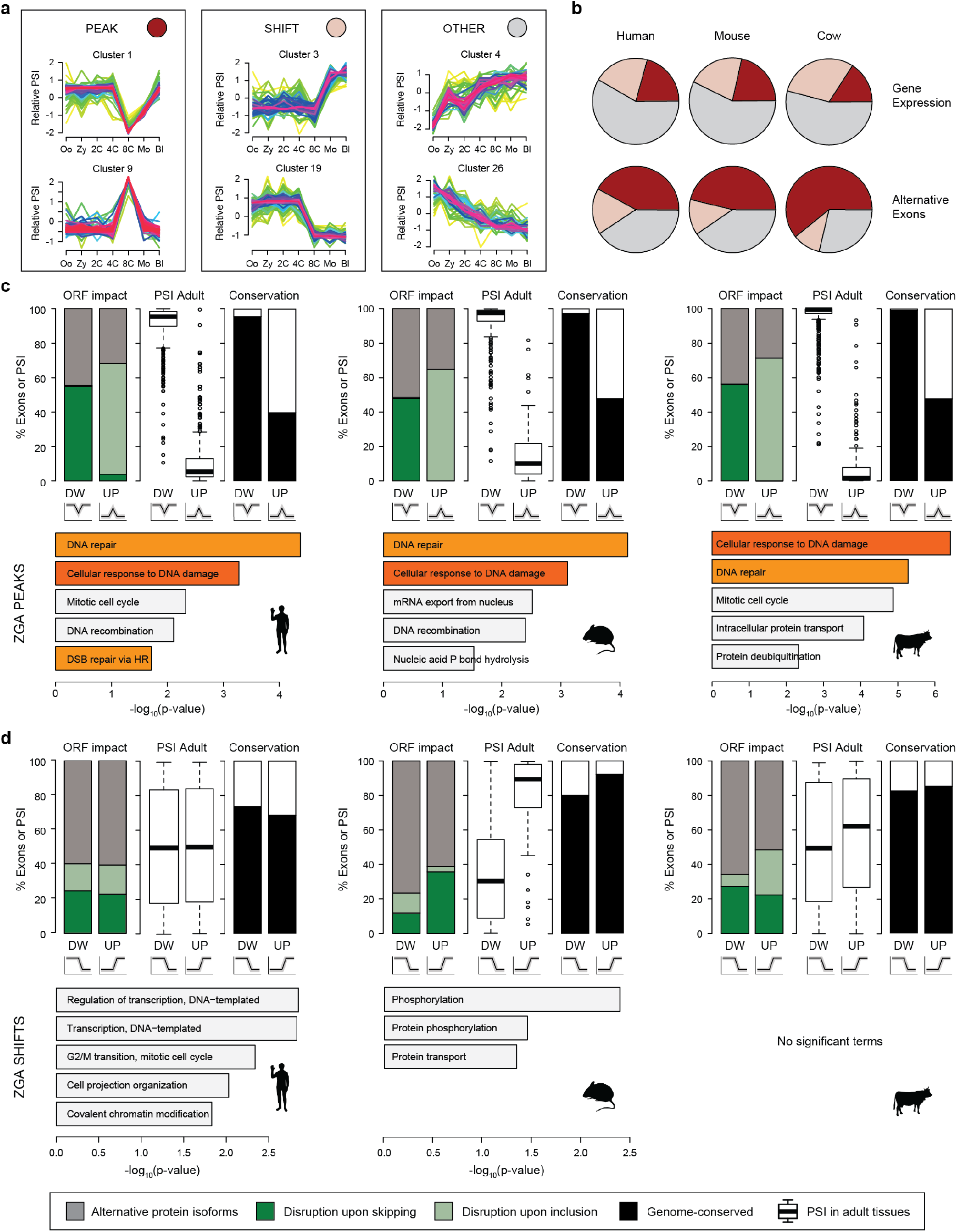
Exons peaking at ZGA disrupt the ORFs and are enriched for DNA damage response genes. **(a)** Representative examples of the three main Mfuzz cluster types (Peak, Shift and Other). **(b)** Proportions of each type of Mfuzz cluster for GE (top) and alternative exons with sufficient read coverage in all stages (bottom). Abbreviations: Oo, oocyte; Zy, zygote; Mo, morula; Bl, blastocyst; Gv, oocyte GV; MII, oocyte MII. **(c,d)** For each species, several features are shown for exons in Mfuzz clusters with Peak (c) or Shift (d) dynamics at ZGA, either with decreased (DW) or increased (UP) PSI. Top: percentage of exons in coding sequences predicted to disrupt the ORF upon inclusion or skipping or to produce alternative protein isoforms (left), distribution of median PSIs in adult tissues (center), and percentage of exons with genomic conservation in any of the other two species (right). Bottom: enriched Gene Ontology (GO) categories. Orange bars indicate DNA damage/repair related categories.

### Changes in GE and AS are maximal at ZGA

We next measured the number of AS events that change between each pair of consecutive stages (hereafter stage transitions) in our time course (Extended Data Fig. 1b and Methods). In total, we found 2711, 1828 and 4748 unique AS events of all types with differential regulation in 2058, 1350 and 2735 genes for human, mouse and cow, respectively (Fig. 1e). For comparison, we also calculated the numbers of differentially expressed genes (Extended Data Fig. 1c) and found 6545, 8895 and 8118 genes in each respective species that showed differential expression in at least one transition. Consistent with the PCA results, we found the largest number of changes for both AS and GE at ZGA in the three species (Fig. 1e). For example, in human, 1130/2711 (41.7%) of AS events with differential regulation and 3826/6545 (58.5%) of genes differentially expressed were observed at ZGA (4C-8C transition). Moreover, a clear bias was observed in the direction of regulation for alternative exons and intron retention in the three species at this transition: whereas the majority of exons showed increased skipping at ZGA, most differentially spliced introns had increased retention (Fig. 1e). Despite AS and GE changes mostly occurring at ZGA, the specific genes with AS and GE changes did not significantly overlap for nearly all transitions, a pattern that was consistent for intron retention and exon skipping separately (Extended Data Fig. 3a-c). Given that a large number of genes become transcribed at this stage for the first time, these results imply that, in many cases, the maternally inherited transcript isoform is (partially) substituted by a new isoform without significantly altering the overall mRNA steady state level of the gene.

**Figure 3.**
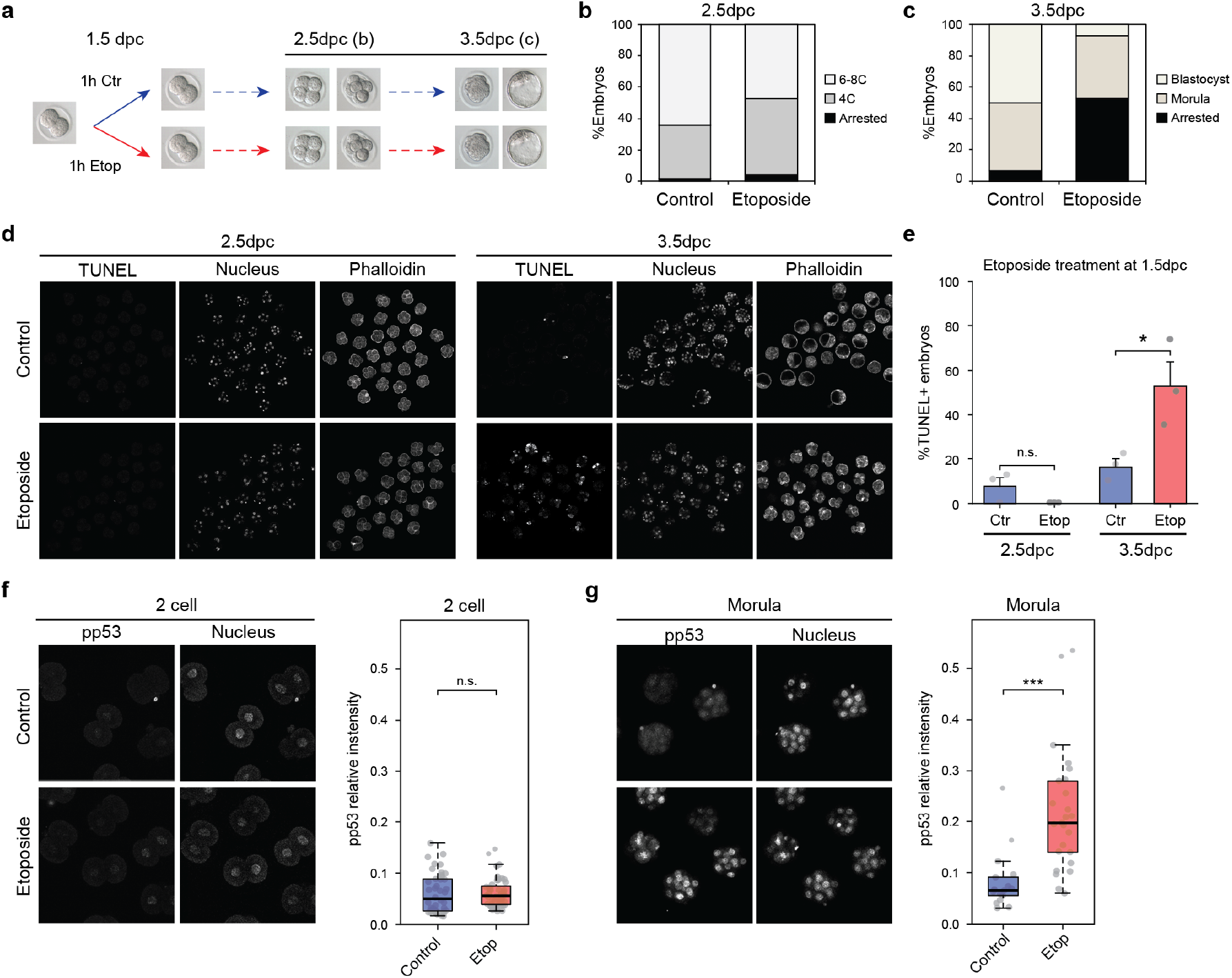
Mouse embryos at ZGA have a lower response to DNA damage induced by etoposide. **(a)** Scheme of etoposide treatment and analysis of embryo recovery: 2C embryos (1.5 days post coitum, dpc) were either left un-treated or treated with 0.5uM etoposide for 1h and then washed and left to recover for further 48h. **(b,c)** Proportion of embryos found arrested or developed to the stated stage 24h (b) or 48h (c) post-treatment. Stack plots show the average of 6 independent experiments with a total number of embryos analysed of n=105 (control) and n=113 (treated). All embryos not reaching the expected developmental stage at a given time point either because of cell death or cell cycle arrest were classified as “Arrested”. **(d)** TUNEL staining of embryos in (b) and (c), showing a high incidence of cell death in treated embryos only 48h, but not 24h, post-etoposide treatment. **(e)** Percentage of embryos showing positive TUNEL staining at each developmental stage. Average of three independent experiments is shown with a total number of embryos of: n=72 (control, 2.5dpc), n=63 (etoposide, 2.5dpc), n=67 (control, 3.5dpc) and n=50 (etoposide, 3.5dpc). * *P*=0.036 two-sided Student’s T-test. **(f,g)** Immunostaining for phospho-p53 (Ser15; pp53) in 2C (f) or early morula (g) stage embryos treated with 0.5uM etoposide for 1h or left untreated. Quantification of phospho-p53 (Ser15) levels from embryos is shown for each stage. Each dot represents the average relative intensity of all cells of a single embryo. Boxplots show staining levels of embryos from three independent experiments. Number of embryos: n=42 (control, 2C), n=47 (etoposide, 2C), n=17 (control, morula) and n=25 (etoposide, morula). *** (0.001 < *P*) based on Wilcoxon Rank Sum tests.

### Conserved and species-specific changes in AS

We next asked whether changes in AS and GE were conserved across species or were specific to individual species. For exons changing in each transition for a given species, we assessed in each other species: (i) if the exons were present in their genome (“genome-conserved” ^27^), (ii) if these orthologous exons had sufficient read coverage, and, if so, (iii) if they changed (|ΔPSI| > 15) in the same direction at any transition (Extended Data Fig. 4a-f; see Methods for details). This analysis showed that most AS events changing in two species change at ZGA, but highlighted overall low levels of conservation. For instance, only 3.6% and 5.1% (31 and 44 out of 859) of ZGA-regulated exons in human also change their inclusion in the same direction at the mouse or cow ZGA, respectively. These percentages increase up to 15.8% and 30.1%, respectively, for exons with an ortholog and sufficient read coverage at the ZGA in the other species. In the case of GE, 27.2% and 35.0% of genes with differential expression at ZGA in human overlapped with those changing at ZGA in mouse and cow, respectively, although a high level of heterochronies was also observed (Supplementary Fig. 4), as previously reported ^5^. To further identify exons that were dynamically regulated during pre-implantation development across mammals, we next searched for orthologous exons whose inclusion levels were different between any two stages in the three species (see Methods). This revealed 259 exons (Extended Data Fig. 4g and Supplementary Table 3), which were significantly enriched for genes involved in key signalling pathways (e.g. Wnt pathway), transcription and chromatin modifiers (e.g. *Tcerg1*, *Dnmt3b*, *Ezh2*), and genes related to morphogenesis (e.g. cadherin binding) (Extended Data Fig. 4h).

**Figure 4.**
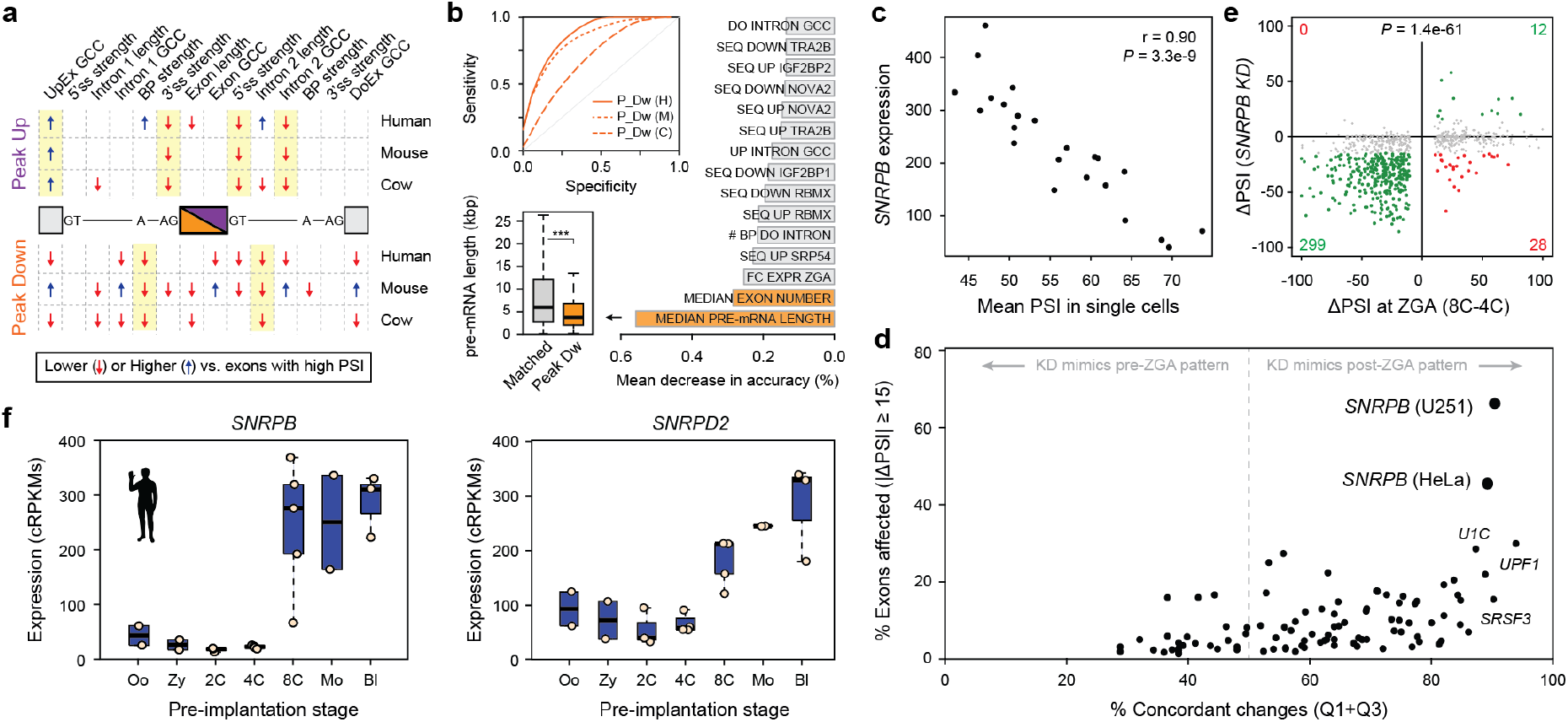
*Snrpb* and *Snrpd2* are activated during ZGA and are associated with peak exons. **(a)** Schematic summary of intron-exon features associated with ZGA peak-down or peak-up exons for the three species. Blue/red arrows indicate features with statistically significantly higher/lower median values compared to exons with high PSI across development and differentiated tissues based on two-sided Mann–Whitney U tests. Full comparisons with exons with high/low PSI as well as control sets with matched pre-ZGA PSIs and value distributions are shown in Supplementary File 1. **(b)** Top inset: receiver operating characteristic curves (ROCs) for Random Forest classifiers for peak-down exons in the three species (H: human, AUC = 0.852; M: mouse, AUC = 0.818; C: cow, AUC = 0.678). Barplot: top ranking features based on mean decrease in accuracy averaged for the three species. Bottom inset: distribution of pre-mRNA lengths for human peak-down exons and a control set of exons with matched pre-ZGA PSI distributions. **(c)** Correlation at the single cell level between human *SNRPB* expression and the mean PSI for peak-down exons. **(d)** Summary of associations between human splicing factor knockdowns and exons from Mfuzz clusters that peak at ZGA (up and down). Y-axis: percentage of exons affected by the knockdown (|ΔPSI| ≥ 15; Y-axis). X-axis: concordance of the direction of change in the knockdown respect to the change at ZGA (in Cartesian axes as per (e), those in quadrants 1 and 3 [Q1+Q3]). **(e)** Scattered plot showing ΔPSI of exons from Mfuzz cluster that peak at ZGA upon *SNRPB* knockdown in U251 cells (Y-axis) versus ΔPSI at ZGA (8C-4C; X-axis). Exons with |ΔPSI| ≥ 15 upon knockdown are highlighted in green/red and numbers indicated. p-value corresponds to a two-sided Binomial test between Q1+Q3 versus Q2+Q4. U251 data from ^67^. **(f)** Expression of *SNRPB* and *SNRPD2* in human early development. Equivalent plots for mouse and cow are shown in Extended Data Fig. 8d.

### Peak profiles dominate AS dynamics

To further characterize the temporal dynamics of AS, we used Mfuzz ^28^ to cluster alternative exons according to their co-regulated inclusion levels throughout the time course. We obtained 28, 18 and 22 exon clusters in human, mouse and cow, respectively (Supplementary Figs. 5-7; Supplementary Table 4). Most of these clusters could be broadly classified into three general patterns: (i) peak-like regulation, in which the exon is highly included or skipped only in a given stage, quickly returning to the initial levels (hereafter “peak exons”), (ii) shift-like regulation, in which the exon goes from high to low inclusion at a specific transition, or vice versa, but does not return to the initial level (“shift exons”), and (iii) other regulation, those that do not fit the previous descriptions (Fig. 2a; see Methods for precise definitions). Remarkably, the peak behaviour was the most common regulation among alternative exons (Fig. 2b; e.g. 42.1% vs 17.3% of shift exons in human). This contrasted with the patterns of Mfuzz clusters based on GE, which were more represented by shift or other behaviours (Fig. 2b). These results were consistent with the V-shape patterns in the PCA (Fig. 1c) and the asymmetric patterns observed for exon skipping and intron retention at ZGA, which were inverted in the post-ZGA transition in the three species (Fig. 1e), suggesting that a large proportion of changes at this time are temporary. Indeed, 427/749 (57%) alternative exons with significant changes at ZGA covered in the Mfuzz cluster analysis showed peak behaviour. Mfuzz clustering patterns were highly validated by RT-PCR using RNA from independent pools of embryos, with 21/23 (91%) of peak, 11/11 (100%) of shift and 19/19 (100%) other alternative exons showing the expected temporal dynamics (Extended Data Fig. 5).

**Figure 5.**
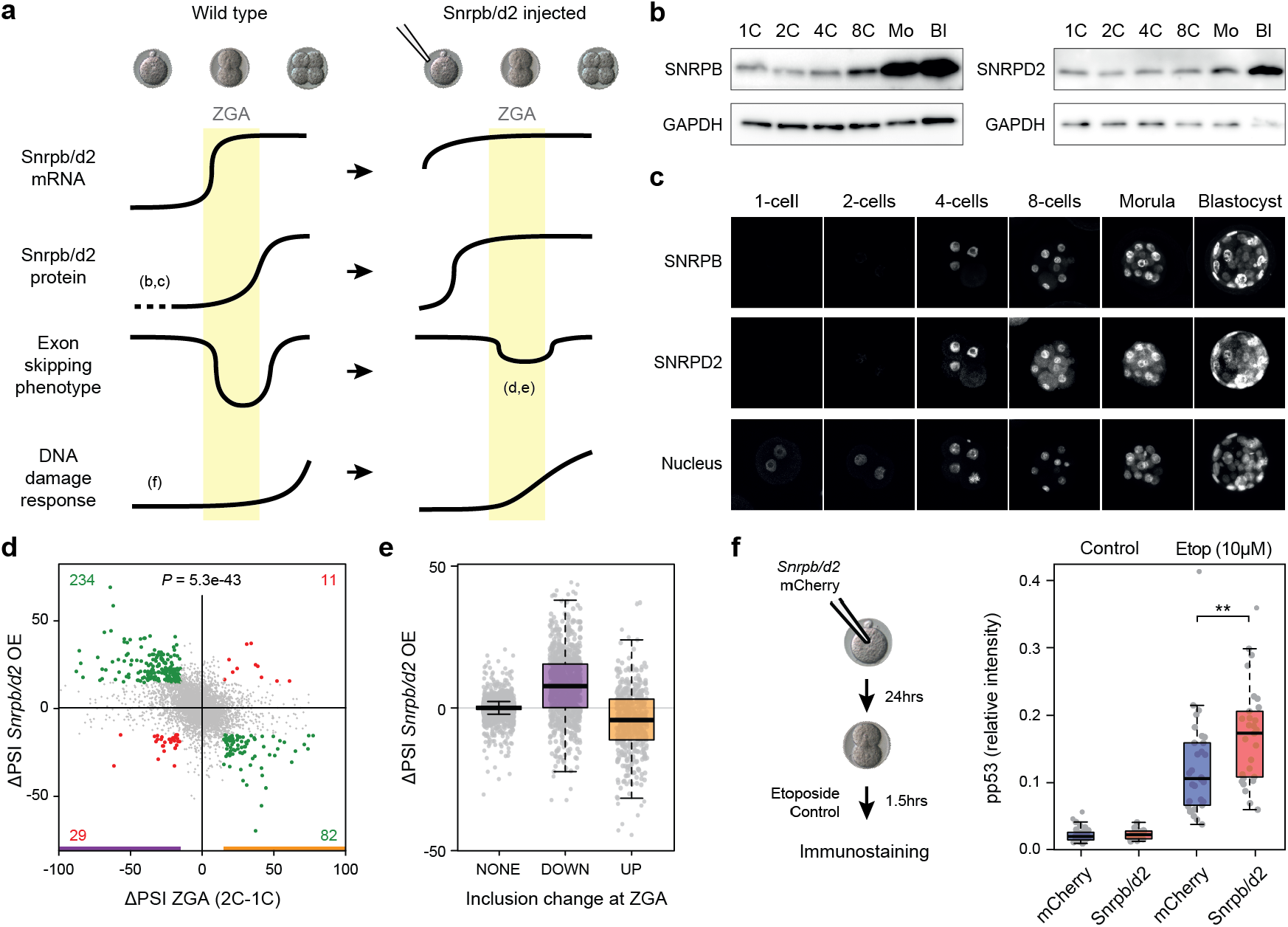
*Snrpb* and *Snrpd2* are partly responsible for peak-down AS patterns affecting DNA damage response at ZGA. **(a)** Working hypothesis for peak-like alternative exon regulation during ZGA in mouse. Left (Wild type): *Snrpb*/*d2* have low mRNA and protein levels before ZGA. At ZGA, their mRNA levels are increased, followed by a rise in protein levels. This period with low levels of *Snrpb*/*d2* protein leaves a time window in which sensitive exons in newly transcribed genes get excluded due to insufficient amount of these regulators. This situation starts being restored after ZGA, when *Snrpb*/*d2* protein levels reach sufficient amounts. Given their specific functional enrichment, aberrant exclusion of exons in DDR genes during ZGA will contribute to a delay in the response to DNA damage at these developmental stages. Right: induced expression of *Snrpb*/*d2* prior to ZGA should prevent the abovementioned exon skipping in DDR genes and therefore allow for an earlier stronger response to DNA damage. **(b,c)** Western blot and immunostaining showing SNRPB and SNRPBD2 levels during pre-implantation development in mouse. Pools of 150 or 60 embryos per stage were used for the SNRPB and SNRPD2 western blots respectively. **(d)** Changes in exon inclusion levels at 2C stage upon early induced expression of *Snrpb/d2* (Y-axis) respect to the observed change at ZGA (2C-1C; X-axis). Exons with |ΔPSI| ≥ 15 upon *Snrpb/d2* expression and at ZGA are highlighted in green/red and numbers indicated. p-value corresponds to a two-sided Binomial test between Q2+Q4 versus Q1+Q3. **(e)** Distribution of ΔPSI upon *Snrpb/d2* induced expression for all exons with increased (UP), decreased (DOWN) or no change (NONE) in PSI at ZGA. **(f)** Pronuclear stage embryos were co-injected with *Snrpb/d2* mRNA or injected with mCherry mRNA and left to develop to the 2C stage, when they were treated for 1.5h with 10uM etoposide or left untreated. Boxplots show the quantification of phospho-p53 (Ser15) relative intensity from immunostaining of embryos in each condition. Each dot represents the average relative intensity of both cells of a single embryo. Data from three independent experiments is shown (individual experiments are shown in Extended Data Fig. 10a). Total number of embryos for each condition: n=29 (mCherry untreated), n=26 (Snrpb/d2, untreated), n=30 (mCherry, etoposide) and n=29 (Snrpb/d2, etoposide). ** *P* = 0.0034, two-sided Wilcoxon Rank Sum test.

### Peak AS changes at ZGA often disrupt the ORF and are enriched for DDR genes

To begin elucidating the potential functional impact of AS changes during early embryogenesis, we investigated the predicted effect that alternative sequence inclusion or exclusion had on ORFs at each stage. Although the fractions of AS events that are predicted to disrupt the ORF were similar across stage transitions and species (Extended Data Fig. 6a), we found strong biases in the direction of the disruption depending on the transition: the vast majority of non-frame preserving AS events changing at ZGA disrupt the ORF specifically at that stage in the three species (Extended Data Fig. 6b). That is, these alternative sequences were more included at ZGA when the inclusion was predicted to disrupt the ORF, and vice versa for exclusion. Consistent with these predictions, isoforms predicted to disrupt the ORF at ZGA showed strong upregulation upon NMD disruption through UPF1 knockdown in HR1 cells (Extended Data Fig. 6c). Moreover, the disruptive impact on ORFs seems further strengthened by a global differential engagement of isoforms in translating ribosomes. Comparison of RNA-Seq data from high and low polysome fractions from human embryonic stem cells ^29^ or HEK293 cells ^30^ revealed a strong bias for ORF-disrupting isoforms at the ZGA to be less engaged by translating ribosomes, whereas the opposite was true for the few ORF-recovering isoforms (Extended Data Fig. 6d,e).

Analysis of the subsets of exons that belonged to different Mfuzz clusters changing at ZGA further informed these patterns. Most peak exons disrupted the ORF at the peaking stage in the three species, whereas shift exons more often generated alternative protein isoforms (Fig. 2c,d, top-left). In addition, peak-down exons were usually constitutively or highly included in differentiated tissues, whereas peak-up exons were normally not or only lowly included in differentiated tissues (Fig. 2c, top-middle). Interestingly, the biased inclusion levels in peak exons occurred for both ORF-preserving and ORF-disrupting events, underscoring a widespread change from canonical to rare isoforms at this stage. In line with this, ZGA isoforms were found to be upregulated upon NMD depletion and depleted in the high polysome fraction in both embryonic stem cells and 293T cells, irrespective of the predicted ORF impact (Extended Data Fig. 6f-h). Two additional lines of evidence supported the opposite nature of peak-down and peak-up exons. First, each cluster type had distinct profiles of overlap with transposable elements: peak-down exons were strongly depleted for transposable elements, whereas peak-up exons were enriched for these genetic elements (Extended Data Fig. 7). Second, peak-down exons had the highest levels of genome conservation among mammals, whereas most peak-up exons were species-specific (Fig. 2c, top-right). Therefore, altogether, these patterns are consistent with most peak-down exons being constitutive exons of major coding relevance and whose skipping at ZGA leads to non-functional isoforms, whereas peak-up exons are cryptic-like exons that do not encode important protein domains and often disrupt the gene’s canonical isoforms when included.

Remarkably, despite the low level of regulatory conservation of individual events (see above), exons in clusters peaking at ZGA were altogether significantly enriched for genes involved in DNA repair and cellular response to DNA damage in human, mouse and cow (Fig. 2c, bottom). These GO enrichments contrast sharply with those for genes with shift events, with terms related to transcription or protein phosphorylation, and whose regulated exons had more intermediate inclusion levels across adult cell and tissues (Fig. 2d). In summary, these results show that a large number of peak exons are likely to disrupt protein function temporarily upon ZGA, impacting genes involved in response to DNA damage.

### Reduced DNA damage response to etoposide during ZGA

The predicted impact of AS on the function of proteins involved in DDR is especially remarkable as previous studies indicate that early pre-implantation embryos present a delayed response to DNA damage induced by irradiation ^31–33^, suggesting that this process may not be fully functional during ZGA. To gain further insights into DDR regulation during ZGA, we first treated 2C embryos with low doses of the topoisomerase II inhibitor etoposide for 1h and analyzed the effect on their developmental progression for further 48h (Fig. 3a). Etoposide treated embryos progressed to the 8C stage at the same rate as control embryos (Fig. 3b) and only became arrested after early morula compaction (Fig. 3c). In addition, TUNEL staining showed cell death was suppressed until morula stage, when a large proportion of etoposide treated embryos were TUNEL positive (Fig. 3d,e). Etoposide induces DNA double strand breaks (DSBs), which primarily get resolved though the activation of the ATM pathway ^34^. P53 is one of the main downstream targets that get phosphorylated by ATM in response to DSBs ^35^. We therefore used the levels of p53 phosphorylation at Ser15 to ask whether embryos at the time of ZGA display a correct activation of ATM pathway in response to etoposide. After treatment of 2C embryos or early morulas (8 to 16 cells) for 1h with low dose etoposide, the level of phospo-p53 (Ser15) was significantly higher in treated morulas whereas it remained low in treated 2C embryos (Fig. 3f,g). These results thus support a dysfunction of ATM–p53-mediated DDR to DSBs during ZGA, which only becomes fully active from the early morula stage, when the embryos resolve the accumulated damage by inducing cell cycle arrest and death. Mechanistically, AS-mediated protein disruption at ZGA could contribute to lowering the overall levels of functional DDR proteins at this developmental stage, reducing the response to DNA lesions.

### Peak exon behavior is dependent on low *SNRPB* and *SNRPD2* levels

To obtain insights into the characteristics and regulation of peak exons at ZGA, we first evaluated multiple exonic and intronic features associated with exon skipping using *Matt* ^36^. This analysis revealed some common patterns across the three species, including weaker branch points and shorter downstream introns for exons peaking down at ZGA, and weaker 5′ and 3′ splice sites and differences in GC content for those peaking up (Fig. 4a and Supplementary File 1). Remarkably, a Random Forest classifier based on multiple genomic features was able to discriminate human and mouse peak exons from sets of exons with matched pre-ZGA PSIs with high sensitivity and specificity (e.g. AUC = 0.852 and 0.885 for human peak-down and peak-up exons, respectively; Fig. 4b and Extended Data Fig. 8a). Investigation of the features that contributed the most to this discrimination revealed the length of pre-mRNA and intron number as top-ranking characteristics for peak-down exons, which were located in significantly shorter genes in the three species (Fig. 4b, Extended Data Fig. 8b,c and Supplementary File 1).

We next investigated which splicing regulators may be responsible for the specific temporal dynamics of peak exons at ZGA. For this purpose, we took several complementary approaches (see Methods for details). First, for each species, we looked for enrichment of known binding motifs for RNA binding proteins ^37^ in the exonic and neighboring intronic regions for exons from Mfuzz clusters peaking up or down at ZGA. Although a few significantly enriched motifs were found for individual species, the associated RNA binding proteins did not show changes in GE at ZGA and/or the enrichments were not evolutionarily conserved (Supplementary Table 5). Second, we correlated average PSIs of peak-down or peak-up exons with GE of known splicing regulators at the ZGA at the single cell level (Supplementary Table 6). This identified multiple significant correlations at each transition, including several AS factors (e.g. *SRSF2*, *TIAL1*) as well as core spliceosomal components (e.g. *SNRPB*; Fig. 4c). Third, since these correlations may be indirect, we collected RNA-Seq data from 119 available experimental perturbations for 84 unique splicing regulators in different human cell or tissue types (Supplementary Table 1). For each regulator and experiment, we calculated the average change in PSI (or ΔPSI) between knockdown and control for all alternative exons, and overlapped those with significant changes with exons peaking at ZGA (Supplementary Table 7). Strikingly, exons changing upon *SNRPB* knockdown showed the strongest association with ZGA peak exons in two independent available experiments (Fig. 4d). Noticeably, the majority of overlapping exons corresponded to peak-down exons that are skipped upon *SRNPB* depletion (Fig. 4e; lower left quadrant).

Given this strong association, we decided to investigate the potential role of *SNRPB* at ZGA in more detail. *SNRPB*, also known as SmB, is part of the Sm heptameric ring, which is required for the biogenesis of the U1, U2, U4/U6, and U5 snRNP molecules on the pre-mRNA ^38^. Contrary to our initial expectation, *SNRPB* was very lowly expressed at pre-ZGA stages and showed a sharp increase in expression at the ZGA stage in human, mouse and cow (Fig. 4f and Extended Data Fig. 8d). In addition, another gene encoding a subunit of the Sm ring, *SNRPD2*, had similar activation patterns at ZGA in the three species (Fig. 4f and Extended Data Fig. 8d). Considering all these data together, we hypothesized the following scenario (Fig. 5a, left). Genes with peak-down *SNRPB/D2*-sensitive exons become actively transcribed at ZGA, when the maternally inherited levels of the two Sm proteins are low. This results in the production of transcripts that skip these exons, leading to a net decrease in their inclusion levels compared to pre-ZGA stages. As development proceeds, the burst of expression of *SNRPB/D2* genes at ZGA leads to a subsequent increase in Sm protein levels, which, in turn, allows the correct inclusion of peak exons in nascent transcripts at post-ZGA stages, thereby eventually restoring pre-ZGA high inclusion levels. In this way, the dynamic production and turnover of transcripts from genes containing *SNRPB/D2*-sensitive exons, together with the expression dynamics of the *SNRPB/D2* genes themselves, is thus predicted to result in temporary peak-like patterns of exon skipping around ZGA stages.

To assess this hypothesis, we tested the major predictions made by this model. First, we checked whether or not *Snrpb* and *Snrpd2* were depleted at the protein level at pre-ZGA stages. Western blot and immunostaining at different stages confirmed very low levels of both proteins until the late 2C/4C stage, with a sharp increase by the 8C/morula stage (Fig. 5b,c). Next, we evaluated the effect that increasing SNRPB and SNRPD2 levels prior to ZGA had on exons changing at this particular stage (Fig. 5a, right). For this, we injected *in vitro* transcribed mRNA from either *Snrpb* and *Snrpd2* together (*Snrpb/d2*) or mCherry, as a control, into pronuclear stage embryos, and sequenced polyA+ mRNA from zygotes (5h post mRNA injection) and 2C stage embryos (24h post mRNA injection). We identified 878 and 484 alternative exons decreasing and increasing their inclusion levels (|ΔPSI| >= 15), respectively, between our zygote and 2C stage embryos. Remarkably, of these, 234/878 (26.7%) exons with decreased PSI were less skipped after *Snrpb/d2* overexpression compared to the control condition (in contrast to only 29 with further skipping). Moreover, 82/484 (16.9%) exons that increased their inclusion from 1C to 2C decrease their PSI in *Snrpb/d2* injected embryos (compared to 11 in the opposite direction) (Fig. 5d,e). The negative association between the changes at ZGA and upon *Snrpb/d2* overexpression is highly significant (*P* = 1.1e-30, binomial test), suggesting that higher *Snrpb/d2* levels before ZGA maintain alternative exon patterns in a more pre-ZGA state. A similar, but milder, reversion of ZGA AS patterns was observed upon early expression of *Snrpb* or *Snrpd2* alone (Extended Data Fig. 9a,b). Importantly, this pattern was not observed for GE (Extended Data Fig. 9c), for which very few changes were identified upon *Snrpb/d2* overexpression (29/5,048 [0.58%] of ZGA changing genes), indicating that the reversion was specific to exon inclusion.

### Earlier *Snrpb/d2* expression leads to increased DNA damage response in 2C embryos

Our results show that peak-like exon skipping at ZGA can be in part prevented by combined overexpression of *Snrpb* and *Snrpd2* (Fig. 5d,e). Given that such exons are predicted to impact the function of proteins involved in DDR (Fig. 2c), we next evaluated the effect of *Snrpb/d2* overexpression on the ability of the 2C embryo to respond to DNA damage induced by etoposide. For this, we injected mRNA from *Snrpb*/*d2* or mCherry into pronuclear stage embryos. Injected embryos were left to develop in culture up to the 2C stage and then treated for 1.5h with a high dose of etoposide (10uM), which ensures induction of DDR at this early stage. Activation of DDR was evaluated by phospho-p53 (Ser15) immunostaining. Strikingly, while *Snrpb/d2* overexpression did not induce a significant change on phospho-p53 levels on basal conditions, these levels were significantly higher after etoposide treatment in *Snrpb/d2* injected embryos (Fig. 5f and Extended Data Fig. 10a). This suggests that the temporal skipping of exons sensitive to *Snrpb*/*d2* at least in part modulates p53-mediated DDR occurring during ZGA, contributing to the low response observed at this stage.

## DISCUSSION

We have combined multiple datasets and applied a strict quality control to generate a comprehensive and highly validated atlas of AS events during pre-implantation development in three mammalian species. Previous studies revealed that AS is very dynamic during mouse pre-implantation development, with most changes in isoform usage occurring at the ZGA ^12,23–25^. Our transcriptomic analysis confirms this observation not only for mouse but also for human and cow, despite the marked differences in the relative timing of ZGA in the three species ^7,15^. Strikingly, blastomeres undergoing ZGA in the three species showed the highest levels of isoform diversity generated by exon skipping so far reported for any other tissue, cell type or developmental stage. This is particularly surprising given the low morphological complexity of these cells, especially when compared to complex organs as intricate as the brain, which had the previously highest levels of AS ^20,39^. Remarkably, this exceptional exon skipping complexity was observed during only one or two stages, lasting 24-48 hours in development, and was due to hundreds of alternative exons whose inclusion levels displayed a sharp peak-like temporal behavior. Although the specific peak exons are not highly conserved, they overall have an equivalent molecular impact (ORF disruption at ZGA), are in genes enriched for similar gene functions (DDR), and are likely regulated by the same factors (Sm ring components) in the three studied species. Therefore, we suggest that these peak-like AS patterns are the result of a specific splicing failure at ZGA that is evolutionarily conserved and developmentally programmed to contribute to attenuate DDR during early embryogenesis.

The molecular mechanism behind this programmed splicing failure is at least in part related to the unique developmental dynamics of two Sm ring components. Unlike most adult and embryonic cells, pre-ZGA blastomeres inherit low mRNA and protein levels of *SNRPB* and *SNRPD2*, leading to skipping of sensitive exons in genes that are transcribed in the major ZGA wave. Since the *SNRPB*/*D*2 genes themselves are also strongly transcribed in this wave, their mRNA levels increase quickly, which is followed by a rise in protein levels that prompts high exon inclusion in the subsequent stages, restoring the canonical splicing patterns. Consistent with this model, zygotic injection of *Snrpb*/*d*2 mRNA led to a partial “rescue” of the splicing failure observed at ZGA. Importantly, only a subset of all exons is sensitive to low *SNRPB*/*D*2 levels, which is enriched among ZGA peak exons. These exons showed common genomic features and could be accurately discriminated from exons with matched pre-ZGA inclusion levels by a Random Forest model. Interestingly, this programmed splicing failure is reminiscent of the widespread intron detention reported during mouse spermatogenesis ^40^. In that case, an excess of transcription was proposed to overload the available spliceosomal machinery, leading to reduced splicing efficiency of a subset of introns with weak splice sites, which are then properly spliced and translated at later stages. Therefore, despite their different molecular mechanisms, associated targets and physiological consequences, these processes suggest that developmentally programmed specific splicing failures may be exploited by multicellular organisms as a strategy to regulate their development and physiology more often than previously anticipated.

The enrichment for DDR functions among genes with peak exons at ZGA in the three studied species provides a new piece to understand the special regulation of DDR during early mammalian embryogenesis. It has been shown that the DDR to irradiation increases during pre-implantation development ^31,32^, however the mechanisms behind this change in sensitivity to DNA damage are mostly unknown. We have shown that embryos treated with etoposide during ZGA fail to properly activate a p53-mediated response and neither arrest nor die until they reach morula stage. This points to the ATM pathway not being fully functional during early cleavage stages, in line with previous observations ^41,42^. Remarkably, reverting the peak-like exon skipping pattern at ZGA by *Snrpb/d2* injection led to an increase in phospho-p53 levels in response to etoposide at this stage (Fig. 5f), whereas the levels of gamma-H2AX, a mark common for the activation of different DDR pathways, increased to a similar extent upon etoposide treatment in control and *Snrpb/d2* injected embryos (Extended Data Fig. 10b,c). It is known that H2AX phosphorylation can happen independently of ATM ^41,43^; therefore, these results suggest that *Snrpb*/*d2*-dependent exon skipping occurring at ZGA has a significant impact on DDR occurring through the ATM-p53 pathway, although other pathways may not be affected by this type of regulation. In fact, *Atm* itself has an exon that gets temporarily skipped at ZGA and whose exclusion is predicted to disrupt the ORF (Extended Data Fig. 5).

Intriguingly, a delayed DDR would seemingly make early pre-implantation embryos highly sensitive to DNA damage, as it implies the accumulation of DNA lesions that will not be resolved until morula stage. In fact, DNA damage produced by DSBs at these early stages has been associated to low embryo survival ^44^. Why would this be beneficial for developing embryos? Although there could be many non-mutually exclusive explanations, we could envision a scenario in which severe DNA damage occurring during the first cell divisions is not resolved and instead gets amplified to provoke embryo arrest and death before implantation, lowering the tolerance for DNA lesions at a time when any mutation will be transmitted to all embryo lineages. However, not all DDR pathways can be lower at this developmental point, as genome stability must be ensured during ZGA, a particularly disruptive stage at the molecular level, with global transcription occurring in a largely epigenetically naive and distinct context ^45^, and with transcription itself producing DNA breaks ^46^. DNA damage is mostly sensed by the ATM and ATR kinases that activate downstream responses depending on the lesion. Hampering DDR though the ATM pathway will decrease the tolerance to particularly damaging DNA lesions such are DSBs. However, ATR responds to replication stress and is essential to coordinate replication and transcription; therefore, it is likely to be fully implemented in early embryos to be able to cope with the activation of the genome. Consistently, it has been reported that mouse embryos are particularly sensitive to the replication stress agent aphidicolin during ZGA ^47^. Moreover, ATR, but not ATM, was recently shown to be required for the conversion of mouse embryonic stem cells into 2C-like cells, which are characterized by the activation of various ZGA markers ^48^. Thus, hampering DDR through ATM but not ATR pathway could allow lowering the tolerance for DNA damage whilst ensuring genome stability during ZGA. Although other factors are likely involved, here we provide a mechanism by which a specific programmed splicing failure dependent on the levels of Sm proteins results in extensive protein disruption at the time of ZGA and contributes to a delay in the ATM-mediated response to DNA damage.

## METHODS

### RNA-Seq datasets

Publicly available Illumina RNA-Seq samples from oocyte (Oo), zygote (Zy), 2-cell (2C), 4-cell (4C), 8-cell (8C), 16-cell/morula (Mo) and blastocyst (Bl), were downloaded from NCBI Short Read Archive (SRA). All samples and associated information are provided in Supplementary Table 1. The selected datasets comprise three, four and three independent experiments with single cells or bulk embryo samples in human, mouse and cow, respectively (Fig. 1a). To ensure the selection of high quality data representative of each stage and with sufficient read coverage to measure AS, we performed the following filtering steps (the reason for exclusion of each individual sample is detailed in Supplementary Table 1): (i) Only RNA-Seq runs of at least 50 nucleotides were used; (ii) Individual cells (“outliers”) were discarded from the analyses if they did not cluster with other samples of the expected stage based on GE clustering profiles (see below; Supplementary Figs. 1-3). In particular, mouse 2C and 4C samples from ^10^ were excluded as they did not cluster with those from ^3^ likely due to slightly different timings, and cow 8C from ^49^ were discarded since they did not yet undergo ZGA; (iii) Experiments for which pooling replicates together did not provide sufficient reads for each stage were discarded (see below); (iv) Samples with strong 3′ bias, as assessed by *vast-tools align* though mapping to the five 3′-most 500-nt segments of mRNAs longer than 2,500 nts ^18^, were also removed; and (v) Other miscellaneous reasons stated in Supplementary Table 1. In total, we selected 135, 183 and 28 individual single-cell or bulk embryo samples in human, mouse and cow, respectively. Finally, to assess relative AS levels (Fig. 1e and Extended Data Fig. 2a) or PSIs in differentiated adult cell and tissue types (Fig. 2c,d), we compiled another set of samples for the three species from VastDB (http://vastdb.crg.eu/; Supplementary Table 1).

### Quantification of AS and GE levels

To calculate the percent of inclusion for a given alternative sequence (either an exon, an intron or an exon truncation/extension due to alternative 3′ or 5′ splice site choices) we used *vast-tools* v1 ^18^. *vast-tools* relies on a comprehensive database of exon-exon and exon-intron junctions for the identification and quantification of different types of AS events, and has been used to quantify inclusion levels in multiple species with high validation rates (e.g. ^18,50–55^). *vast-tools* provides a table with percent inclusion levels (using the metrics percent-spliced-in, PSI) for each AS event and sample, as well as a series of quality scores on the reliability of the estimate. In this study, we have used a minimum read coverage of VLOW for all event types (for details, see ^18^), and also filtered out intron retention events with a significant read imbalance at the two retention junctions (option --p_IR). For each species, we have used the following VASTDB libraries: human (hg19, hsa.16.02.18), mouse (mm9, mmu.16.02.18) and cow (bosTau6, bta.20.12.19). To measure GE of each gene in each cell/pool, we also used the *align* module of *vast-tools*, which provides a normalized count measure for each gene (cRPKM, corrected [for mappability] Reads Per Kilobase of transcript per Million mapped reads; see ^56^) and raw read counts for each gene.

### De-novo exon skipping events

To ensure that pre-implantation-specific exons could also be quantified by *vast-tools*, we conducted a *de novo* search for alternative cassette exons for each species using our early development RNA-Seq data (Supplementary Table 1), and created an additional *vast-tools* library with the exon-exon junctions from those exons. For this purpose, we mapped these RNA-Seq samples to their respective genomes (hg19, mm9 and bosTau6 assemblies) using *tophat2* ^57^ and built gene models through *cufflinks* ^58^. The resulting GTFs were merged using *cuffmerge* ^59^ for each species and processed using *SUPPA* ^60^ to extract all identified cassette exons. Custom scripts were then used to identify novel exons, which corresponded to internal alternative exons that were not present in *vast-tools* and had at least one upstream (C1) or downstream (C2) exon annotated in Ensembl. Using this approach, we identified 3,468 new alternative exons not present in *vast-tools* v1 for human (508 not annotated in Ensembl), 2,206 (267 not annotated) for mouse and 727 (180 not annotated) for cow (reported in Supplementary Table 2). Next, for each species, we created a library with exon-exon junctions for these exons, by joining the upstream and alternative exons (inclusion, C1A), the alternative and the downstream exons (inclusion, AC2) and the upstream and downstream exons (skipping, C1C2), with a minimum of eight mapping positions from each exon for 50-nt reads (for details, see ^61^). This additional library was incorporated into the *vast-tools* workflow as an additional module.

### Determination of cell identities, merging of samples and principal component analyses

GE values for individual cells were normalized with *DESeq* ^62^ and clustered using heatmap.2 with default parameters. Raw counts obtained from *vast-tools align* were normalized using size factors and a variance stabilizing transformation (“blind” option) before plotting the top 500 most variable genes as a heatmap with z-scores for rows (Supplementary File 1). As stated above, individual cells that did not fit into their expected stage and may represent dying, damaged or mislabelled cells were removed from further study (Supplementary Table 1). Next, since single-cell libraries have low molecular complexity and are often sequenced at low depth, resulting in low coverage across exon-exon junctions, we created pools of samples representative of specific stages, aiming at generating pools of samples with a total of >150 million reads, when possible. In all cases, cells from the same embryo were kept in the same pool, and where two embryos needed to be merged, this was based on the hierarchical clustering results. The pooling of samples was performed with the *merge* module of *vast-tools*, and the composition of the pools is described in Supplementary Table 1.

Principal Component Analysis (PCA) was conducted in R using the function *princomp* on either normalized gene expression raw counts (for GE) or PSIs (for AS). For AS, all exons with sufficient read coverage (VLOW or higher) in >80% of the merged samples and standard deviation ≥5 were considered. GE measurements took all genes with standard deviation ≥5 across merged samples as input.

### Estimation of AS complexity

We used a simple measure of the transcriptional complexity generated by AS at each developmental stage or differentiated tissue. For those AS events with sufficient read coverage (VLOW or higher) in at least 50% of all the compared samples, we calculated for each stage or tissue the fraction of AS events with sufficient read coverage in that sample whose PSI was 20<PSI<80 (i.e. was predicted to generate two prevalent isoforms). Modifying the range of PSI to define prevalent isoforms, the minimum coverage per event, the minimum fraction of samples with coverage, or using individual single cells/samples (instead of the pooled samples) did not qualitatively changed the results (Extended Data Fig. 2b,c).

### Definition of differentially spliced AS events and expressed genes at consecutive stages

Given that the number of replicates per stage and species was relatively low and uneven, we made the following definitions to call differentially spliced AS events at each developmental transition (i.e. two consecutive developmental stages) based on differences in mean PSI between stages (Extended Data Fig. 1b):

(1) Both consecutive stages must have at least two samples with sufficient read coverage (VLOW or higher) for human and mouse, or one for cow.
(2) In all cases, the overlap between the PSI distributions of the two compared stages had to be ≤ 10 (i.e. ‘range diff’ ≥ −10).
(3a) Depending on the intra-stage PSI range (i.e. the difference in PSI between the sample with the highest PSI and the one with the lowest for a given stage), a minimum mean change in PSI (ΔPSI) between the two stages was required:

3a.i) if the PSI range in both stages was ≤ 30, then |ΔPSI| ≥ 20, else
3a.ii) if the PSI range in both stages was ≤ 50, then |ΔPSI| ≥ 30, else
3a.iii) if the PSI range in any stage was > 50, then |ΔPSI| ≥ 55.
(3b) In addition, if the mean PSI of any of the two stages was ≥ 99.5 or ≤ 0.5 (i.e. either near complete inclusion or skipping), then |ΔPSI| ≥ 10.

All exons that were differentially spliced at any transition are provided in Supplementary Table 4.

For a more direct comparison with these differentially spliced definitions, we called differentially expressed genes between consecutive transitions also using qualitative definitions based on fold changes between mean cRPKM values for each stage (Extended Data Fig. 1c). First, we filtered out genes with low expression in both stages (mean cRPKM < 2). Then, for genes with mean expression cRPKM < 10 in both stages, we required that one stage had a mean cRPKM < 1 and another ≥ 5 to consider it as differentially expressed. Finally, for genes with mean expression cRPKM ≥ 10 in at least one stage, we imposed an absolute fold change ≥ 2 for a gene to be considered differentially expressed.

To assess the overlap between differentially spliced AS events and differentially expressed genes per transition (Extended Data Fig. 3a-c), we categorized each differentially spliced AS event based on whether the host gene was up- or down-regulated at the GE level, not differentially expressed or had too low expression, as defined above. To obtain the expected overlap between both types of transcriptomic change at each transition (triangles in Extended Data Fig. 3a-c), we calculated the fraction of differentially expressed genes among those hosting AS events that fulfil the minimum coverage criteria in both consecutive stages (see above). Statistical significance of the overlap was calculated through two-sided Fisher’s Exact tests using a contingency table for differentially spliced and differentially expressed genes out of the total number of genes tested in both analyses.

### Clustering AS and GE by temporal dynamics and functional enrichment analysis

We used the soft clustering algorithm Mfuzz ^28^ to group alternative exons and genes based on their PSI or expression profiles. For exons, we first selected those that had sufficient read coverage (VLOW or higher) in at least one sample for each time point, and that were differentially spliced (as described above) between any pair of stages (including non-consecutive stages). This yielded a total of 1,904, 968 and 1,083 exons for human, mouse and cow, respectively. Next, we provided a mean PSI per stage for each valid exon as input for Mfuzz. Default settings were employed, and the optimum number of clusters was automatically determined for each species using the *Dmin* function. Next, we selected those exons that were differentially spliced between any pair of stages and that had sufficient read coverage in all but one or two stages (for human and mouse) or all but one stage (for cow), and imputed the missing values in two ways: (i) if the missing point is at the beginning or end of the time course, it was assigned a value identical to the second or previous value, respectively; (ii) if the missing point is in between known values, it was assigned the mean between the two neighbouring values. These additional exons (1,050, 689 and 649 exons for human, mouse and cow, respectively) were then assigned to the previously defined Mfuzz clusters for exons with complete coverage using the ‘Mfuzz: membership’ function. It should be noted that exons in the original clusters might be reassigned to other clusters through this process. Mfuzz clusters were also generated for mean GE values in the same way that was described for alternative exons with complete coverage.

Mfuzz profiles were then classified into ‘Peak’, ‘Shift’ and ‘Other’ based on the following definitions. First, for each exon or gene, the values across the time course were classified as being in the upper, mid and lower tercile given segments of size = (Max-Mix)/3, producing an array of seven elements, one per stage (e.g. 1113111; six for cow). A profile was classified as ‘Other’ if it had either: (i) two or more values in the mid tercile, (ii) a change from upper to lower tercile in the first or last transition (or viceversa), in which case potential peak and shift patterns cannot be discriminated, (iii) the first or last value in the mid tercile; or (iv) a profile gradually changing from high to low PSI (or viceversa), defined as having the first stage in the first tercile and the last stage in the third tercile (or viceversa) and the maximum change in PSI at any consecutive stage divided by the total PSI range < 0.4; as ‘Peak’ if its first and last values were both in the first (peak-up) or third (peak-down) tercile and it was not defined as Other; and ‘Shift’ for any other profile. Profiles of each cluster were then manually inspected to ensure the accurateness of these definitions. Finally, to calculate the fraction of alternative exons that belonged to each type of cluster (Fig. 2b), only exons with coverage in all time points were considered. All exons that were included in any Mfuzz cluster as well as their associated features are provided in Supplementary Table 4.

Statistical significance of Gene Ontology (GO) term enrichments was calculated using proportion tests (*prop.test* in R) given a foreground and a background list of genes. GO annotations for the three species were downloaded from Biomart (Ensembl v91) and GO terms from each species were transferred to the orthologs in the other species to standardize the comparisons and improve cow annotations. For the GO enrichment analysis of exons in different Mfuzz clusters changing at ZGA (either Peak or Shift), we used as background all multiexonic genes that had at least one AS event (of any type) that changed in any pairwise comparison (a total of 11,096, 9,550 and 12,229 genes for human, mouse and cow, respectively).

### Assessment of evolutionary conservation

To compare each set of exons changing at a given transition in one species (Species 1) against another (Species 2), we performed the following steps to assess the evolutionary conservation at different levels, as previously described ^27,51,63^. First, to find which exons are conserved at the genomic level, we use the *liftOver* tool ^64^ with -minMatch=0.10 -multiple - minChainT=200 -minChainQ=200 parameters. Next, for those coordinates lifted to Species 2, we extracted the two neighbouring dinucleotides. Lifted exons with at least one canonical 5′ or 3′ splice site (GT/C or AG) were considered Genome-conserved. Then, we matched Genome-conserved exons to *vast-tools* v1 identifiers based on coordinate overlap and selected those that had sufficient read coverage in at least the equivalent transition of Species 2 respect to the ZGA. Exons were considered to have a “PSI change in the same direction” if they displayed a |ΔPSI| > 15 in the same direction as those in Species 1 in at least one transition. The equivalence between transitions in Species 1 and 2 was displayed using alluvial plots for each pair of species (Extended Data Fig. 4a-f). For Species 2, only the transition with the largest ΔPSI in the same direction was selected.

In addition, we performed the following steps to identify a set of orthologous exons that were regulated during pre-implantation development of the three studied mammalian species (Extended Data Fig. 3g). First, for each species, we identified differentially spliced exons between any pair of developmental stages (whether consecutive or not) as described above (Extended Data Fig. 1b). From these, we identified 93 orthologous exons that were differentially spliced in the three species. Furthermore, for those identified as differentially spliced in two species, we then asked whether there was an ortholog exon in the third species and, if so, whether it had an average |ΔPSI| > 15 between any pair of stages. This resulted in a total of 259 orthologous exons (Supplementary Table 3). To assess the enrichment of GO terms among the genes hosting these exons, we used as background multiexonic genes with 1:1:1 orthologs in the three species and with at least one exon skipping event with coverage in two stages in any species (15,132 genes).

Finally, to assess regulatory conservation of genes differentially expressed at each transition in a given species (Species 1) against another (Species 2) (Supplementary Fig. 4), we first identified one-to-one orthologs based on Ensembl-BioMart information, and then checked whether any differentially up- or downregulated gene in Species 1 was also identified as up- or down-regulated in Species 2 (“GE change same direction”). The equivalence between transitions was shown using alluvial plots, in which only the transition with the largest fold change in the same direction was selected for Species 2.

### Predicted impact on open reading frames, NMD and ribosome-engagement analyses

The predicted impact on the ORF of the inclusion/exclusion of each alternatively spliced sequence was obtained from *VastDB* (release 3) and it was inferred essentially as described in ^26^. Four major categories are reported: (i) AS events that are predicted to generate alternative protein isoforms both upon inclusion and skipping of the alternative sequence (i.e. are located in the coding sequence [CDS] and maintain the ORF and/or are located towards the end of the CDS and are not predicted to trigger NMD or to create a large protein truncation [>20% of the reference isoform and/or >300 aa]); (ii) AS events that disrupt the ORF upon sequence inclusion (e.g. most introns, exons that are usually not included and whose length is not multiple of three and/or contain in-frame stop codons); (iii) AS events that disrupt the ORF upon sequence exclusion (e.g. exons that are normally constitutive and whose length is not multiple of three); (iv) AS events in the 5′ or 3′ untranslated regions (UTRs). Comparisons among ZGA Mfuzz clusters (Fig. 2C,D) included only those exons that were labelled as alternative protein isoform (i), disruptive upon inclusion (ii) or disruptive upon exclusion (iii).

To compare the relative engagement of the inclusion and exclusion isoforms on ribosomes for different exon subsets (Extended Data Fig. 6d,e,g,h), we employed Transcript Isoforms in Polysomes sequencing (TrIP-seq) data for human embryonic stem cells ^29^ or HEK 293T cells ^30^. We used *vast-tools* to obtain PSI values for each exon in the cytosolic and high-polysome fractions and calculated the ΔPSI between the two fractions, which gave a measure of the difference in ribosomal engagement (positive ΔPSI means higher engagement upon inclusion, and the opposite for negative ΔPSI and exclusion). For each exon category, we plotted events with sufficient read coverage (VLOW or higher) in both the cytosolic and high-polysome fractions. For differentially spliced exons at the ZGA transition (Extended Data Fig. 6d,e), we plotted the ΔPSI with respect to the ZGA isoform grouped by the predicted impact on the ORF at ZGA (i.e. positive/negative ΔPSI values imply the ZGA isoform is more/less engaged). For each type of temporal dynamics with change at ZGA (Extended Data Fig. 6g,h), we plotted directly the ΔPSI between cytosolic and high-polysome fractions (i.e. a negative/positive ΔPSI implies that the inclusion/skipping isoform, that is the ZGA isoform for peak-up/peak-down exons, respectively, is less engaged). In addition, to assess the impact of NMD disruption on these sets of exons (Extended Data Fig. 6c,f), we performed an equivalent analysis calculating the ΔPSI upon UPF1 knockdown in HR1 cells for each category (data from ^65^).

Finally, to study the enrichment or depletion of transposable elements in different Mfuzz exon clusters (Extended Data Fig. 7), we overlapped the coordinates of these elements for each species as defined by RepeatMasker (excluding simple repeats) and of the alternative exons and neighbouring intronic regions (500 bp upstream and downstream the exon) using *bedtools intersect*, and counted the number of exons with at least 1-nt overlap with any transposable element family. For each Mfuzz cluster, we plotted the relative fraction of exons overlapping transposable elements with respect to the fraction of all exons included in Mfuzz clusters (“ALL”).

### Evaluation of intron-exon features of ZGA-Peak exons

We used *Matt* v1.3.0 ^36^ to compare exon and intron features associated with splicing regulation between exons with Peak dynamics at ZGA and different reference exon sets. For each group of exons being compared, *Matt cmpr_exons* automatically extracts and compares 69 genomic features associated with AS regulation, including exon and intron length and GC content, splice site strength, branch point number, strength and distance to the 3′ splice site using different predictions, length and position of the polypyrimidine tract, etc. For the calculation of splice site and branch point strength, we used the available human models. Comparisons among groups are performed using Mann-Whitney U tests and visualized using boxplots (Supplementary File 1). For this analysis, we defined the following six exon sets for each species (Supplementary Table 8):

i. P_Dw: exons with a peak-down profile at ZGA, sufficient read coverage (VLOW or higher) in the pre-ZGA and ZGA stage, and a ΔPSI ≤ −10 at the ZGA transition. Given the broader time of ZGA in cow, the ZGA ΔPSI was defined as the largest ΔPSI from either 4C-8C or 4C-Morula (also in (ii)). N of exons: Human = 526, mouse = 345, cow = 646.
ii. P_Up: exons with a peak-up profile at ZGA, sufficient read coverage in the pre-ZGA and ZGA stage, and a ΔPSI ≥ 10 at the ZGA transition. N of exons: Human = 236, mouse = 32, cow = 152.
iii. HIGH_PSI: exons with PSI > 90 across both pre-implantation and differentiated samples with sufficient read coverage. N of exons: Human = 11878, mouse = 6768, cow = 5108.
iv. LOW_PSI: exons with PSI < 10 across both pre-implantation and differentiated samples with sufficient read coverage. N of exons: Human = 4077, mouse = 1213, cow = 955.

In addition, we extracted 33710, 16999 and 9228 background exons with sufficient read coverage in the pre-ZGA and ZGA stage, |ΔPSI| < 5 at the ZGA transition, and that are not part of the foreground sets (i) and (ii) for human, mouse and cow, respectively. Since the pre-ZGA PSI distribution of these background exons differ from those of (i) and (ii), we constructed two stratified background sets. For this purpose, for each exon of each foreground set (i and ii), we randomly selected one background exon whose pre-ZGA PSI deviates from the pre-ZGA PSI of the foreground exon by no more than 5. Selected background exons were then excluded from the pool of background exons and the process was repeated a total of four times. The stratified background exon sets after the four iterations ([v] Bg_Dwstrat and [vi] Bg_Upstrat) contained 2104/1224/2584 and 944/128/604 for sets (i) and (ii) for human/mouse/cow, and matched at least 89% of the foreground exons.

#### Classification and feature analysis with Random Forest

To consolidate and extend results obtained by *Matt*’s discriminative feature analysis, we applied a Random Forest model to the classification of peak-down exons versus a matched background and peak-up versus a matched background for each of the three species, and extracted the variable importance with the goal to identify features most relevant for these classifications. First, we constructed a comprehensive set of 746 features for the exons as follows: a) All 69 intron-exon related genomic features extracted with function *get_efeatures* of splicing toolkit *Matt* v1.3.0, as mentioned above; b) GE fold change at ZGA (ZGA/pre-ZGA) of the gene the exon belongs to; c) For each 338 regular-expression RNA-binding motifs from CisBP-RNA v0.6, we scanned the 200 nt upstream and 200 nt downstream (150 nt intronic + 50 exonic) sequences for each exon for motif hits and added the number of hits as individual features.

We then employed the R package *randomForest* v4.6-12 to train models for each foreground exon set. Because the foreground and background sets differ in size by at least one order of magnitude, we down-sampled the background exon set anew for each Random Forest and trained multiple models in an iterative manner. In each iteration, we randomly chose for each foreground exon one background exon with |ΔPSI| < 5 at ZGA, ensuring that the characteristics of the foreground and background exons have similar weight when training Random Forests. We held fixed the Random Forest parameters ntry=25 and nodesize=1, whose recommended values for classification tasks are sqrt(N features) and 1. We optimized the number of trees (ntree) by iterating it from 100 to 3100 by 300. For each ntree value we trained 100 Random Forest and determined average classification accuracies measures by comparing the OOB classification votes to the ground truth. We chose the ntree value that gave overall best average OOB error, area under ROC and precision recall curves, average sensitivities for specificities 90% and 95%. At the same time, we chose ntree as small as possible to guard against overfitting effects. Finally, we trained 1000 Random Forests with the optimal ntree value and with different down-sampled stratified background sets in each run, and reported average ROC and precision recall curves, as well rankings of the features according to their average feature importance (i.e. mean decrease of accuracy as reported by the Random Forest model).

### Identification of potential regulators of exons with Peak dynamics

To identify potential regulators of exons with peak-up and peak-down Mfuzz dynamics at ZGA in the three species, we took three main approaches:

- <u>Sequence motif enrichment analyses</u>: from the foreground sets (i, P_Dw) and (ii, P_Up) described above, we extracted 300 nt of the upstream and of the downstream intronic regions as well as the exon sequences and employed *Matt* v1.3 ^36^ to test enrichment of all available motifs for RNA binding proteins (RBPs) in CisBP-RNA ^37^ in each of three regions separately. The background set corresponded to those exons that have sufficient read coverage in all or all but one stage and that were not part of P_Dw or P_Up sets (human: 34452 exons, mouse: 20232 exons, cow: 11177 exons).
- <u>Correlation between RBP expression and exon PSIs at the single cell level</u>: For each foreground exon set in human and in mouse, we correlated (Pearson) at the single-cell level the GE values of 196 manually curated splicing factors (Supplementary Table 6) with the mean PSIs of those peak-down or peak-up exons (i and ii sets above) with sufficient read coverage in each blastomere of the ZGA stage (8C in human and 2C in mouse). This was performed using *corr.test* from the *psych* package in R.
- <u>Overlap with specific splicing factor-dependent exons</u>: To identify peak exons whose inclusion levels were dependent on specific splicing factors, we first collected publicly available RNA-Seq data from specific splicing factor depletion experiments (knockdown or knockout) in any cell or tissue type (Supplementary Table 1). In total, we compiled data for 119 available experimental perturbations for 84 splicing factors comprising 64 independent studies. These data were processed using *vast-tools* to obtain ΔPSIs for each experiment and splicing factor for all exons with sufficient read coverage in control and experimental condition. Next, for each individual splicing factor and experiment, we scored the number of ZGA peak exons (i and ii sets above) that show a |ΔPSI| ≥ 15 upon splicing factor depletion. To identify the most promising splicing factors (Fig. 4d), we calculated: (i) the percent of exons with sufficient read coverage affected by the depletion, and (ii) the consistency of these changes (e.g. downregulation of peak-down and upregulation of peak-up), as estimated by the percentage of exons in quadrants Q1 and Q3 and evaluated using a two-sided Binomial test (Supplementary Table 7).

### Embryo collection and manipulation

All embryos were obtained from B6CBA F1 crosses. Zygotes were collected from the oviduct of superovulated females 20h post HCG injection and cultured in EmbryoMax® KSOM Mouse Embryo Media (Millipore) at 37°C, 5%CO2 up to 2 cell, 4 cell, 8 cell, morula or blastocyst stage. For the experiments in Fig. 3f,g early morulas were collected at 2.5dpc by flushing of the oviduct in M2 media (Sigma). For the overexpression experiments (Fig. 5d-f), *Snrpb* and *Snrpd2* cDNA were cloned into a modified pCS2+8NmCherry vector lacking mCherry tag (Addgene) for their *in vitro* transcription. The mCherry used as control was transcribed from the pCS2+8NmCherry vector (Addgene). *In vitro* transcription was performed using the mMESSAGE mMACHINE® SP6 Transcription Kit (Ambion) according to manufacturers’ instructions. For all overexpression experiments 1 cell embryos at pronuclear stage were microinjected with 300ng/ul of mCherry mRNA (control) or 150ng/ul of *Snrpb* mRNA and 150ng/ul *Snrpd2* mRNA (*Snrpb/d2*).

### Etoposide treatment

For experiments in Fig. 3, embryos were either left untreated or treated for 1h at the stated stage with 0.5uM etoposide (Sigma). Following treatment embryos were either fixed directly for immunostaining or washed in KSOM media and kept in culture to be fixed at the desired stage. For *Snrpb* and *Snrpd2* overexpression experiments (Fig. 5f), injected one-cell embryos were left in culture and either treated with 10uM etoposide for 1.5h or left untreated. Following the treatment embryos were fixed for immunostaining.

### RT-PCR validation assays and RNA-Seq experiments

RT-PCR assays for alternative exon validations were performed on pools of embryos at different developmental stages. The Arcturus Pico Pure RNA extraction kit (Thermo Fisher) was used for RNA extraction and cDNA was transcribed with Superscript III Reverse Transcriptase (Thermo Fisher). RT-PCR primers can be found in Supplementary Table 9. PSI quantification from RT-PCR was performed using the Fiji software ^66^.

For RNA-Seq experiments following *Snrpb* and *Snrpd2* overexpression, one cell embryos were microinjected as described above and collected for RNA extraction at either 5h post injection (zygote stage) or 24h post injection (2C stage). For each condition 40 embryos coming from three independent experiments were pooled to extract RNA for sequencing. RNA was extracted using the Qiagen RNeasy Micro Kit. SMARTer Stranded RNA-Seq Kit was used for library preparation prior to Illumina sequencing. Libraries were sequenced in a HiSeq2500 machine, generating an average of ~69 million 125-nt paired-end reads per sample. Read numbers and mapping statistics are provided in Supplementary Table 10.

### Snrpb *and* Snrpd2 *overexpression RNA-Seq analysis*

RNA-Seq data for control or *Snrpb* and/or *Snrpd2* overexpressing embryos at 1C or 2C stages was processed using *vast-tools*. For both AS and GE analyses, control 1C and 2C embryos were compared to identify differentially spliced exons or expressed genes at ZGA, and 2C embryos overexpressing mRNA from *Snrpb*, *Snrpd2* or both genes were compared with the control condition. For AS analyses (Fig. 5c,d and Extended Data Fig. 9a,b), only exons with sufficient read coverage in 1C and 2C controls as well as the tested experimental condition were used, and a cut-off of |ΔPSI| ≥ 15 was employed. For GE analyses, cRPKM values were normalized using quantile normalization in R, and genes with fewer than 50 read counts or expression levels lower than cRPKM < 5 in the three conditions (1C and 2C controls as well as the tested experimental condition) were discarded. To calculate log2 fold changes, 0.01 was added to each normalized cRPKM value, and a minimum fold change of |FC| ≥ 2 was set as a cut-off.

### Embryo immunostaining, TUNEL staining and western blot

Embryos were fixed in 4%PFA/PBS for 10 min. Following fixation they were permeabilised in 0.5% Triton X-100 for 15min, blocked in 10%BSA/0,1% Triton X-100/PBS and incubated overnight in primary antibody: anti-SNRPB (Thermo Fisher), anti-SNRPD2 (Thermo Fisher), anti-phospho H2AX Ser 139 (Cell Signaling), anti-phospho p53 Ser15 (Cell Signaling). Hoechst was used for nuclear staining and CytoPainter Phalloidin-iFluor 647 Reagent (Abcam) for membrane staining. Imaging was conducted in a Leica SP5 inverted confocal and images processed with Fiji software ^66^. For quantification, relative intensity represents the mean fluorescent intensity of the nucleus relative to nucleus area along the biggest nuclear section, measured with Fiji. Each dot represents average relative intensity of all cells of the embryo. Each plot represents quantification of embryos from six (Fig. 3b,c) or three (Fig. 3-g and 5f, and Extended Data Fig. 10) independent experiments.

The Apop-Tag Fluorescein Kit (Millipore) was used for TUNEL staining according to manufacturer’s instruction with minor modifications. Briefly, embryos were fixed in 4% paraformaldehyde for 10 minutes at room temperature, washed in 0.1% Triton X-100/PBS, permeabilised for 15 minutes in 0.4% Triton X-100/PBS. Following washes in PBS embryos were equilibrated and stained according to the Apop-Tag kit’s protocol. Hoechst and Cytopainter Phalloidin-iFluor 647 Reagent (Abcam) were used for counterstaining following TUNEL staining. For western blot embryos at different developmental stages were collected as described above. For protein extraction pools of embryos at each stage were collected in Laemmli Buffer. Proteins were separated by 14% SDS-PAGE and transferred to PVDF membranes (Biorad). Following blocking in 5% Milk (Sigma) in TBST, membranes were incubated overnight with the primary antibody in 5% BSA (Sigma) in TBST. The following antibodies and dilutions were used: anti-SNRPB (Thermo Fisher, 1/200), anti SNRPB2 (Abcam, 1/2000) and anti-GAPDH (Abcam, 1/10000).

## Acknowledgements

We thank Jernej Ule and Rupert Faraway for feedback on the analyses, and Federica Mantica for assistance on R plotting. The research has been funded by the Spanish Ministerio de Ciencia (BFU2014-55076-P and BFU2017-89201-P to MI), Marie Skłodowska-Curie actions grants (H2020-MSCA-IF-2014_ST-656843 to BP), La Caixa PhD fellowship to CDRW, and the ‘Centro de Excelencia Severo Ochoa 2013-2017’ (SEV-2012-0208). We acknowledge the support of the CERCA Programme/Generalitat de Catalunya and of the Spanish Ministry of Economy, Industry and Competitiveness (MEIC) to the EMBL partnership.

