## Supplementary Materials for "A developmentally programmed splicing failure attenuates the DNA damage response during mammalian zygotic genome activation"

**Supplementary Table Legends**

**Supplementary Table 1 - RNA-Seq samples used in this study.** Several spreadsheets containing the publicly available RNA-Seq datasets used in this study for human, mouse and cow. For each species, we provide a sheet containing the early development data and another the cell and tissue types. Each table includes the SRA identifier, number of reads, length, source, etc. Furthermore, the groups of samples that were pooled together to increase read depth are indicated. For discarded samples (red font), the reason for removal is indicated. An additional sheet ("RBP_experiments_Human") contains the information about the experiments of splicing factor depletion, an another one ("Pool_read_stats") contains the summary statistics of the sample pooling.

**Supplementary Table 2 - Alternative exons not included in vast-tools.** For each species, the list of alternative exons that were not present in vast-tools and that were added as an additional module. For each exon, the basic information is provided, including coordinates (hg19, mm9 or bosTau6), whether or not they are annotated in Ensembl ("Non_annot"), whether they are part of any Mfuzz cluster and the type of cluster ("ClusterID" and "ClusterType"), and the average PSI in each developmental stage (NA, insufficient read coverage).

**Supplementary Table 3 - Alternative exons dynamically regulated during early embryogenesis of the three species.** Exons that showed dynamic inclusion levels in the three studied species (Extended Data Fig. 4g). Each row corresponds to an orthologous exon and the information is provided for the three species, including coordinates (hg19, mm9 and bosTau6), vast-tools ID, PSI in each developmental stage and species and predicted impact on the ORF.

**Supplementary Table 4 - Differentially regulated exons in early development.** For each species, the list of exons that showed differential regulation in any transition and/or that were included in the Mfuzz clusters. In addition to the basic exon information, the table includes: summary statistics of the inclusion levels across differentiated tissues and early development samples, conservation status in the other two species and predicted protein impact.

**Supplementary Table 5 - Significantly enriched motifs for RNA binding proteins.** For each genomic region (upstream intron, exon or downstream intron), the motifs from cisBP-RNA that are significantly enriched over the background (as estimated by Matt) for peak-down (P_Dw) and peak-up (P_Up) exons at ZGA are shown for each species. Color code: dark red, the motif is depleted in P_Dw exons; light red, the motif is enriched in P_Dw exons; dark blue, the motif is depleted in P_Up exons; light blue, the motif is enriched in P_Up exons.

**Supplementary Table 6 - Correlations between GE of splicing factors and PSIs at the single-cell level.** Summary statistics (correlation coefficient and p-value) for the correlation analysis between the expression of 196 splicing factors and the inclusion levels of peak-down and peak-up exons at ZGA in individual ZGA blastomeres (8C in human and 2C in mouse, respectively).

**Supplementary Table 7 - Association between splicing factor depletion and ZGA.** Summary statistics for the overlap between peak-down and peak-up exons at ZGA and exons changing ($|\Delta PSI| > 15$) upon depletion of specific splicing factors. For each experiment (X*), the number of exons in quadrants Q1-Q4 (based on the direction of the overlap) is provided, as well as the percentage of those in Q1 and Q3 ("Perc_Q1_Q3") and the p-value of a two-sided Binomial associated this percentage. Moreover, for the total number of peak exons at ZGA with read coverage ("Cov_P_Up" and "Cov_P_Dw"), it provides the percentage of exons in each quadrant.

**Supplementary Table 8 - Exons used for genomic feature analyses.** For each species, the list of genes used as input for *Matt cmpr_exons* and associated information (including PSI in the pre-ZGA and post-ZGA stages and the group to which they belong).

**Supplementary Table 9 - Primers used for RT-PCR validations.**

**Supplementary Table 10 - Statistics for in-house RNA-Seq samples.** Statistics associated with the RNA-Seq data for embryos injected with *Snrpb* and/or *Snrpd2* or a control.

**Supplementary File 1 - Output files from *Matt cmpr_exons* analyses for human, mouse and cow exon sets.** Full PDF reports obtained from *Matt* for each species. The reports are concatenated, and the species investigated is stated in the fifth page of each report (pages 5, 112 and 205; highlighted in yellow).

### Extended Data Figures

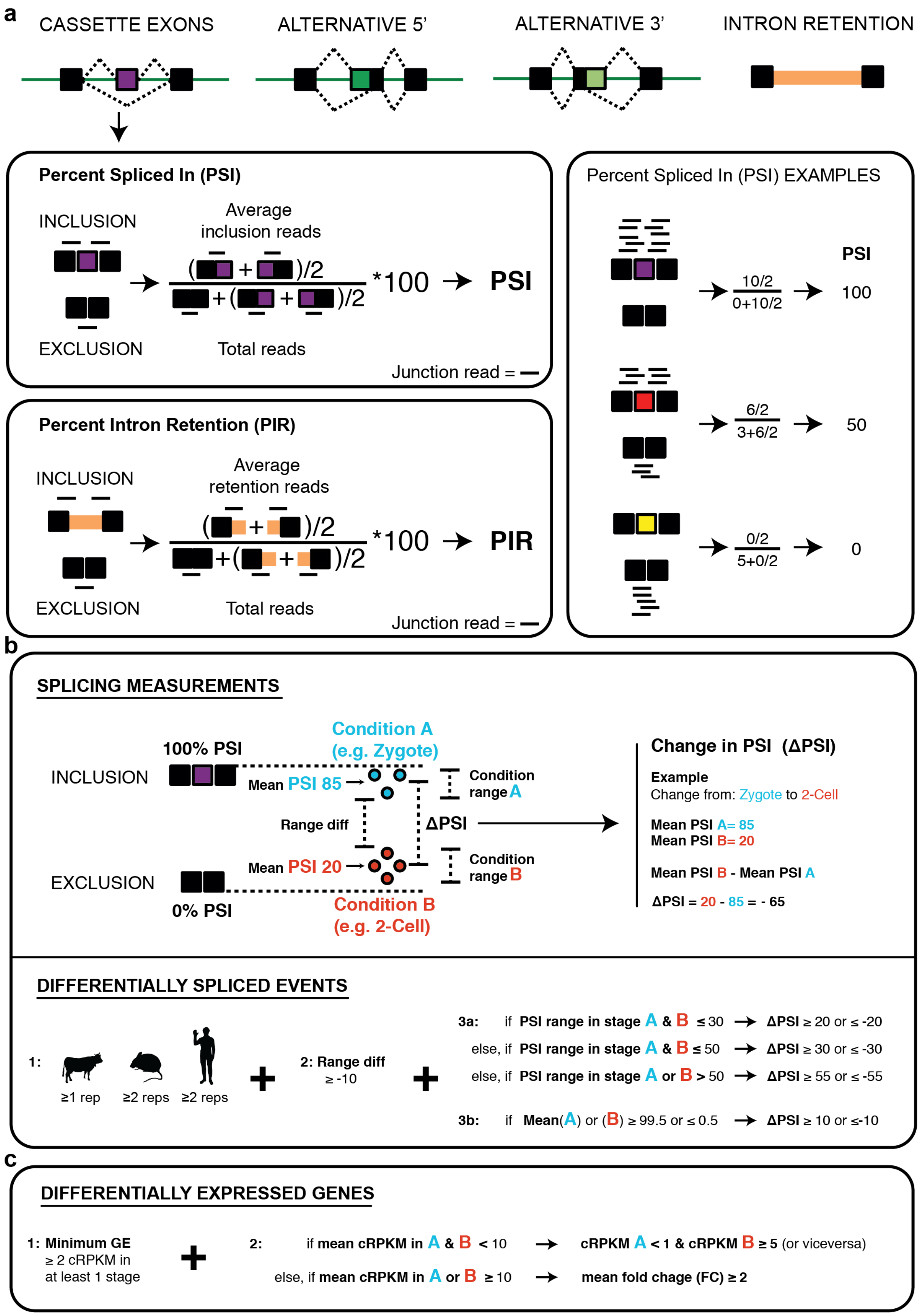

**Extended Data Fig. 1 - Schematic summary of AS quantification and differential AS and GE analyses.** **(a)** Simplified schematic representation of how Percent Spliced In (PSI) and Percent Intron Retention (PIR) are calculated in *vast-tools* using solely RNA-Seq reads mapping to exon-exon (or exon-intron) junctions. See [18,68] for further details. **(b)** Top: schematic representation of the main descriptive statistics considered for each stage (mean PSI and range of PSI of samples with sufficient read coverage) and the parameters used to call differential AS between two conditions (change in mean PSI between the two conditions [ΔPSI] and difference between the minimum and the maximum PSI in the two conditions [Range diff]). Bottom: definition of differentially spliced events between two stages. 1: minimum number of required samples with sufficient read coverage per stage, depending on the species (human: 2, mouse: 2, cow: 1). 2: Range diff $\geq$ -10. Either of two conditions, 3a: a minimum mean |ΔPSI| depending on the PSI range in each stage, or 3b: if mean PSI in either stage is very high (PSI $\geq$ 99.5) or very low (PSI $\leq$ 0.5), then a |ΔPSI| $\geq$ 10 is required. **(c)** Schematic representation of the definition of differentially expressed genes between two stages based on mean GE levels at each stage and mean fold change in GE.

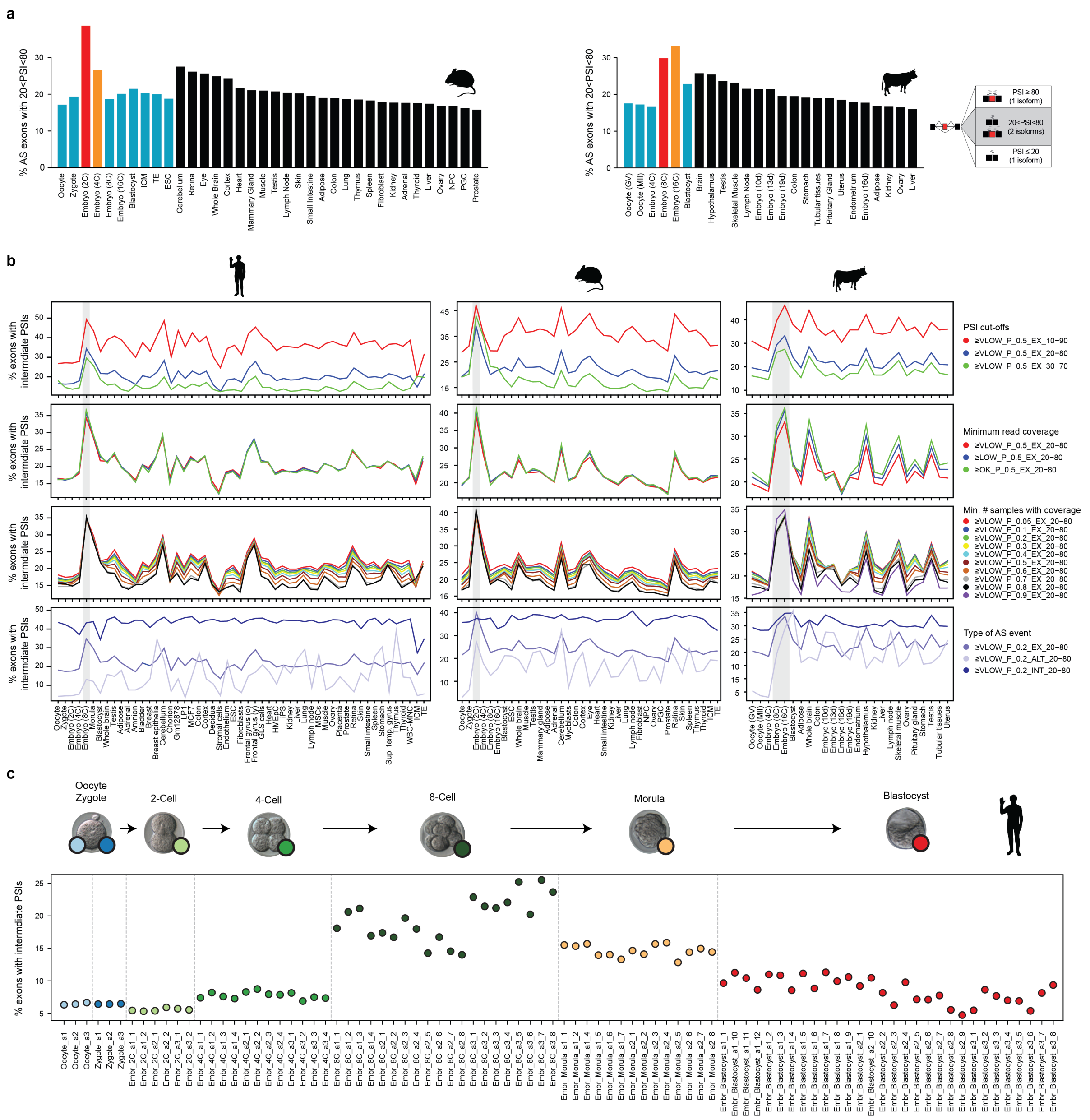

**Extended Data Fig. 2 - Relative exon skipping complexity is maximal at ZGA. (a)** Percentage of exons with 20<PSI<80 (i.e. generating two substantial isoforms) in each sample was calculated for every stage and differentiated tissue, showing the highest relative exon skipping levels for the ZGA stage for mouse (left) and cow (right). **(b)** Percentage of AS events with intermediate PSI levels for each species assessing the impact of different variables and cut-offs. From top to bottom: (i) different PSI cut-offs: 10-90, 20-80 (as in (A)) or 30-70. (ii) Minimum read coverage, as defined by *vast-tools* quality scores (≥VLOW, ≥LOW or ≥ OK). (iii) Minimum number of samples with coverage across the panel, from a minimum fraction of 0.05 to 0.9. (iv) Type of AS event, including exons (EX), introns (INT) or alternative 3′ and 5′ splice sites (ALT). **(c)** Percentage of exons with 20<PSI<80 in individual blastomeres.

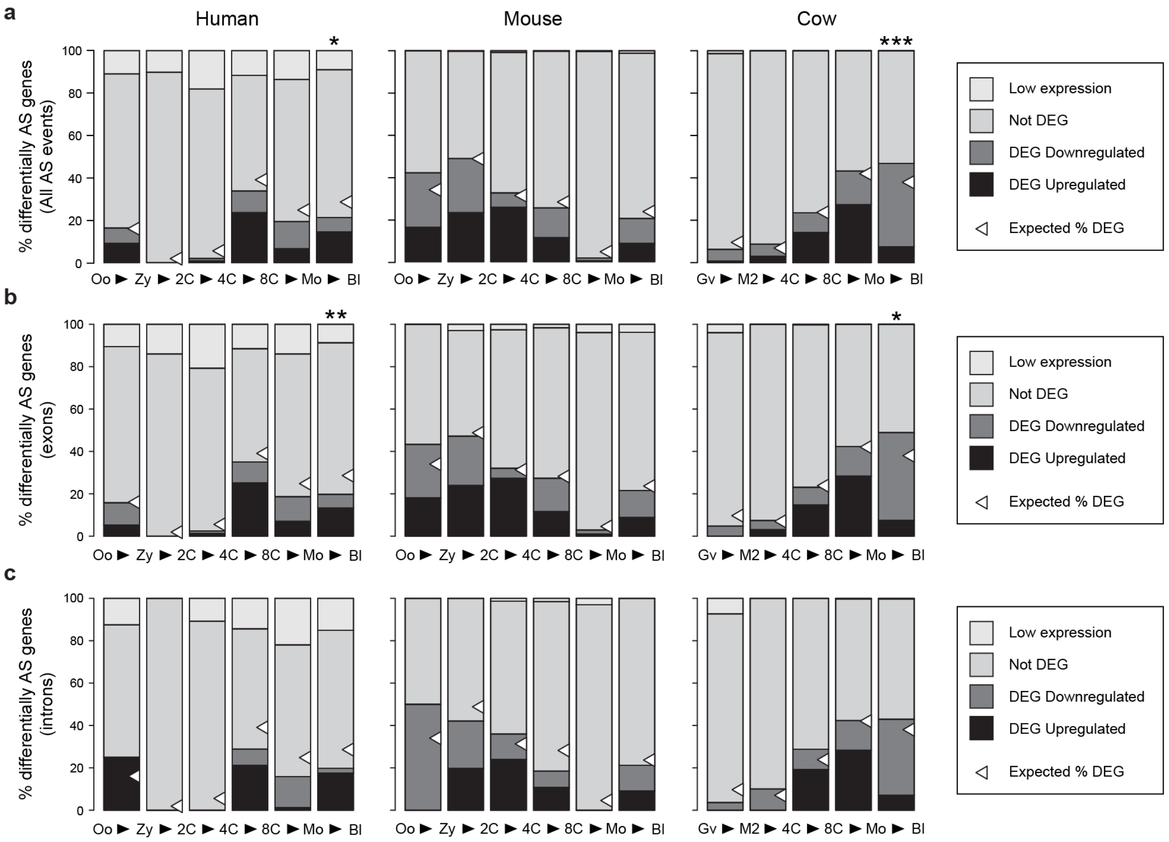


**Extended Data Fig. 3 - Overlap between genes regulated at the GE and AS levels. (a-c)** Percentage of genes with differentially spliced AS events (a), exons (b) or introns (c) at each transition with differential GE in the same transition. White triangles correspond to the percentage of differentially expressed genes (DEGs) expected by chance. * (0.01 ≤ $P$ < 0.05), ** (0.001 ≤ $P$ < 0.01) and *** ($P$ < 0.001) indicate statistical significance (observed vs. expected) based on Proportion tests. "Low expression": genes with cRPKM < 2 in both stages of the transition and not tested for differential expression.

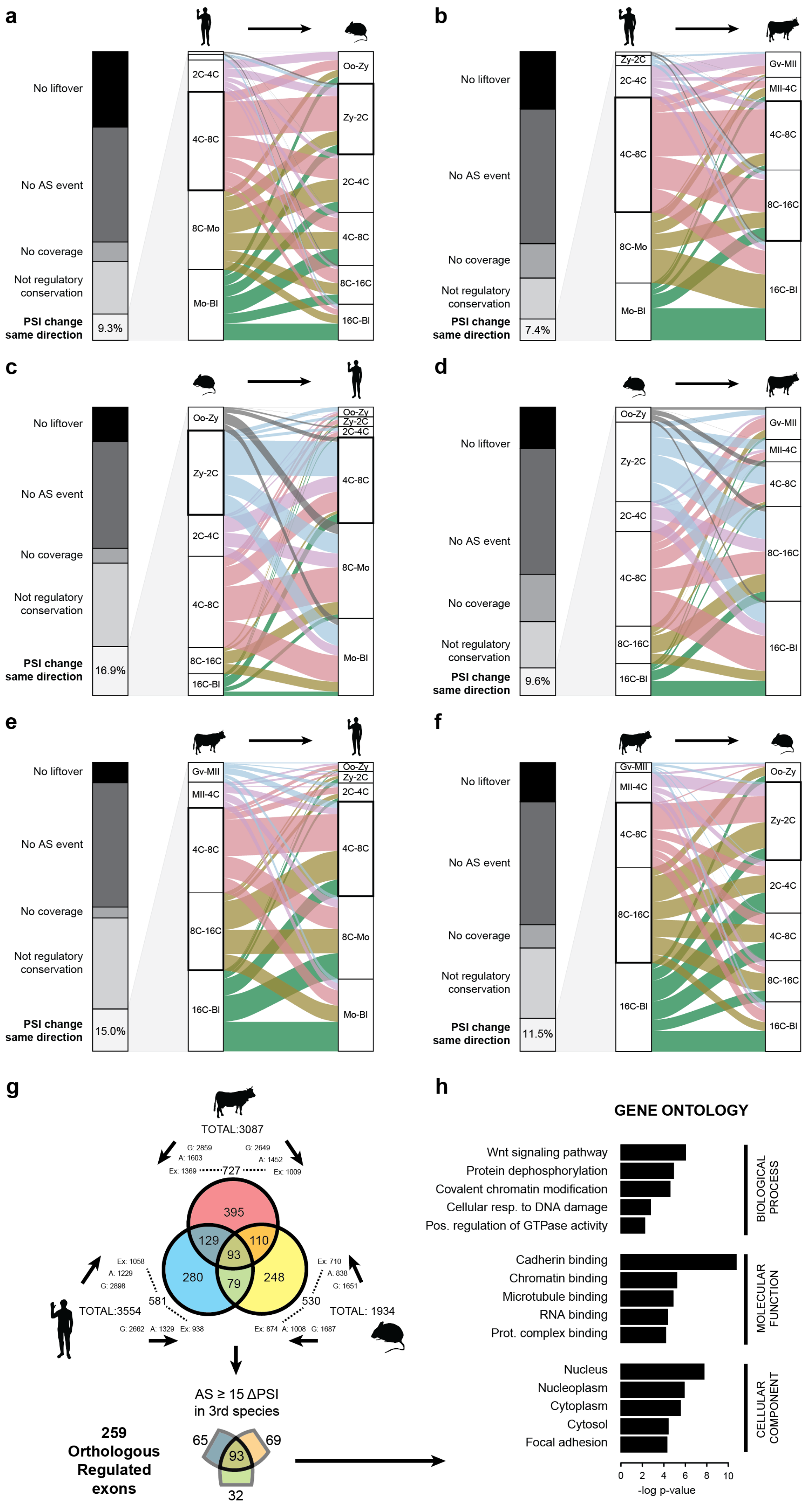
a
No liftover
No AS event
No coverage
Not regulatory conservation
PSI change same direction
9.3%
2C-4C
4C-8C
8C-Mo
Mo-Bl
Oo-Zy
Zy-2C
2C-4C
4C-8C
8C-16C
16C-Bl
b
No liftover
No AS event
No coverage
Not regulatory conservation
PSI change same direction
7.4%
Zy-2C
2C-4C
4C-8C
8C-Mo
Mo-Bl
Gv-MII
MII-4C
4C-8C
8C-16C
16C-Bl
c
No liftover
No AS event
No coverage
Not regulatory conservation
PSI change same direction
16.9%
Oo-Zy
Zy-2C
2C-4C
4C-8C
8C-16C
16C-Bl
Oo-Zy
Zy-2C
2C-4C
4C-8C
8C-Mo
Mo-Bl
d
No liftover
No AS event
No coverage
Not regulatory conservation
PSI change same direction
9.6%
Oo-Zy
Zy-2C
2C-4C
4C-8C
8C-16C
16C-Bl
Gv-MII
MII-4C
4C-8C
8C-16C
16C-Bl
e
No liftover
No AS event
No coverage
Not regulatory conservation
PSI change same direction
15.0%
Gv-MII
MII-4C
4C-8C
8C-16C
16C-Bl
Oo-Zy
Zy-2C
2C-4C
4C-8C
8C-Mo
Mo-Bl
f
No liftover
No AS event
No coverage
Not regulatory conservation
PSI change same direction
11.5%
Gv-MII
MII-4C
4C-8C
8C-16C
16C-Bl
Oo-Zy
Zy-2C
2C-4C
4C-8C
8C-16C
16C-Bl
g
TOTAL:3087
G: 2859
A: 1603
Ex: 1369
727
G: 2649
A: 1452
Ex: 1009
395
129
110
93
280
79
248
Ex: 1058
A: 1229
G: 2898
581
Ex: 710
A: 838
G: 1651
530
TOTAL:3554
TOTAL: 1934
G: 2662
A: 1329
Ex: 938
Ex: 874
A: 1008
G: 1687
AS ≥ 15 ΔPSI
in 3rd species
259
Orthologous
Regulated
exons
65
69
93
32
h
GENE ONTOLOGY
Wnt signaling pathway
Protein dephosphorylation
Covalent chromatin modification
Cellular resp. to DNA damage
Pos. regulation of GTPase activity
BIOLOGICAL PROCESS
Cadherin binding
Chromatin binding
Microtubule binding
RNA binding
Prot. complex binding
MOLECULAR FUNCTION
Nucleus
Nucleoplasm
Cytoplasm
Cytosol
Focal adhesion
CELLULAR COMPONENT
0
2
4
6
8
10
-log p-value

**Extended Data Fig. 4 - Evolutionary conservation analysis of alternative exon and GE changes. (a-f)** For each query and target species, stack plot of the level of conservation for each alternative exon changing at each transition in the query species. No liftover: absence of genomic conservation; No AS event: although the exon is found in the genome, no alternative exon is present in the *vast-tools* v1 annotation (i.e. likely a constitutive exon); No coverage: the exon does not have sufficient read coverage in at least the equivalent stages in the target species; No regulatory conservation: the exon does not change in the same direction ($|\Delta PSI| \geq 15$) at any of the transitions in the target species. From the percentage of exons that change in the same direction in at least one transition of the target species, alluvial plot showing the transition of change in query species (left) and the observed highest change in the same direction in the target species (right). ZGA transition in each species is highlighted by a thicker block. (a) Human to mouse, (b) human to cow, (c) mouse to human, (d) mouse to cow, (e) cow to human, (f) cow to mouse. **(g)** For all exons differentially regulated in any pairwise stage comparison for each species (TOTAL), number of them with genome conservation [G], *vast-tools* AS identifier [A] and sufficient read coverage [Ex] in the target species (arrows). The intersect between Ex for both target species is shown at the edge of each circle of the Venn diagram (human: 581, mouse: 530, cow: 727). For these exons, the Venn diagram shows the overlap of exons that are differentially spliced in at least one pairwise stage comparison. For those exons regulated in two species, we then asked (bottom) whether the orthologous exon had a $|\Delta PSI| \geq 15$ in any pairwise stage comparison in the third species. This yielded a total of 259 orthologous exons with dynamic regulation in the three species. **(h)** GO term enrichment analysis for the genes harboring these 259 exons in human. Abbreviations: Oo, oocyte; Zy, zygote; Mo, morula; Bl, blastocyst; Gv, oocyte GV; MII, oocyte MII.

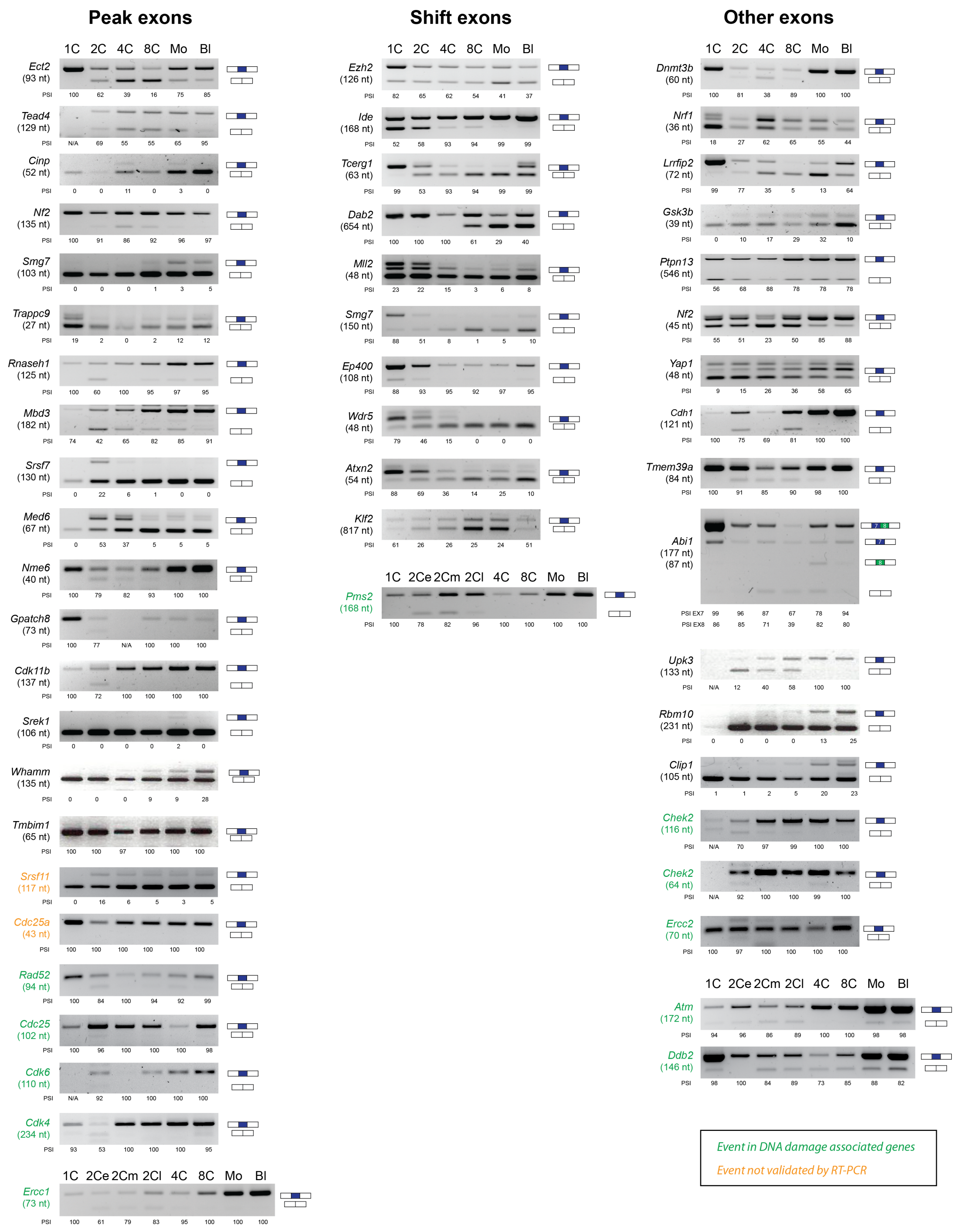


**Extended Data Fig. 5 - RT-PCR validation of exons during early development.** RT-PCR assays were performed using independent RNA samples for each developmental stage to validate different types of exon profiles. Exon length in nts is indicated, as well as PSI quantifications based on relative band intensity (as measured with ImageJ). Exons in DDR genes are highlighted in green, and the two exons whose profiles were not validated, in orange. Note that variable amounts of cDNA template were used for different stages and PCR reactions; therefore changes in GE cannot be inferred from the variations in band intensity between stages.

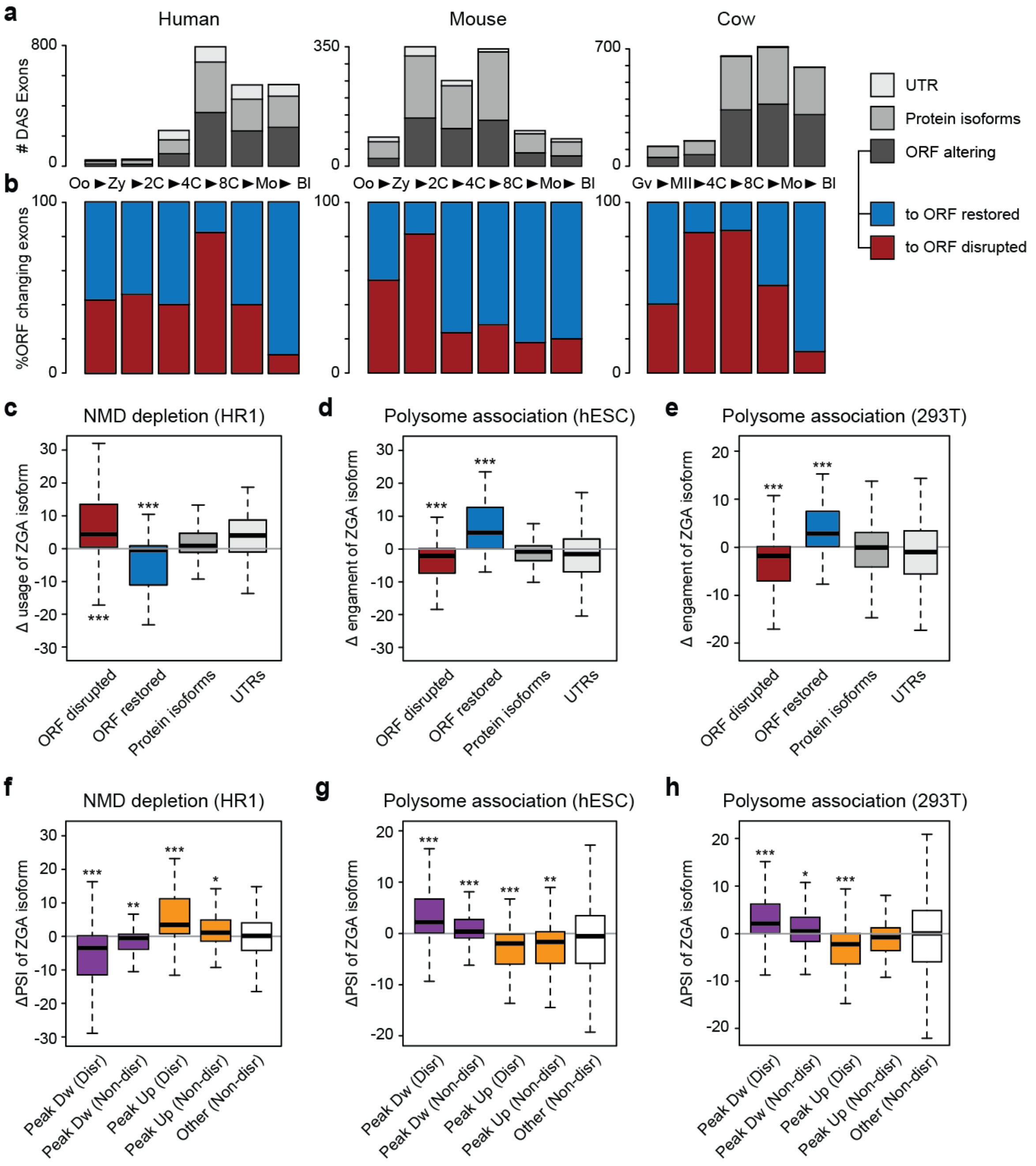


**Extended Data Fig. 6 - Differentially spliced exons at ZGA often generate non-functional proteins. (a)** Number of differentially spliced exons at each transition by predicted impact on the ORF. "UTR", exon located in the 5′ or 3′ UTR; "Protein isoforms", both inclusion and exclusion isoforms are predicted to lead to functional protein isoforms; "ORF altering", either the inclusion or exclusion lead to ORF disruption. Oo, oocyte; Zy, zygote; Mo, morula; Bl, blastocyst; Gv, oocyte GV; MII, oocyte MII. **(b)** For ORF altering exons, the percentage of those that disrupt (red) or restore (blue) the ORF specifically in that transition. **(c)** Change in usage of the ZGA isoform (whether inclusion or exclusion) upon NMD depletion in HR1 cells based on the ORF impact predicted at the ZGA stage. **(d-e)** Change in ribosome engagement (PSI in high polysome fraction - PSI cytoplasmic fraction) of the ZGA isoform in embryonic stem cells (ESCs)(d) or HEK293 cells (e). **(f)** ΔPSI upon NMD depletion in HR1 cells of exons of different types of Mfuzz clusters changing at ZGA based on the impact on the ORF ("Disr", ORF disruptive; "Non-disr", non ORF disruptive). **(g,h)** Change in ribosome engagement in ESCs (g) or HEK293 cells (h) of different types of Mfuzz clusters changing at ZGA based on the impact on the ORF. * ($0.01 \leq P < 0.05$), ** ($0.001 \leq P < 0.01$) and *** ($0.001 < P$) indicate statistically significant differences respect to "Protein isoforms" (for c-e) or "Other (Non-disr)" (for f-g) based on Wilcoxon Sum-Rank tests. Data from [65] [c,f], [29] [d,g] and [30] [e,h].

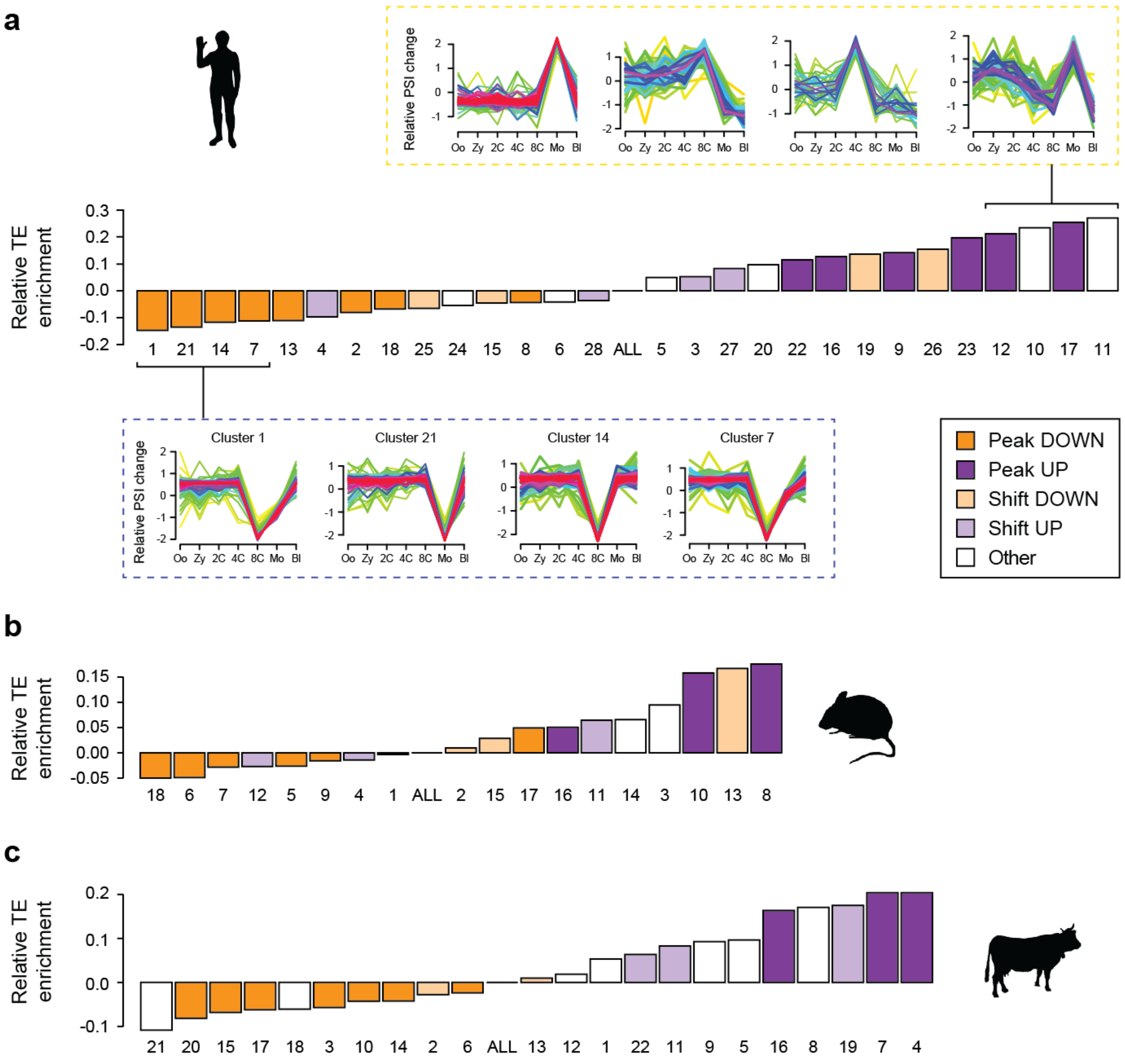


**Extended Data Fig. 7 - Overlap between exons and transposable elements. (a-c)** Relative enrichment of exons overlapping RepeatMasker repeats for each Mfuzz cluster for human (a), mouse (b) and cow (c). Peak, Shift or Other dynamics are indicated. Cluster IDs correspond to those in Supplementary Figs. 5-7.

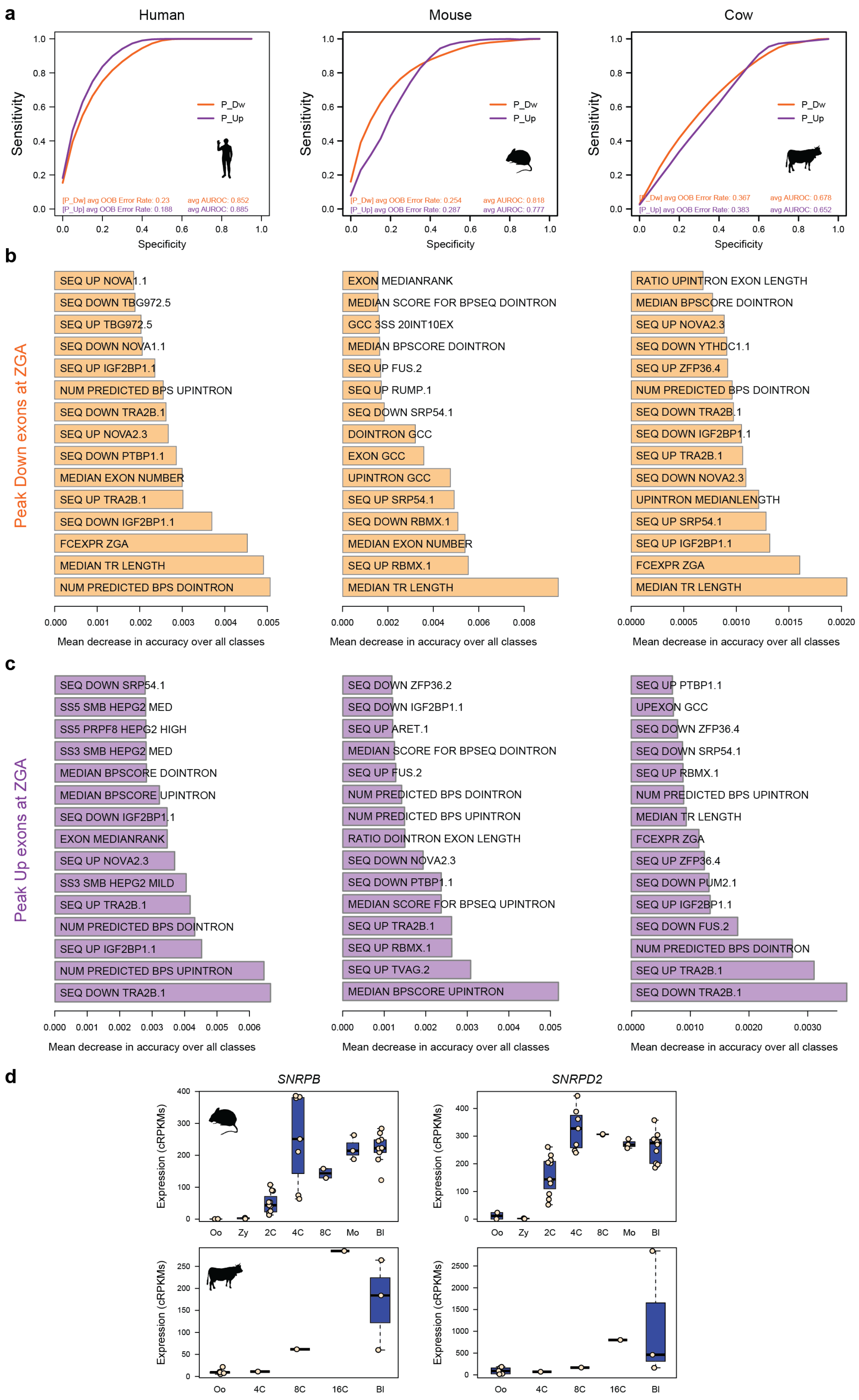
a
Human
Mouse
Cow
Sensitivity
Specificity
P_Dw
P_Up
[P_Dw] avg OOB Error Rate: 0.23 avg AUROC: 0.852
[P_Up] avg OOB Error Rate: 0.188 avg AUROC: 0.885
[P_Dw] avg OOB Error Rate: 0.254 avg AUROC: 0.818
[P_Up] avg OOB Error Rate: 0.287 avg AUROC: 0.777
[P_Dw] avg OOB Error Rate: 0.367 avg AUROC: 0.678
[P_Up] avg OOB Error Rate: 0.383 avg AUROC: 0.652
b
Peak Down exons at ZGA
SEQ UP NOVA1.1
SEQ DOWN TBG972.5
SEQ UP TBG972.5
SEQ DOWN NOVA1.1
SEQ UP IGF2BP1.1
NUM PREDICTED BPS UPINTRON
SEQ DOWN TRA2B.1
SEQ UP NOVA2.3
SEQ DOWN PTBP1.1
MEDIAN EXON NUMBER
SEQ UP TRA2B.1
SEQ DOWN IGF2BP1.1
FCEXPR ZGA
MEDIAN TR LENGTH
NUM PREDICTED BPS DOINTRON
EXON MEDIANRANK
MEDIAN SCORE FOR BPSEQ DOINTRON
GCC 3SS 20INT10EX
MEDIAN BPSCORE DOINTRON
SEQ UP FUS.2
SEQ UP RUMP.1
SEQ DOWN SRP54.1
DOINTRON GCC
EXON GCC
UPINTRON GCC
SEQ UP SRP54.1
SEQ DOWN RBMX.1
MEDIAN EXON NUMBER
SEQ UP RBMX.1
MEDIAN TR LENGTH
RATIO UPINTRON EXON LENGTH
MEDIAN BPSCORE DOINTRON
SEQ UP NOVA2.3
SEQ DOWN YTHDC1.1
SEQ UP ZFP36.4
NUM PREDICTED BPS DOINTRON
SEQ DOWN TRA2B.1
SEQ DOWN IGF2BP1.1
SEQ UP TRA2B.1
SEQ DOWN NOVA2.3
UPINTRON MEDIANLENGTH
SEQ UP SRP54.1
SEQ UP IGF2BP1.1
FCEXPR ZGA
MEDIAN TR LENGTH
Mean decrease in accuracy over all classes
c
Peak Up exons at ZGA
SEQ DOWN SRP54.1
SS5 SMB HEPG2 MED
SS5 PRPF8 HEPG2 HIGH
SS3 SMB HEPG2 MED
MEDIAN BPSCORE DOINTRON
MEDIAN BPSCORE UPINTRON
SEQ DOWN IGF2BP1.1
EXON MEDIANRANK
SEQ UP NOVA2.3
SS3 SMB HEPG2 MILD
SEQ UP TRA2B.1
NUM PREDICTED BPS DOINTRON
SEQ UP IGF2BP1.1
NUM PREDICTED BPS UPINTRON
SEQ DOWN TRA2B.1
SEQ DOWN ZFP36.2
SEQ DOWN IGF2BP1.1
SEQ UP ARET.1
MEDIAN SCORE FOR BPSEQ DOINTRON
SEQ UP FUS.2
NUM PREDICTED BPS DOINTRON
NUM PREDICTED BPS UPINTRON
RATIO DOINTRON EXON LENGTH
SEQ DOWN NOVA2.3
SEQ DOWN PTBP1.1
MEDIAN SCORE FOR BPSEQ UPINTRON
SEQ UP TRA2B.1
SEQ UP RBMX.1
SEQ UP TVAG.2
MEDIAN BPSCORE UPINTRON
SEQ UP PTBP1.1
UPEXON GCC
SEQ DOWN ZFP36.4
SEQ DOWN SRP54.1
SEQ UP RBMX.1
NUM PREDICTED BPS UPINTRON
MEDIAN TR LENGTH
FCEXPR ZGA
SEQ UP ZFP36.4
SEQ DOWN PUM2.1
SEQ UP IGF2BP1.1
SEQ DOWN FUS.2
NUM PREDICTED BPS DOINTRON
SEQ UP TRA2B.1
SEQ DOWN TRA2B.1
Mean decrease in accuracy over all classes
d
SNRPB
SNRPD2
Expression (cRPKMs)
Oo Zy 2C 4C 8C Mo Bl
Oo 4C 8C 16C Bl

**Extended Data Fig. 8 - Random Forest classification of exons with peak dynamics at ZGA and *SNRPB/D2* expression. (a)** Receiver operating characteristic curve (ROC) for the Random Forest classification of exons with peak-down (orange, P_Dw) or peak-up (purple, P_Up) versus a matched set of background exons with similar pre-ZGA PSI distributions for each species. Average OOB error rate and area under the ROC are indicated for each set for 1000 random Random Forest models. **(b,c)** Most discriminative features for peak-down (b) or peak-up (c) exons for each species quantified as mean decrease in accuracy. Precise definitions for each feature can be obtained on http://matt.crg.eu. **(d)** cRPKM metric is used to quantify GE for *Snrpb* and *Snrpd2* in mouse and cow.

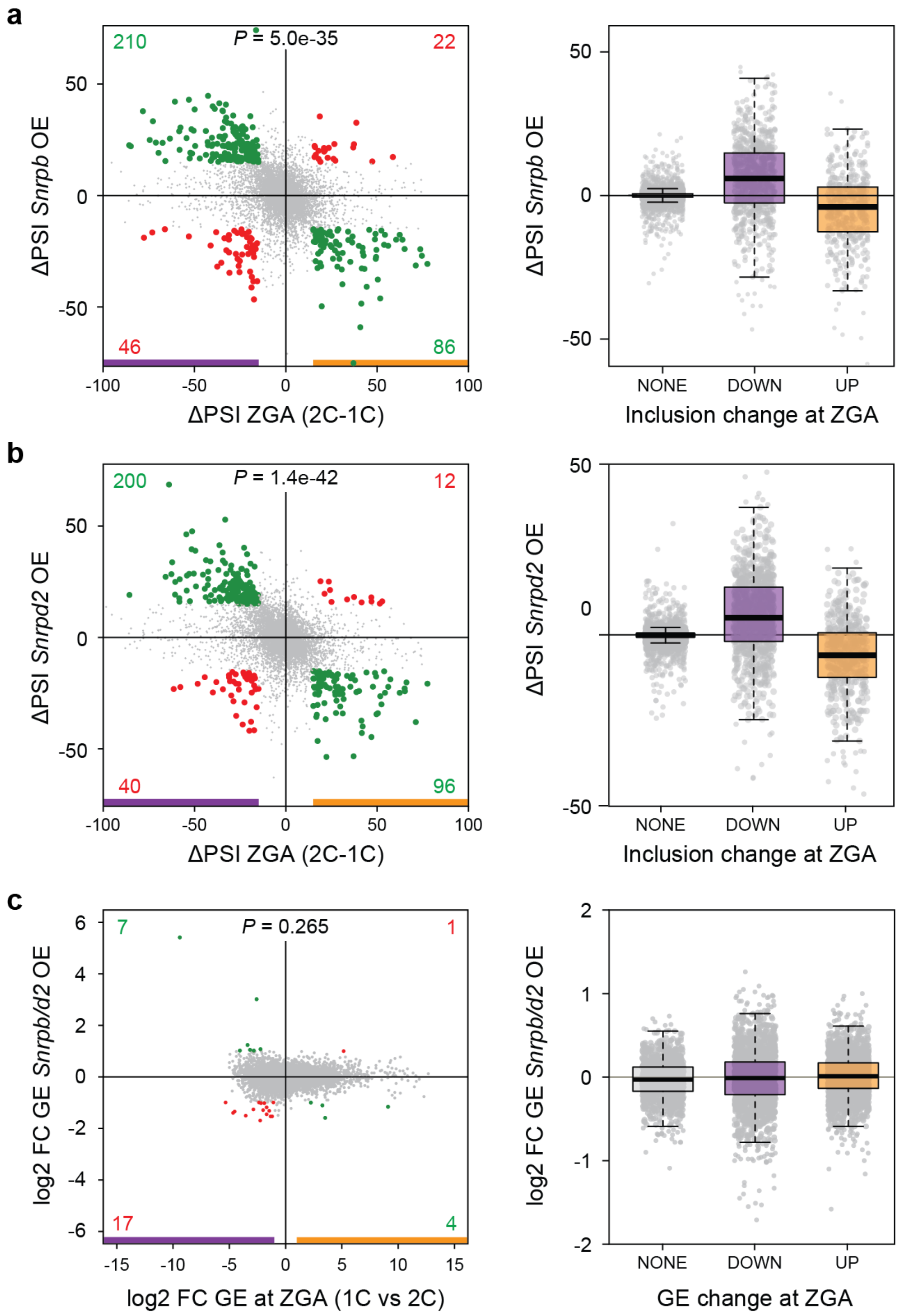


**Extended Data Fig. 9 - Reversion of ZGA transcriptomic changes by *Snrpb/d2* earlier expression. (a,b)** Left: Changes in exon inclusion levels at 2C stage upon early-induced expression of *Snrpb* (a) or *Snrpd2* (b) (Y-axis) respect to the observed change at ZGA (2C-1C; X-axis). Exons with |ΔPSI| ≥ 15 upon *Snrpb* or *Snrpd2* expression and at ZGA are highlighted in green/red and numbers indicated. Right: Distribution of ΔPSI upon *Snrpb* or *Snrpd2* induced expression for all exons with increased (UP), decreased (DOWN) or no change (NONE) in PSI at ZGA. **(c)** Left: Changes in GE levels at 2C stage upon early induced expression of *Snrpb/d2* (Y-axis) respect to the observed change at ZGA (2C-1C; X-axis). Genes with |log2(FC)| ≥ 1 upon *Snrpb/d2* expression and at ZGA are highlighted in green/red and numbers indicated. Right: Distribution of log2(FC) upon *Snrpb/d2* induced expression for all exons with increased (UP), decreased (DOWN) or no change (NONE) in GE at ZGA. *P*-values correspond to a two-sided Binomial test between Q2+Q4 versus Q1+Q3.

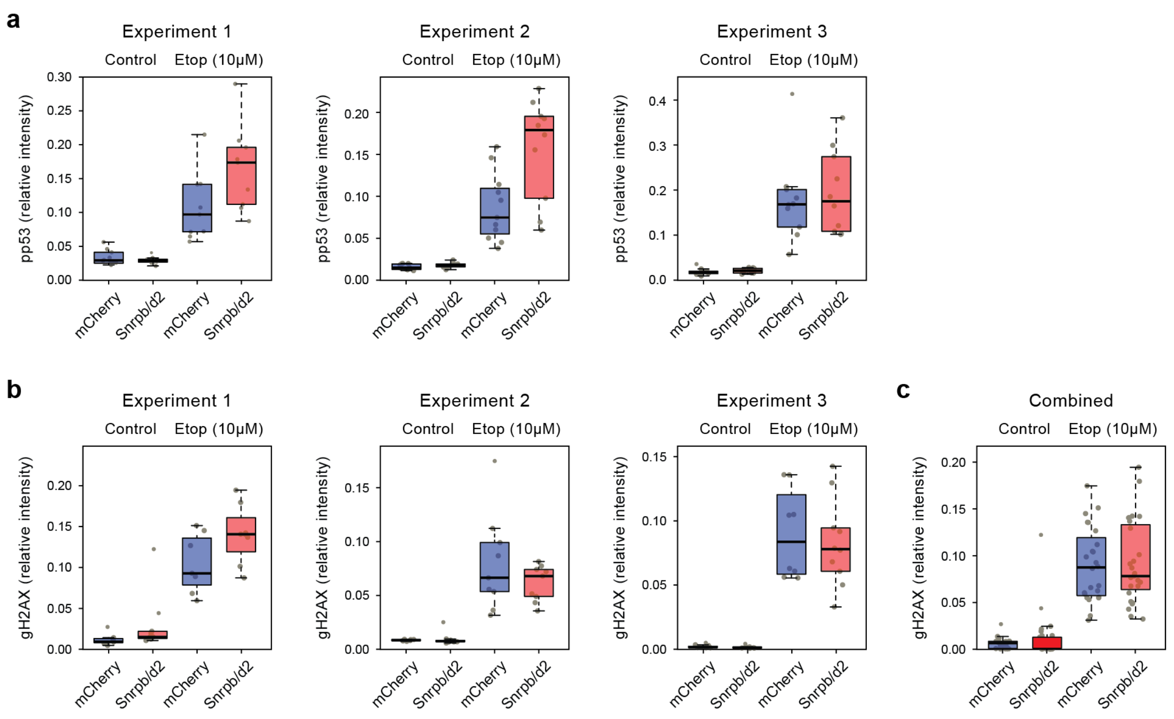


**Extended Data Fig. 10 - DNA damage response is modulated by Snrpb/d2 earlier expression. (a,b)** Quantification of phospho-p53 (Ser15, pp53) (a) or gamma-H2AX (b) immunostaining levels from 2C embryos injected at pronuclear stage with *Snrpb/d2* or mCherry mRNA and treated for 1.5h with 10uM etoposide or left untreated. Each plot represents an independent experiment and each dot represents the average relative intensity of both cells of a single embryo. **(c)** Combined data from all three individual experiments in (b). Total number of embryos for each condition and experiment in (a): Experiment 1: n=10 (mCherry untreated), n=9 (Snrpb/d2, untreated), n=9 (mCherry, etoposide) and n=9 (Snrpb/d2, etoposide); Experiment 2: n=9 (mCherry untreated), n=8 (Snrpb/d2, untreated), n=11 (mCherry, etoposide) and n=10 (Snrpb/d2, etoposide); Experiment 3: n=10 (mCherry untreated), n=9 (Snrpb/d2, untreated), n=10 (mCherry, etoposide) and n=10 (Snrpb/d2, etoposide). Total number of embryos for each condition and experiment in (b): Experiment 1: n=7 (mCherry untreated), n=9 (Snrpb/d2, untreated), n=7 (mCherry, etoposide) and n=8 (Snrpb/d2, etoposide); Experiment 2: n=8 (mCherry untreated), n=11 (Snrpb/d2, untreated), n=9 (mCherry, etoposide) and n=9 (Snrpb/d2, etoposide); Experiment 3: n=11 (mCherry untreated), n=10 (Snrpb/d2, untreated), n=8 (mCherry, etoposide) and n=10 (Snrpb/d2, etoposide); Combined (c): n=26 (mCherry untreated), n=30 (Snrpb/d2, untreated), n=24 (mCherry, etoposide) and n=27 (Snrpb/d2, etoposide).

#### Supplementary Figures

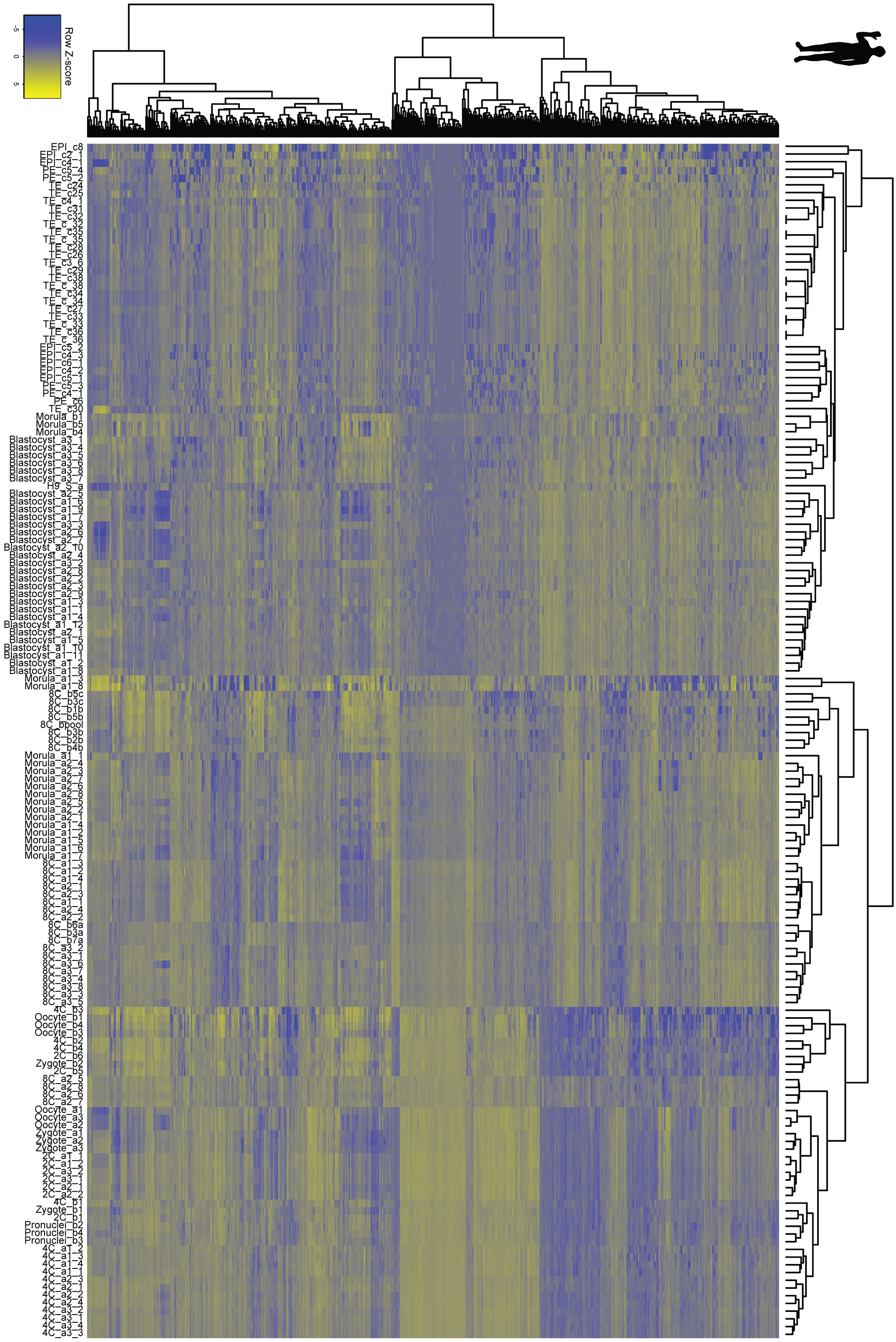

**Supplementary Fig. 1 - Hierarchical clustering of human early development samples.** Heatmaps and unsupervised hierarchical clustering using default *heatmap.2* parameters of the top 500 most variably expressed genes as determined by *DESeq* for each studied species. These clustering outputs were used to determine cell identity and locate anomalous cells (see Supplementary Table 1 for details).

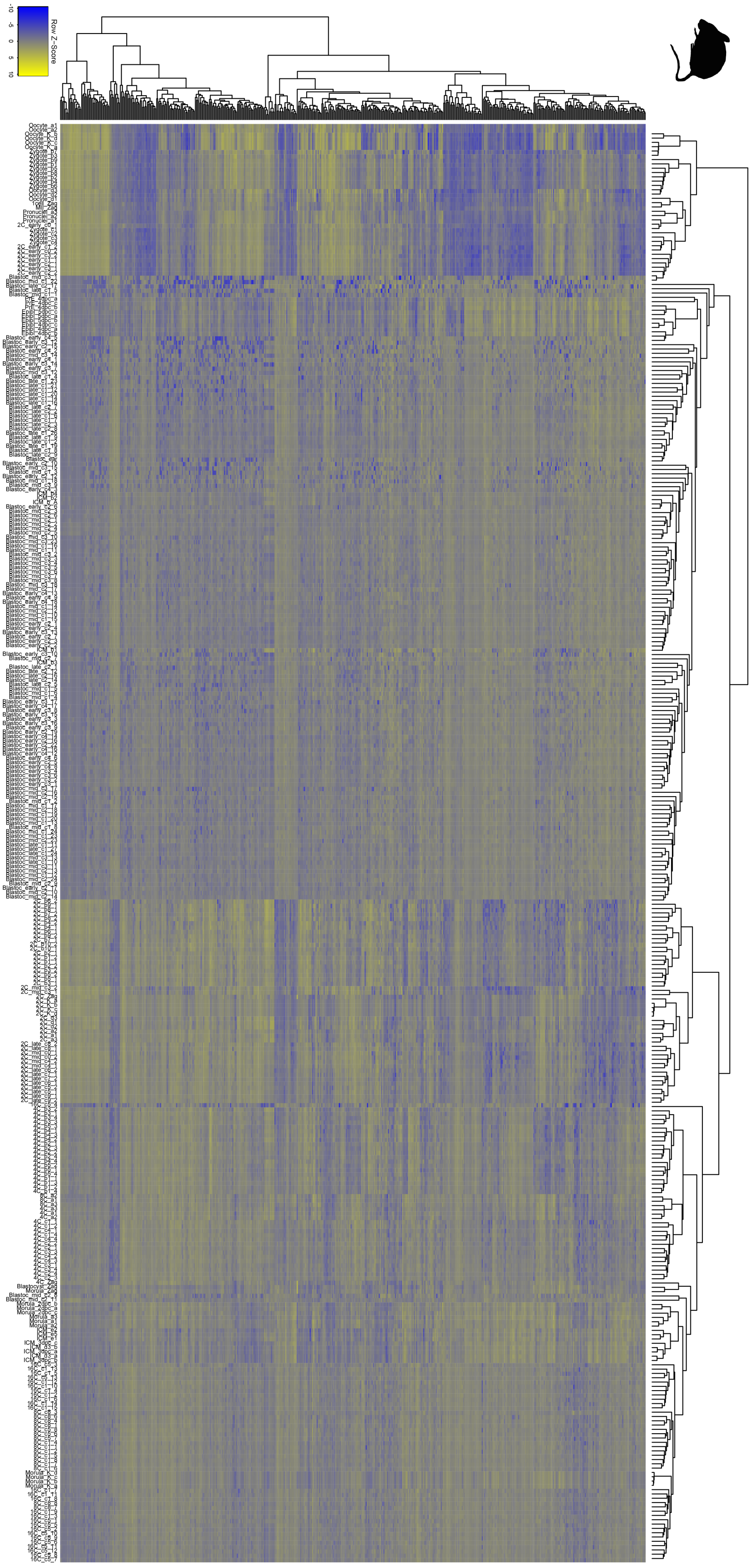
Row Z-Score

**Supplementary Fig. 2 - Hierarchical clustering of mouse early development samples.** Heatmaps and unsupervised hierarchical clustering using default *heatmap.2* parameters of the top 500 most variably expressed genes as determined by *DESeq* for each studied species. These clustering outputs were used to determine cell identity and locate anomalous cells (see Supplementary Table 1 for details).

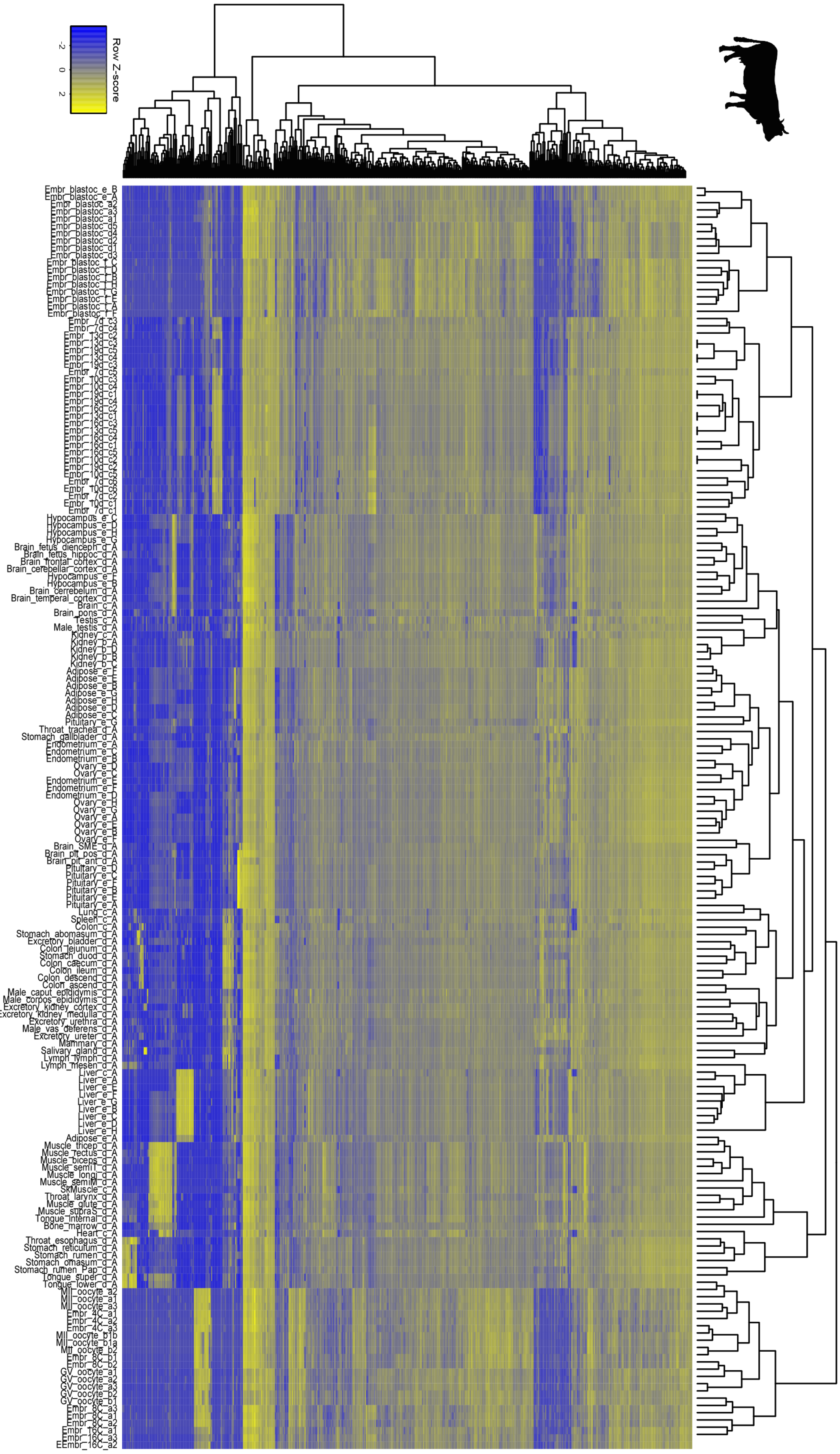
Row Z-score
-2
0
2

**Supplementary Fig. 3 - Hierarchical clustering of cow early development samples.** Heatmaps and unsupervised hierarchical clustering using default *heatmap.2* parameters of the top 500 most variably expressed genes as determined by *DESeq* for each studied species. These clustering outputs were used to determine cell identity and locate anomalous cells (see Supplementary Table 1 for details).

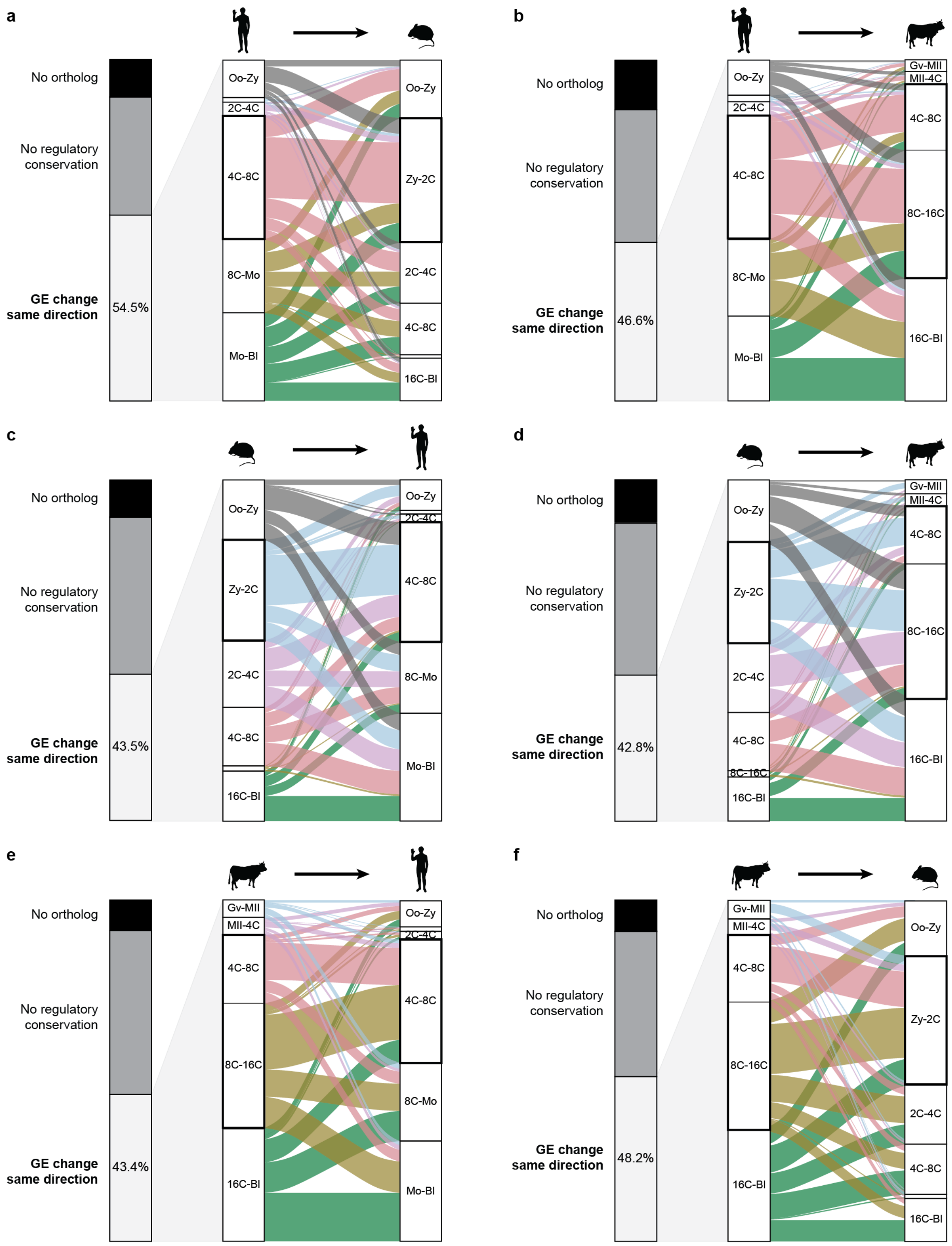


**Supplementary Fig. 4 - Pairwise evolutionary conservation of GE changes. (a-f)** For each query and target species, stack plot of the level of conservation for each gene whose GE levels change at any transition in the query species. No ortholog: absence of one to one orthologs in the target species; No regulatory conservation: the orthologous gene does not significantly change in the same direction at any of the transitions in the target species. From the percentage of genes that change expression in the same direction in at least one transition of the target species, alluvial plot showing the transition of change in query species (left) and the observed highest change in the same direction in the target species (right). (a) Human to mouse, (b) human to cow, (c) mouse to human, (d) mouse to cow, (e) cow to human, (f) cow to mouse.

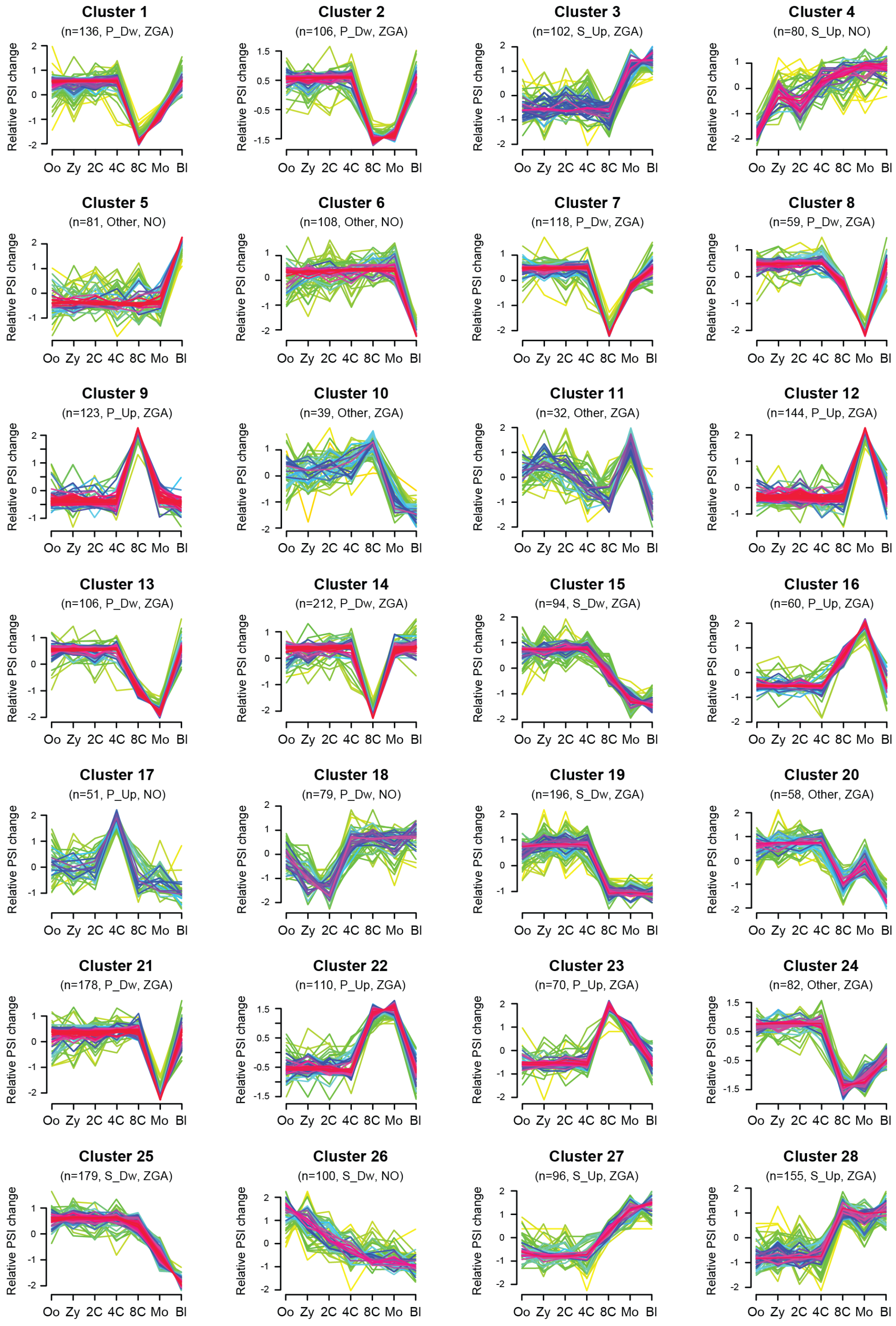


**Supplementary Fig. 5 - Mfuzz exon clusters in human, mouse and cow.** Profiles of scaled PSIs for each exon within each Mfuzz cluster in each studied species. For each cluster, the number of exons (n), the most common pattern (Peak, Shift or Other, and direction) and whether or not it is considered to change at ZGA. P_Dw, peak-down; P_Up, peak-up; S_Dw, shift-down; S_Up, shift-up.

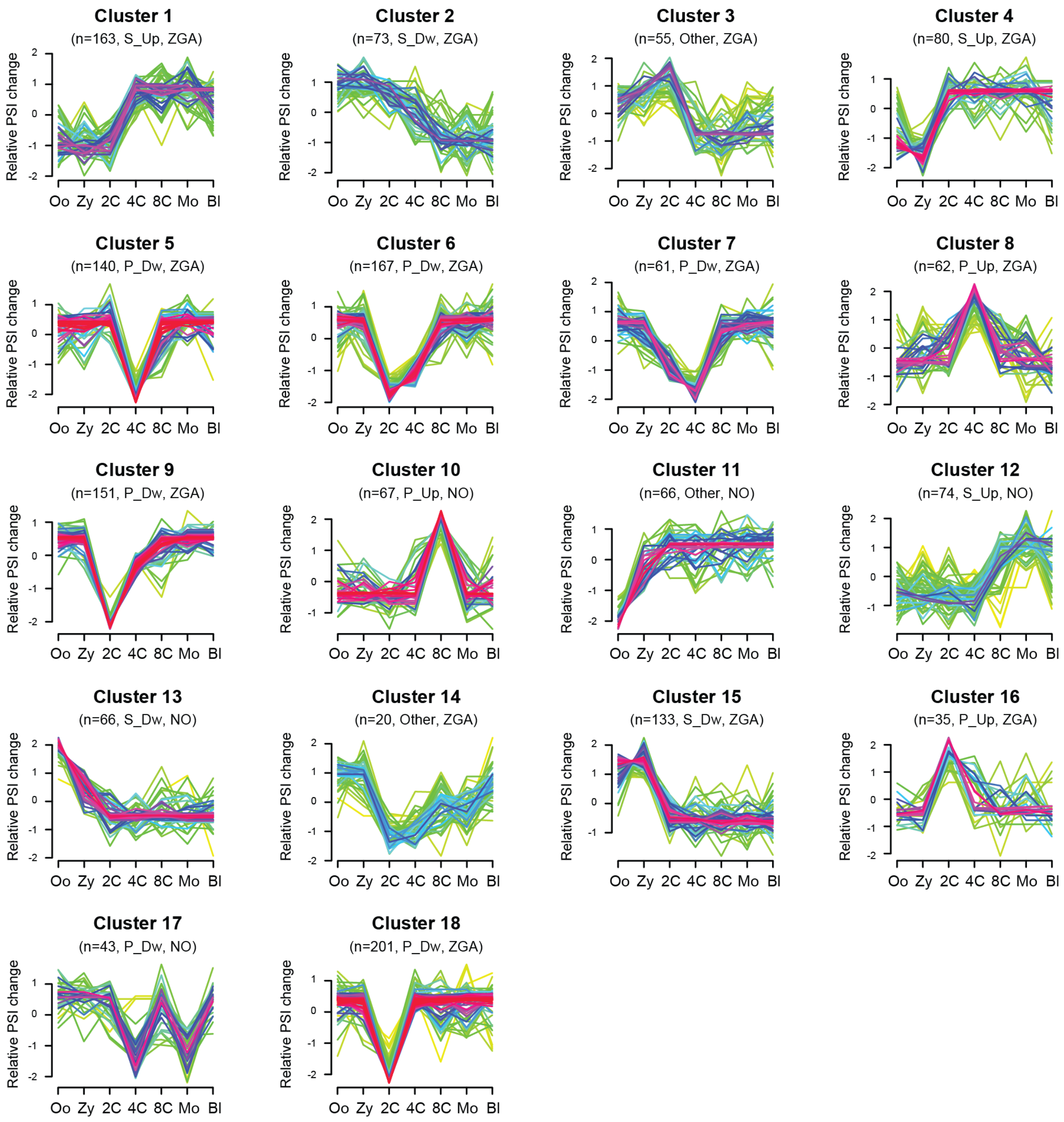


**Supplementary Fig. 6 - Mfuzz exon clusters in human, mouse and cow.** Profiles of scaled PSIs for each exon within each Mfuzz cluster in each studied species. For each cluster, the number of exons (n), the most common pattern (Peak, Shift or Other, and direction) and whether or not it is considered to change at ZGA. P_Dw, peak-down; P_Up, peak-up; S_Dw, shift-down; S_Up, shift-up.

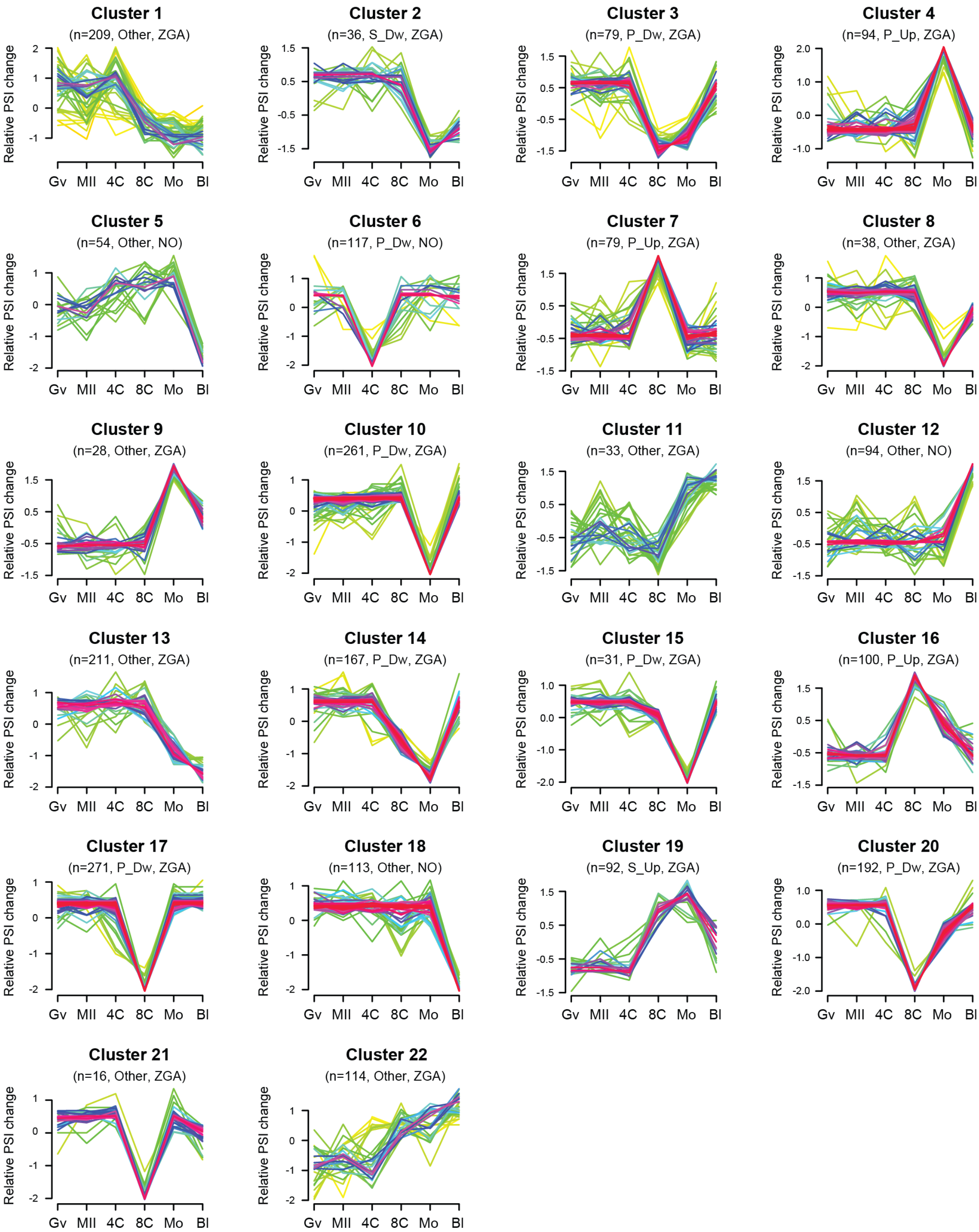


**Supplementary Fig. 7 - Mfuzz exon clusters in human, mouse and cow.** Profiles of scaled PSIs for each exon within each Mfuzz cluster in each studied species. For each cluster, the number of exons (n), the most common pattern (Peak, Shift or Other, and direction) and whether or not it is considered to change at ZGA. P_Dw, peak-down; P_Up, peak-up; S_Dw, shift-down; S_Up, shift-up.
