## Supplementary File 1 for "A developmentally programmed splicing failure attenuates the DNA damage response during mammalian zygotic genome activation"

### Comparison of exons grouped into: Bg-Dwstrat, P-Dw, Bg-Upstrat, P-Up, HIGH-PSI, LOW-PSI

September 16, 2020
Matt version 1.3.0

#### Contents

**1 Infos** 4

**2 Warning: Please read this note carefully** 4

**3 Notes for publishing results** 4

**4 Data sets** 5

**5 Overview: Features with statistically significant differences (p-val $\leq 0.05$)** 6

**6 Details: Box plots and statistical assessments for all features** 23

### 1 Infos

Visualizations of exon features for different groups of exons. Each exon occurs in exactly one gene, but might occur in several transcripts of that gene. Hence, for some features like the exon length, there is exactly one value for each exon. For other features, e.g., length of the up-stream exon(s), which could be different in different transcripts, there might be several values for each exon. Consequently, in the latter cases, the median of these value gets reported.

### 2 Warning: Please read this note carefully

Please keep in mind that some features might affect other features. Especially: all branch-point features get extracted from sub-sequences of introns, by standard the last 150 nt at the 3' end of each intron (if you haven't changed this) always neglecting the first 20 nt at their 5' end. If introns of one set are especially short, i.e., many are shorter then these 150 nt, then the shorter intron length might affect branch-point features. For example, there might be less branch points found in shorter introns or their distance to the 3' intron ends might be generally shorter simply because of their shorter intron length.

### 3 Notes for publishing results

The Matt paper: *Matt: Unix tools for alternative splicing analysis, A. Gohr, M. Irimia, Bioinformatics, 2018, bty606, DOI: 10.1093/bioinformatics/bty606*
When publishing results wrt. splice site strengths which you determined for your data using matt, please cite: *Maximum entropy modeling of short sequence motifs with applications to RNA splicing signals, Yeo et al., 2003, DOI: 10.1089/1066527041410418*
When publishing results wrt. branch point features which you determined for your data with matt, please cite: *Genome-wide association between branch point properties and alternative splicing, Corvelo et al., 2010, DOI: 10.1371/journal.pcbi.1001016*
When publishing results with respect to the binding strength of the human Sf1 splicing factor, you might refert to where the Sf1 binding motif comes from: *Analysis of in situ pre-mRNA targets of human splicing factor SF1 reveals a function in alternative splicing, Margherita Corioni, Nicolas Antih, Goranka Tanackovic, Mihaela Zavolan, and Angela Kramer, 2011, DOI: 10.1093/nar/gkq1042*
The Sf1 binding motif is described in supplement, page 13, table S2: Weight matrix of the binding specificity of SF1.

### 4 Data sets

Input file:
../hsa_final.tab
Selection criteria for defining exons groups:
Bg_Dwstrat : having value Bg_Dwstrat in column GROUP
P_Dw : having value P_Dw in column GROUP
Bg_Upstrat : having value Bg_Upstrat in column GROUP
P_Up : having value P_Up in column GROUP
HIGH_PSI : having value HIGH_PSI in column GROUP
LOW_PSI : having value LOW_PSI in column GROUP
Exon duplicates removal: yes
Numbers of exons per group before / after neglecting exons which were not found in GTF file (gene annotation). For the comparisons only exons which were found in the gene annotation are used. These numbers might change slightly for each feature if NAs occur.
Bg_Dwstrat: 2104 / 2094
P_Dw: 526 / 526
Bg_Upstrat: 944 / 937
P_Up: 236 / 232
HIGH_PSI: 11878 / 11841
LOW_PSI: 4077 / 4065

### 5 Overview: Features with statistically significant differences (p-val $\leq 0.05$)

#### MAXENTSCR HSAMODEL 5SS

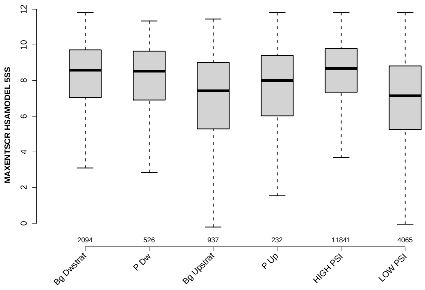

#### MAXENTSCR HSAMODEL 3SS

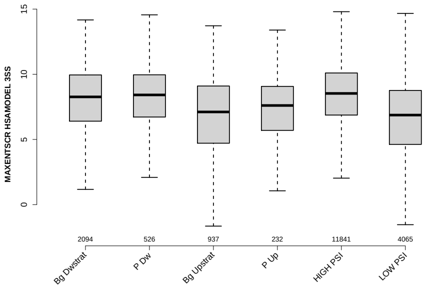

#### PROP EXON IN UTR

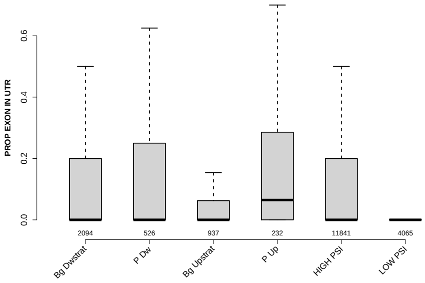

#### PROP INTERNAL EXON

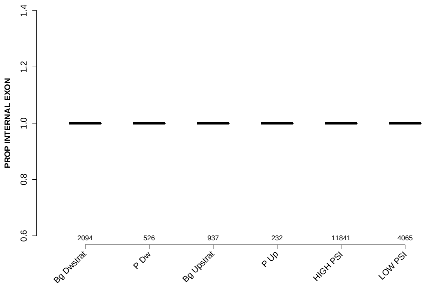

### PROP LAST EXON

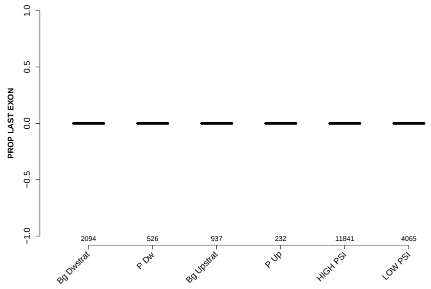

### MEDIAN EXON NUMBER

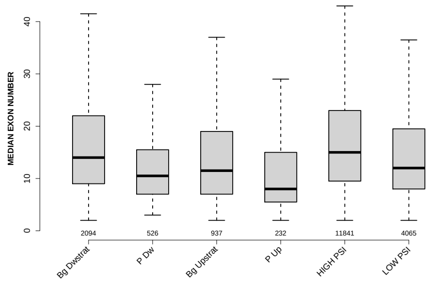

### EXON LENGTH

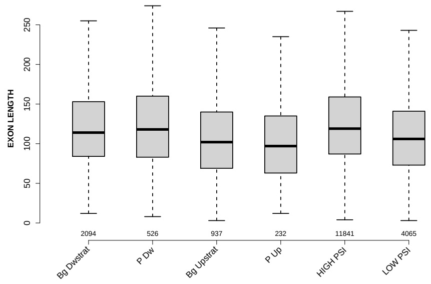

### PROP FIRST EXON

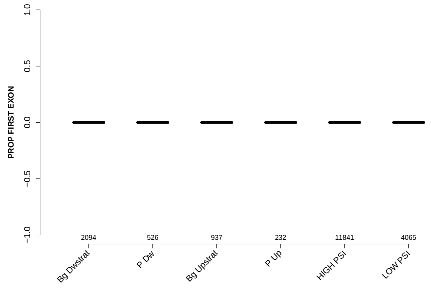

### RATIO DOINTRON EXON GCC

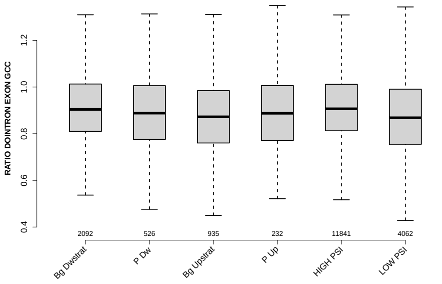

### MEDIAN TR LENGTH

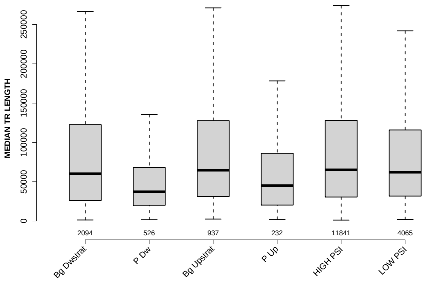

### RATIO DOEXON EXON LENGTH

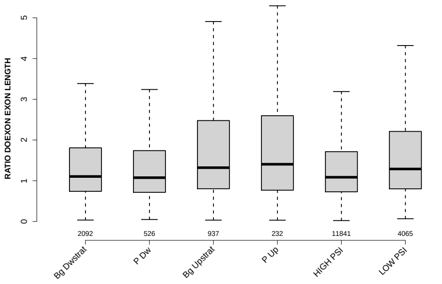

### RATIO UPEXON EXON LENGTH

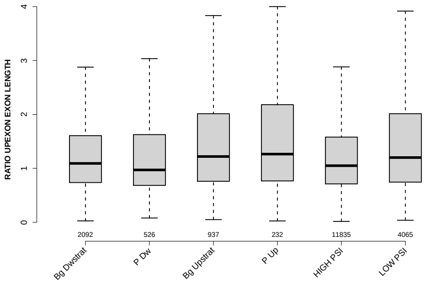

### RATIO UPINTRON EXON GCC

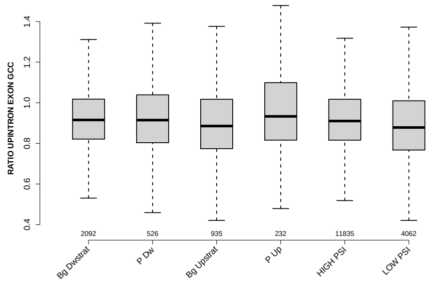

### GCC 3SS 20INT10EX

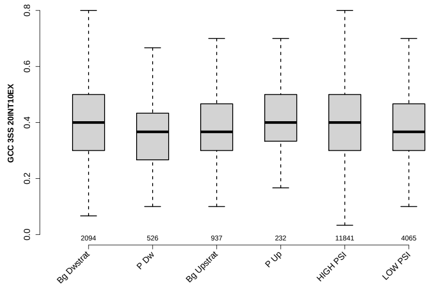

### MEDIAN DIST FROM BP TO 3SS UPINTRON

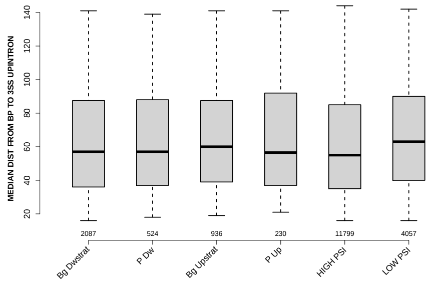

### RATIO DOINTRON EXON LENGTH

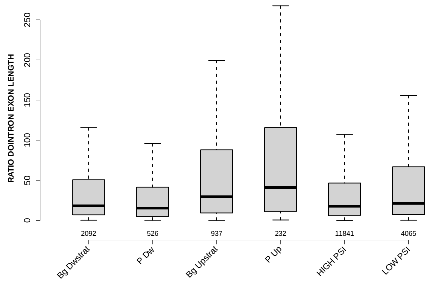

### DOEXON GCC

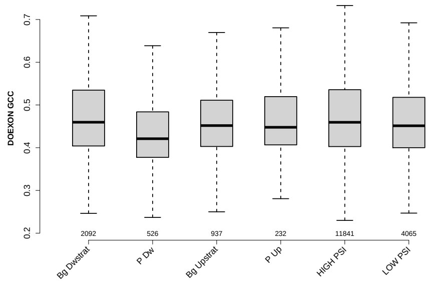

### SCORE FOR MAXBP SEQ UPINTRON

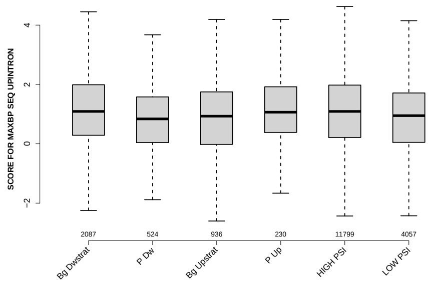

### DIST FROM MAXBP TO 3SS UPINTRON

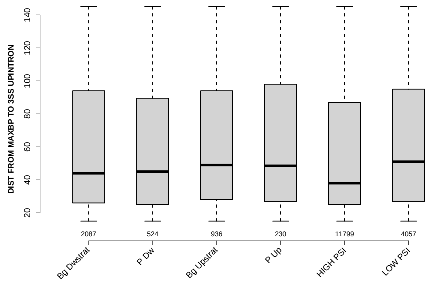

### GCC 5SS 20INT10EX

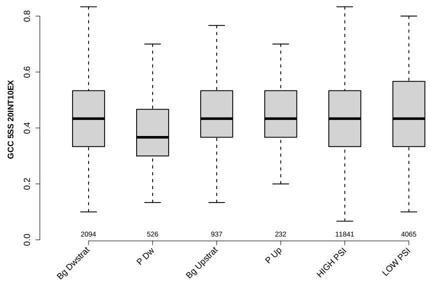

#### DOINTRON GCC

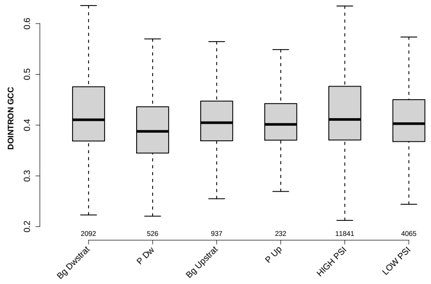

#### EXON GCC

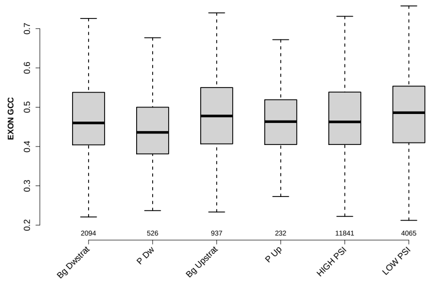

#### DO 3SS 20INT10EX GCC

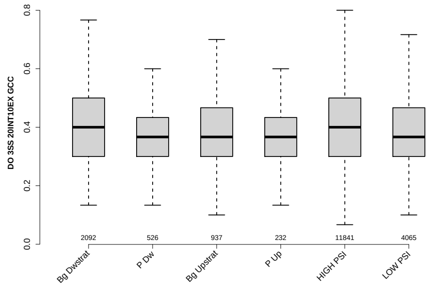

#### RATIO UPINTRON EXON LENGTH

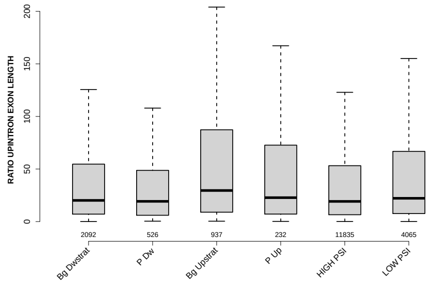

#### UPEXON GCC

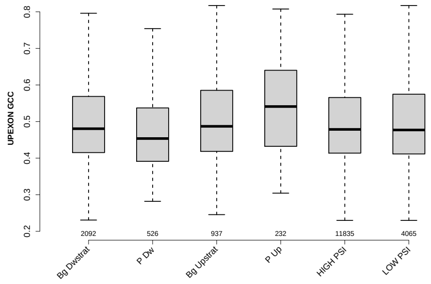

#### BPSCORE MAXBP UPINTRON

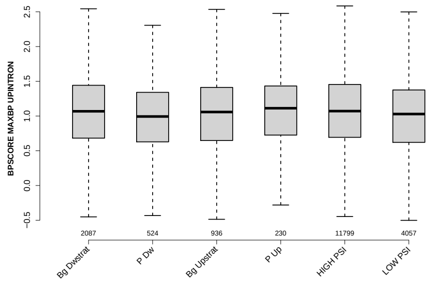

#### RATIO DOEXON EXON GCC

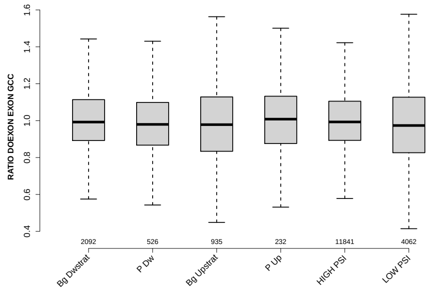

#### MEDIAN SCORE FOR BPSEQ UPINTRON

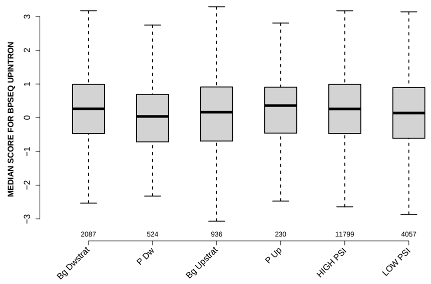

#### RATIO UPEXON EXON GCC

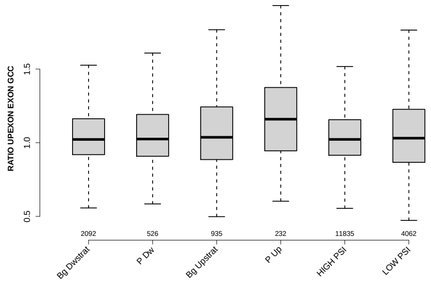

#### PYRIMIDINECONT MAXBP UPINTRON

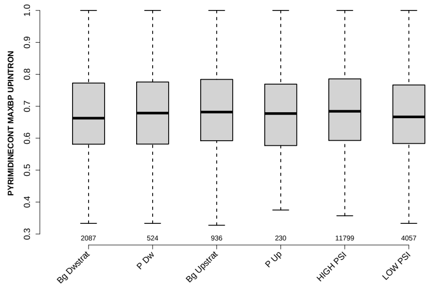

#### DOINTRON MEDIANLENGTH

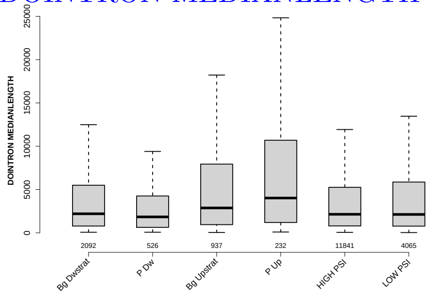

#### UPINTRON GCC

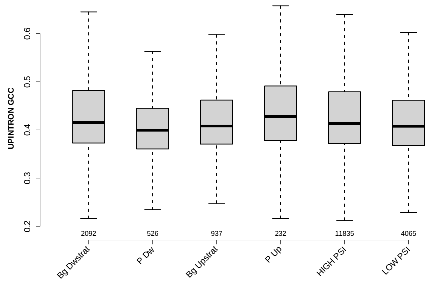

#### UP 5SS 20INT10EX GCC

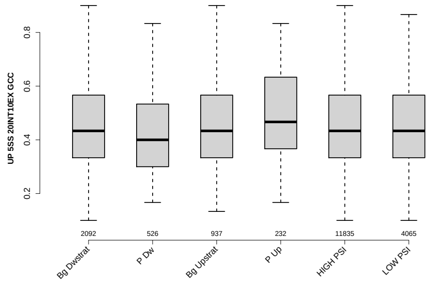

#### SCORE FOR MAXBP SEQ DOINTRON

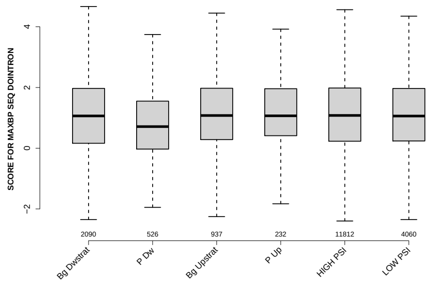

#### SF1 HIGHESTSCORE 3SS UPINTRON

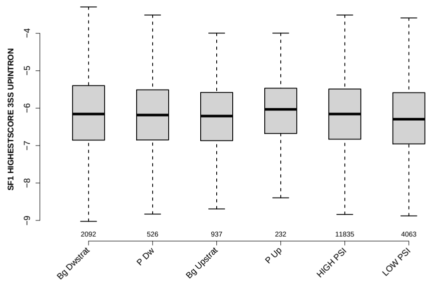

#### NTRS ALL FOR GENE

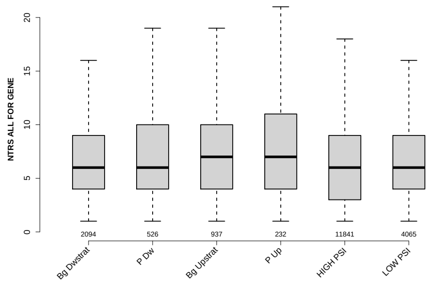

#### MAXENTSCR HSAMODEL UPSTRM 5SS

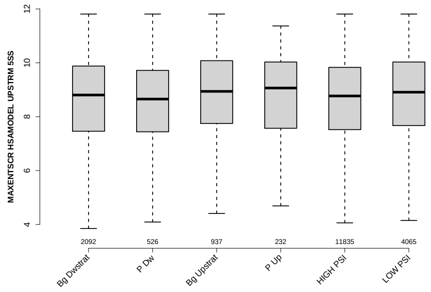

#### MEDIAN PYRIMIDINECONT UPINTRON

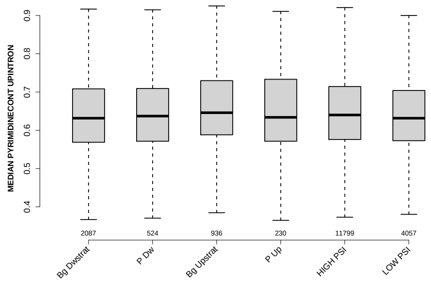

#### EXON MEDIANRELATIVERANK 5BINS

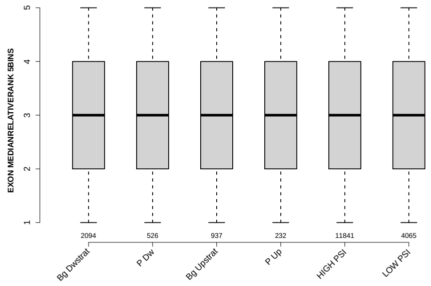

#### POLYPYRITRAC SCORE MAXBP UPINTRON

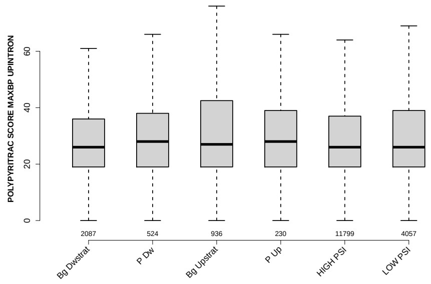

#### EXON MEDIANRELATIVERANK

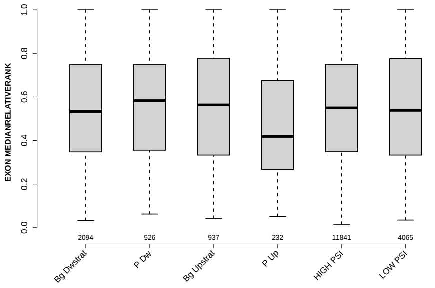

#### EXON MEDIANRELATIVERANK 10BINS

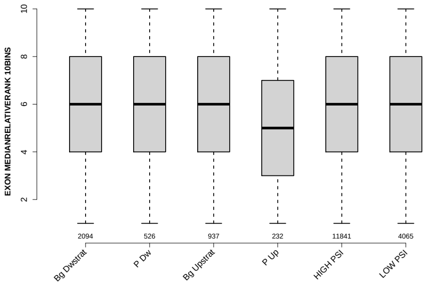

#### POLYPYRITRAC LEN MAXBP UPINTRON

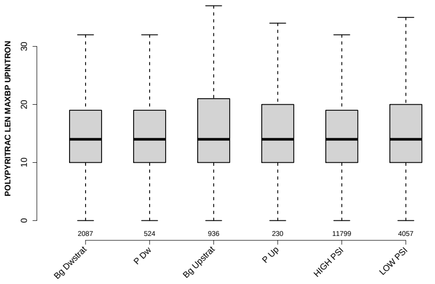

#### BPSCORE MAXBP DOINTRON

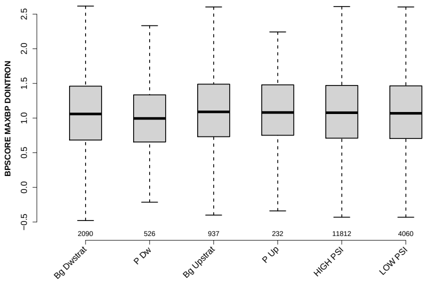

#### MEDIAN BPSCORE UPINTRON

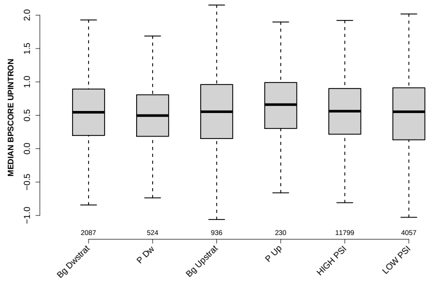

#### SF1 HIGHESTSCORE 3SS DOINTRON

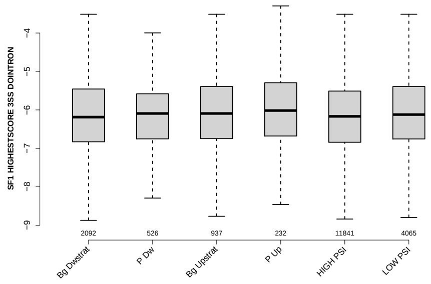

#### NUM PREDICTED BPS UPINTRON

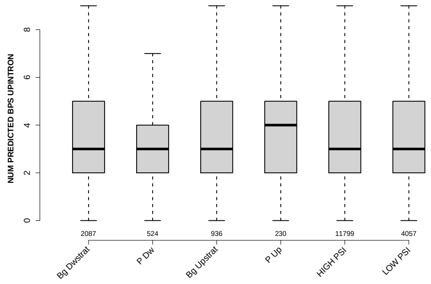

#### MEDIAN POLYPYRITRAC LEN UPINTRON

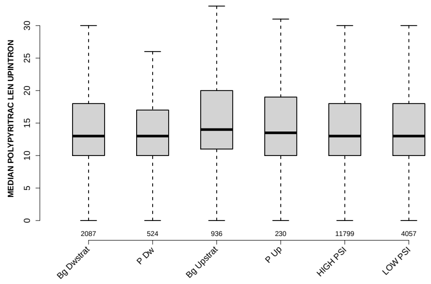

### UPINTRON MEDIANLENGTH

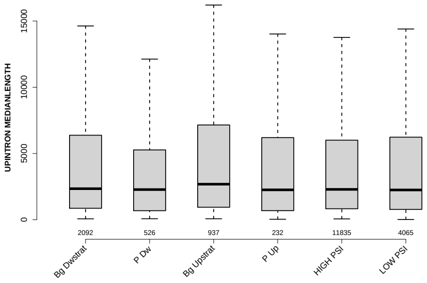

### EXON MEDIANRELATIVERANK 3BINS

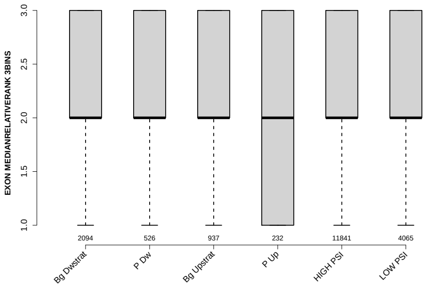

### MEDIAN POLYPYRITRAC SCORE UPINTRON

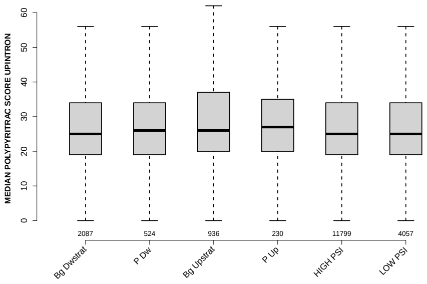

### POLYPYRITRAC OFFSET MAXBP DOINTRON

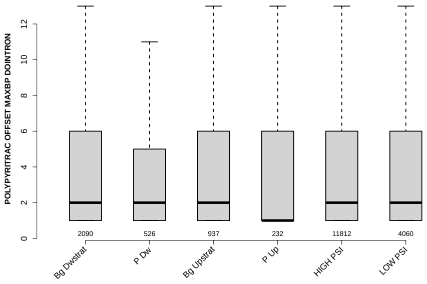

#### MAXENTSCR HSAMODEL DOWNSTRM 3SS

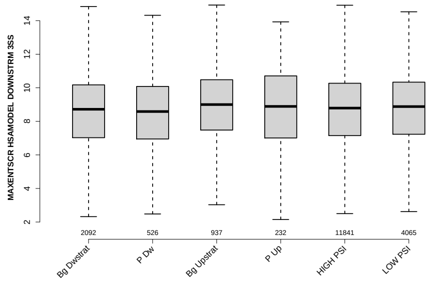

#### NUM PREDICTED BPS DOINTRON

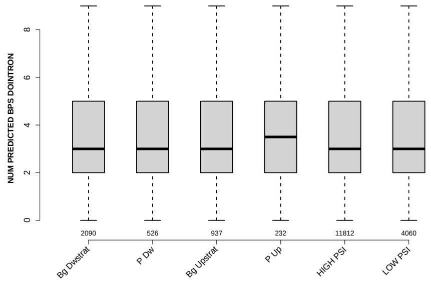

#### MEDIAN SCORE FOR BPSEQ DOINTRON

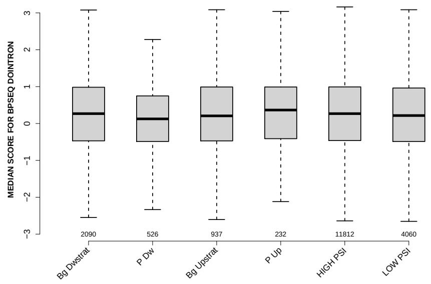

#### UPEXON MEDIANLENGTH

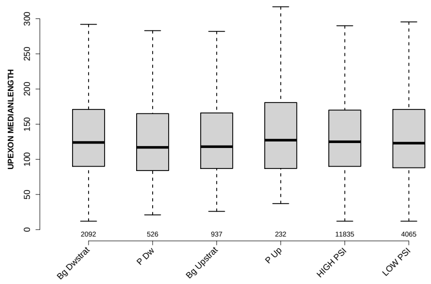

### MEDIAN POLYPYRITRAC SCORE DOINTRON

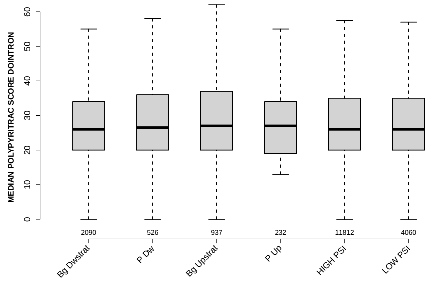

### MEDIAN PYRIMIDINECONT DOINTRON

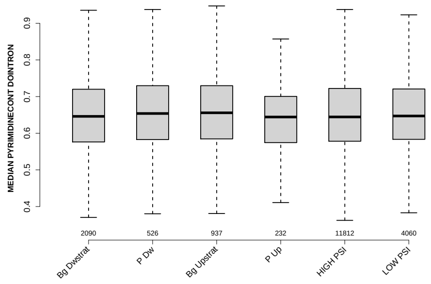

### MEDIAN POLYPYRITRAC OFFSET DOINTRON

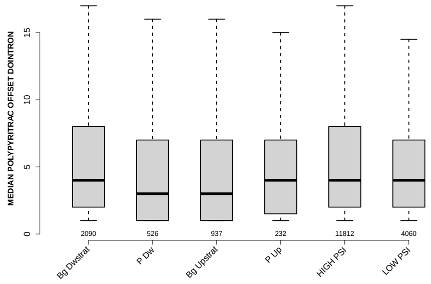

### DOEXON MEDIANLENGTH

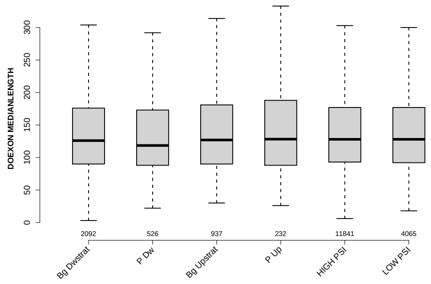

### POLYPYRITRAC OFFSET MAXBP UPINTRON

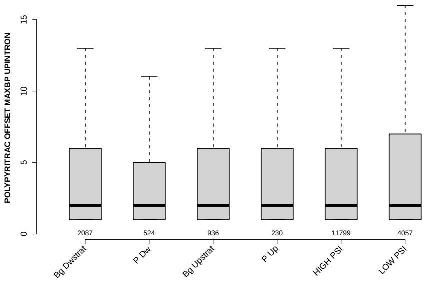

### POLYPYRITRAC SCORE MAXBP DOINTRON

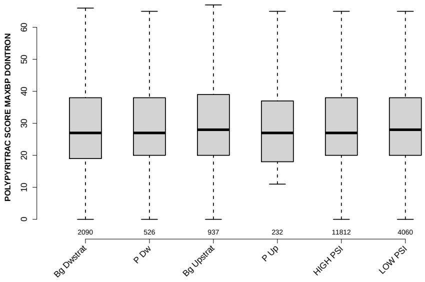

### PYRIMIDINECONT MAXBP DOINTRON

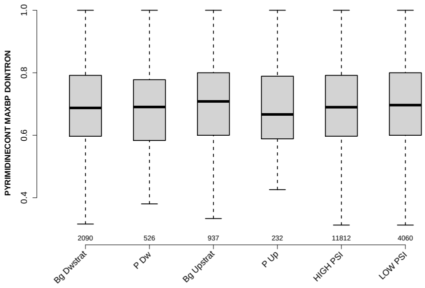

### MEDIAN POLYPYRITRAC OFFSET UPINTRON

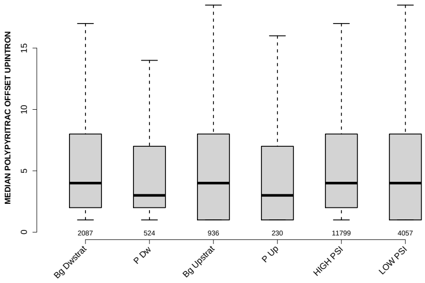

### DIST FROM MAXBP TO 3SS DOINTRON

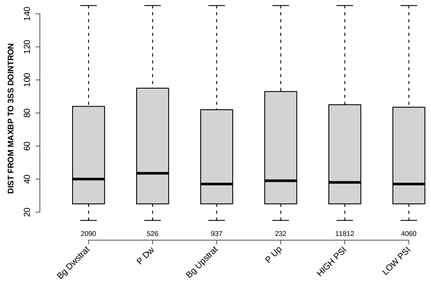

### 6 Details: Box plots and statistical assessments for all features

#### 6.1 EXON LENGTH

Back to: Overview | ToC

Meaning:

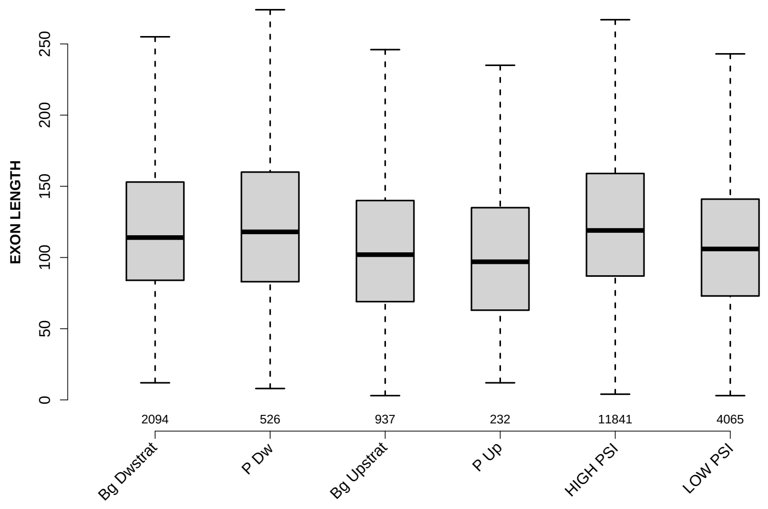

Significant results from Mann-Whitney U test:

- `Bg_Dwstrat` vs `Bg_Upstrat` : 2.46732e-10
  mean: 139.4365 > 121.5219 , median: 114 > 102
- `Bg_Dwstrat` vs `P_Up` : 4.58365e-06
  mean: 139.4365 < 148.6293 , median: 114 > 97
- `Bg_Dwstrat` vs `HIGH_PSI` : 0.00105241
  mean: 139.4365 < 140.4399 , median: 114 < 119
- `Bg_Dwstrat` vs `LOW_PSI` : 2.21673e-13
  mean: 139.4365 > 116.045 , median: 114 > 106
- `P_Dw` vs `Bg_Upstrat` : 3.02178e-06
  mean: 129.0665 > 121.5219 , median: 118 > 102
- `P_Dw` vs `P_Up` : 5.22511e-05
  mean: 129.0665 < 148.6293 , median: 118 > 97
- `P_Dw` vs `LOW_PSI` : 8.73518e-06
  mean: 129.0665 > 116.045 , median: 118 > 106
- `Bg_Upstrat` vs `HIGH_PSI` : 5.99591e-21
  mean: 121.5219 < 140.4399 , median: 102 < 119

- `P_Up` vs `HIGH_PSI` : 7.0238e-09
  mean: 148.6293 $>$ 140.4399 , median: 97 $<$ 119
- `HIGH_PSI` vs `LOW_PSI` : 2.37328e-50
  mean: 140.4399 $>$ 116.045 , median: 119 $>$ 106

#### 6.2 UPEXON MEDIANLENGTH

Back to: Overview | ToC

Meaning: median length of up-stream exon

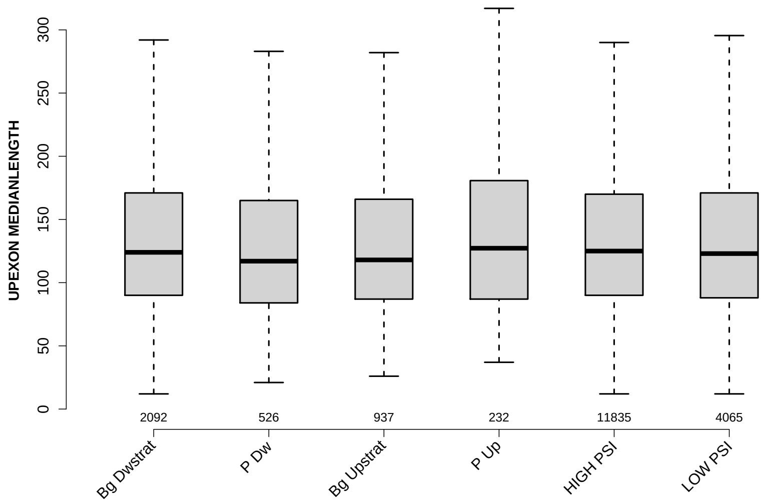

Significant results from Mann-Whitney U test:

- `Bg_Dwstrat` vs `P_Dw` : 0.0155546
  mean: 157.6554 > 144.2082 , median: 124 > 117
- `Bg_Dwstrat` vs `Bg_Upstrat` : 0.0477937
  mean: 157.6554 > 149.9226 , median: 124 > 118
- `P_Dw` vs `HIGH_PSI` : 0.00609642
  mean: 144.2082 < 159.4839 , median: 117 < 125
- `P_Dw` vs `LOW_PSI` : 0.0462092
  mean: 144.2082 < 157.607 , median: 117 < 123
- `Bg_Upstrat` vs `HIGH_PSI` : 0.015867
  mean: 149.9226 < 159.4839 , median: 118 < 125

#### 6.3 DOEXON MEDIANLENGTH

Back to: Overview | ToC

Meaning: median length of down-stream exon

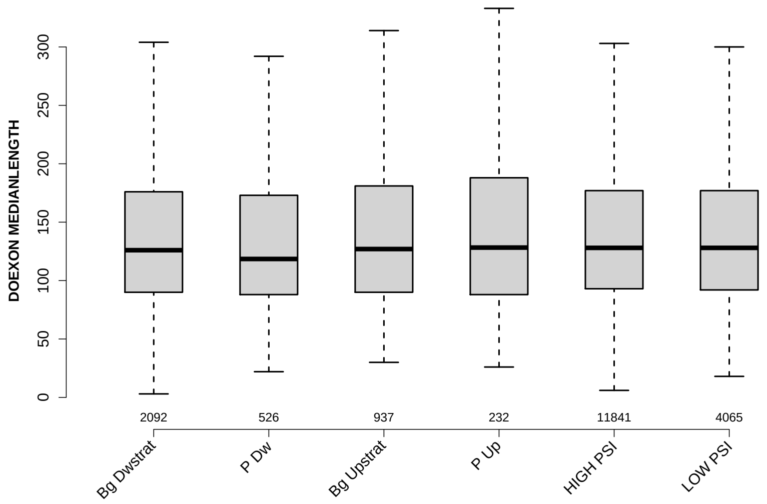

Significant results from Mann-Whitney U test:

- `P_Dw` vs `Bg_Upstrat` : 0.0430755
  mean: 248.8308 > 236.6862 , median: 118.5 < 127
- `P_Dw` vs `HIGH_PSI` : 0.016745
  mean: 248.8308 < 256.9639 , median: 118.5 < 128
- `P_Dw` vs `LOW_PSI` : 0.0184861
  mean: 248.8308 > 242.2478 , median: 118.5 < 128

#### 6.4 RATIO UPEXON EXON LENGTH

Back to: Overview | ToC

Meaning: median up-stream exon length / exon length

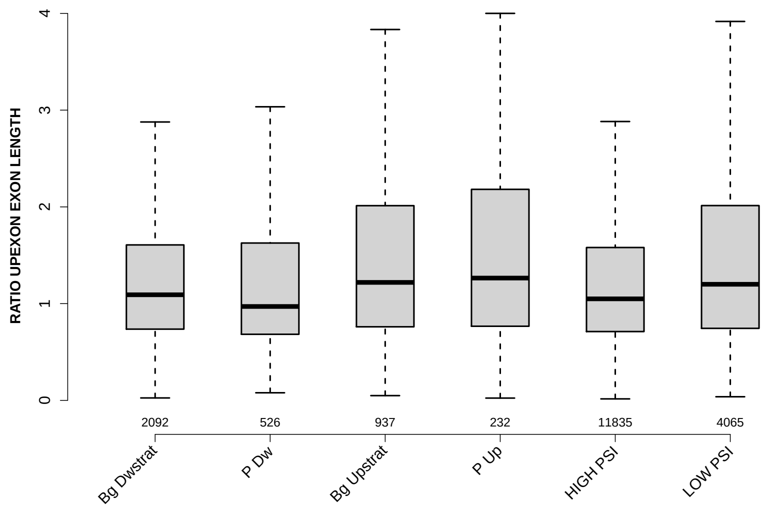

Significant results from Mann-Whitney U test:

- Bg_Dwstrat vs P_Dw : 0.0244612
  mean: 1.4319 < 1.491 , median: 1.0916 > 0.970588
- Bg_Dwstrat vs Bg_Upstrat : 2.00652e-05
  mean: 1.4319 < 2.156 , median: 1.0916 < 1.2194
- Bg_Dwstrat vs P_Up : 0.00125606
  mean: 1.4319 < 2.0866 , median: 1.0916 < 1.2642
- Bg_Dwstrat vs HIGH_PSI : 0.0339988
  mean: 1.4319 < 1.4332 , median: 1.0916 > 1.0495
- Bg_Dwstrat vs LOW_PSI : 6.04415e-09
  mean: 1.4319 < 2.0583 , median: 1.0916 < 1.2
- P_Dw vs Bg_Upstrat : 2.15202e-06
  mean: 1.491 < 2.156 , median: 0.970588 < 1.2194
- P_Dw vs P_Up : 9.38688e-05
  mean: 1.491 < 2.0866 , median: 0.970588 < 1.2642
- P_Dw vs LOW_PSI : 9.86414e-08
  mean: 1.491 < 2.0583 , median: 0.970588 < 1.2
- Bg_Upstrat vs HIGH_PSI : 4.58181e-10
  mean: 2.156 > 1.4332 , median: 1.2194 > 1.0495
- P_Up vs HIGH_PSI : 6.53531e-05
  mean: 2.0866 > 1.4332 , median: 1.2642 > 1.0495

- `HIGH_PSI` vs `LOW_PSI` : 3.02038e-28
  mean: 1.4332 < 2.0583 , median: 1.0495 < 1.2

#### 6.5 RATIO DOEXON EXON LENGTH

Back to: Overview | ToC

Meaning: median down-stream exon length / exon length

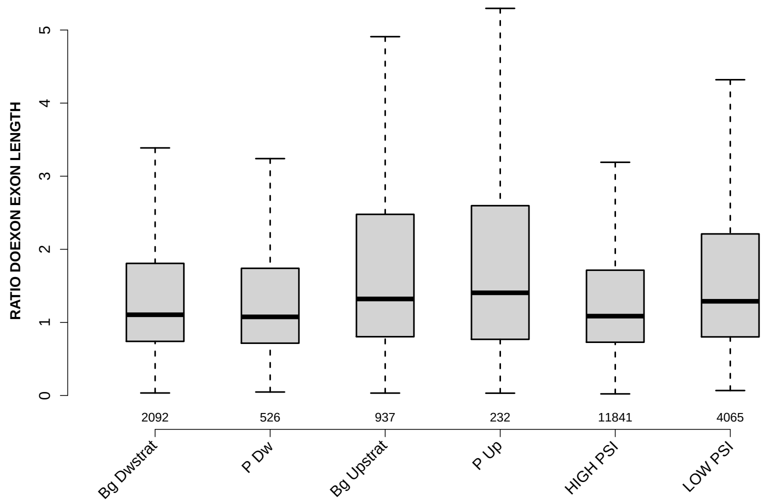

Significant results from Mann-Whitney U test:

- Bg_Dwstrat vs Bg_Upstrat : 1.15199e-07
  mean: 2.3076 < 4.0177 , median: 1.1045 < 1.3208
- Bg_Dwstrat vs P_Up : 0.00130935
  mean: 2.3076 < 3.4238 , median: 1.1045 < 1.4048
- Bg_Dwstrat vs LOW_PSI : 2.18811e-09
  mean: 2.3076 < 3.4596 , median: 1.1045 < 1.2887
- P_Dw vs Bg_Upstrat : 6.08649e-06
  mean: 2.1962 < 4.0177 , median: 1.0755 < 1.3208
- P_Dw vs P_Up : 0.00117648
  mean: 2.1962 < 3.4238 , median: 1.0755 < 1.4048
- P_Dw vs LOW_PSI : 1.14008e-05
  mean: 2.1962 < 3.4596 , median: 1.0755 < 1.2887
- Bg_Upstrat vs HIGH_PSI : 3.24385e-13
  mean: 4.0177 > 2.2864 , median: 1.3208 > 1.0857
- P_Up vs HIGH_PSI : 9.64806e-05
  mean: 3.4238 > 2.2864 , median: 1.4048 > 1.0857
- HIGH_PSI vs LOW_PSI : 8.21358e-29
  mean: 2.2864 < 3.4596 , median: 1.0857 < 1.2887

#### 6.6 UPINTRON MEDIANLENGTH

Back to: Overview | ToC

Meaning: median length of up-stream introns

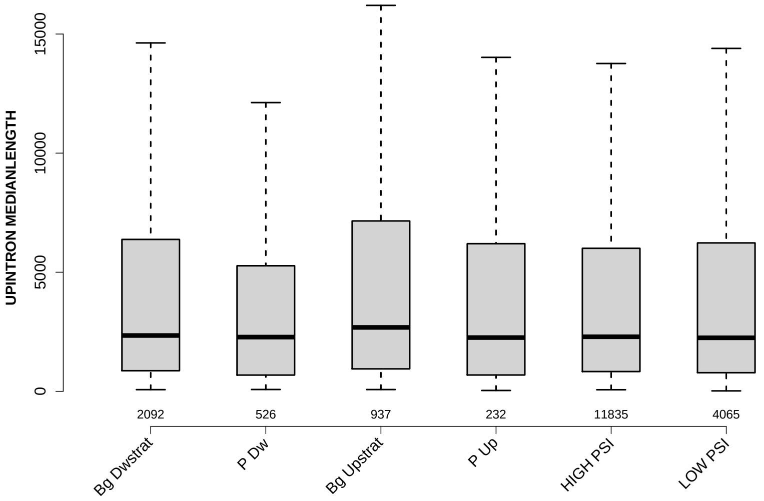

Significant results from Mann-Whitney U test:

- `Bg_Dwstrat` vs `Bg_Upstrat` : 0.0154362
  mean: 7830.1456 < 8599.8431 , median: 2344 < 2684
- `P_Dw` vs `Bg_Upstrat` : 0.00176615
  mean: 5221.634 < 8599.8431 , median: 2277 < 2684
- `Bg_Upstrat` vs `P_Up` : 0.0428827
  mean: 8599.8431 > 6275.9073 , median: 2684 > 2256.5
- `Bg_Upstrat` vs `HIGH_PSI` : 0.00137953
  mean: 8599.8431 > 7546.014 , median: 2684 > 2291
- `Bg_Upstrat` vs `LOW_PSI` : 0.00117753
  mean: 8599.8431 > 7078.3458 , median: 2684 > 2248

#### 6.7 DOINTRON MEDIANLENGTH

Back to: Overview | ToC

Meaning: median length of down-stream introns

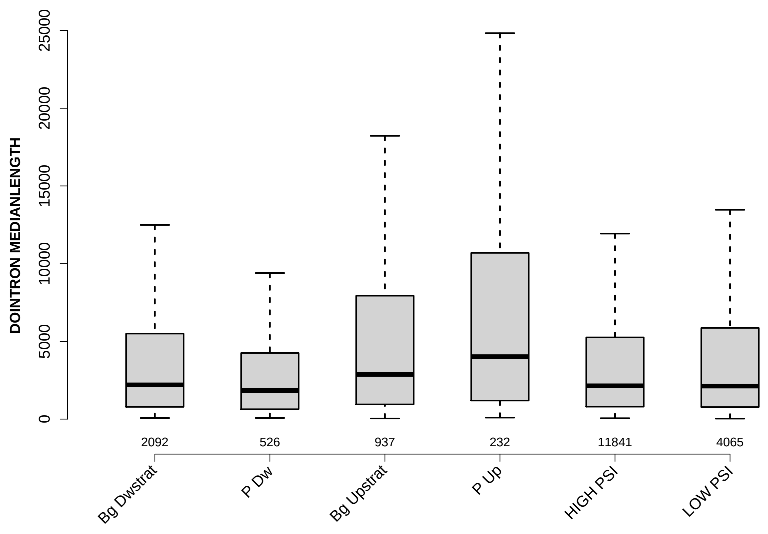

Significant results from Mann-Whitney U test:

- Bg_Dwstrat vs P_Dw : 0.000626304
  mean: 6458.6489 > 4113.4373 , median: 2201 > 1841
- Bg_Dwstrat vs Bg_Upstrat : 1.23586e-05
  mean: 6458.6489 < 8904.7412 , median: 2201 < 2875
- Bg_Dwstrat vs P_Up : 1.81377e-06
  mean: 6458.6489 < 9425.2047 , median: 2201 < 4018.5
- P_Dw vs Bg_Upstrat : 4.17756e-10
  mean: 4113.4373 < 8904.7412 , median: 1841 < 2875
- P_Dw vs P_Up : 2.41554e-10
  mean: 4113.4373 < 9425.2047 , median: 1841 < 4018.5
- P_Dw vs HIGH_PSI : 0.000976207
  mean: 4113.4373 < 6065.1019 , median: 1841 < 2142
- P_Dw vs LOW_PSI : 0.000221386
  mean: 4113.4373 < 6992.2823 , median: 1841 < 2126
- Bg_Upstrat vs P_Up : 0.033434
  mean: 8904.7412 < 9425.2047 , median: 2875 < 4018.5
- Bg_Upstrat vs HIGH_PSI : 1.48993e-08
  mean: 8904.7412 > 6065.1019 , median: 2875 > 2142
- Bg_Upstrat vs LOW_PSI : 1.27888e-05
  mean: 8904.7412 > 6992.2823 , median: 2875 > 2126

- P_Up vs HIGH_PSI : 1.05501e-07
  mean: 9425.2047 > 6065.1019 , median: 4018.5 > 2142
- P_Up vs LOW_PSI : 4.89761e-06
  mean: 9425.2047 > 6992.2823 , median: 4018.5 > 2126

#### 6.8 RATIO UPINTRON EXON LENGTH

Back to: Overview | ToC

Meaning: median up-stream intron length / exon length

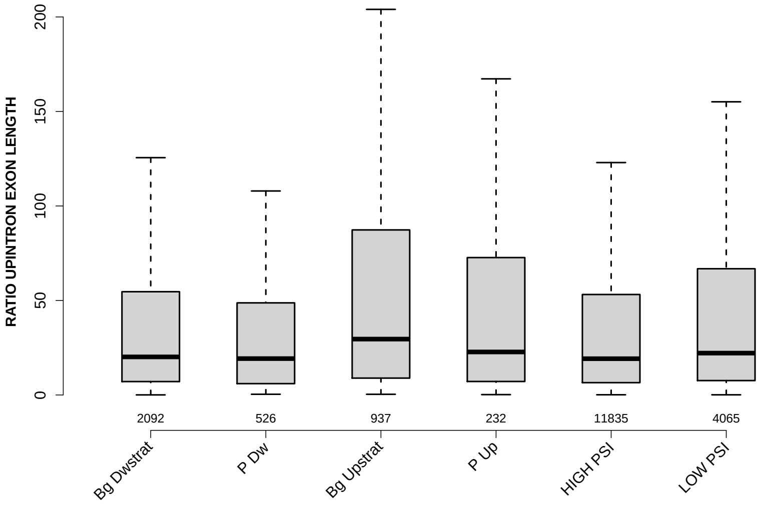

Significant results from Mann-Whitney U test:

- `Bg_Dwstrat` vs `Bg_Upstrat` : 9.84489e-08
  mean: 75.4137 < 106.4315 , median: 20.1437 < 29.5638
- `Bg_Dwstrat` vs `LOW_PSI` : 0.00693228
  mean: 75.4137 < 89.7353 , median: 20.1437 < 22.152
- `P_Dw` vs `Bg_Upstrat` : 3.29376e-07
  mean: 55.7631 < 106.4315 , median: 19.2481 < 29.5638
- `P_Dw` vs `LOW_PSI` : 0.00214903
  mean: 55.7631 < 89.7353 , median: 19.2481 < 22.152
- `Bg_Upstrat` vs `HIGH_PSI` : 2.35202e-12
  mean: 106.4315 > 69.3002 , median: 29.5638 > 19.1942
- `Bg_Upstrat` vs `LOW_PSI` : 0.000163006
  mean: 106.4315 > 89.7353 , median: 29.5638 > 22.152
- `HIGH_PSI` vs `LOW_PSI` : 1.87978e-08
  mean: 69.3002 < 89.7353 , median: 19.1942 < 22.152

#### 6.9 RATIO DOINTRON EXON LENGTH

Back to: Overview | ToC

Meaning: median down-stream intron length / exon length

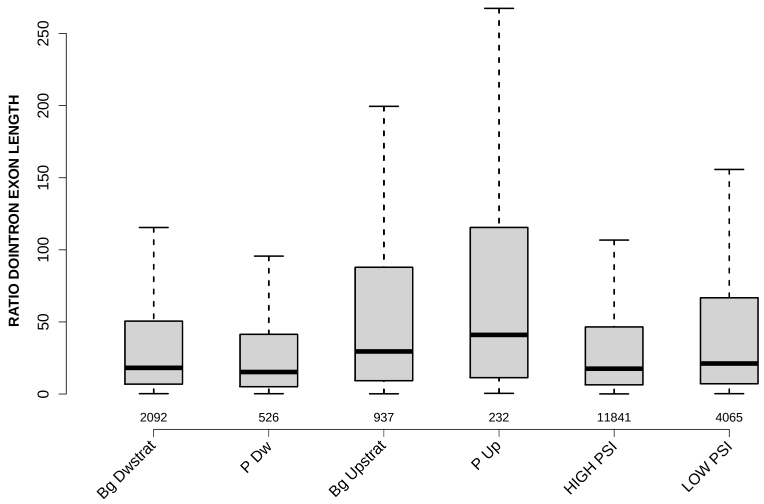

Significant results from Mann-Whitney U test:

- Bg_Dwstrat vs P_Dw : 0.00367679
  mean: 62.4944 > 46.0234 , median: 18.1649 > 15.2712
- Bg_Dwstrat vs Bg_Upstrat : 1.06103e-12
  mean: 62.4944 < 114.544 , median: 18.1649 < 29.5238
- Bg_Dwstrat vs P_Up : 1.15637e-08
  mean: 62.4944 < 122.5259 , median: 18.1649 < 41.0051
- Bg_Dwstrat vs LOW_PSI : 9.3933e-05
  mean: 62.4944 < 94.5224 , median: 18.1649 < 21.1702
- P_Dw vs Bg_Upstrat : 1.36838e-14
  mean: 46.0234 < 114.544 , median: 15.2712 < 29.5238
- P_Dw vs P_Up : 1.58671e-11
  mean: 46.0234 < 122.5259 , median: 15.2712 < 41.0051
- P_Dw vs HIGH_PSI : 0.0148633
  mean: 46.0234 < 56.7889 , median: 15.2712 < 17.5762
- P_Dw vs LOW_PSI : 1.39017e-07
  mean: 46.0234 < 94.5224 , median: 15.2712 < 21.1702
- Bg_Upstrat vs HIGH_PSI : 1.49839e-20
  mean: 114.544 > 56.7889 , median: 29.5238 > 17.5762
- Bg_Upstrat vs LOW_PSI : 3.49212e-06
  mean: 114.544 > 94.5224 , median: 29.5238 > 21.1702

- `P_Up` vs `HIGH_PSI` : 8.73714e-11
  mean: $122.5259 > 56.7889$ , median: $41.0051 > 17.5762$
- `P_Up` vs `LOW_PSI` : 2.99123e-05
  mean: $122.5259 > 94.5224$ , median: $41.0051 > 21.1702$
- `HIGH_PSI` vs `LOW_PSI` : 2.6134e-14
  mean: $56.7889 < 94.5224$ , median: $17.5762 < 21.1702$

#### 6.10 EXON GCC

Back to: Overview | ToC

Meaning: GC content of entire exon sequence

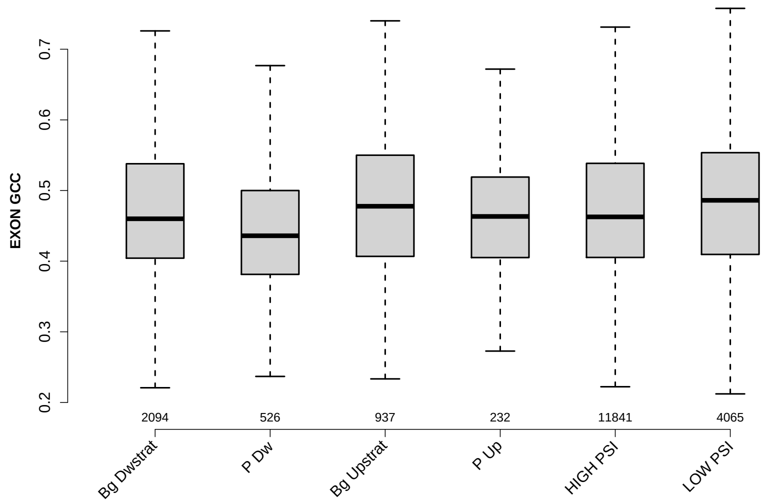

Significant results from Mann-Whitney U test:

- Bg_Dwstrat vs P_Dw : 2.29095e-09
  mean: 0.473315 > 0.446595 , median: 0.46 > 0.435962
- Bg_Dwstrat vs LOW_PSI : 0.000119758
  mean: 0.473315 < 0.478917 , median: 0.46 < 0.486111
- P_Dw vs Bg_Upstrat : 4.72506e-11
  mean: 0.446595 < 0.475821 , median: 0.435962 < 0.477717
- P_Dw vs P_Up : 0.000906747
  mean: 0.446595 < 0.463566 , median: 0.435962 < 0.463302
- P_Dw vs HIGH_PSI : 4.16541e-12
  mean: 0.446595 < 0.473651 , median: 0.435962 < 0.462687
- P_Dw vs LOW_PSI : 6.08139e-17
  mean: 0.446595 < 0.478917 , median: 0.435962 < 0.486111
- Bg_Upstrat vs P_Up : 0.0349417
  mean: 0.475821 > 0.463566 , median: 0.477717 > 0.463302
- P_Up vs LOW_PSI : 0.00303638
  mean: 0.463566 < 0.478917 , median: 0.463302 < 0.486111
- HIGH_PSI vs LOW_PSI : 8.33522e-08
  mean: 0.473651 < 0.478917 , median: 0.462687 < 0.486111

#### 6.11 UPINTRON GCC

Back to: Overview | ToC

Meaning: GC content of entire up-stream intron sequence

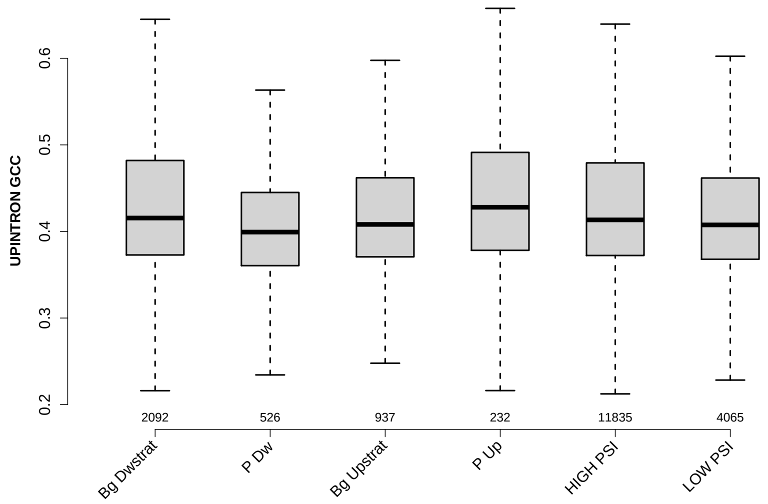

Significant results from Mann-Whitney U test:

- `Bg_Dwstrat` vs `P_Dw` : 1.22492e-08
  mean: 0.434658 > 0.406972 , median: 0.415517 > 0.39927
- `Bg_Dwstrat` vs `Bg_Upstrat` : 0.00305207
  mean: 0.434658 > 0.423061 , median: 0.415517 > 0.408115
- `Bg_Dwstrat` vs `LOW_PSI` : 8.34497e-07
  mean: 0.434658 > 0.420695 , median: 0.415517 > 0.407568
- `P_Dw` vs `Bg_Upstrat` : 0.00220971
  mean: 0.406972 < 0.423061 , median: 0.39927 < 0.408115
- `P_Dw` vs `P_Up` : 2.22931e-06
  mean: 0.406972 < 0.442645 , median: 0.39927 < 0.427952
- `P_Dw` vs `HIGH_PSI` : 6.18628e-09
  mean: 0.406972 < 0.43335 , median: 0.39927 < 0.413381
- `P_Dw` vs `LOW_PSI` : 0.00151407
  mean: 0.406972 < 0.420695 , median: 0.39927 < 0.407568
- `Bg_Upstrat` vs `P_Up` : 0.00393024
  mean: 0.423061 < 0.442645 , median: 0.408115 < 0.427952
- `Bg_Upstrat` vs `HIGH_PSI` : 0.00385299
  mean: 0.423061 < 0.43335 , median: 0.408115 < 0.413381
- `P_Up` vs `LOW_PSI` : 0.00073654
  mean: 0.442645 > 0.420695 , median: 0.427952 > 0.407568

- `HIGH_PSI` vs `LOW_PSI` : 2.478e-10
  mean: 0.43335 > 0.420695 , median: 0.413381 > 0.407568

#### 6.12 UPEXON GCC

Back to: Overview | ToC

Meaning: GC content of entire up-stream exon sequence

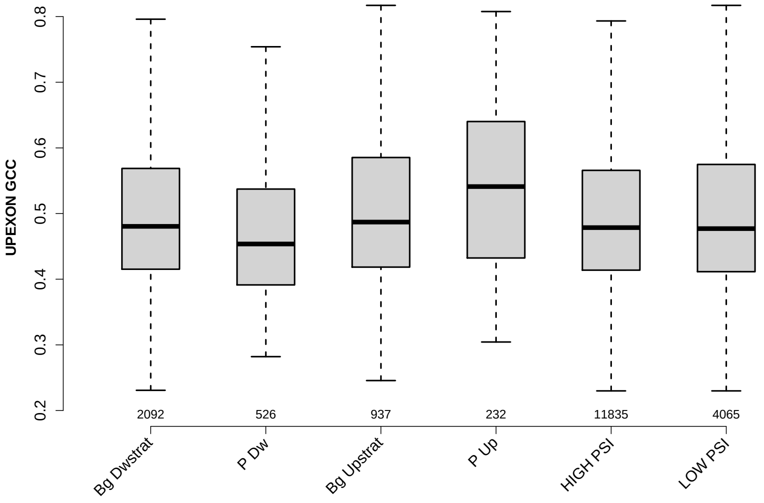

Significant results from Mann-Whitney U test:

- `Bg_Dwstrat` vs `P_Dw` : 7.67912e-07
  mean: 0.494879 > 0.470597 , median: 0.480487 > 0.45364
- `Bg_Dwstrat` vs `Bg_Upstrat` : 0.0330922
  mean: 0.494879 < 0.506272 , median: 0.480487 < 0.487006
- `Bg_Dwstrat` vs `P_Up` : 4.40419e-07
  mean: 0.494879 < 0.539031 , median: 0.480487 < 0.541099
- `P_Dw` vs `Bg_Upstrat` : 5.51916e-09
  mean: 0.470597 < 0.506272 , median: 0.45364 < 0.487006
- `P_Dw` vs `P_Up` : 3.07105e-12
  mean: 0.470597 < 0.539031 , median: 0.45364 < 0.541099
- `P_Dw` vs `HIGH_PSI` : 2.42096e-07
  mean: 0.470597 < 0.493304 , median: 0.45364 < 0.478571
- `P_Dw` vs `LOW_PSI` : 1.71736e-07
  mean: 0.470597 < 0.498352 , median: 0.45364 < 0.477046
- `Bg_Upstrat` vs `P_Up` : 0.000434501
  mean: 0.506272 < 0.539031 , median: 0.487006 < 0.541099
- `Bg_Upstrat` vs `HIGH_PSI` : 0.00476105
  mean: 0.506272 > 0.493304 , median: 0.487006 > 0.478571
- `Bg_Upstrat` vs `LOW_PSI` : 0.0408349
  mean: 0.506272 > 0.498352 , median: 0.487006 > 0.477046

- `P_Up` vs `HIGH_PSI` : 3.93187e-08
  mean: 0.539031 > 0.493304 , median: 0.541099 > 0.478571
- `P_Up` vs `LOW_PSI` : 1.40597e-06
  mean: 0.539031 > 0.498352 , median: 0.541099 > 0.477046

#### 6.13 DOINTRON GCC

Back to: Overview | ToC

Meaning: GC content of entire down-stream intron sequence

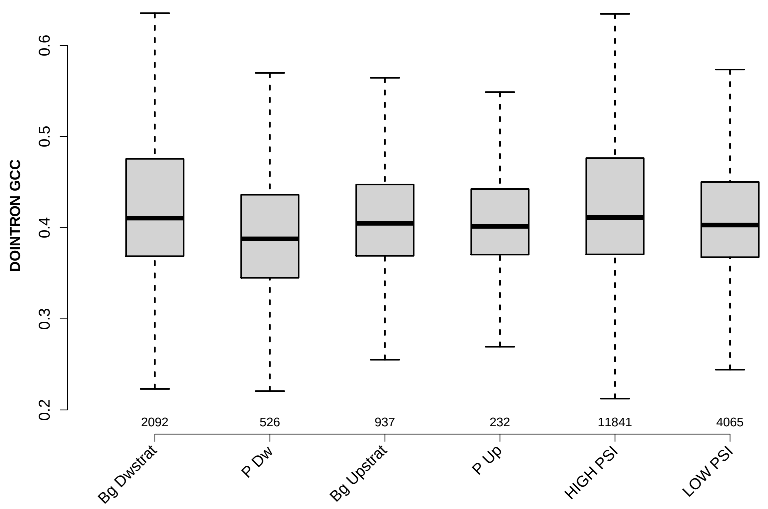

Significant results from Mann-Whitney U test:

- Bg_Dwstrat vs P_Dw : 1.46806e-13
  mean: 0.428827 > 0.395518 , median: 0.410523 > 0.387706
- Bg_Dwstrat vs Bg_Upstrat : 0.00179893
  mean: 0.428827 > 0.413933 , median: 0.410523 > 0.404762
- Bg_Dwstrat vs P_Up : 0.0300803
  mean: 0.428827 > 0.411404 , median: 0.410523 > 0.401428
- Bg_Dwstrat vs LOW_PSI : 5.04169e-06
  mean: 0.428827 > 0.414825 , median: 0.410523 > 0.402908
- P_Dw vs Bg_Upstrat : 2.05361e-06
  mean: 0.395518 < 0.413933 , median: 0.387706 < 0.404762
- P_Dw vs P_Up : 0.00355082
  mean: 0.395518 < 0.411404 , median: 0.387706 < 0.401428
- P_Dw vs HIGH_PSI : 2.52746e-17
  mean: 0.395518 < 0.430754 , median: 0.387706 < 0.41115
- P_Dw vs LOW_PSI : 4.49174e-08
  mean: 0.395518 < 0.414825 , median: 0.387706 < 0.402908
- Bg_Upstrat vs HIGH_PSI : 2.86218e-05
  mean: 0.413933 < 0.430754 , median: 0.404762 < 0.41115
- P_Up vs HIGH_PSI : 0.0100977
  mean: 0.411404 < 0.430754 , median: 0.401428 < 0.41115

- `HIGH_PSI` vs `LOW_PSI` : 4.36602e-15
  mean: 0.430754 > 0.414825 , median: 0.41115 > 0.402908

#### 6.14 DOEXON GCC

Back to: Overview | ToC

Meaning: GC content of entire down-stream exon sequence

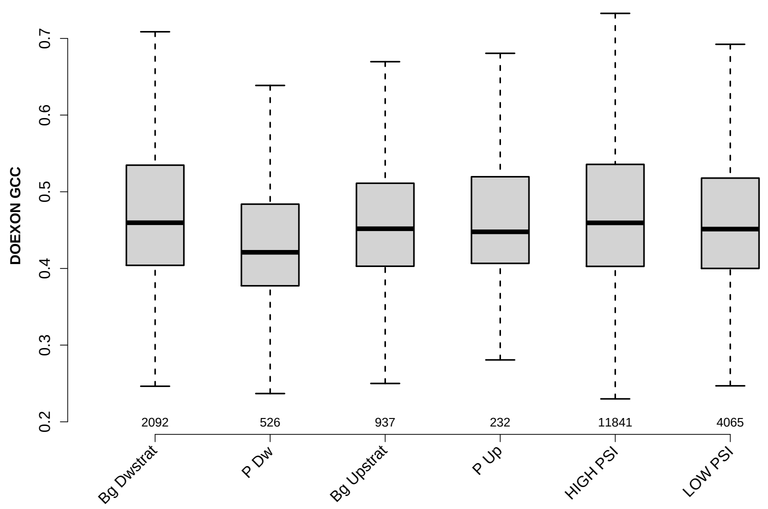

Significant results from Mann-Whitney U test:

- `Bg_Dwstrat` vs `P_Dw` : 3.93724e-17
  mean: 0.470431 > 0.434284 , median: 0.459615 > 0.421053
- `Bg_Dwstrat` vs `Bg_Upstrat` : 0.00909812
  mean: 0.470431 > 0.459761 , median: 0.459615 > 0.451777
- `Bg_Dwstrat` vs `LOW_PSI` : 0.000453389
  mean: 0.470431 > 0.461668 , median: 0.459615 > 0.451327
- `P_Dw` vs `Bg_Upstrat` : 1.33576e-09
  mean: 0.434284 < 0.459761 , median: 0.421053 < 0.451777
- `P_Dw` vs `P_Up` : 1.9657e-05
  mean: 0.434284 < 0.460427 , median: 0.421053 < 0.447768
- `P_Dw` vs `HIGH_PSI` : 5.95084e-20
  mean: 0.434284 < 0.470424 , median: 0.421053 < 0.459459
- `P_Dw` vs `LOW_PSI` : 1.19581e-12
  mean: 0.434284 < 0.461668 , median: 0.421053 < 0.451327
- `Bg_Upstrat` vs `HIGH_PSI` : 0.00312275
  mean: 0.459761 < 0.470424 , median: 0.451777 < 0.459459
- `HIGH_PSI` vs `LOW_PSI` : 3.83483e-07
  mean: 0.470424 > 0.461668 , median: 0.459459 > 0.451327

#### 6.15 RATIO UPEXON EXON GCC

Back to: Overview | ToC

Meaning: UPEXON GCC / EXON GCC

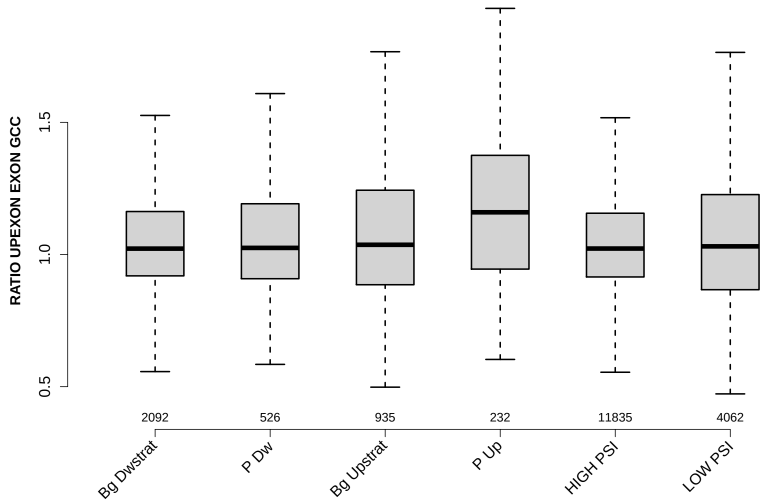

Significant results from Mann-Whitney U test:

- `Bg_Dwstrat` vs `P_Up` : 4.5193e-09
  mean: 1.0651 < 1.1885 , median: 1.0223 < 1.1597
- `P_Dw` vs `P_Up` : 2.41421e-06
  mean: 1.0772 < 1.1885 , median: 1.025 < 1.1597
- `Bg_Upstrat` vs `P_Up` : 7.96986e-06
  mean: 1.0932 < 1.1885 , median: 1.0368 < 1.1597
- `P_Up` vs `HIGH_PSI` : 1.31032e-10
  mean: 1.1885 > 1.0593 , median: 1.1597 > 1.0225
- `P_Up` vs `LOW_PSI` : 8.10441e-09
  mean: 1.1885 > 1.0707 , median: 1.1597 > 1.0308

#### 6.16 RATIO UPINTRON EXON GCC

Back to: Overview | ToC

Meaning: UPINTRON GCC / EXON GCC

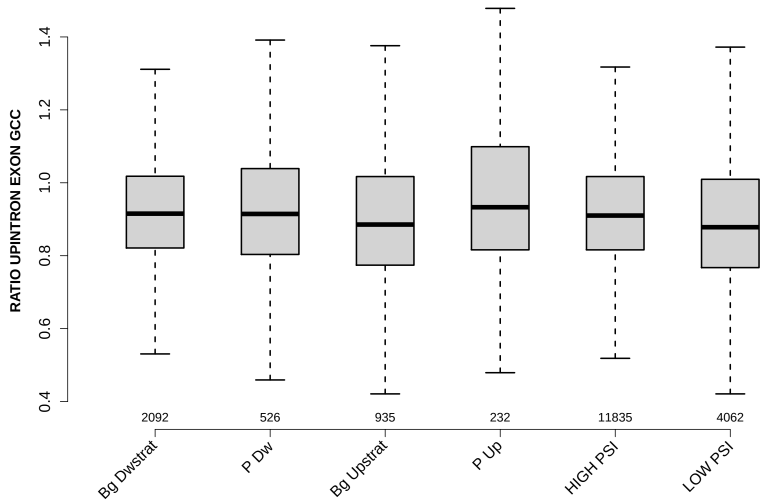

Significant results from Mann-Whitney U test:

- Bg_Dwstrat vs Bg_Upstrat : 2.03581e-05
  mean: 0.930372 > 0.910414 , median: 0.915358 > 0.885429
- Bg_Dwstrat vs P_Up : 0.0079079
  mean: 0.930372 < 0.97067 , median: 0.915358 < 0.932925
- Bg_Dwstrat vs LOW_PSI : 4.57466e-16
  mean: 0.930372 > 0.899451 , median: 0.915358 > 0.878145
- P_Dw vs Bg_Upstrat : 0.00612003
  mean: 0.928069 > 0.910414 , median: 0.914524 > 0.885429
- P_Dw vs P_Up : 0.0182768
  mean: 0.928069 < 0.97067 , median: 0.914524 < 0.932925
- P_Dw vs LOW_PSI : 2.3673e-05
  mean: 0.928069 > 0.899451 , median: 0.914524 > 0.878145
- Bg_Upstrat vs P_Up : 1.58615e-05
  mean: 0.910414 < 0.97067 , median: 0.885429 < 0.932925
- Bg_Upstrat vs HIGH_PSI : 1.88976e-05
  mean: 0.910414 < 0.925148 , median: 0.885429 < 0.910107
- P_Up vs HIGH_PSI : 0.00186689
  mean: 0.97067 > 0.925148 , median: 0.932925 > 0.910107
- P_Up vs LOW_PSI : 9.00261e-08
  mean: 0.97067 > 0.899451 , median: 0.932925 > 0.878145

- `HIGH_PSI` vs `LOW_PSI` : 5.86882e-27
  mean: 0.925148 > 0.899451 , median: 0.910107 > 0.878145

#### 6.17 RATIO DOINTRON EXON GCC

Back to: Overview | ToC

Meaning: DOINTRON GCC / EXON GCC

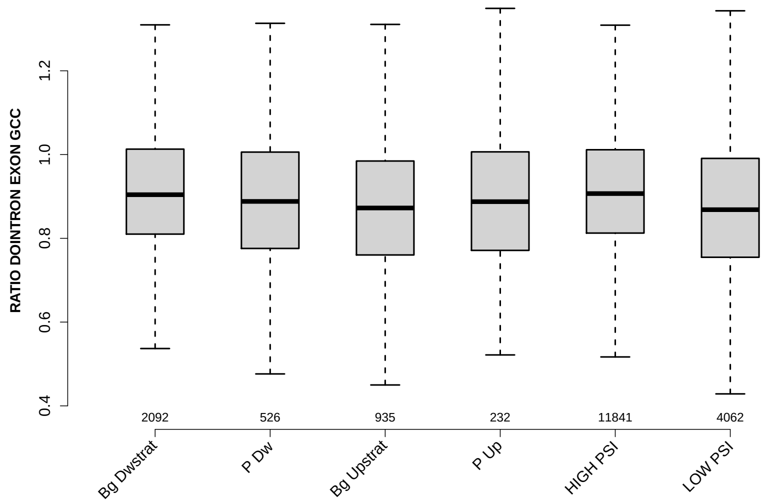

Significant results from Mann-Whitney U test:

- `Bg_Dwstrat` vs `P_Dw` : 0.0200385
  mean: 0.918227 > 0.902744 , median: 0.904153 > 0.888113
- `Bg_Dwstrat` vs `Bg_Upstrat` : 7.36119e-08
  mean: 0.918227 > 0.889576 , median: 0.904153 > 0.872424
- `Bg_Dwstrat` vs `LOW_PSI` : 6.7841e-16
  mean: 0.918227 > 0.887036 , median: 0.904153 > 0.868362
- `P_Dw` vs `HIGH_PSI` : 0.00411919
  mean: 0.902744 < 0.919539 , median: 0.888113 < 0.906832
- `P_Dw` vs `LOW_PSI` : 0.0282417
  mean: 0.902744 > 0.887036 , median: 0.888113 > 0.868362
- `Bg_Upstrat` vs `HIGH_PSI` : 2.48565e-11
  mean: 0.889576 < 0.919539 , median: 0.872424 < 0.906832
- `HIGH_PSI` vs `LOW_PSI` : 3.07029e-37
  mean: 0.919539 > 0.887036 , median: 0.906832 > 0.868362

#### 6.18 RATIO DOEXON EXON GCC

Back to: Overview | ToC

Meaning: DOEXON GCC / EXON GCC

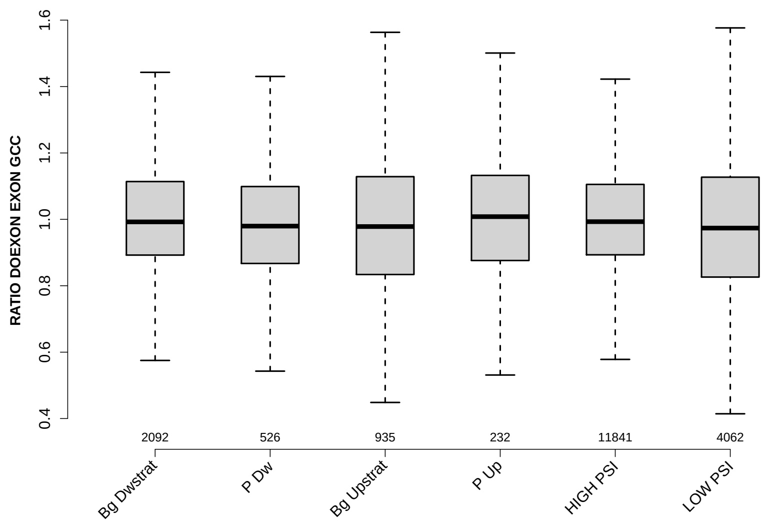

Significant results from Mann-Whitney U test:

- `Bg_Dwstrat` vs `P_Dw` : 0.0408523
  mean: 1.0118 > 0.991822 , median: 0.992351 > 0.979641
- `Bg_Dwstrat` vs `Bg_Upstrat` : 0.00230633
  mean: 1.0118 > 0.994632 , median: 0.992351 > 0.978318
- `Bg_Dwstrat` vs `LOW_PSI` : 1.06254e-06
  mean: 1.0118 > 0.993107 , median: 0.992351 > 0.973687
- `P_Dw` vs `HIGH_PSI` : 0.0453911
  mean: 0.991822 < 1.008 , median: 0.979641 < 0.993012
- `Bg_Upstrat` vs `HIGH_PSI` : 0.00106141
  mean: 0.994632 < 1.008 , median: 0.978318 < 0.993012
- `P_Up` vs `LOW_PSI` : 0.0490429
  mean: 1.0136 > 0.993107 , median: 1.0081 > 0.973687
- `HIGH_PSI` vs `LOW_PSI` : 1.78678e-11
  mean: 1.008 > 0.993107 , median: 0.993012 > 0.973687

##### 6.19 SF1 HIGHESTSCORE 3SS UPINTRON

Back to: Overview | ToC

Meaning: highest score of a SF1 positon weight matrix trained with human data in the last 150 nt 3 prime intron positons of up-stream intron

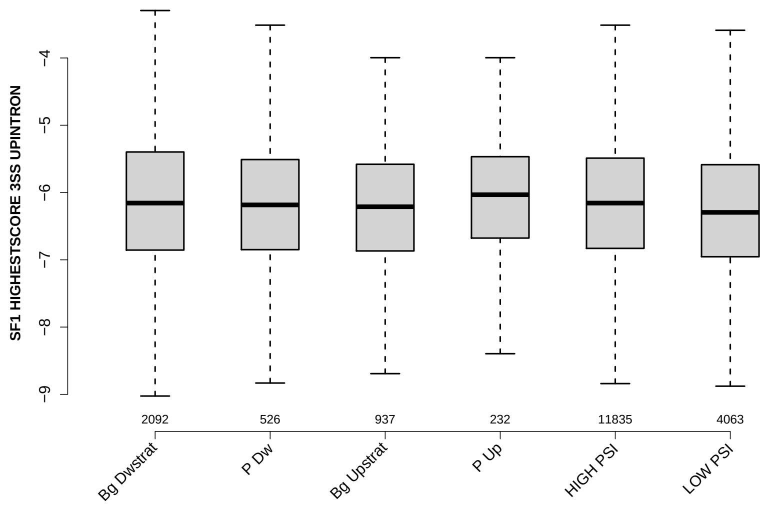

Significant results from Mann-Whitney U test:

- `Bg_Dwstrat` vs `LOW_PSI` : 5.62514e-05
  mean: -6.14092 > -6.23278 , median: -6.15674 > -6.29538
- `P_Up` vs `LOW_PSI` : 0.00272087
  mean: -6.05467 > -6.23278 , median: -6.03223 > -6.29538
- `HIGH_PSI` vs `LOW_PSI` : 2.23916e-08
  mean: -6.14789 > -6.23278 , median: -6.15674 > -6.29538

#### 6.20 SF1 HIGHESTSCORE 3SS DOINTRON

Back to: Overview | ToC

Meaning: highest score of a SF1 positon weight matrix trained with human data in the last 150 nt 3 prime intron positons of down-stream intron

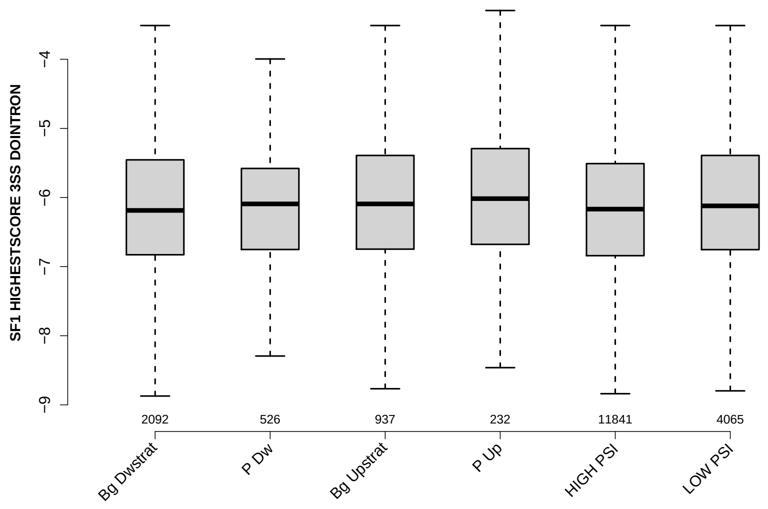

Significant results from Mann-Whitney U test:

- Bg_Dwstrat vs Bg_Upstrat : 0.0439411
  mean: -6.14534 < -6.06197 , median: -6.18725 < -6.09302
- Bg_Dwstrat vs P_Up : 0.00504558
  mean: -6.14534 < -5.93242 , median: -6.18725 < -6.01665
- Bg_Dwstrat vs LOW_PSI : 0.023972
  mean: -6.14534 < -6.08497 , median: -6.18725 < -6.12133
- P_Dw vs P_Up : 0.0336393
  mean: -6.10638 < -5.93242 , median: -6.09302 < -6.01665
- Bg_Upstrat vs HIGH_PSI : 0.0155456
  mean: -6.06197 > -6.15302 , median: -6.09302 > -6.1683
- P_Up vs HIGH_PSI : 0.00287
  mean: -5.93242 > -6.15302 , median: -6.01665 > -6.1683
- P_Up vs LOW_PSI : 0.0413543
  mean: -5.93242 > -6.08497 , median: -6.01665 > -6.12133
- HIGH_PSI vs LOW_PSI : 0.000400909
  mean: -6.15302 < -6.08497 , median: -6.1683 < -6.12133

#### 6.21 UP 5SS 20INT10EX GCC

Back to: Overview | ToC

Meaning: GC content of up-stream 5ss sequence (20int+10ex positions)

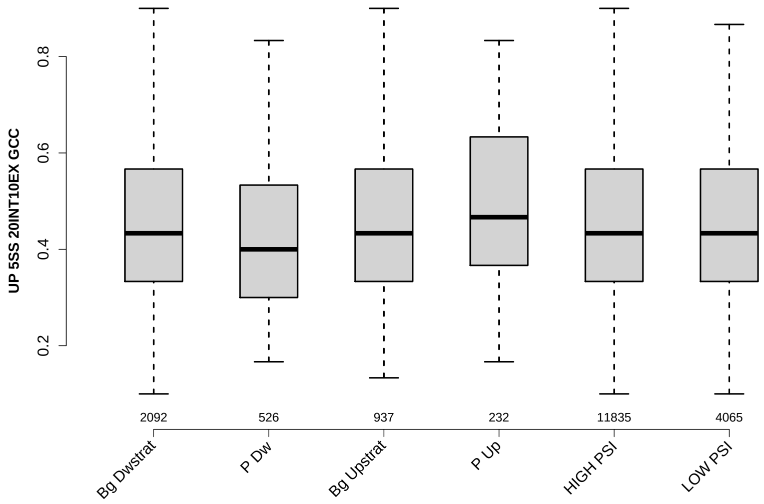

Significant results from Mann-Whitney U test:

- `Bg_Dwstrat` vs `P_Dw` : 4.11659e-08
  mean: 0.459544 > 0.4218 , median: 0.433333 > 0.4
- `Bg_Dwstrat` vs `P_Up` : 0.002034
  mean: 0.459544 < 0.491092 , median: 0.433333 < 0.466667
- `P_Dw` vs `Bg_Upstrat` : 9.886e-07
  mean: 0.4218 < 0.460263 , median: 0.4 < 0.433333
- `P_Dw` vs `P_Up` : 2.83303e-09
  mean: 0.4218 < 0.491092 , median: 0.4 < 0.466667
- `P_Dw` vs `HIGH_PSI` : 3.3081e-09
  mean: 0.4218 < 0.458328 , median: 0.4 < 0.433333
- `P_Dw` vs `LOW_PSI` : 5.86164e-08
  mean: 0.4218 < 0.458938 , median: 0.4 < 0.433333
- `Bg_Upstrat` vs `P_Up` : 0.00394957
  mean: 0.460263 < 0.491092 , median: 0.433333 < 0.466667
- `P_Up` vs `HIGH_PSI` : 0.000910911
  mean: 0.491092 > 0.458328 , median: 0.466667 > 0.433333
- `P_Up` vs `LOW_PSI` : 0.0011662
  mean: 0.491092 > 0.458938 , median: 0.466667 > 0.433333

#### 6.22 GCC 3SS 20INT10EX

Back to: Overview | ToC

Meaning: GC content of 3ss sequence (20int+10ex positions)

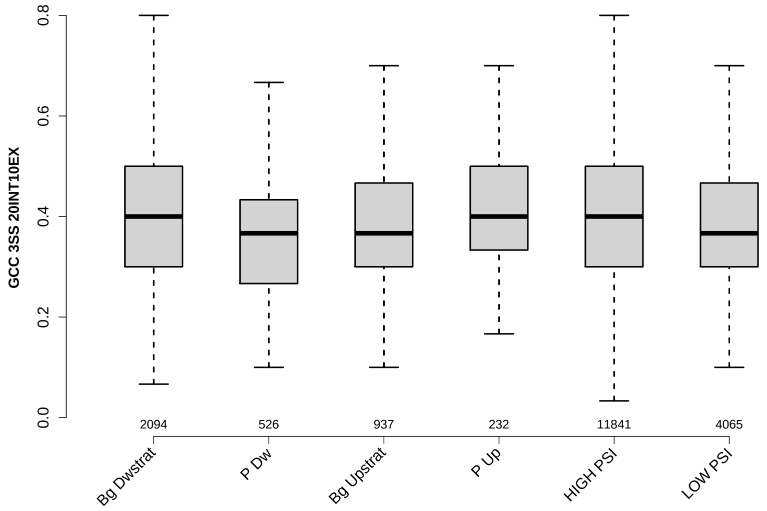

Significant results from Mann-Whitney U test:

- `Bg_Dwstrat` vs `P_Dw` : 3.06882e-11
  mean: 0.40979 > 0.367871 , median: 0.4 > 0.366667
- `Bg_Dwstrat` vs `Bg_Upstrat` : 0.014293
  mean: 0.40979 > 0.396585 , median: 0.4 > 0.366667
- `Bg_Dwstrat` vs `LOW_PSI` : 3.75487e-11
  mean: 0.40979 > 0.385691 , median: 0.4 > 0.366667
- `P_Dw` vs `Bg_Upstrat` : 7.23121e-06
  mean: 0.367871 < 0.396585 , median: 0.366667 = 0.366667
- `P_Dw` vs `P_Up` : 1.86019e-06
  mean: 0.367871 < 0.408764 , median: 0.366667 < 0.4
- `P_Dw` vs `HIGH_PSI` : 1.93908e-13
  mean: 0.367871 < 0.411122 , median: 0.366667 < 0.4
- `P_Dw` vs `LOW_PSI` : 0.000659861
  mean: 0.367871 < 0.385691 , median: 0.366667 = 0.366667
- `Bg_Upstrat` vs `HIGH_PSI` : 0.00315817
  mean: 0.396585 < 0.411122 , median: 0.366667 < 0.4
- `Bg_Upstrat` vs `LOW_PSI` : 0.0158561
  mean: 0.396585 > 0.385691 , median: 0.366667 = 0.366667
- `P_Up` vs `LOW_PSI` : 0.00142732
  mean: 0.408764 > 0.385691 , median: 0.4 > 0.366667

- `HIGH_PSI` vs `LOW_PSI` : 1.32567e-23
  mean: 0.411122 > 0.385691 , median: 0.4 > 0.366667

#### 6.23 GCC 5SS 20INT10EX

Back to: Overview | ToC

Meaning: GC content of 5ss sequence (20int+10ex positions)

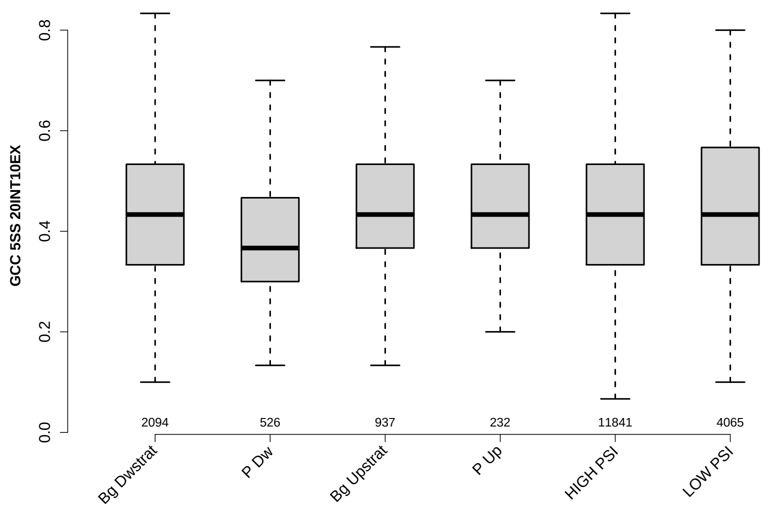

Significant results from Mann-Whitney U test:

- `Bg_Dwstrat` vs `P_Dw` : 8.50674e-15
  mean: 0.439303 > 0.391191 , median: 0.433333 > 0.366667
- `P_Dw` vs `Bg_Upstrat` : 3.8011e-15
  mean: 0.391191 < 0.441764 , median: 0.366667 < 0.433333
- `P_Dw` vs `P_Up` : 3.31907e-08
  mean: 0.391191 < 0.43592 , median: 0.366667 < 0.433333
- `P_Dw` vs `HIGH_PSI` : 2.39048e-18
  mean: 0.391191 < 0.441835 , median: 0.366667 < 0.433333
- `P_Dw` vs `LOW_PSI` : 3.09985e-19
  mean: 0.391191 < 0.442845 , median: 0.366667 < 0.433333

#### 6.24 DO 3SS 20INT10EX GCC

Back to: Overview | ToC

Meaning: GC content of down-stream 3ss sequence (20int+10ex positions)

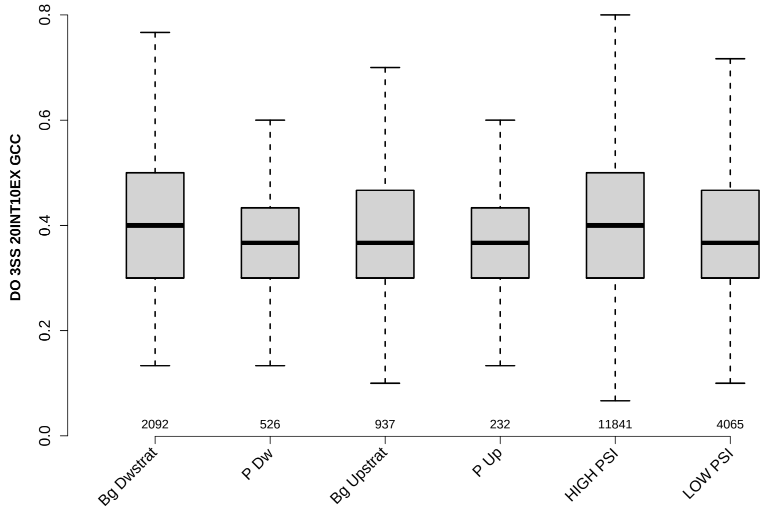

Significant results from Mann-Whitney U test:

- `Bg_Dwstrat` vs `P_Dw` : 2.48724e-10
  mean: 0.408556 > 0.368156 , median: 0.4 > 0.366667
- `Bg_Dwstrat` vs `Bg_Upstrat` : 3.6064e-05
  mean: 0.408556 > 0.385681 , median: 0.4 > 0.366667
- `Bg_Dwstrat` vs `P_Up` : 0.000438257
  mean: 0.408556 > 0.376652 , median: 0.4 > 0.366667
- `Bg_Dwstrat` vs `LOW_PSI` : 2.17121e-05
  mean: 0.408556 > 0.392792 , median: 0.4 > 0.366667
- `P_Dw` vs `Bg_Upstrat` : 0.00438982
  mean: 0.368156 < 0.385681 , median: 0.366667 = 0.366667
- `P_Dw` vs `HIGH_PSI` : 2.54217e-13
  mean: 0.368156 < 0.410495 , median: 0.366667 < 0.4
- `P_Dw` vs `LOW_PSI` : 1.92024e-05
  mean: 0.368156 < 0.392792 , median: 0.366667 = 0.366667
- `Bg_Upstrat` vs `HIGH_PSI` : 8.7574e-08
  mean: 0.385681 < 0.410495 , median: 0.366667 < 0.4
- `P_Up` vs `HIGH_PSI` : 9.31197e-05
  mean: 0.376652 < 0.410495 , median: 0.366667 < 0.4
- `HIGH_PSI` vs `LOW_PSI` : 3.10728e-13
  mean: 0.410495 > 0.392792 , median: 0.4 > 0.366667

#### 6.25 MAXENTSCR HSAMODEL UPSTRM 5SS

Back to: Overview | ToC

Meaning: maximum entropy score of 5ss of up-stream exon using a model trained with human splice sites

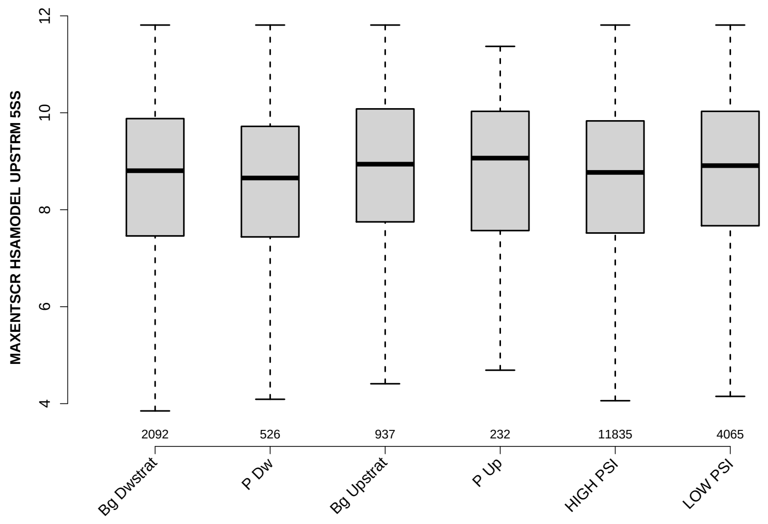

Significant results from Mann-Whitney U test:

- Bg_Dwstrat vs Bg_Upstrat : 0.000982193
  mean: 7.9521 < 8.4016 , median: 8.805 < 8.94
- Bg_Dwstrat vs LOW_PSI : 0.000497619
  mean: 7.9521 < 8.5968 , median: 8.805 < 8.91
- P_Dw vs Bg_Upstrat : 0.000179286
  mean: 8.1606 < 8.4016 , median: 8.655 < 8.94
- P_Dw vs P_Up : 0.0149779
  mean: 8.1606 < 8.6147 , median: 8.655 < 9.065
- P_Dw vs LOW_PSI : 0.000272825
  mean: 8.1606 < 8.5968 , median: 8.655 < 8.91
- Bg_Upstrat vs HIGH_PSI : 0.000281921
  mean: 8.4016 < 8.4164 , median: 8.94 > 8.77
- HIGH_PSI vs LOW_PSI : 1.6809e-06
  mean: 8.4164 < 8.5968 , median: 8.77 < 8.91

#### 6.26 MAXENTSCR HSAMODEL 3SS

Back to: Overview | ToC

Meaning: maximum entropy score of 3ss using a model trained with human splice sites

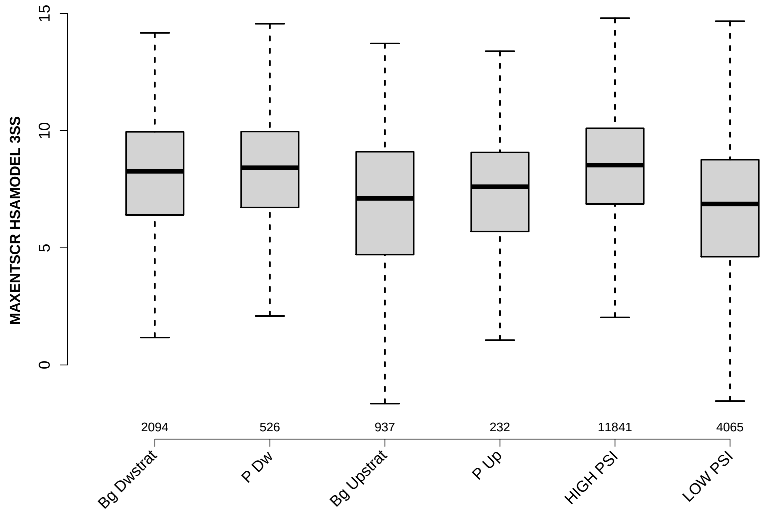

Significant results from Mann-Whitney U test:

- Bg_Dwstrat vs Bg_Upstrat : 5.85642e-20
  mean: 7.545 > 6.4599 , median: 8.265 > 7.11
- Bg_Dwstrat vs P_Up : 4.88841e-05
  mean: 7.545 > 7.0096 , median: 8.265 > 7.605
- Bg_Dwstrat vs HIGH_PSI : 1.71461e-07
  mean: 7.545 < 8.3636 , median: 8.265 < 8.53
- Bg_Dwstrat vs LOW_PSI : 2.78373e-65
  mean: 7.545 > 6.4194 , median: 8.265 > 6.87
- P_Dw vs Bg_Upstrat : 1.24067e-15
  mean: 8.0902 > 6.4599 , median: 8.415 > 7.11
- P_Dw vs P_Up : 7.15469e-06
  mean: 8.0902 > 7.0096 , median: 8.415 > 7.605
- P_Dw vs LOW_PSI : 1.00762e-31
  mean: 8.0902 > 6.4194 , median: 8.415 > 6.87
- Bg_Upstrat vs HIGH_PSI : 7.11214e-49
  mean: 6.4599 < 8.3636 , median: 7.11 < 8.53
- Bg_Upstrat vs LOW_PSI : 0.0202678
  mean: 6.4599 > 6.4194 , median: 7.11 > 6.87
- P_Up vs HIGH_PSI : 2.46866e-10
  mean: 7.0096 < 8.3636 , median: 7.605 < 8.53

- P_Up vs LOW_PSI : 0.00285101
  mean: 7.0096 > 6.4194 , median: 7.605 > 6.87
- HIGH_PSI vs LOW_PSI : 1.16019e-242
  mean: 8.3636 > 6.4194 , median: 8.53 > 6.87

#### 6.27 MAXENTSCR HSAMODEL 5SS

Back to: Overview | ToC

Meaning: maximum entropy score of 5ss using a model trained with human splice sites

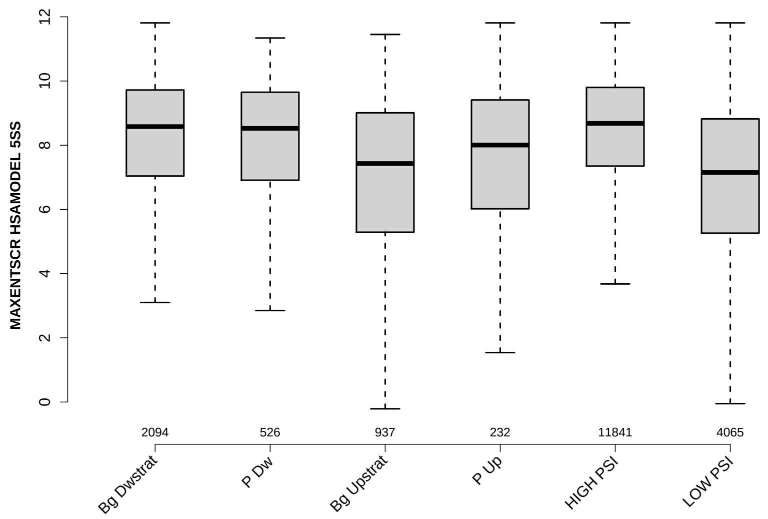

Significant results from Mann-Whitney U test:

- Bg_Dwstrat vs Bg_Upstrat : 1.06102e-28
  mean: 7.5602 > 6.4371 , median: 8.58 > 7.43
- Bg_Dwstrat vs P_Up : 0.000509523
  mean: 7.5602 > 7.1469 , median: 8.58 > 8.005
- Bg_Dwstrat vs HIGH_PSI : 0.000278896
  mean: 7.5602 < 8.2685 , median: 8.58 < 8.68
- Bg_Dwstrat vs LOW_PSI : 1.45834e-81
  mean: 7.5602 > 6.6078 , median: 8.58 > 7.15
- P_Dw vs Bg_Upstrat : 1.1415e-13
  mean: 7.833 > 6.4371 , median: 8.525 > 7.43
- P_Dw vs P_Up : 0.0107204
  mean: 7.833 > 7.1469 , median: 8.525 > 8.005
- P_Dw vs HIGH_PSI : 0.00220286
  mean: 7.833 < 8.2685 , median: 8.525 < 8.68
- P_Dw vs LOW_PSI : 1.65847e-25
  mean: 7.833 > 6.6078 , median: 8.525 > 7.15
- Bg_Upstrat vs P_Up : 0.00532544
  mean: 6.4371 < 7.1469 , median: 7.43 < 8.005
- Bg_Upstrat vs HIGH_PSI : 1.0074e-55
  mean: 6.4371 < 8.2685 , median: 7.43 < 8.68

- `P_Up` vs `HIGH_PSI` : 5.11853e-07
  mean: 7.1469 < 8.2685 , median: 8.005 < 8.68
- `P_Up` vs `LOW_PSI` : 4.59167e-05
  mean: 7.1469 > 6.6078 , median: 8.005 > 7.15
- `HIGH_PSI` vs `LOW_PSI` : 2.02321e-251
  mean: 8.2685 > 6.6078 , median: 8.68 > 7.15

#### 6.28 MAXENTSCR HSAMODEL DOWNSTRM 3SS

Back to: Overview | ToC

Meaning: maximum entropy score of 3ss of down-stream exon using a model trained with human splice sites

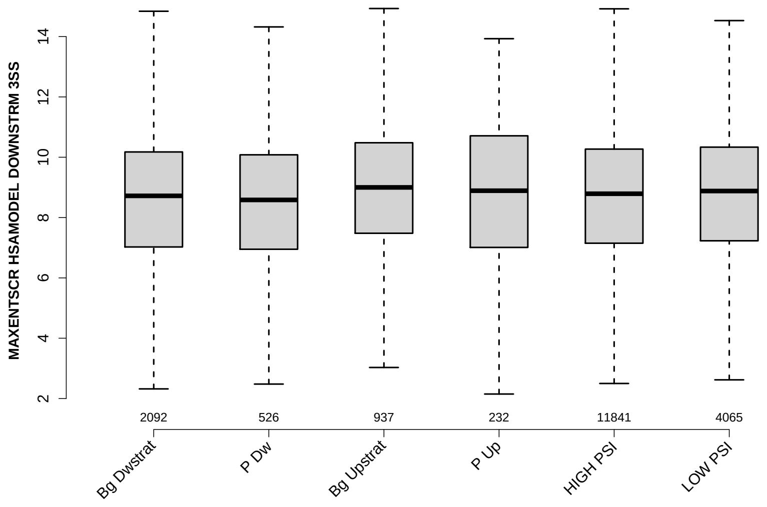

Significant results from Mann-Whitney U test:

- Bg_Dwstrat vs Bg_Upstrat : 0.00225467
  mean: 8.1915 < 8.612 , median: 8.72 < 9
- Bg_Dwstrat vs P_Up : 0.046527
  mean: 8.1915 < 8.8524 , median: 8.72 < 8.89
- Bg_Dwstrat vs LOW_PSI : 0.0154713
  mean: 8.1915 < 8.6738 , median: 8.72 < 8.88
- P_Dw vs Bg_Upstrat : 0.00271797
  mean: 8.4118 < 8.612 , median: 8.585 < 9
- P_Dw vs P_Up : 0.0281167
  mean: 8.4118 < 8.8524 , median: 8.585 < 8.89
- P_Dw vs LOW_PSI : 0.0208653
  mean: 8.4118 < 8.6738 , median: 8.585 < 8.88
- Bg_Upstrat vs HIGH_PSI : 0.0117709
  mean: 8.612 > 8.5977 , median: 9 > 8.79

#### 6.29 DIST FROM MAXBP TO 3SS UPINTRON

Back to: Overview | ToC

Meaning: distance to 3ss of best precited BP

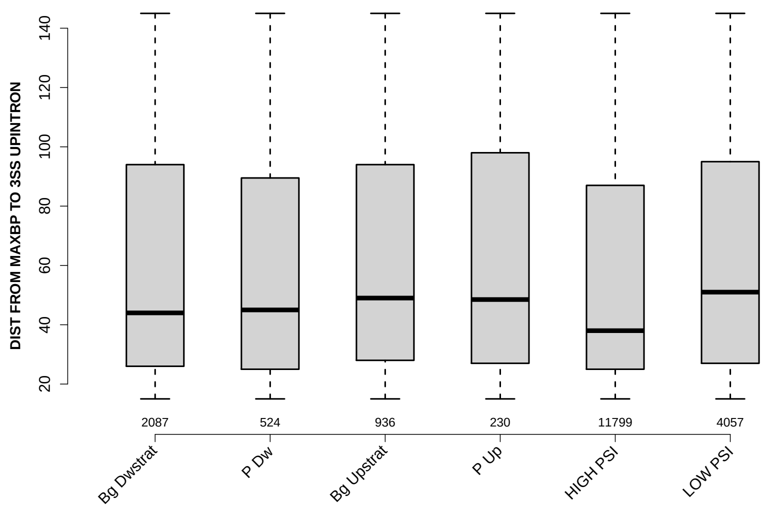

Significant results from Mann-Whitney U test:

- `Bg_Dwstrat` vs `HIGH_PSI` : 5.35358e-05
  mean: 60.1672 > 56.752 , median: 44 > 38
- `Bg_Dwstrat` vs `LOW_PSI` : 0.0108856
  mean: 60.1672 < 62.5943 , median: 44 < 51
- `P_Dw` vs `LOW_PSI` : 0.0497452
  mean: 59.5744 < 62.5943 , median: 45 < 51
- `Bg_Upstrat` vs `HIGH_PSI` : 1.02915e-06
  mean: 62.1004 > 56.752 , median: 49 > 38
- `P_Up` vs `HIGH_PSI` : 0.016019
  mean: 61.8826 > 56.752 , median: 48.5 > 38
- `HIGH_PSI` vs `LOW_PSI` : 2.20787e-19
  mean: 56.752 < 62.5943 , median: 38 < 51

#### 6.30 SCORE FOR MAXBP SEQ UPINTRON

Back to: Overview | ToC

Meaning: BP sequence score of best predicted BP

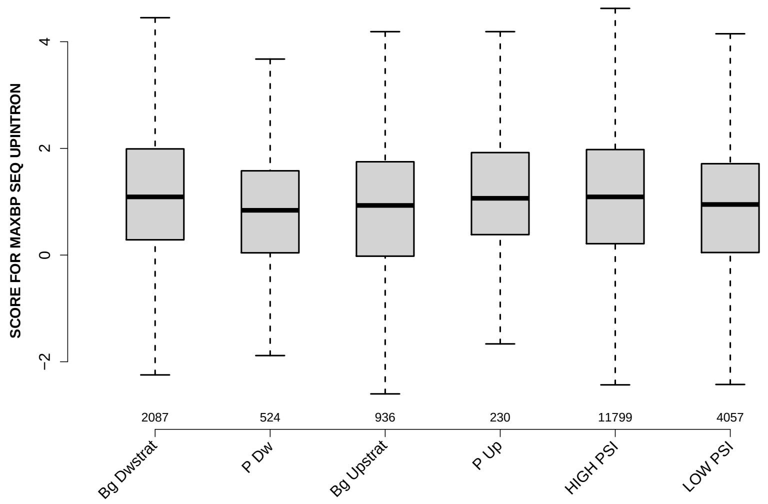

Significant results from Mann-Whitney U test:

- `Bg_Dwstrat` vs `P_Dw` : 7.34514e-06
  mean: 1.1367 > 0.857972 , median: 1.0909 > 0.839114
- `Bg_Dwstrat` vs `Bg_Upstrat` : 1.14964e-06
  mean: 1.1367 > 0.876834 , median: 1.0909 > 0.930543
- `Bg_Dwstrat` vs `LOW_PSI` : 2.94878e-12
  mean: 1.1367 > 0.872678 , median: 1.0909 > 0.948451
- `P_Dw` vs `P_Up` : 0.00935245
  mean: 0.857972 < 1.1042 , median: 0.839114 < 1.0644
- `P_Dw` vs `HIGH_PSI` : 1.13505e-05
  mean: 0.857972 < 1.1082 , median: 0.839114 < 1.0909
- `Bg_Upstrat` vs `P_Up` : 0.0170309
  mean: 0.876834 < 1.1042 , median: 0.930543 < 1.0644
- `Bg_Upstrat` vs `HIGH_PSI` : 6.85952e-07
  mean: 0.876834 < 1.1082 , median: 0.930543 < 1.0909
- `P_Up` vs `LOW_PSI` : 0.0111145
  mean: 1.1042 > 0.872678 , median: 1.0644 > 0.948451
- `HIGH_PSI` vs `LOW_PSI` : 1.12089e-19
  mean: 1.1082 > 0.872678 , median: 1.0909 > 0.948451

#### 6.31 PYRIMIDINECONT MAXBP UPINTRON

Back to: Overview | ToC

Meaning: Pyrimidine content between the BP adenine and the 3 prime splice site for best BP

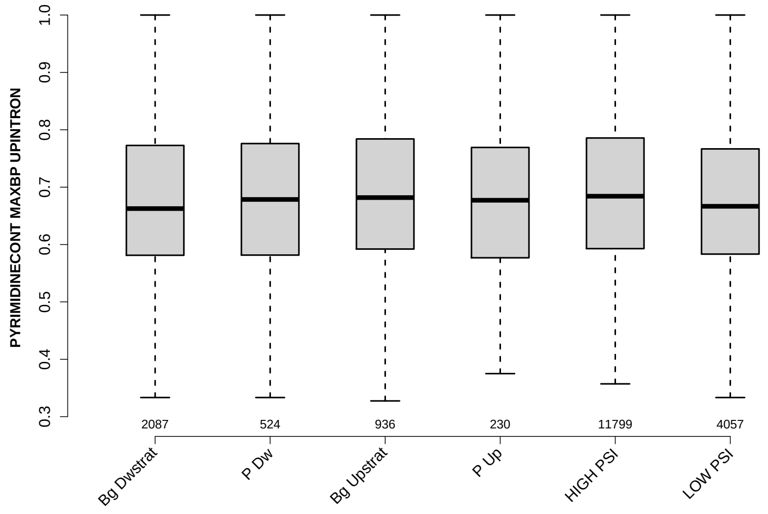

Significant results from Mann-Whitney U test:

- Bg_Dwstrat vs Bg_Upstrat : 0.0247588
  mean: 0.677415 $<$ 0.687378 , median: 0.662651 $<$ 0.681818
- Bg_Dwstrat vs HIGH_PSI : 8.5559e-06
  mean: 0.677415 $<$ 0.690566 , median: 0.662651 $<$ 0.684211
- Bg_Upstrat vs LOW_PSI : 0.00683573
  mean: 0.687378 $>$ 0.675846 , median: 0.681818 $>$ 0.666667
- HIGH_PSI vs LOW_PSI : 1.46861e-10
  mean: 0.690566 $>$ 0.675846 , median: 0.684211 $>$ 0.666667

#### 6.32 POLYPYRITRAC OFFSET MAXBP UPINTRON

Back to: Overview | ToC

Meaning: Polypyrimidine track offset relative to the BP adenine for best BP

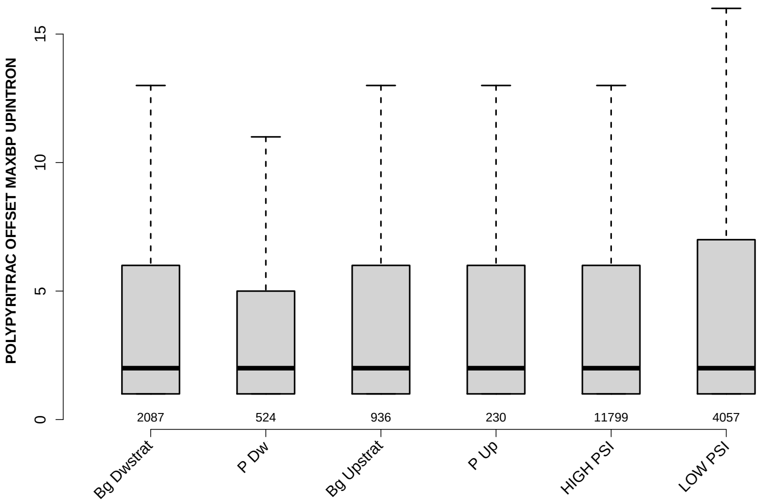

Significant results from Mann-Whitney U test:

- Bg_Dwstrat vs Bg_Upstrat : 0.0181389
  mean: 4.6766 > 4.4124 , median: 2 = 2
- Bg_Dwstrat vs HIGH_PSI : 0.0434821
  mean: 4.6766 > 4.4329 , median: 2 = 2
- Bg_Upstrat vs LOW_PSI : 0.0396919
  mean: 4.4124 < 4.667 , median: 2 = 2

#### 6.33 POLYPYRITRAC LEN MAXBP UPINTRON

Back to: Overview | ToC

Meaning: Polypyrimidine track length for best BP

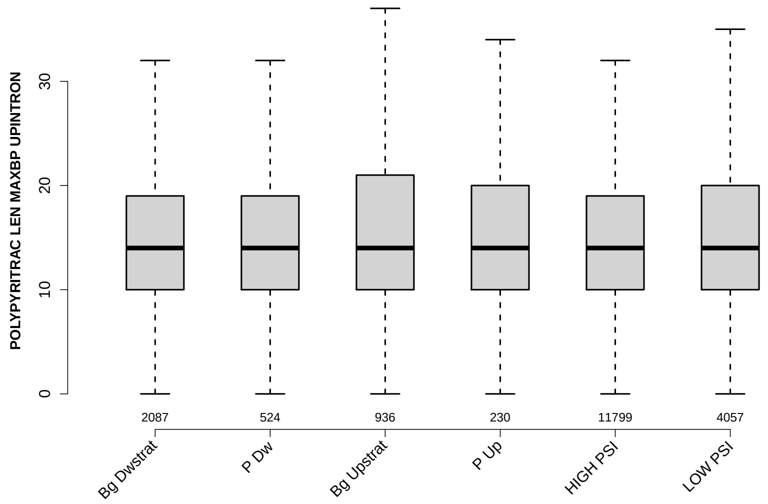

Significant results from Mann-Whitney U test:

- Bg_Dwstrat vs Bg_Upstrat : 0.000274923
  mean: 15.4212 < 17.3291 , median: 14 = 14
- P_Dw vs Bg_Upstrat : 0.0181075
  mean: 15.1851 < 17.3291 , median: 14 = 14
- Bg_Upstrat vs HIGH_PSI : 0.000558746
  mean: 17.3291 > 15.4354 , median: 14 = 14
- Bg_Upstrat vs LOW_PSI : 0.0128633
  mean: 17.3291 > 16.1122 , median: 14 = 14

#### 6.34 POLYPYRITRAC SCORE MAXBP UPINTRON

Back to: Overview | ToC

Meaning: Polypyrimidine track score for best BP

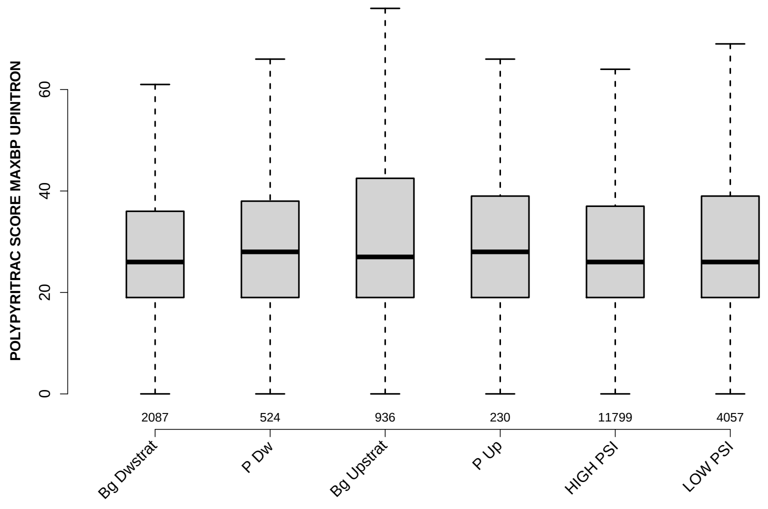

Significant results from Mann-Whitney U test:

- Bg_Dwstrat vs Bg_Upstrat : 4.94394e-05
  mean: 29.7331 < 34.578 , median: 26 < 27
- Bg_Dwstrat vs HIGH_PSI : 0.0350731
  mean: 29.7331 < 29.8694 , median: 26 = 26
- Bg_Dwstrat vs LOW_PSI : 0.00405806
  mean: 29.7331 < 32.4831 , median: 26 = 26
- Bg_Upstrat vs HIGH_PSI : 0.000614922
  mean: 34.578 > 29.8694 , median: 27 > 26
- Bg_Upstrat vs LOW_PSI : 0.0330169
  mean: 34.578 > 32.4831 , median: 27 > 26

#### 6.35 BPSCORE MAXBP UPINTRON

Back to: Overview | ToC

Meaning: SVM classification score of best BP

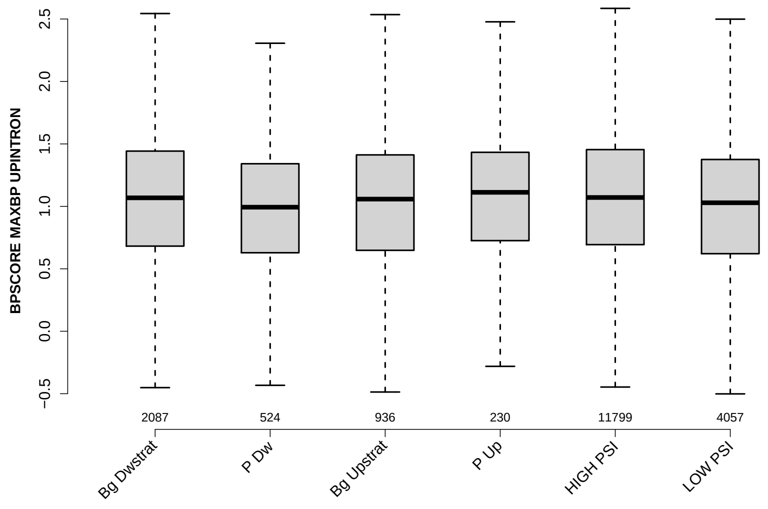

Significant results from Mann-Whitney U test:

- Bg_Dwstrat vs P_Dw : 0.00614938
  mean: 1.0466 > 0.976865 , median: 1.0683 > 0.993271
- Bg_Dwstrat vs LOW_PSI : 5.16637e-05
  mean: 1.0466 > 0.968946 , median: 1.0683 > 1.0288
- P_Dw vs P_Up : 0.00692198
  mean: 0.976865 < 1.0801 , median: 0.993271 < 1.1124
- P_Dw vs HIGH_PSI : 0.00099627
  mean: 0.976865 < 1.0564 , median: 0.993271 < 1.071
- P_Up vs LOW_PSI : 0.00724808
  mean: 1.0801 > 0.968946 , median: 1.1124 > 1.0288
- HIGH_PSI vs LOW_PSI : 1.70594e-11
  mean: 1.0564 > 0.968946 , median: 1.071 > 1.0288

#### 6.36 NUM PREDICTED BPS UPINTRON

Back to: Overview | ToC

Meaning: number of all predicted BPs which have a positive BP score

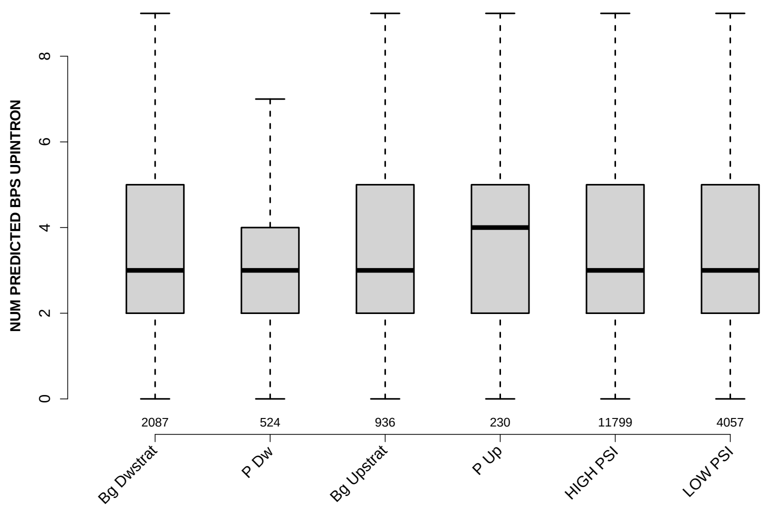

Significant results from Mann-Whitney U test:

- Bg_Dwstrat vs P_Up : 0.00230097
  mean: 3.3517 < 3.8174 , median: 3 < 4
- P_Dw vs P_Up : 0.00430557
  mean: 3.3359 < 3.8174 , median: 3 < 4
- Bg_Upstrat vs P_Up : 0.00715629
  mean: 3.3921 < 3.8174 , median: 3 < 4
- P_Up vs HIGH_PSI : 0.00674717
  mean: 3.8174 > 3.4326 , median: 4 > 3
- P_Up vs LOW_PSI : 0.00072975
  mean: 3.8174 > 3.314 , median: 4 > 3
- HIGH_PSI vs LOW_PSI : 0.00555472
  mean: 3.4326 > 3.314 , median: 3 = 3

#### 6.37 MEDIAN DIST FROM BP TO 3SS UPINTRON

Back to: Overview | ToC

Meaning: like DIST FROM MAXBP TO 3SS but median over top-3 predicted BPs

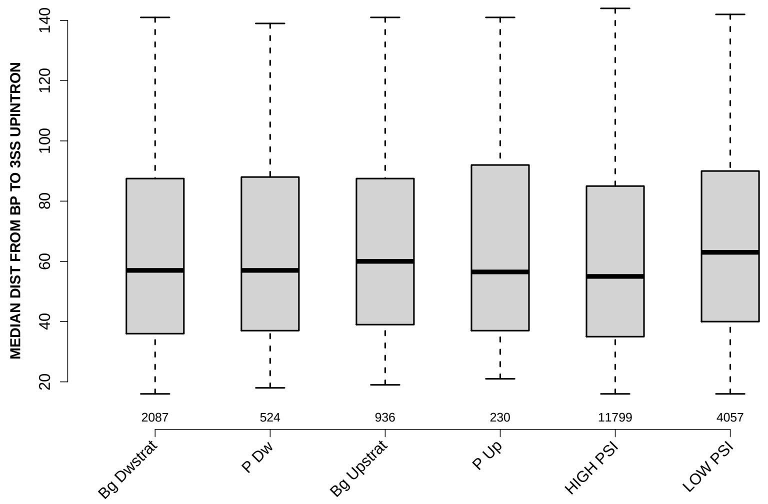

Significant results from Mann-Whitney U test:

- `Bg_Dwstrat` vs `HIGH_PSI` : 0.0403047
  mean: 63.425 > 62.0241 , median: 57 > 55
- `Bg_Dwstrat` vs `LOW_PSI` : 5.18137e-06
  mean: 63.425 < 66.7893 , median: 57 < 63
- `P_Dw` vs `LOW_PSI` : 0.0294431
  mean: 64.1221 < 66.7893 , median: 57 < 63
- `Bg_Upstrat` vs `HIGH_PSI` : 0.000434502
  mean: 65.0946 > 62.0241 , median: 60 > 55
- `HIGH_PSI` vs `LOW_PSI` : 6.16559e-21
  mean: 62.0241 < 66.7893 , median: 55 < 63

#### 6.38 MEDIAN SCORE FOR BPSEQ UPINTRON

Back to: Overview | ToC

Meaning: like SCORE FOR MAXBP SEQ but median over top-3 predicted BPs

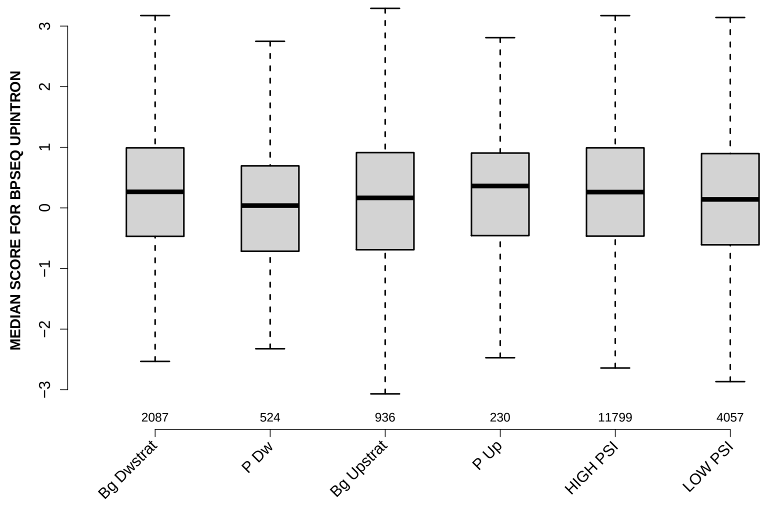

Significant results from Mann-Whitney U test:

- Bg_Dwstrat vs P_Dw : 3.69893e-05
  mean: 0.268167 > 0.043438 , median: 0.265045 > 0.0383488
- Bg_Dwstrat vs Bg_Upstrat : 0.0115236
  mean: 0.268167 > 0.14437 , median: 0.265045 > 0.165271
- Bg_Dwstrat vs LOW_PSI : 1.82619e-05
  mean: 0.268167 > 0.116244 , median: 0.265045 > 0.140235
- P_Dw vs P_Up : 0.0114867
  mean: 0.043438 < 0.24078 , median: 0.0383488 < 0.361763
- P_Dw vs HIGH_PSI : 3.24761e-06
  mean: 0.043438 < 0.270691 , median: 0.0383488 < 0.261123
- Bg_Upstrat vs HIGH_PSI : 0.00212852
  mean: 0.14437 < 0.270691 , median: 0.165271 < 0.261123
- HIGH_PSI vs LOW_PSI : 4.18483e-11
  mean: 0.270691 > 0.116244 , median: 0.261123 > 0.140235

#### 6.39 MEDIAN PYRIMIDINECONT UPINTRON

Back to: Overview | ToC

Meaning: like PYRIMIDINECONT MAXBP but median over top-3 predicted BPs

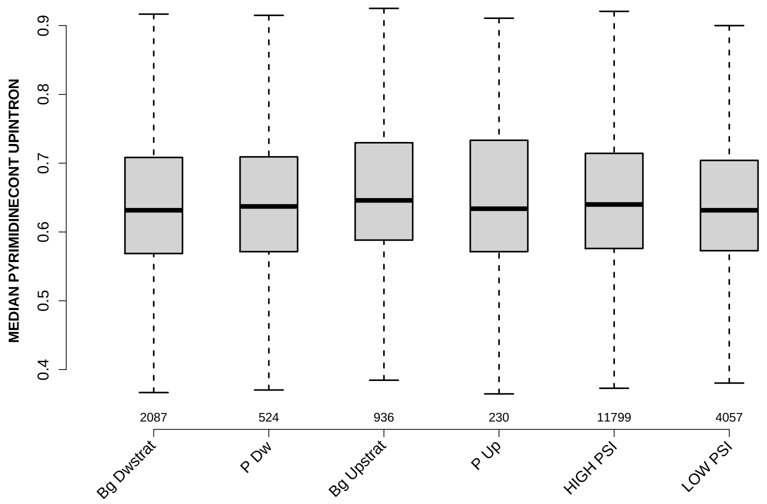

Significant results from Mann-Whitney U test:

- Bg_Dwstrat vs Bg_Upstrat : 0.000118346
  mean: 0.640799 < 0.654972 , median: 0.631579 < 0.645926
- Bg_Dwstrat vs HIGH_PSI : 0.00329623
  mean: 0.640799 < 0.647691 , median: 0.631579 < 0.64
- P_Dw vs Bg_Upstrat : 0.0221465
  mean: 0.643838 < 0.654972 , median: 0.637147 < 0.645926
- Bg_Upstrat vs HIGH_PSI : 0.0137469
  mean: 0.654972 > 0.647691 , median: 0.645926 > 0.64
- Bg_Upstrat vs LOW_PSI : 1.50889e-05
  mean: 0.654972 > 0.638956 , median: 0.645926 > 0.631579
- HIGH_PSI vs LOW_PSI : 4.74247e-05
  mean: 0.647691 > 0.638956 , median: 0.64 > 0.631579

#### 6.40 MEDIAN POLYPYRITRAC OFFSET UPINTRON

Back to: Overview | ToC

Meaning: like POLYPYRITRAC OFFSET MAXBP but median over top-3 predicted BPs

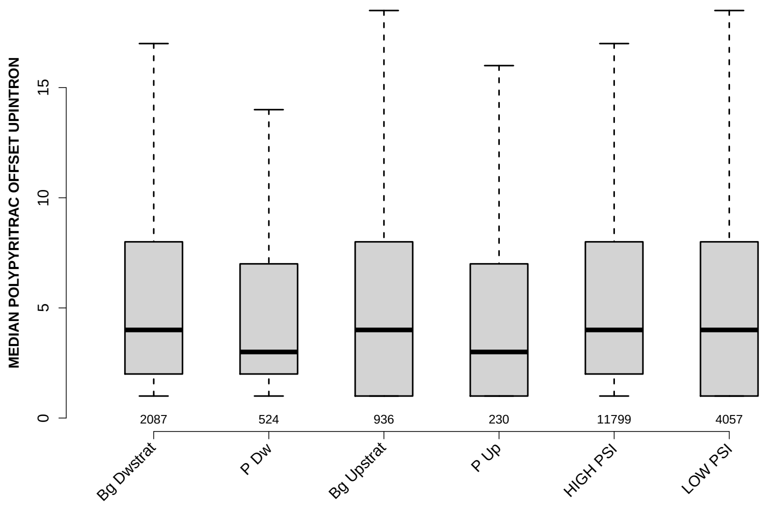

Significant results from Mann-Whitney U test:

- `Bg_Dwstrat` vs `P_Dw` : 0.0412185
  mean: 6.6021 > 5.7586 , median: 4 > 3
- `Bg_Dwstrat` vs `P_Up` : 0.0239183
  mean: 6.6021 > 5.8348 , median: 4 > 3

#### 6.41 MEDIAN POLYPYRITRAC LEN UPINTRON

Back to: Overview | ToC

Meaning: like POLYPYRITRAC LEN MAXBP but median over top-3 predicted BPs

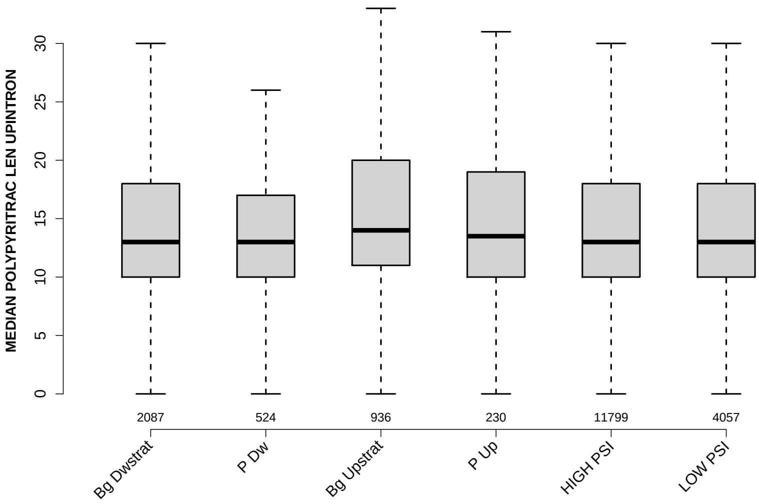

Significant results from Mann-Whitney U test:

- `Bg_Dwstrat` vs `Bg_Upstrat` : 0.00229029
  mean: 14.8857 < 16.3665 , median: 13 < 14
- `P_Dw` vs `Bg_Upstrat` : 0.00627707
  mean: 14.4084 < 16.3665 , median: 13 < 14
- `Bg_Upstrat` vs `HIGH_PSI` : 0.000778358
  mean: 16.3665 > 14.8453 , median: 14 > 13
- `Bg_Upstrat` vs `LOW_PSI` : 0.0203157
  mean: 16.3665 > 15.4028 , median: 14 > 13

#### 6.42 MEDIAN POLYPYRITRAC SCORE UPINTRON

Back to: Overview | ToC

Meaning: like POLYPYRITRAC SCORE MAXBP but median over top-3 predicted BPs

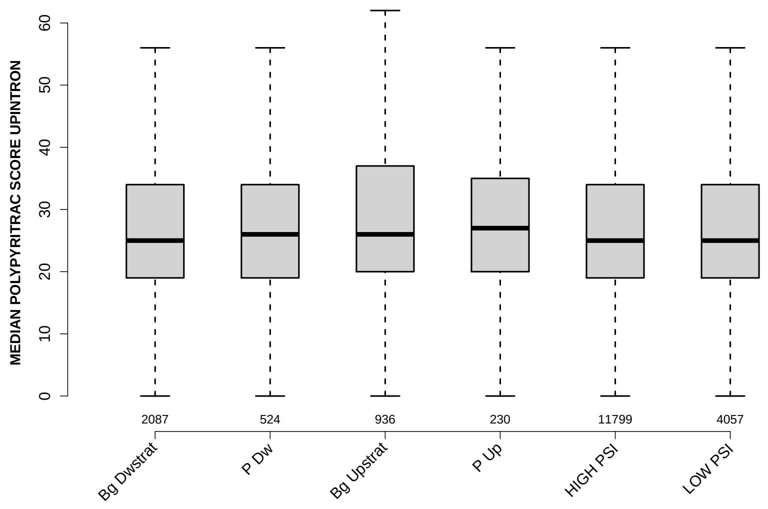

Significant results from Mann-Whitney U test:

- `Bg_Dwstrat` vs `Bg_Upstrat` : 0.00152568
  mean: 28.4789 < 32.3718 , median: 25 < 26
- `Bg_Upstrat` vs `HIGH_PSI` : 0.0029066
  mean: 32.3718 > 28.5696 , median: 26 > 25

#### 6.43 MEDIAN BPSCORE UPINTRON

Back to: Overview | ToC

Meaning: like BPSCORE MAXBP but median over top-3 predicted BPs

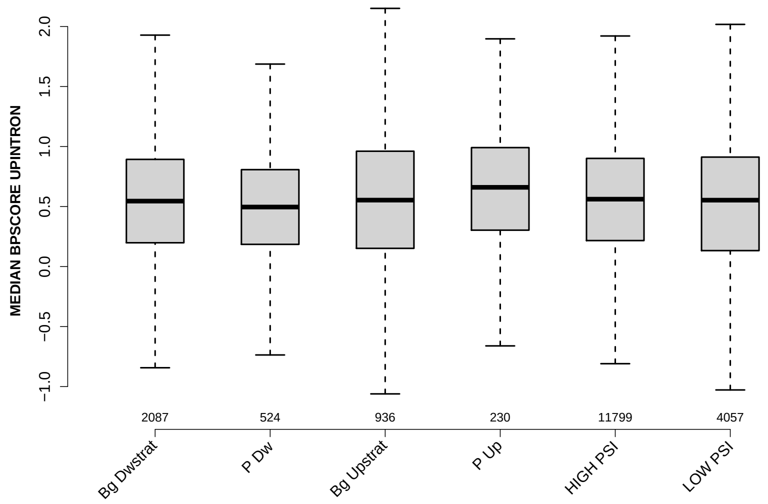

Significant results from Mann-Whitney U test:

- Bg_Dwstrat vs P_Dw : 0.0429416
  mean: 0.4722 > 0.426673 , median: 0.545311 > 0.495957
- Bg_Dwstrat vs P_Up : 0.0109371
  mean: 0.4722 < 0.557884 , median: 0.545311 < 0.66009
- P_Dw vs P_Up : 0.000300781
  mean: 0.426673 < 0.557884 , median: 0.495957 < 0.66009
- P_Dw vs HIGH_PSI : 0.00416372
  mean: 0.426673 < 0.47743 , median: 0.495957 < 0.561507
- Bg_Upstrat vs P_Up : 0.0292457
  mean: 0.463458 < 0.557884 , median: 0.553525 < 0.66009
- P_Up vs HIGH_PSI : 0.0209824
  mean: 0.557884 > 0.47743 , median: 0.66009 > 0.561507
- P_Up vs LOW_PSI : 0.00563963
  mean: 0.557884 > 0.446658 , median: 0.66009 > 0.553232
- HIGH_PSI vs LOW_PSI : 0.0278463
  mean: 0.47743 > 0.446658 , median: 0.561507 > 0.553232

#### 6.44 DIST FROM MAXBP TO 3SS DOINTRON

Back to: Overview | ToC

Meaning: distance to 3ss of best precited BP

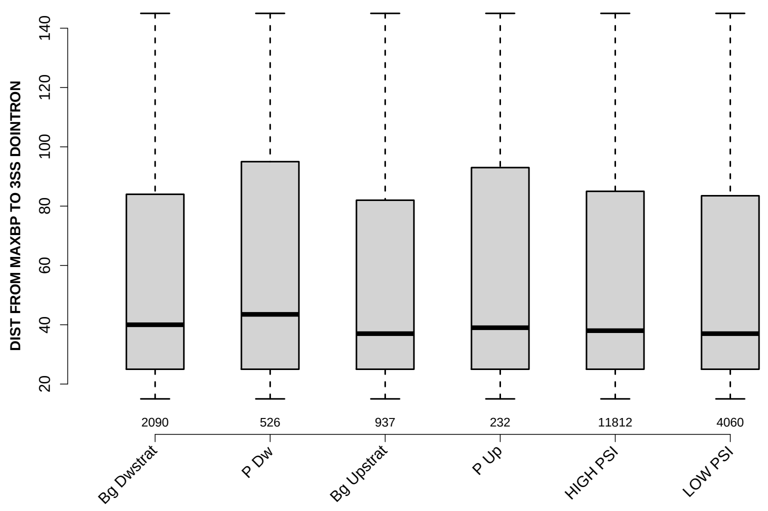

Significant results from Mann-Whitney U test:

- `P_Dw` vs `LOW_PSI` : 0.0299561
  mean: 59.7376 > 55.2894 , median: 43.5 > 37

#### 6.45 SCORE FOR MAXBP SEQ DOINTRON

Back to: Overview | ToC

Meaning: BP sequence score of best predicted BP

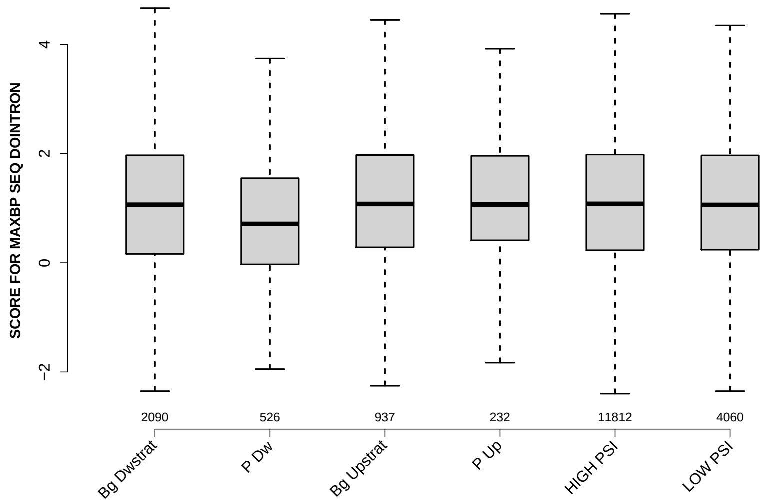

Significant results from Mann-Whitney U test:

- `Bg_Dwstrat` vs `P_Dw` : 1.62217e-06
  mean: 1.0878 > 0.798855 , median: 1.0633 > 0.711689
- `P_Dw` vs `Bg_Upstrat` : 3.55529e-06
  mean: 0.798855 < 1.0959 , median: 0.711689 < 1.0773
- `P_Dw` vs `P_Up` : 0.000451178
  mean: 0.798855 < 1.0955 , median: 0.711689 < 1.0673
- `P_Dw` vs `HIGH_PSI` : 5.6048e-09
  mean: 0.798855 < 1.1124 , median: 0.711689 < 1.0796
- `P_Dw` vs `LOW_PSI` : 2.23585e-07
  mean: 0.798855 < 1.0886 , median: 0.711689 < 1.06

#### 6.46 PYRIMIDINECONT MAXBP DOINTRON

Back to: Overview | ToC

Meaning: Pyrimidine content between the BP adenine and the 3 prime splice site for best BP

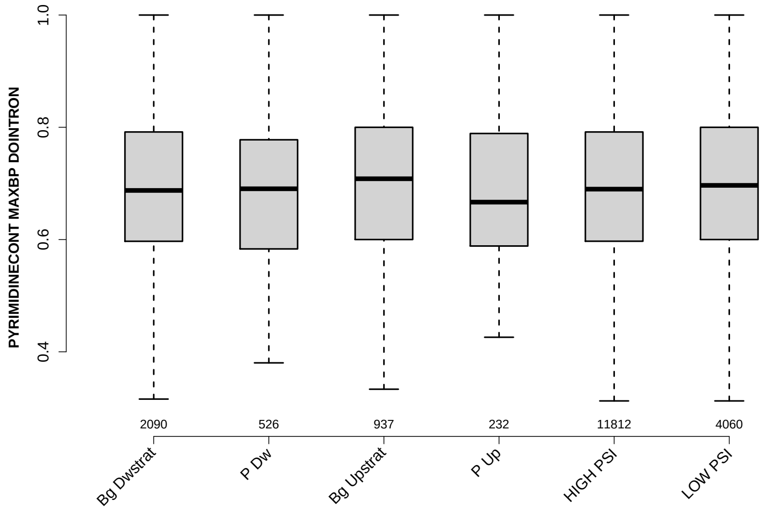

Significant results from Mann-Whitney U test:

- `Bg_Dwstrat` vs `Bg_Upstrat` : 0.0262578
  mean: 0.693046 < 0.706042 , median: 0.6875 < 0.708333
- `P_Dw` vs `Bg_Upstrat` : 0.0217364
  mean: 0.690307 < 0.706042 , median: 0.690476 < 0.708333
- `Bg_Upstrat` vs `HIGH_PSI` : 0.0298414
  mean: 0.706042 > 0.696049 , median: 0.708333 > 0.689828
- `HIGH_PSI` vs `LOW_PSI` : 0.0490788
  mean: 0.696049 < 0.700693 , median: 0.689828 < 0.696529

#### 6.47 POLYPYRITRAC OFFSET MAXBP DOINTRON

Back to: Overview | ToC

Meaning: Polypyrimidine track offset relative to the BP adenine for best BP

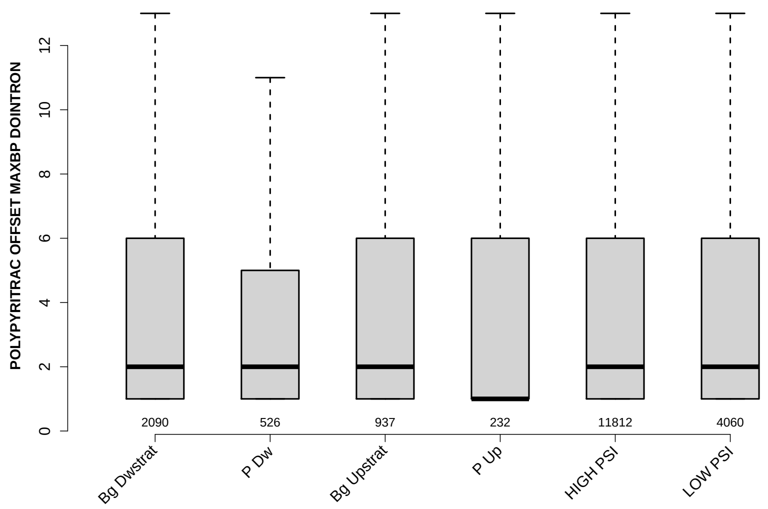

Significant results from Mann-Whitney U test:

- Bg_Dwstrat vs P_Dw : 0.00224718
  mean: 4.5861 > 3.7167 , median: 2 = 2
- Bg_Dwstrat vs LOW_PSI : 0.0432933
  mean: 4.5861 > 4.1729 , median: 2 = 2
- P_Dw vs HIGH_PSI : 0.0165101
  mean: 3.7167 < 4.3718 , median: 2 = 2
- P_Dw vs LOW_PSI : 0.0428592
  mean: 3.7167 < 4.1729 , median: 2 = 2

#### 6.48 POLYPYRITRAC LEN MAXBP DOINTRON

Back to: Overview | ToC

Meaning: Polypyrimidine track length for best BP

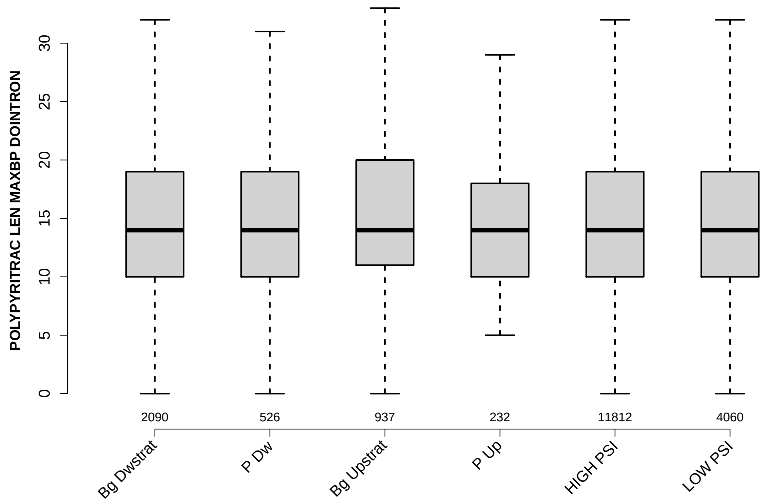

Significant results from Mann-Whitney U test:

- none

#### 6.49 POLYPYRITRAC SCORE MAXBP DOINTRON

Back to: Overview | ToC

Meaning: Polypyrimidine track score for best BP

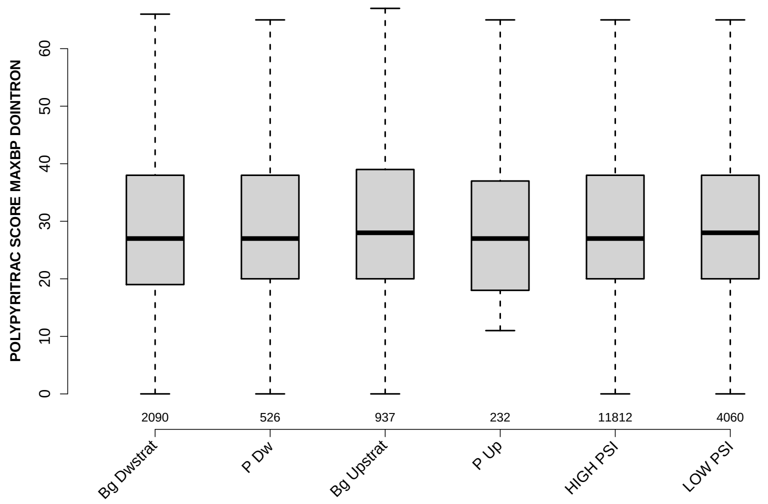

Significant results from Mann-Whitney U test:

- `Bg_Dwstrat` vs `Bg_Upstrat` : 0.0266479
  mean: 30.4837 < 31.9157 , median: 27 < 28
- `Bg_Dwstrat` vs `LOW_PSI` : 0.0186998
  mean: 30.4837 < 31.1062 , median: 27 < 28
- `HIGH_PSI` vs `LOW_PSI` : 0.0483246
  mean: 30.6173 < 31.1062 , median: 27 < 28

#### 6.50 BPSCORE MAXBP DOINTRON

Back to: Overview | ToC

Meaning: SVM classification score of best BP

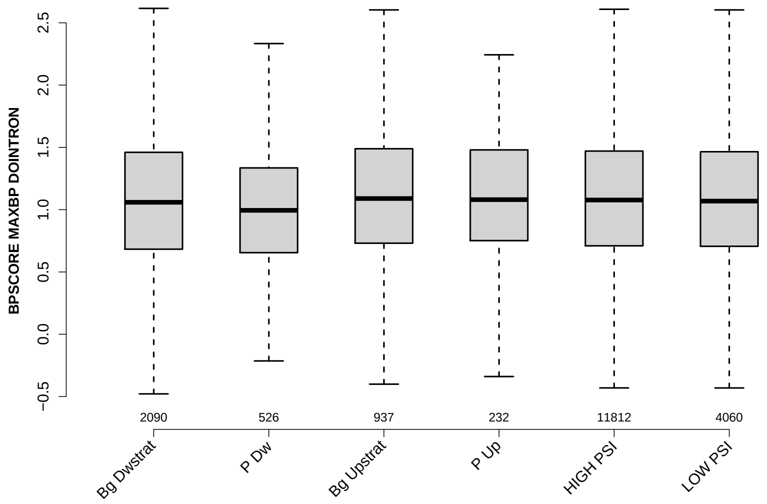

Significant results from Mann-Whitney U test:

- Bg_Dwstrat vs P_Dw : 0.0155295
  mean: 1.0452 > 0.988494 , median: 1.0591 > 0.99467
- P_Dw vs Bg_Upstrat : 0.000351892
  mean: 0.988494 < 1.105 , median: 0.99467 < 1.0894
- P_Dw vs P_Up : 0.0077813
  mean: 0.988494 < 1.0798 , median: 0.99467 < 1.0805
- P_Dw vs HIGH_PSI : 0.000290586
  mean: 0.988494 < 1.0706 , median: 0.99467 < 1.0773
- P_Dw vs LOW_PSI : 0.000692142
  mean: 0.988494 < 1.08 , median: 0.99467 < 1.0686

#### 6.51 NUM PREDICTED BPS DOINTRON

Back to: Overview | ToC

Meaning: number of all predicted BPs which have a positive BP score

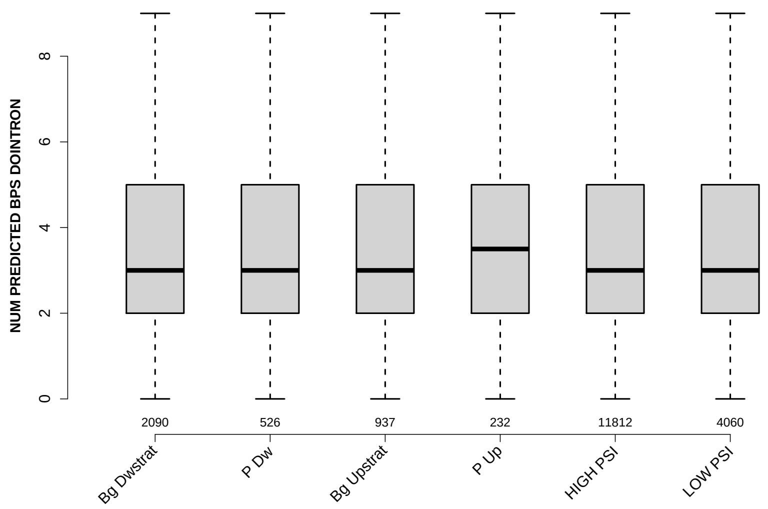

Significant results from Mann-Whitney U test:

- Bg_Dwstrat vs P_Dw : 0.00286986
  mean: 3.4196 < 3.6958 , median: 3 = 3
- Bg_Dwstrat vs Bg_Upstrat : 0.00862634
  mean: 3.4196 < 3.6147 , median: 3 = 3
- Bg_Dwstrat vs P_Up : 0.0291059
  mean: 3.4196 < 3.7284 , median: 3 < 3.5
- Bg_Dwstrat vs LOW_PSI : 0.0109064
  mean: 3.4196 < 3.5421 , median: 3 = 3
- P_Dw vs HIGH_PSI : 0.00378885
  mean: 3.6958 > 3.4495 , median: 3 = 3
- Bg_Upstrat vs HIGH_PSI : 0.0106364
  mean: 3.6147 > 3.4495 , median: 3 = 3
- P_Up vs HIGH_PSI : 0.0407418
  mean: 3.7284 > 3.4495 , median: 3.5 > 3
- HIGH_PSI vs LOW_PSI : 0.00423624
  mean: 3.4495 < 3.5421 , median: 3 = 3

#### 6.52 MEDIAN DIST FROM BP TO 3SS DOINTRON

Back to: Overview | ToC

Meaning: like DIST FROM MAXBP TO 3SS but median over top-3 predicted BPs

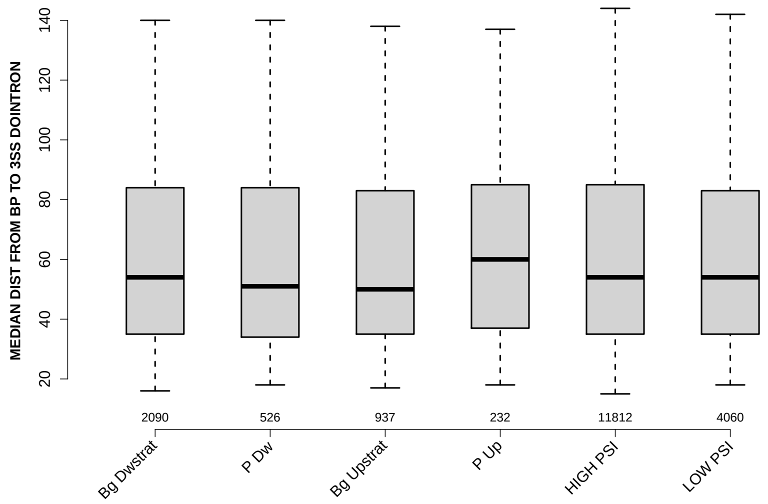

Significant results from Mann-Whitney U test:

- none

#### 6.53 MEDIAN SCORE FOR BPSEQ DOINTRON

Back to: Overview | ToC

Meaning: like SCORE FOR MAXBP SEQ but median over top-3 predicted BPs

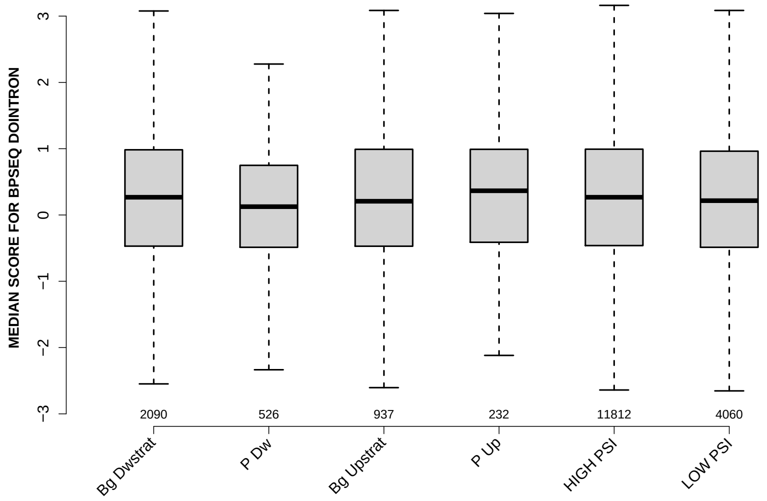

Significant results from Mann-Whitney U test:

- `Bg_Dwstrat` vs `P_Dw` : 0.0158903
  mean: 0.268725 > 0.134946 , median: 0.267625 > 0.125394
- `P_Dw` vs `P_Up` : 0.0236604
  mean: 0.134946 < 0.318468 , median: 0.125394 < 0.365417
- `P_Dw` vs `HIGH_PSI` : 0.00434327
  mean: 0.134946 < 0.272885 , median: 0.125394 < 0.267625
- `P_Dw` vs `LOW_PSI` : 0.045491
  mean: 0.134946 < 0.233802 , median: 0.125394 < 0.215885

#### 6.54 MEDIAN PYRIMIDINECONT DOINTRON

Back to: Overview | ToC

Meaning: like PYRIMIDINECONT MAXBP but median over top-3 predicted BPs

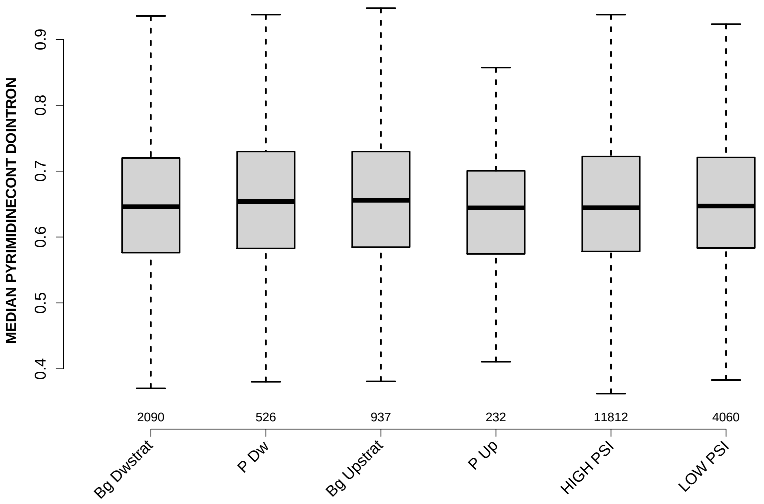

Significant results from Mann-Whitney U test:

- `Bg_Dwstrat` vs `Bg_Upstrat` : 0.0139507
  mean: 0.649558 < 0.660024 , median: 0.645994 < 0.655738
- `Bg_Upstrat` vs `P_Up` : 0.0347205
  mean: 0.660024 > 0.646059 , median: 0.655738 > 0.644256
- `Bg_Upstrat` vs `HIGH_PSI` : 0.0126978
  mean: 0.660024 > 0.651893 , median: 0.655738 > 0.644444

#### 6.55 MEDIAN POLYPYRITRAC OFFSET DOINTRON

Back to: Overview | ToC

Meaning: like POLYPYRITRAC OFFSET MAXBP but median over top-3 predicted BPs

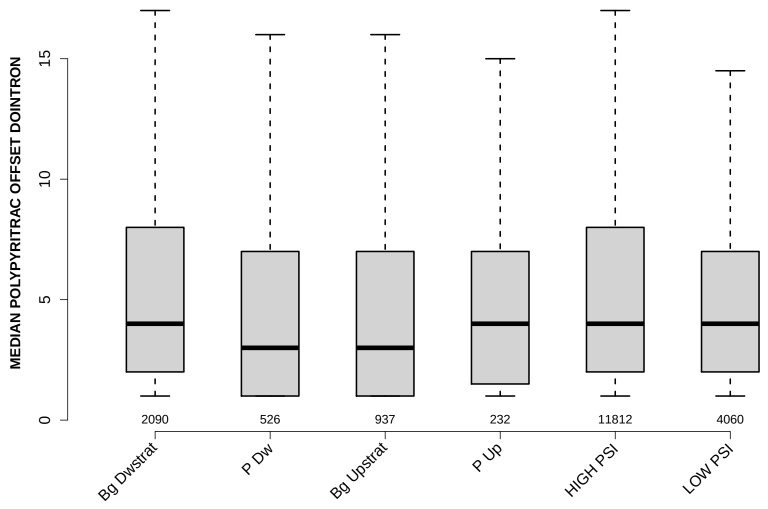

Significant results from Mann-Whitney U test:

- `Bg_Dwstrat` vs `P_Dw` : 0.0268411
  mean: 6.7672 > 5.4876 , median: 4 > 3
- `Bg_Dwstrat` vs `Bg_Upstrat` : 0.0136868
  mean: 6.7672 > 5.5891 , median: 4 > 3
- `Bg_Dwstrat` vs `LOW_PSI` : 0.0168207
  mean: 6.7672 > 5.9451 , median: 4 = 4

#### 6.56 MEDIAN POLYPYRITRAC LEN DOINTRON

Back to: Overview | ToC

Meaning: like POLYPYRITRAC LEN MAXBP but median over top-3 predicted BPs

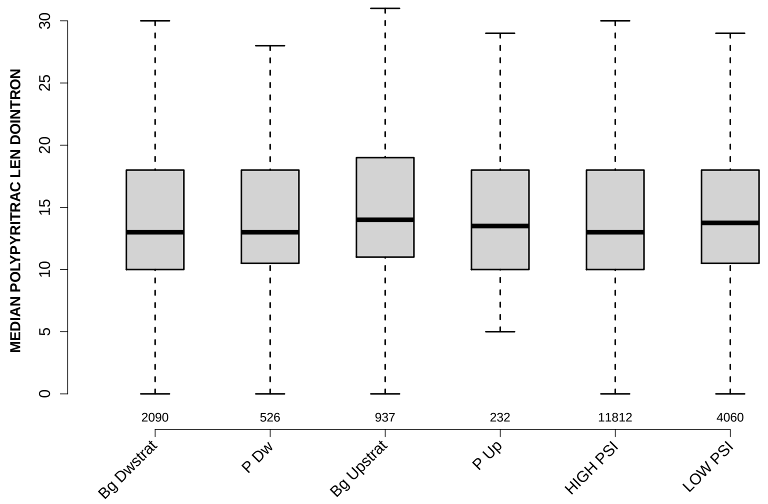

Significant results from Mann-Whitney U test:

- none

#### 6.57 MEDIAN POLYPYRITRAC SCORE DOINTRON

Back to: Overview | ToC

Meaning: like POLYPYRITRAC SCORE MAXBP but median over top-3 predicted BPs

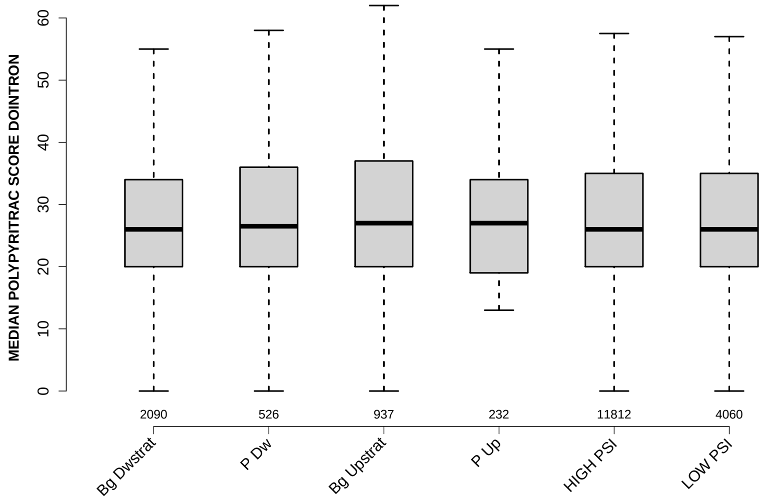

Significant results from Mann-Whitney U test:

- `Bg_Dwstrat` vs `Bg_Upstrat` : 0.0158292
  mean: 29.1347 < 30.8943 , median: 26 < 27
- `Bg_Upstrat` vs `HIGH_PSI` : 0.0103973
  mean: 30.8943 > 29.1302 , median: 27 > 26

#### 6.58 MEDIAN BPSCORE DOINTRON

Back to: Overview | ToC

Meaning: like BPSCORE MAXBP but median over top-3 predicted BPs

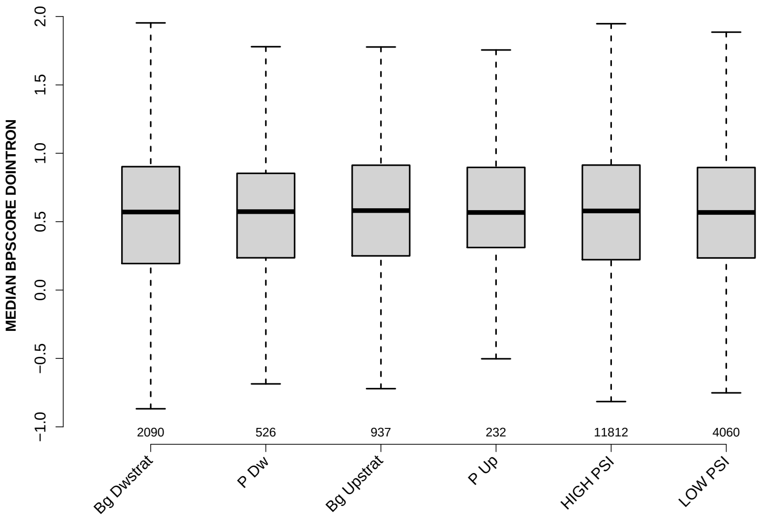

Significant results from Mann-Whitney U test:

- none

#### 6.59 MEDIAN TR LENGTH

Back to: Overview | ToC

Meaning: median length of transcripts the exon occurs in

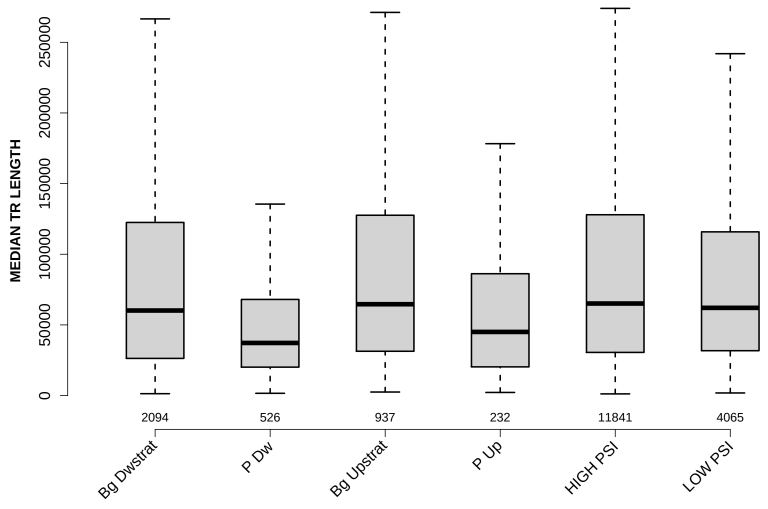

Significant results from Mann-Whitney U test:

- Bg_Dwstrat vs P_Dw : 1.39904e-18
  mean: 98713.5592 > 55503.9753 , median: 60169.5 > 37220.5
- Bg_Dwstrat vs Bg_Upstrat : 0.0386633
  mean: 98713.5592 < 103873.5224 , median: 60169.5 < 64633
- Bg_Dwstrat vs P_Up : 0.000216755
  mean: 98713.5592 > 68879.4741 , median: 60169.5 > 45020.5
- Bg_Dwstrat vs HIGH_PSI : 0.00345048
  mean: 98713.5592 < 103328.8065 , median: 60169.5 < 65127.5
- P_Dw vs Bg_Upstrat : 7.81053e-23
  mean: 55503.9753 < 103873.5224 , median: 37220.5 < 64633
- P_Dw vs P_Up : 0.0241195
  mean: 55503.9753 < 68879.4741 , median: 37220.5 < 45020.5
- P_Dw vs HIGH_PSI : 1.17756e-30
  mean: 55503.9753 < 103328.8065 , median: 37220.5 < 65127.5
- P_Dw vs LOW_PSI : 9.55341e-28
  mean: 55503.9753 < 94287.6311 , median: 37220.5 < 62069
- Bg_Upstrat vs P_Up : 1.49259e-06
  mean: 103873.5224 > 68879.4741 , median: 64633 > 45020.5
- P_Up vs HIGH_PSI : 5.07094e-07
  mean: 68879.4741 < 103328.8065 , median: 45020.5 < 65127.5

- `P_Up` vs `LOW_PSI` : 2.89453e-06
  mean: 68879.4741 < 94287.6311 , median: 45020.5 < 62069
- `HIGH_PSI` vs `LOW_PSI` : 0.0486732
  mean: 103328.8065 > 94287.6311 , median: 65127.5 > 62069

#### 6.60 MEDIAN EXON NUMBER

Back to: Overview | ToC

Meaning: ... of transcripts where exon was found in

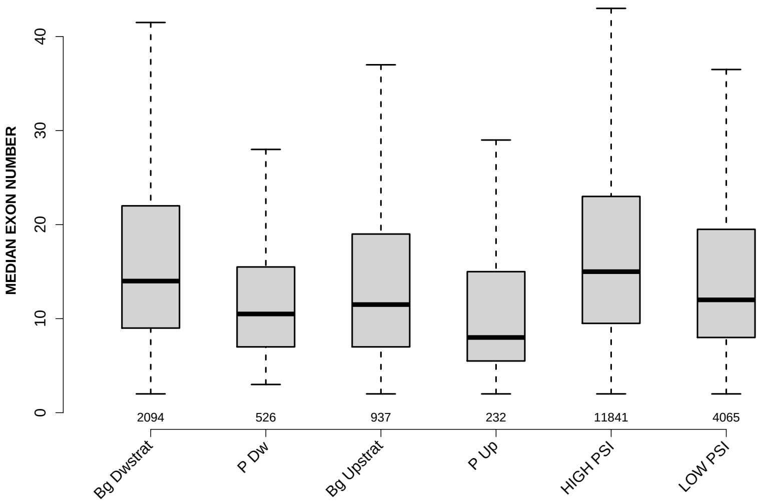

Significant results from Mann-Whitney U test:

- Bg_Dwstrat vs P_Dw : 5.65105e-18
  mean: 17.4248 > 12.3213 , median: 14 > 10.5
- Bg_Dwstrat vs Bg_Upstrat : 9.89014e-11
  mean: 17.4248 > 14.6745 , median: 14 > 11.5
- Bg_Dwstrat vs P_Up : 3.95214e-19
  mean: 17.4248 > 11.1444 , median: 14 > 8
- Bg_Dwstrat vs HIGH_PSI : 9.77143e-06
  mean: 17.4248 < 18.2213 , median: 14 < 15
- Bg_Dwstrat vs LOW_PSI : 4.96177e-10
  mean: 17.4248 > 15.1827 , median: 14 > 12
- P_Dw vs Bg_Upstrat : 0.0127603
  mean: 12.3213 < 14.6745 , median: 10.5 < 11.5
- P_Dw vs P_Up : 0.000319817
  mean: 12.3213 > 11.1444 , median: 10.5 > 8
- P_Dw vs HIGH_PSI : 2.33651e-33
  mean: 12.3213 < 18.2213 , median: 10.5 < 15
- P_Dw vs LOW_PSI : 6.75391e-08
  mean: 12.3213 < 15.1827 , median: 10.5 < 12
- Bg_Upstrat vs P_Up : 3.21617e-07
  mean: 14.6745 > 11.1444 , median: 11.5 > 8

- Bg_Upstrat vs HIGH_PSI : 1.21263e-25
  mean: $14.6745 < 18.2213$ , median: $11.5 < 15$
- Bg_Upstrat vs LOW_PSI : 0.00946795
  mean: $14.6745 < 15.1827$ , median: $11.5 < 12$
- P_Up vs HIGH_PSI : 8.78985e-27
  mean: $11.1444 < 18.2213$ , median: $8 < 15$
- P_Up vs LOW_PSI : 2.62563e-12
  mean: $11.1444 < 15.1827$ , median: $8 < 12$
- HIGH_PSI vs LOW_PSI : 2.03386e-51
  mean: $18.2213 > 15.1827$ , median: $15 > 12$

#### 6.61 EXON MEDIANRELATIVERANK

Back to: Overview | ToC

Meaning: relative rank = rank / number of all exons in transcript, is between 0 and 1

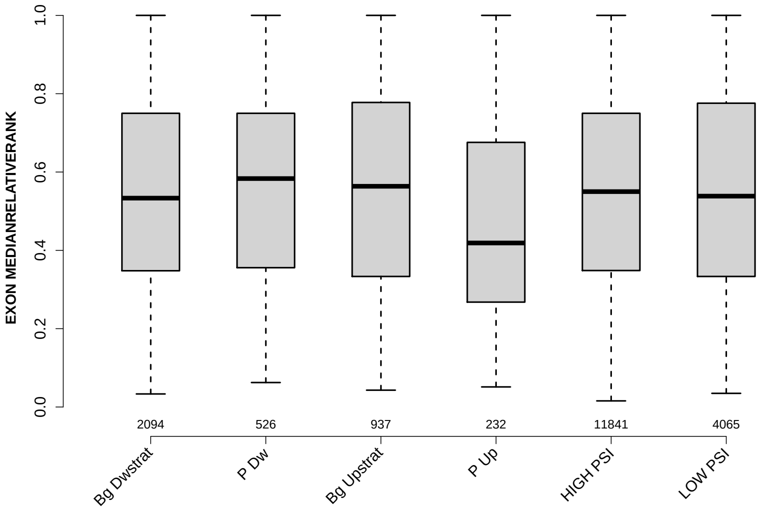

Significant results from Mann-Whitney U test:

- `Bg_Dwstrat` vs `P_Up` : 0.000240693
  mean: 0.541279 > 0.485384 , median: 0.533333 > 0.41886
- `P_Dw` vs `P_Up` : 0.000116859
  mean: 0.554665 > 0.485384 , median: 0.583333 > 0.41886
- `Bg_Upstrat` vs `P_Up` : 0.000357904
  mean: 0.555331 > 0.485384 , median: 0.563636 > 0.41886
- `P_Up` vs `HIGH_PSI` : 7.59871e-05
  mean: 0.485384 < 0.54462 , median: 0.41886 < 0.55
- `P_Up` vs `LOW_PSI` : 0.000158959
  mean: 0.485384 < 0.551282 , median: 0.41886 < 0.538462

#### 6.62 EXON MEDIANRELATIVERANK 3BINS

Back to: Overview | ToC

Meaning: median bin into which EXON MEDIANRELATIVERANK falls when binning 0-1 into 3 bins

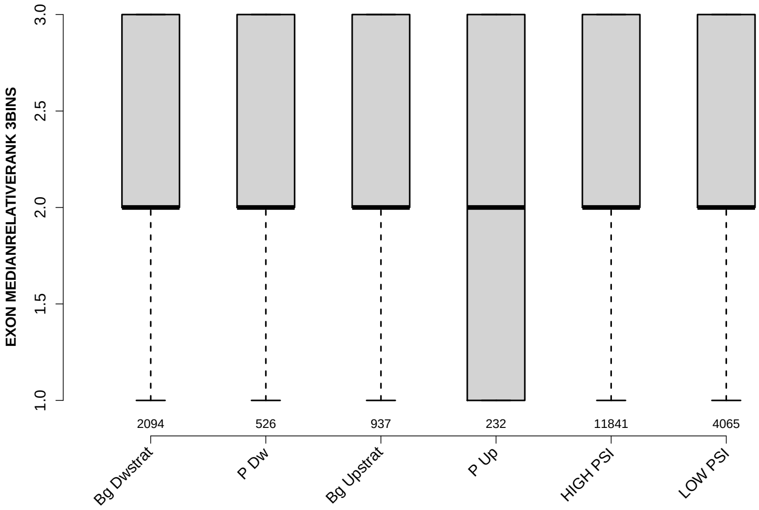

Significant results from Mann-Whitney U test:

- `Bg_Dwstrat` vs `P_Up` : 0.00303641
  mean: 2.1404 > 1.9784 , median: 2 = 2
- `P_Dw` vs `P_Up` : 0.00304376
  mean: 2.1635 > 1.9784 , median: 2 = 2
- `Bg_Upstrat` vs `P_Up` : 0.00366067
  mean: 2.1483 > 1.9784 , median: 2 = 2
- `P_Up` vs `HIGH_PSI` : 0.00146003
  mean: 1.9784 < 2.1448 , median: 2 = 2
- `P_Up` vs `LOW_PSI` : 0.0039
  mean: 1.9784 < 2.1336 , median: 2 = 2

#### 6.63 EXON MEDIANRELATIVERANK 5BINS

Back to: Overview | ToC

Meaning: similar to EXON MEDIANRELATIVERANK 3BINS with 5 bins

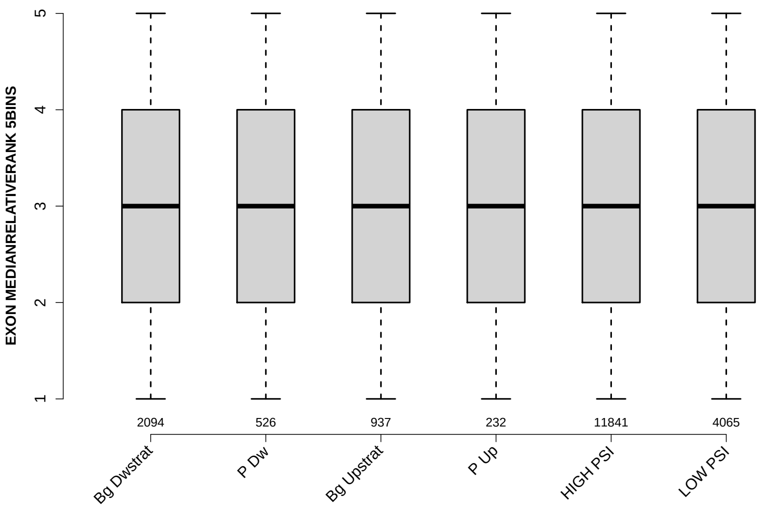

Significant results from Mann-Whitney U test:

- Bg_Dwstrat vs P_Up : 6.9058e-05
  mean: 3.2254 > 2.8879 , median: 3 = 3
- P_Dw vs P_Up : 4.62902e-05
  mean: 3.2814 > 2.8879 , median: 3 = 3
- Bg_Upstrat vs P_Up : 0.000270268
  mean: 3.2369 > 2.8879 , median: 3 = 3
- P_Up vs HIGH_PSI : 2.72923e-05
  mean: 2.8879 < 3.2288 , median: 3 = 3
- P_Up vs LOW_PSI : 7.11608e-05
  mean: 2.8879 < 3.2339 , median: 3 = 3

#### 6.64 EXON MEDIANRELATIVERANK 10BINS

Back to: Overview | ToC

Meaning: similar to EXON MEDIANRELATIVERANK 3BINS with 10 bins

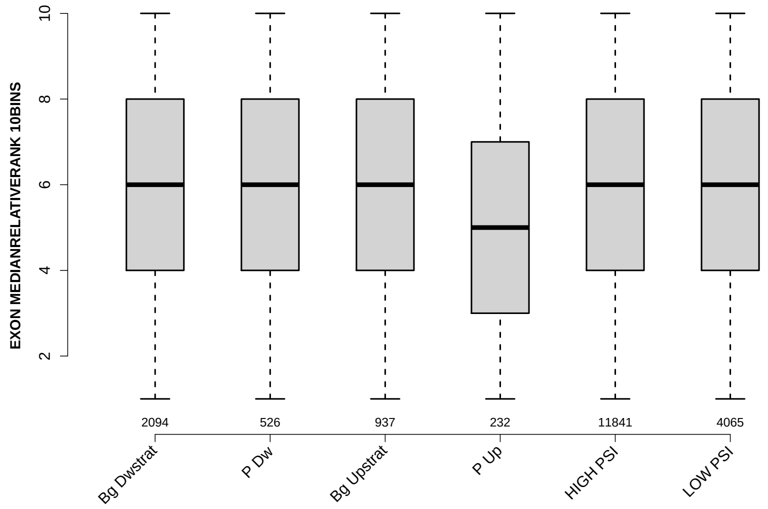

Significant results from Mann-Whitney U test:

- Bg_Dwstrat vs P_Up : 0.000308498
  mean: 5.9475 > 5.3707 , median: 6 > 5
- P_Dw vs P_Up : 9.80214e-05
  mean: 6.0894 > 5.3707 , median: 6 > 5
- Bg_Upstrat vs P_Up : 0.000349191
  mean: 6.0544 > 5.3707 , median: 6 > 5
- P_Up vs HIGH_PSI : 8.91682e-05
  mean: 5.3707 < 5.9764 , median: 5 < 6
- P_Up vs LOW_PSI : 0.000207809
  mean: 5.3707 < 6.0133 , median: 5 < 6

#### 6.65 NTRS ALL FOR GENE

Back to: Overview | ToC

Meaning: number of transcripts of gene where the exon was found in

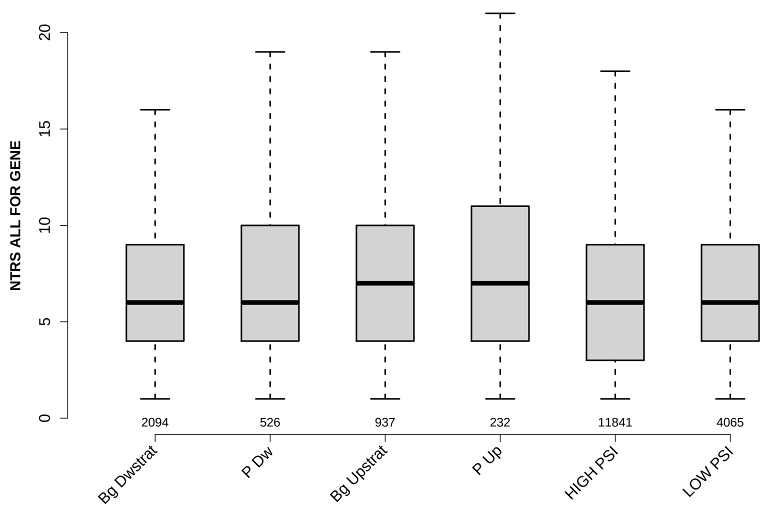

Significant results from Mann-Whitney U test:

- Bg_Dwstrat vs Bg_Upstrat : 0.000497538
  mean: 7.1213 < 7.7311 , median: 6 < 7
- Bg_Dwstrat vs P_Up : 0.000516929
  mean: 7.1213 < 8.2845 , median: 6 < 7
- P_Dw vs P_Up : 0.0413648
  mean: 7.4202 < 8.2845 , median: 6 < 7
- P_Dw vs HIGH_PSI : 0.0129866
  mean: 7.4202 > 6.9061 , median: 6 = 6
- Bg_Upstrat vs HIGH_PSI : 4.80489e-07
  mean: 7.7311 > 6.9061 , median: 7 > 6
- Bg_Upstrat vs LOW_PSI : 0.00354274
  mean: 7.7311 > 7.198 , median: 7 > 6
- P_Up vs HIGH_PSI : 3.32698e-05
  mean: 8.2845 > 6.9061 , median: 7 > 6
- P_Up vs LOW_PSI : 0.00195233
  mean: 8.2845 > 7.198 , median: 7 > 6
- HIGH_PSI vs LOW_PSI : 0.000231589
  mean: 6.9061 < 7.198 , median: 6 = 6

#### 6.66 PROP FIRST EXON

Back to: Overview | ToC

Meaning: NTRS WITH EXON AS FIRST EXON / NTRS WITH EXON

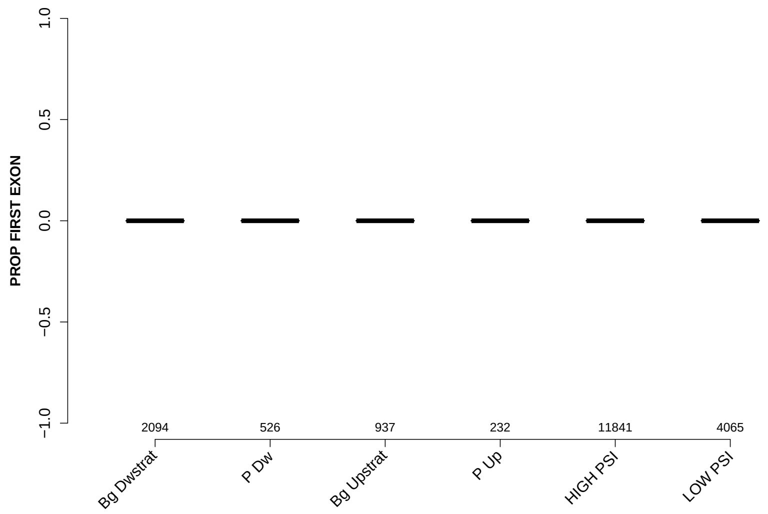

Significant results from Mann-Whitney U test:

- `Bg_Dwstrat` vs `Bg_Upstrat` : 0.00221551
  mean: 0.0197899 > 0.00999861 , median: 0 = 0
- `Bg_Dwstrat` vs `LOW_PSI` : 3.82425e-30
  mean: 0.0197899 > 0.00351649 , median: 0 = 0
- `P_Dw` vs `Bg_Upstrat` : 0.00375138
  mean: 0.0191315 > 0.00999861 , median: 0 = 0
- `P_Dw` vs `LOW_PSI` : 5.04245e-19
  mean: 0.0191315 > 0.00351649 , median: 0 = 0
- `Bg_Upstrat` vs `P_Up` : 0.000173506
  mean: 0.00999861 < 0.0225451 , median: 0 = 0
- `Bg_Upstrat` vs `HIGH_PSI` : 0.0030885
  mean: 0.00999861 < 0.0177753 , median: 0 = 0
- `Bg_Upstrat` vs `LOW_PSI` : 1.93575e-07
  mean: 0.00999861 > 0.00351649 , median: 0 = 0
- `P_Up` vs `HIGH_PSI` : 0.0232617
  mean: 0.0225451 > 0.0177753 , median: 0 = 0
- `P_Up` vs `LOW_PSI` : 1.0196e-19
  mean: 0.0225451 > 0.00351649 , median: 0 = 0
- `HIGH_PSI` vs `LOW_PSI` : 7.31012e-38
  mean: 0.0177753 > 0.00351649 , median: 0 = 0

#### 6.67 PROP LAST EXON

Back to: Overview | ToC

Meaning: NTRS WITH EXON AS LAST EXON / NTRS WITH EXON

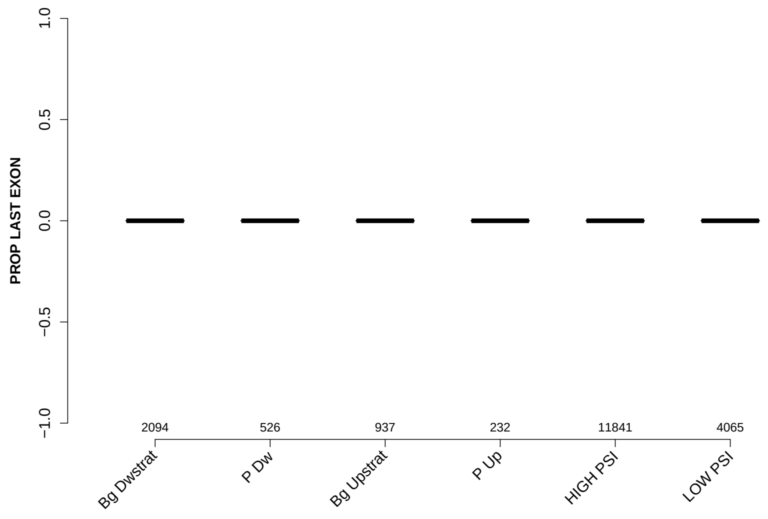

Significant results from Mann-Whitney U test:

- `Bg_Dwstrat` vs `Bg_Upstrat` : 4.58807e-12
  mean: 0.033325 > 0.0128693 , median: 0 = 0
- `Bg_Dwstrat` vs `P_Up` : 0.00240222
  mean: 0.033325 > 0.021764 , median: 0 = 0
- `Bg_Dwstrat` vs `LOW_PSI` : 4.92477e-74
  mean: 0.033325 > 0.00451818 , median: 0 = 0
- `P_Dw` vs `Bg_Upstrat` : 9.54065e-14
  mean: 0.0327417 > 0.0128693 , median: 0 = 0
- `P_Dw` vs `P_Up` : 0.000141715
  mean: 0.0327417 > 0.021764 , median: 0 = 0
- `P_Dw` vs `HIGH_PSI` : 0.0210016
  mean: 0.0327417 > 0.0309625 , median: 0 = 0
- `P_Dw` vs `LOW_PSI` : 3.65165e-65
  mean: 0.0327417 > 0.00451818 , median: 0 = 0
- `Bg_Upstrat` vs `HIGH_PSI` : 1.01612e-12
  mean: 0.0128693 < 0.0309625 , median: 0 = 0
- `Bg_Upstrat` vs `LOW_PSI` : 6.5755e-08
  mean: 0.0128693 > 0.00451818 , median: 0 = 0
- `P_Up` vs `HIGH_PSI` : 0.0032973
  mean: 0.021764 < 0.0309625 , median: 0 = 0

- `P_Up` vs `LOW_PSI` : 2.04083e-05
  mean: $0.021764 > 0.00451818$ , median: $0 = 0$
- `HIGH_PSI` vs `LOW_PSI` : 2.27052e-90
  mean: $0.0309625 > 0.00451818$ , median: $0 = 0$

#### 6.68 PROP INTERNAL EXON

Back to: Overview | ToC

Meaning: NTRS WITH EXON AS INTERNAL EXON / NTRS WITH EXON

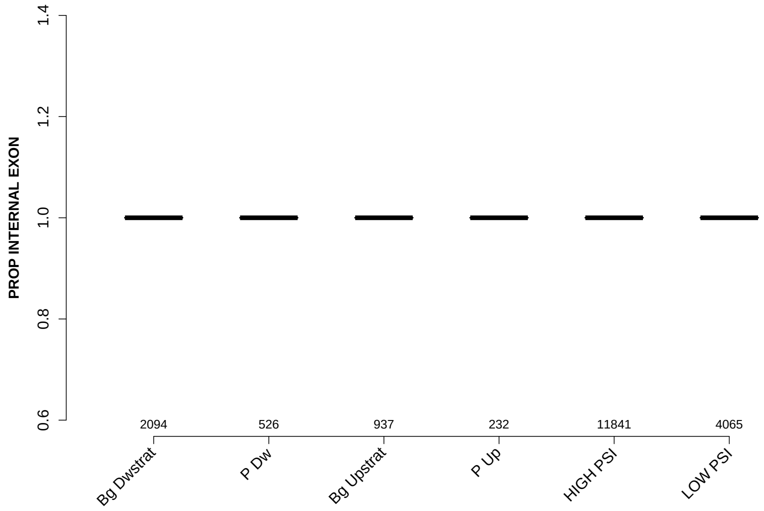

Significant results from Mann-Whitney U test:

- Bg_Dwstrat vs Bg_Upstrat : 1.53191e-12
  mean: 0.947068 < 0.977132 , median: 1 = 1
- Bg_Dwstrat vs LOW_PSI : 2.86505e-97
  mean: 0.947068 < 0.99199 , median: 1 = 1
- P_Dw vs Bg_Upstrat : 6.80182e-13
  mean: 0.948127 < 0.977132 , median: 1 = 1
- P_Dw vs P_Up : 0.0464182
  mean: 0.948127 < 0.957092 , median: 1 = 1
- P_Dw vs HIGH_PSI : 0.0212293
  mean: 0.948127 < 0.951303 , median: 1 = 1
- P_Dw vs LOW_PSI : 1.07819e-72
  mean: 0.948127 < 0.99199 , median: 1 = 1
- Bg_Upstrat vs P_Up : 0.00215498
  mean: 0.977132 > 0.957092 , median: 1 = 1
- Bg_Upstrat vs HIGH_PSI : 9.73823e-13
  mean: 0.977132 > 0.951303 , median: 1 = 1
- Bg_Upstrat vs LOW_PSI : 7.1825e-15
  mean: 0.977132 < 0.99199 , median: 1 = 1
- P_Up vs LOW_PSI : 1.51703e-20
  mean: 0.957092 < 0.99199 , median: 1 = 1

- `HIGH_PSI` vs `LOW_PSI` : 1.44062e-123
  mean: $0.951303 < 0.99199$ , median: $1 = 1$

#### 6.69 PROP EXON IN UTR

Back to: Overview | ToC

Meaning: NTRS WITH EXON IN UTR / NTRS WITH EXON

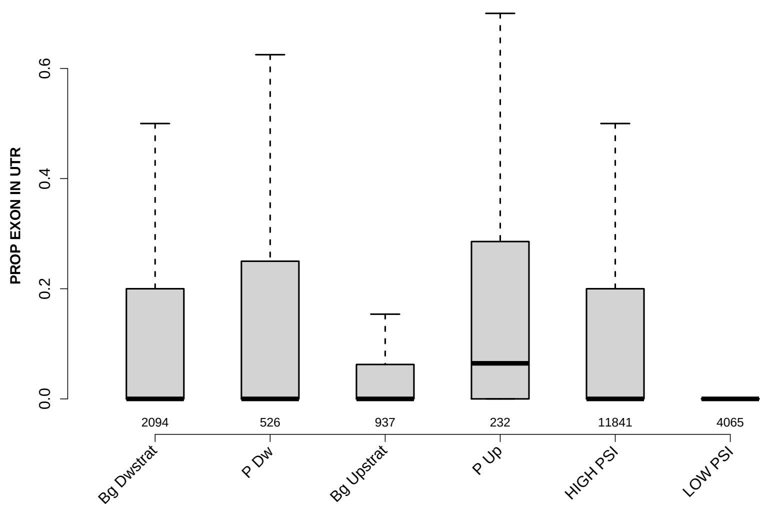

Significant results from Mann-Whitney U test:

- Bg_Dwstrat vs P_Dw : 0.0220437
  mean: 0.141785 < 0.158793 , median: 0 = 0
- Bg_Dwstrat vs Bg_Upstrat : 9.25465e-12
  mean: 0.141785 > 0.0701939 , median: 0 = 0
- Bg_Dwstrat vs P_Up : 0.000130885
  mean: 0.141785 < 0.166916 , median: 0 < 0.0645833
- Bg_Dwstrat vs LOW_PSI : 4.04938e-92
  mean: 0.141785 > 0.0335741 , median: 0 = 0
- P_Dw vs Bg_Upstrat : 2.09002e-13
  mean: 0.158793 > 0.0701939 , median: 0 = 0
- P_Dw vs HIGH_PSI : 0.00562079
  mean: 0.158793 > 0.137886 , median: 0 = 0
- P_Dw vs LOW_PSI : 6.81184e-59
  mean: 0.158793 > 0.0335741 , median: 0 = 0
- Bg_Upstrat vs P_Up : 8.20974e-16
  mean: 0.0701939 < 0.166916 , median: 0 < 0.0645833
- Bg_Upstrat vs HIGH_PSI : 1.20236e-13
  mean: 0.0701939 < 0.137886 , median: 0 = 0
- Bg_Upstrat vs LOW_PSI : 3.22815e-15
  mean: 0.0701939 > 0.0335741 , median: 0 = 0

- `P_Up` vs `HIGH_PSI` : 2.7311e-05
  mean: 0.166916 > 0.137886 , median: 0.0645833 > 0
- `P_Up` vs `LOW_PSI` : 7.0479e-51
  mean: 0.166916 > 0.0335741 , median: 0.0645833 > 0
- `HIGH_PSI` vs `LOW_PSI` : 4.46913e-153
  mean: 0.137886 > 0.0335741 , median: 0 = 0

### Comparison of exons grouped into: Bg-Dwstrat, P-Dw, Bg-Upstrat, P-Up, HIGH-PSI, LOW-PSI

September 16, 2020
Matt version 1.3.0

#### Contents

1 Infos 4

2 Warning: Please read this note carefully 4

3 Notes for publishing results 4

4 Data sets 5

5 Overview: Features with statistically significant differences (p-val $\leq 0.05$) 6

6 Details: Box plots and statistical assessments for all features 22

### 1 Infos

Visualizations of exon features for different groups of exons. Each exon occurs in exactly one gene, but might occur in several transcripts of that gene. Hence, for some features like the exon length, there is exactly one value for each exon. For other features, e.g., length of the up-stream exon(s), which could be different in different transcripts, there might be several values for each exon. Consequently, in the latter cases, the median of these value gets reported.

### 2 Warning: Please read this note carefully

Please keep in mind that some features might affect other features. Especially: all branch-point features get extracted from sub-sequences of introns, by standard the last 150 nt at the 3' end of each intron (if you haven't changed this) always neglecting the first 20 nt at their 5' end. If introns of one set are especially short, i.e., many are shorter then these 150 nt, then the shorter intron length might affect branch-point features. For example, there might be less branch points found in shorter introns or their distance to the 3' intron ends might be generally shorter simply because of their shorter intron length.

### 3 Notes for publishing results

The Matt paper: *Matt: Unix tools for alternative splicing analysis, A. Gohr, M. Irimia, Bioinformatics, 2018, bty606, DOI: 10.1093/bioinformatics/bty606*
When publishing results wrt. splice site strengths which you determined for your data using matt, please cite: *Maximum entropy modeling of short sequence motifs with applications to RNA splicing signals, Yeo et al., 2003, DOI: 10.1089/1066527041410418*
When publishing results wrt. branch point features which you determined for your data with matt, please cite: *Genome-wide association between branch point properties and alternative splicing, Corvelo et al., 2010, DOI: 10.1371/journal.pcbi.1001016*
When publishing results with respect to the binding strength of the human Sf1 splicing factor, you might refert to where the Sf1 binding motif comes from: *Analysis of in situ pre-mRNA targets of human splicing factor SF1 reveals a function in alternative splicing, Margherita Corioni, Nicolas Antih, Goranka Tanackovic, Mihaela Zavolan, and Angela Kramer, 2011, DOI: 10.1093/nar/gkq1042*
The Sf1 binding motif is described in supplement, page 13, table S2: Weight matrix of the binding specificity of SF1.

#### 4 Data sets

Input file:
../mmu_final.tab
Selection criteria for defining exons groups:
Bg_Dwstrat : having value Bg_Dwstrat in column GROUP
P_Dw : having value P_Dw in column GROUP
Bg_Upstrat : having value Bg_Upstrat in column GROUP
P_Up : having value P_Up in column GROUP
HIGH_PSI : having value HIGH_PSI in column GROUP
LOW_PSI : having value LOW_PSI in column GROUP
Exon duplicates removal: yes
Numbers of exons per group before / after neglecting exons which were not found in GTF file (gene annotation). For the comparisons only exons which were found in the gene annotation are used. These numbers might change slightly for each feature if NAs occur.
Bg_Dwstrat: 1224 / 1217
P_Dw: 345 / 344
Bg_Upstrat: 128 / 125
P_Up: 32 / 31
HIGH_PSI: 6768 / 6728
LOW_PSI: 1213 / 1206

#### 5 Overview: Features with statistically significant differences (p-val $\leq 0.05$)

##### MAXENTSCR HSAMODEL 5SS

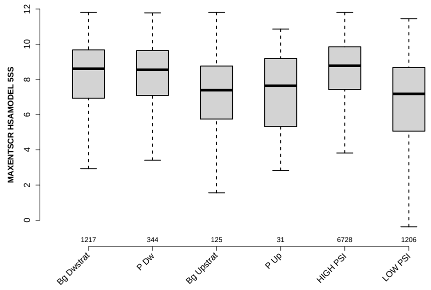

##### MAXENTSCR HSAMODEL 3SS

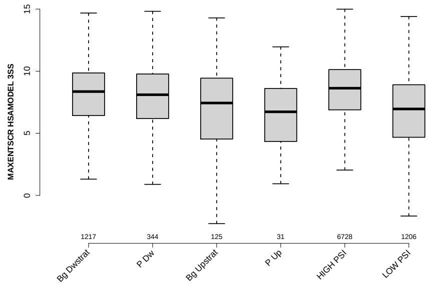

##### MEDIAN TR LENGTH

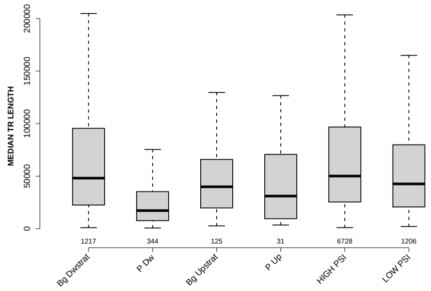

##### MEDIAN EXON NUMBER

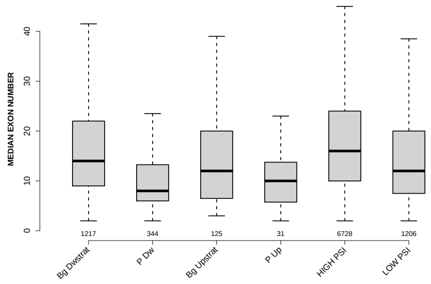

### PROP INTERNAL EXON

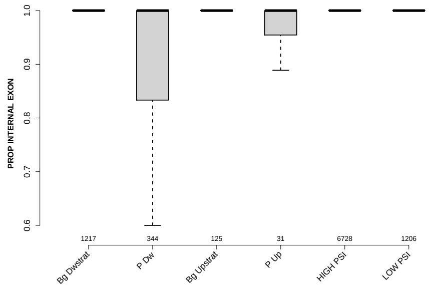

### UPINTRON GCC

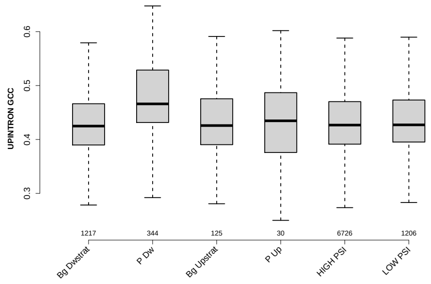

### EXON GCC

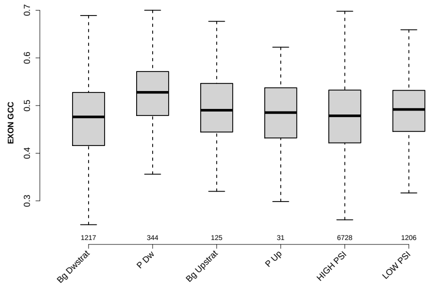

### PROP LAST EXON

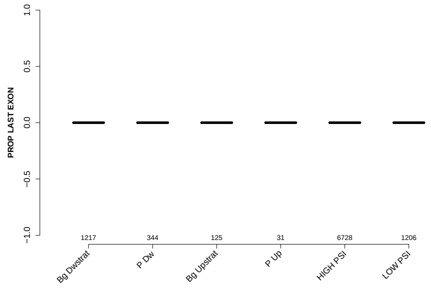

### DOINTRON GCC

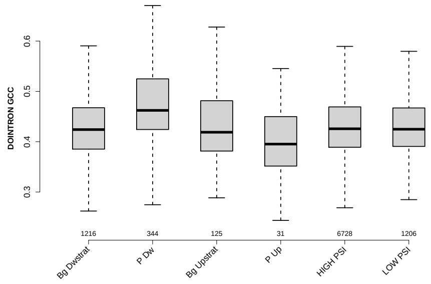

### GCC 3SS 20INT10EX

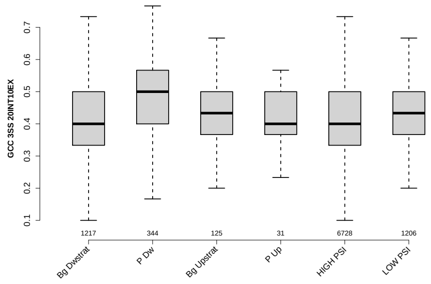

### EXON LENGTH

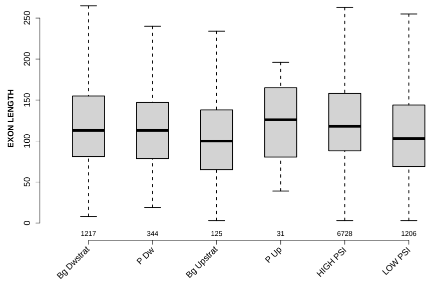

### NUM PREDICTED BPS UPINTRON

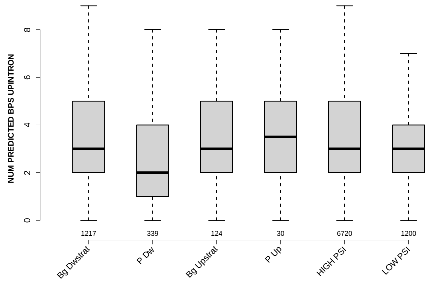

#### PYRIMIDINECONT MAXBP UPINTRON

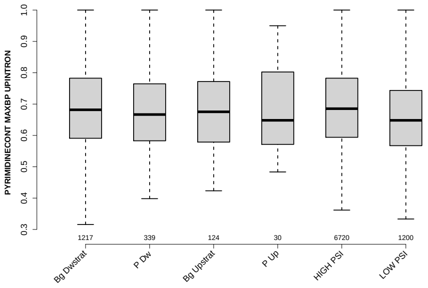

#### GCC 5SS 20INT10EX

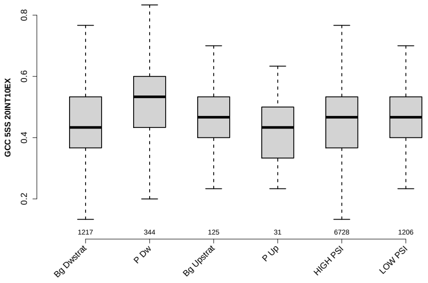

#### BPSCORE MAXBP UPINTRON

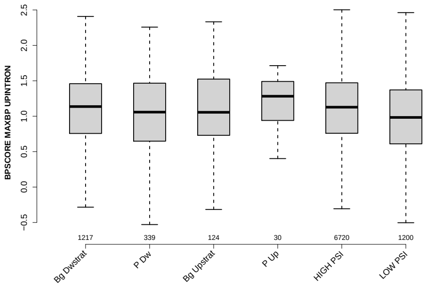

#### UP 5SS 20INT10EX GCC

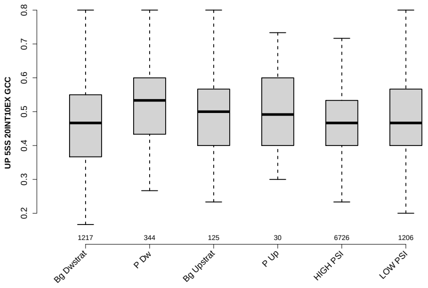

#### UPEXON GCC

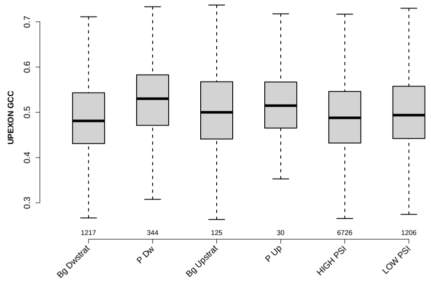

#### DOEXON GCC

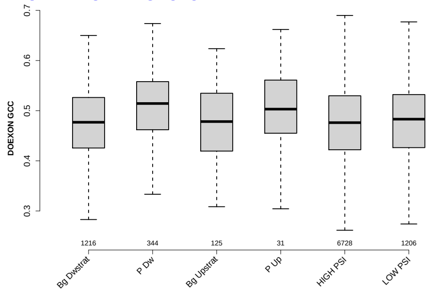

#### UPINTRON MEDIANLENGTH

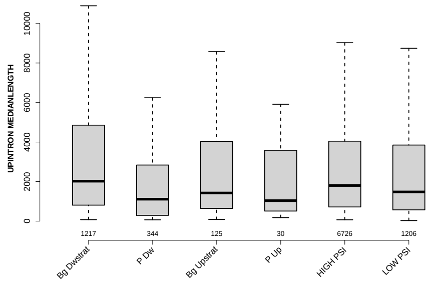

#### DO 3SS 20INT10EX GCC

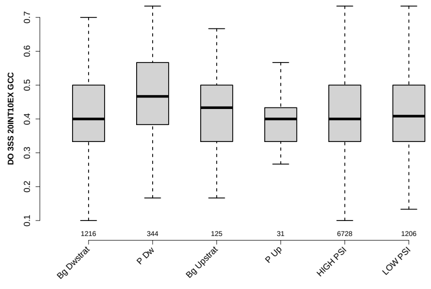

### PROP FIRST EXON

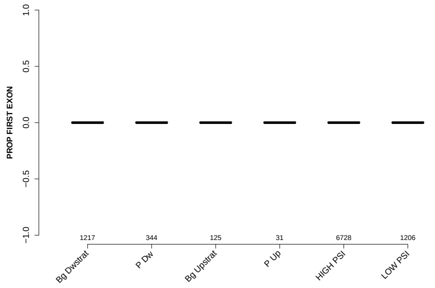

### MEDIAN BPSCORE UPINTRON

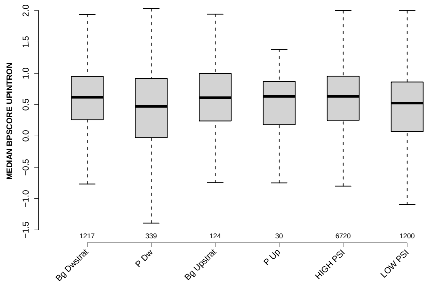

### RATIO UPEXON EXON LENGTH

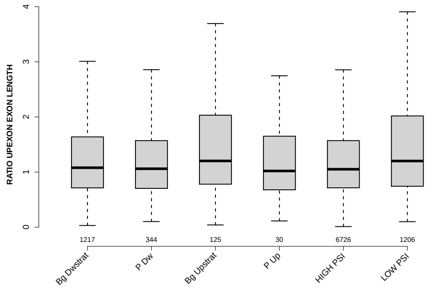

### RATIO UPINTRON EXON LENGTH

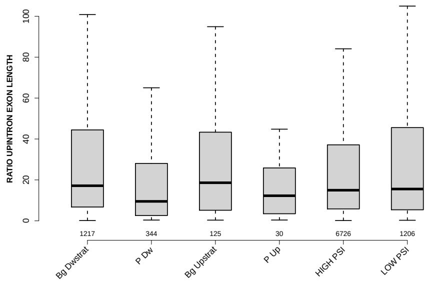

#### MEDIAN POLYPYRITRAC OFFSET UPINTRON

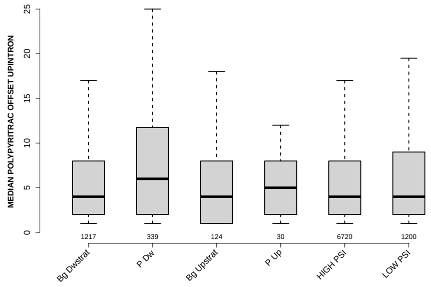

#### SCORE FOR MAXBP SEQ UPINTRON

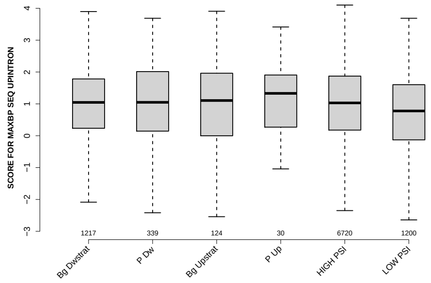

#### NUM PREDICTED BPS DOINTRON

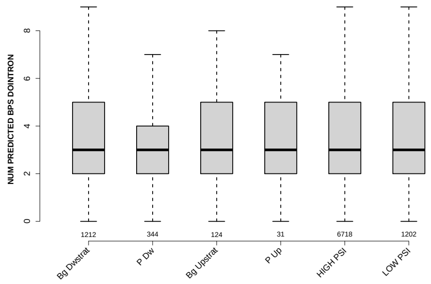

#### DIST FROM MAXBP TO 3SS UPINTRON

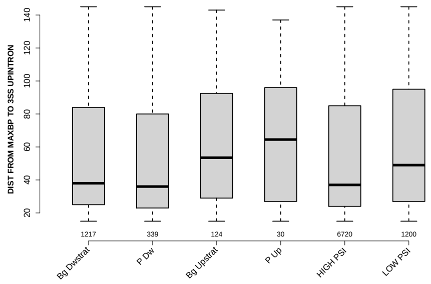

### DOINTRON MEDIANLENGTH

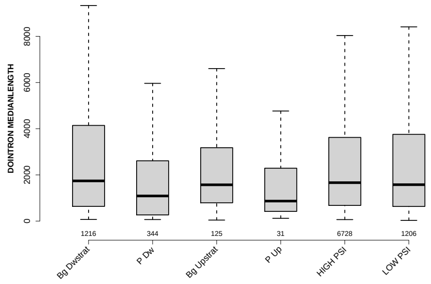

### RATIO DOINTRON EXON GCC

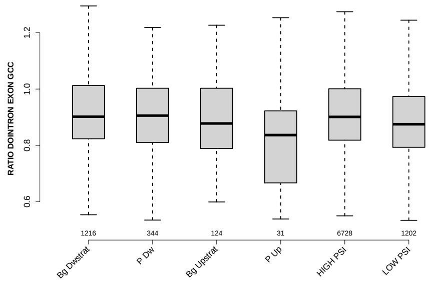

### RATIO DOINTRON EXON LENGTH

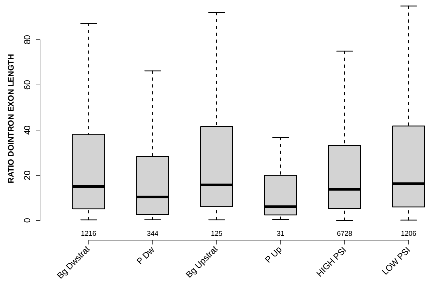

### MEDIAN SCORE FOR BPSEQ UPINTRON

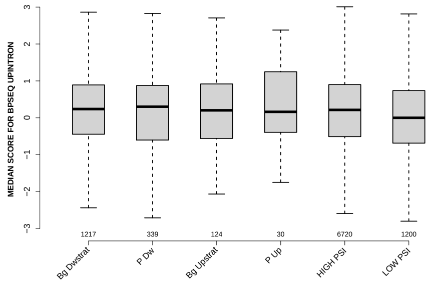

#### SF1 HIGHESTSCORE 3SS UPINTRON

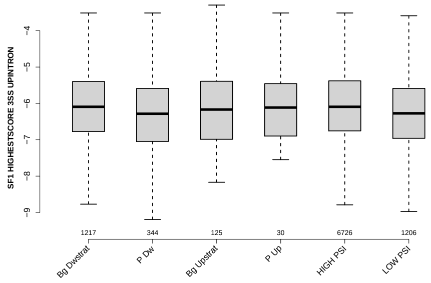

#### EXON MEDIANRELATIVERANK 3BINS

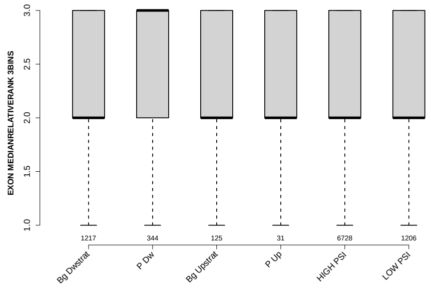

#### RATIO DOEXON EXON LENGTH

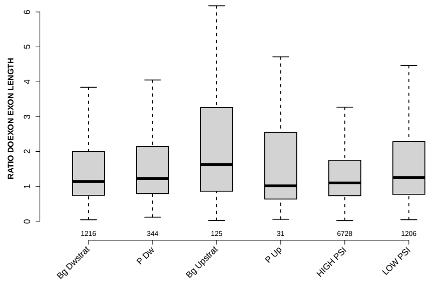

#### MEDIAN PYRIMIDINECONT UPINTRON

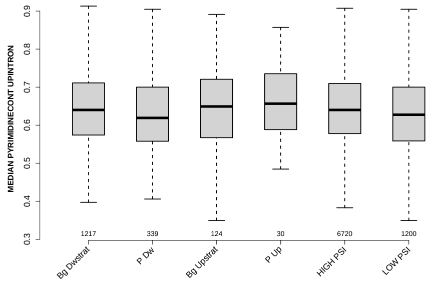

#### MEDIAN POLYPYRITRAC OFFSET DOINTRON

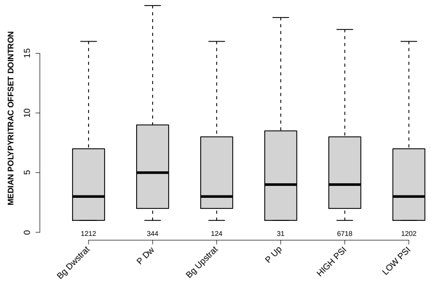

#### RATIO DOEXON EXON GCC

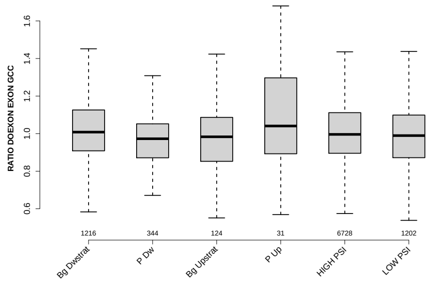

#### EXON MEDIANRELATIVERANK 10BINS

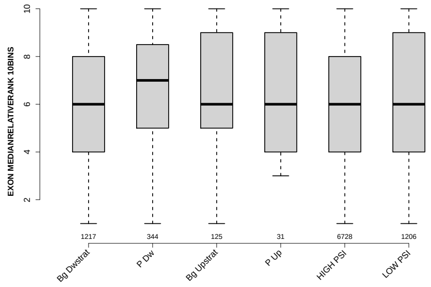

#### EXON MEDIANRELATIVERANK

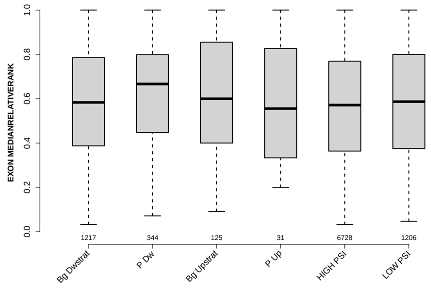

#### EXON MEDIANRELATIVERANK 5BINS

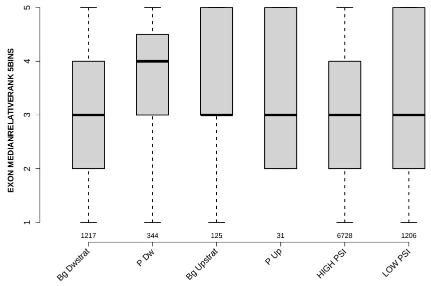

#### POLYPYRITRAC OFFSET MAXBP UPINTRON

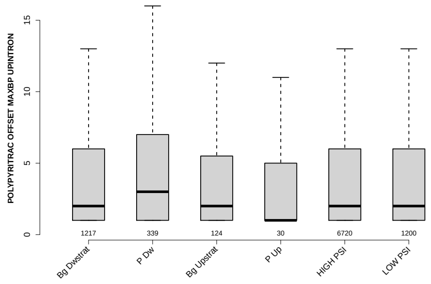

#### RATIO UPINTRON EXON GCC

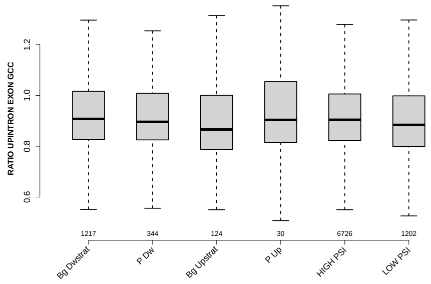

#### NTRS ALL FOR GENE

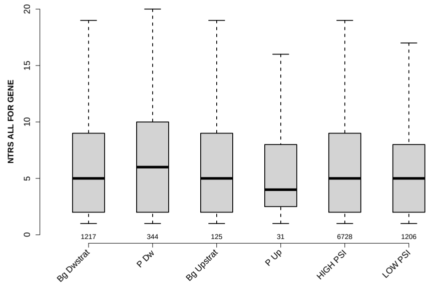

#### POLYPYRITRAC SCORE MAXBP UPINTRON

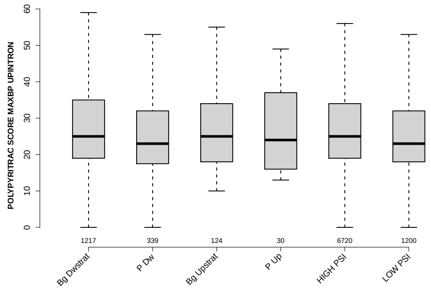

#### MEDIAN DIST FROM BP TO 3SS UPINTRON

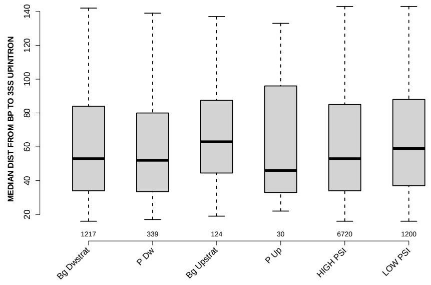

#### UPEXON MEDIANLENGTH

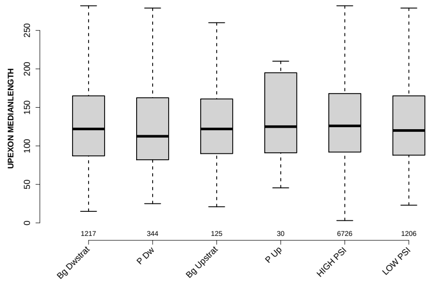

#### POLYPYRITRAC OFFSET MAXBP DOINTRON

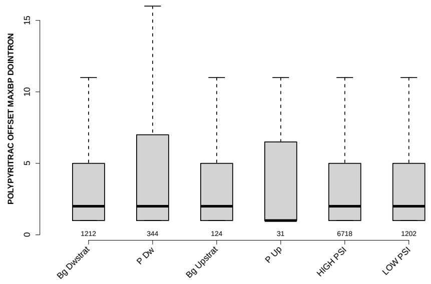

### MAXENTSCR HSAMODEL DOWNSTRM 3SS

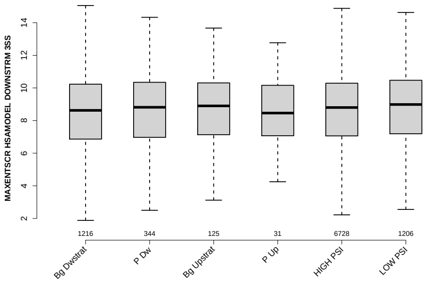

### POLYPYRITRAC LEN MAXBP UPINTRON

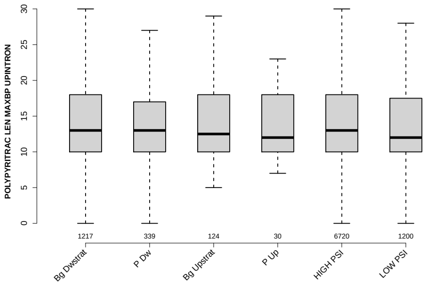

### MEDIAN POLYPYRITRAC LEN DOINTRON

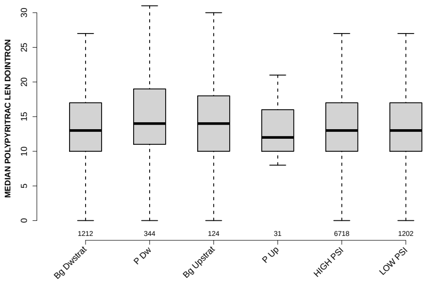

### MAXENTSCR HSAMODEL UPSTRM 5SS

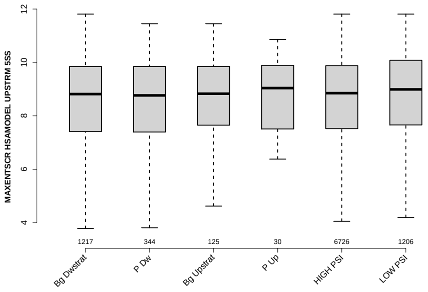

### DOEXON MEDIANLENGTH

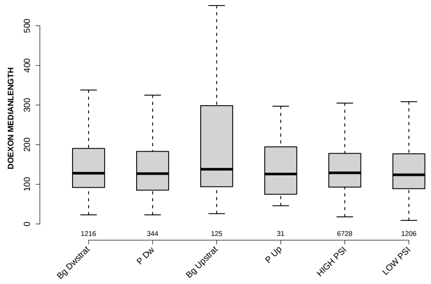

### SF1 HIGHESTSCORE 3SS DOINTRON

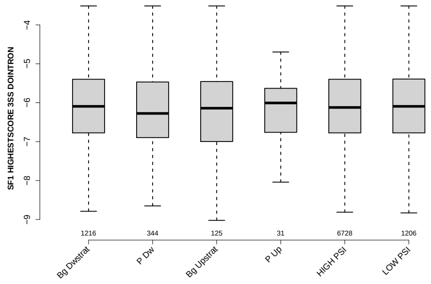

### POLYPYRITRAC LEN MAXBP DOINTRON

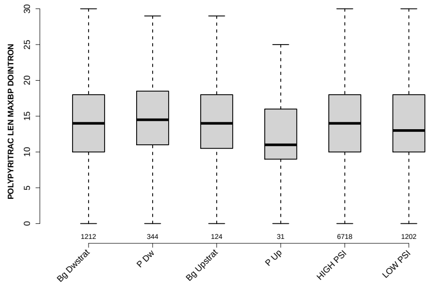

### MEDIAN BPSCORE DOINTRON

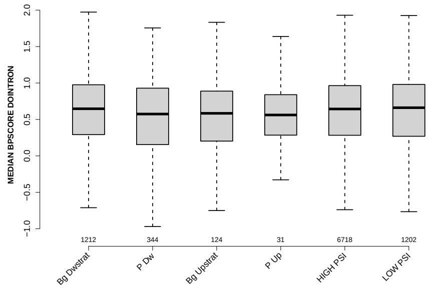

#### MEDIAN POLYPYRITRAC SCORE UPINTRON

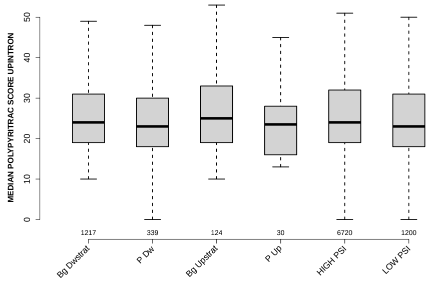

#### RATIO UPEXON EXON GCC

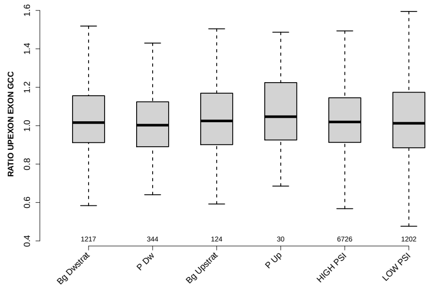

#### PYRIMIDINECONT MAXBP DOINTRON

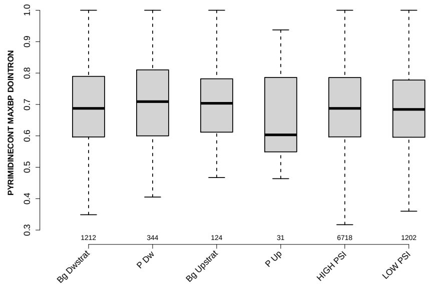

#### DIST FROM MAXBP TO 3SS DOINTRON

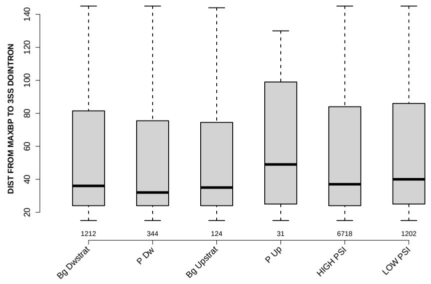

#### MEDIAN SCORE FOR BPSEQ DOINTRON

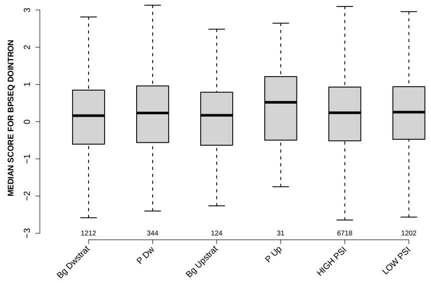

### 6 Details: Box plots and statistical assessments for all features

#### 6.1 EXON LENGTH

Back to: Overview | ToC

Meaning:

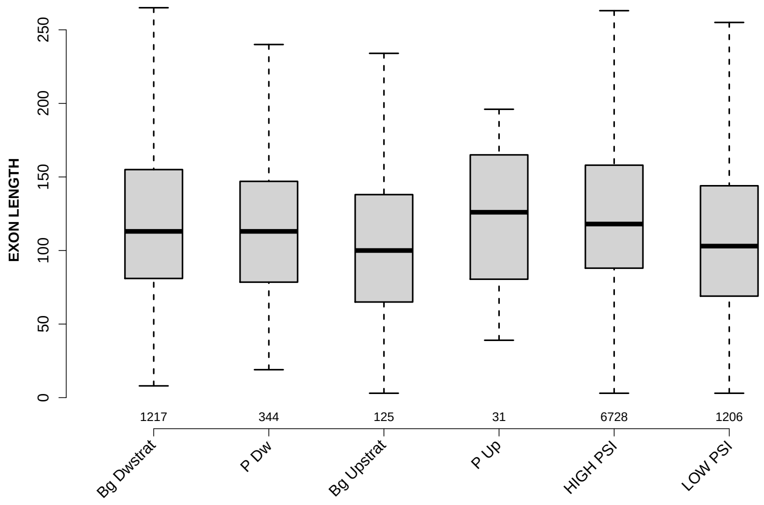

Significant results from Mann-Whitney U test:

- `Bg_Dwstrat` vs `Bg_Upstrat` : 0.00686808
  mean: 132.0205 < 153.48 , median: 113 > 100
- `Bg_Dwstrat` vs `HIGH_PSI` : 0.00362747
  mean: 132.0205 < 137.8043 , median: 113 < 118
- `Bg_Dwstrat` vs `LOW_PSI` : 2.17464e-07
  mean: 132.0205 > 114.8308 , median: 113 > 103
- `P_Dw` vs `HIGH_PSI` : 0.00394084
  mean: 119.311 < 137.8043 , median: 113 < 118
- `P_Dw` vs `LOW_PSI` : 0.0154909
  mean: 119.311 > 114.8308 , median: 113 > 103
- `Bg_Upstrat` vs `HIGH_PSI` : 0.000182856
  mean: 153.48 > 137.8043 , median: 100 < 118
- `HIGH_PSI` vs `LOW_PSI` : 3.70467e-21
  mean: 137.8043 > 114.8308 , median: 118 > 103

#### 6.2 UPEXON MEDIANLENGTH

Back to: Overview | ToC

Meaning: median length of up-stream exon

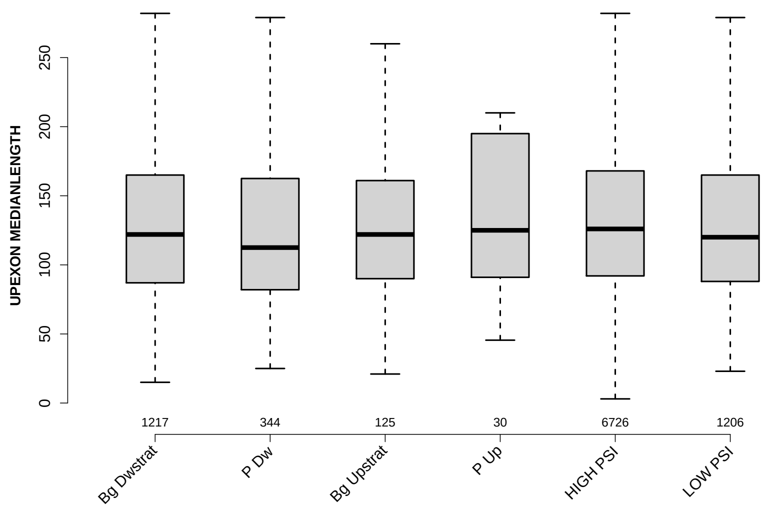

Significant results from Mann-Whitney U test:

- `Bg_Dwstrat` vs `HIGH_PSI` : 0.0358523
  mean: 162.3209 > 155.0394 , median: 122 < 126
- `P_Dw` vs `HIGH_PSI` : 0.00122724
  mean: 131.6061 < 155.0394 , median: 112.5 < 126
- `HIGH_PSI` vs `LOW_PSI` : 0.0329745
  mean: 155.0394 < 164.8964 , median: 126 > 120

#### 6.3 DOEXON MEDIANLENGTH

Back to: Overview | ToC

Meaning: median length of down-stream exon

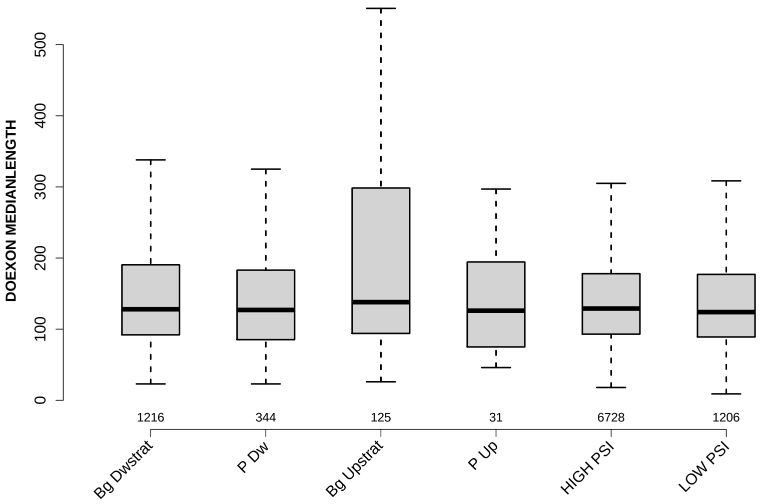

Significant results from Mann-Whitney U test:

- `Bg_Upstrat` vs `HIGH_PSI` : 0.0470757
  mean: 283.48 > 265.4267 , median: 138 > 129
- `Bg_Upstrat` vs `LOW_PSI` : 0.0118354
  mean: 283.48 > 236.694 , median: 138 > 124
- `HIGH_PSI` vs `LOW_PSI` : 0.0497438
  mean: 265.4267 > 236.694 , median: 129 > 124

#### 6.4 RATIO UPEXON EXON LENGTH

Back to: Overview | ToC

Meaning: median up-stream exon length / exon length

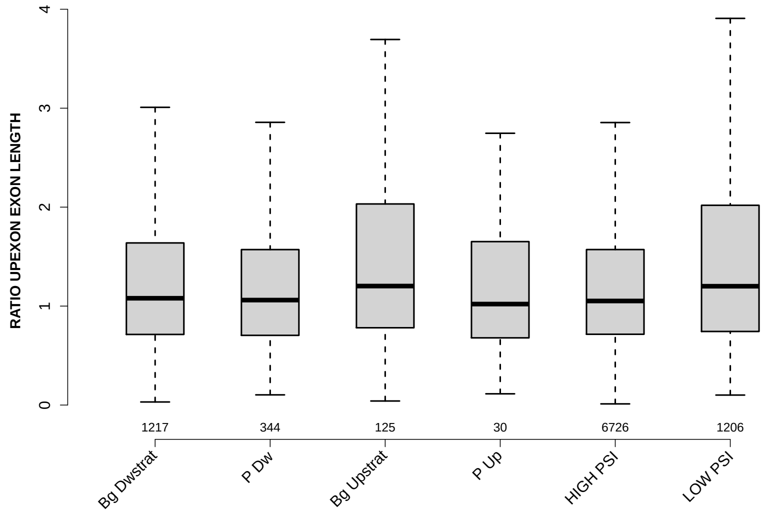

Significant results from Mann-Whitney U test:

- `Bg_Dwstrat` vs `Bg_Upstrat` : 0.0462738
  mean: 1.5712 < 1.8403 , median: 1.0794 < 1.2026
- `Bg_Dwstrat` vs `LOW_PSI` : 2.29245e-05
  mean: 1.5712 < 2.7547 , median: 1.0794 < 1.2009
- `P_Dw` vs `Bg_Upstrat` : 0.028894
  mean: 1.332 < 1.8403 , median: 1.0608 < 1.2026
- `P_Dw` vs `LOW_PSI` : 0.000551896
  mean: 1.332 < 2.7547 , median: 1.0608 < 1.2009
- `Bg_Upstrat` vs `HIGH_PSI` : 0.0133481
  mean: 1.8403 > 1.4109 , median: 1.2026 > 1.0518
- `HIGH_PSI` vs `LOW_PSI` : 3.67068e-11
  mean: 1.4109 < 2.7547 , median: 1.0518 < 1.2009

#### 6.5 RATIO DOEXON EXON LENGTH

Back to: Overview | ToC

Meaning: median down-stream exon length / exon length

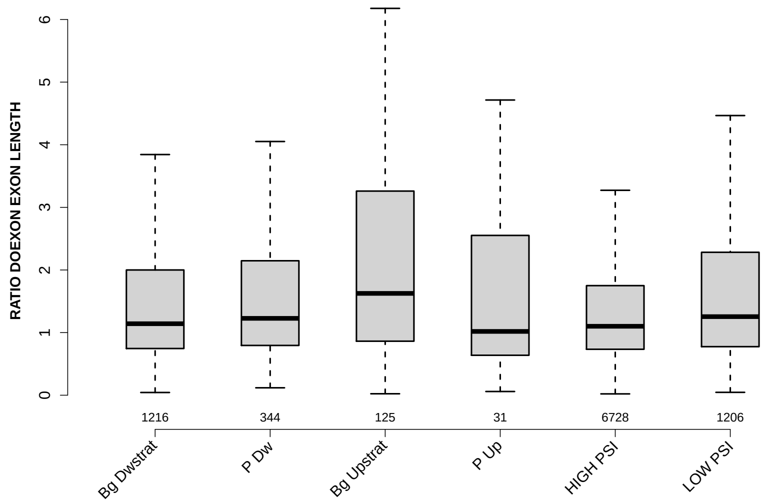

Significant results from Mann-Whitney U test:

- Bg_Dwstrat vs Bg_Upstrat : 0.000844486
  mean: 2.5917 < 3.7439 , median: 1.1419 < 1.626
- Bg_Dwstrat vs HIGH_PSI : 0.0378248
  mean: 2.5917 > 2.4376 , median: 1.1419 > 1.1011
- Bg_Dwstrat vs LOW_PSI : 0.0138136
  mean: 2.5917 < 6.1955 , median: 1.1419 < 1.255
- P_Dw vs Bg_Upstrat : 0.0139676
  mean: 2.6228 < 3.7439 , median: 1.2275 < 1.626
- P_Dw vs HIGH_PSI : 0.0197784
  mean: 2.6228 > 2.4376 , median: 1.2275 > 1.1011
- Bg_Upstrat vs HIGH_PSI : 2.74769e-05
  mean: 3.7439 > 2.4376 , median: 1.626 > 1.1011
- Bg_Upstrat vs LOW_PSI : 0.0232852
  mean: 3.7439 < 6.1955 , median: 1.626 > 1.255
- HIGH_PSI vs LOW_PSI : 9.86499e-08
  mean: 2.4376 < 6.1955 , median: 1.1011 < 1.255

#### 6.6 UPINTRON MEDIANLENGTH

Back to: Overview | ToC

Meaning: median length of up-stream introns

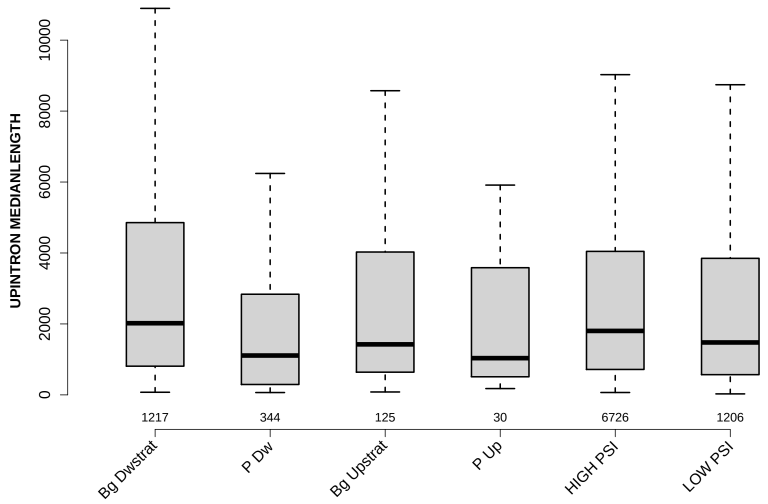

Significant results from Mann-Whitney U test:

- Bg_Dwstrat vs P_Dw : 5.59907e-14
  mean: 5749.1056 > 2636.327 , median: 2020 > 1109.5
- Bg_Dwstrat vs HIGH_PSI : 0.005115
  mean: 5749.1056 > 5051.1353 , median: 2020 > 1803
- Bg_Dwstrat vs LOW_PSI : 1.49461e-06
  mean: 5749.1056 > 4150.5609 , median: 2020 > 1477.5
- P_Dw vs Bg_Upstrat : 0.00243116
  mean: 2636.327 < 3701.156 , median: 1109.5 < 1424
- P_Dw vs HIGH_PSI : 4.08731e-12
  mean: 2636.327 < 5051.1353 , median: 1109.5 < 1803
- P_Dw vs LOW_PSI : 5.34213e-06
  mean: 2636.327 < 4150.5609 , median: 1109.5 < 1477.5
- HIGH_PSI vs LOW_PSI : 0.000256596
  mean: 5051.1353 > 4150.5609 , median: 1803 > 1477.5

#### 6.7 DOINTRON MEDIANLENGTH

Back to: Overview | ToC

Meaning: median length of down-stream introns

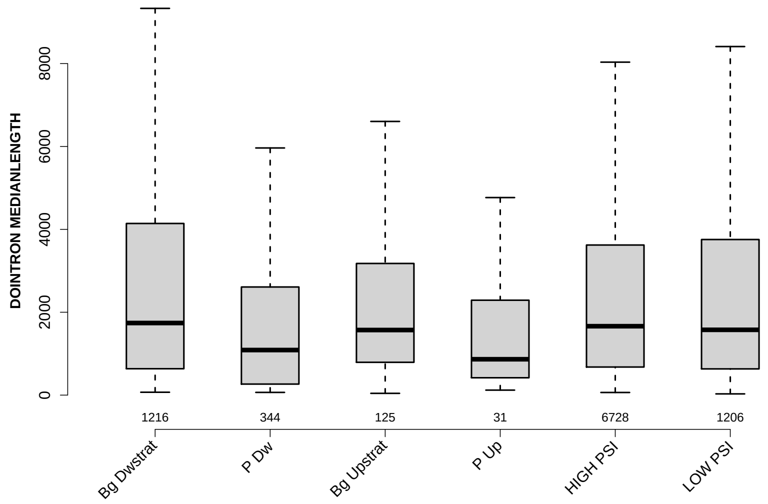

Significant results from Mann-Whitney U test:

- `Bg_Dwstrat` vs `P_Dw` : 3.2563e-08
  mean: 4484.801 > 2817.2849 , median: 1740 > 1088
- `Bg_Dwstrat` vs `P_Up` : 0.0485837
  mean: 4484.801 > 3523.3871 , median: 1740 > 866
- `P_Dw` vs `Bg_Upstrat` : 0.0047812
  mean: 2817.2849 < 3036.008 , median: 1088 < 1571
- `P_Dw` vs `HIGH_PSI` : 5.61988e-09
  mean: 2817.2849 < 4219.6223 , median: 1088 < 1663
- `P_Dw` vs `LOW_PSI` : 4.65742e-07
  mean: 2817.2849 < 4549.9992 , median: 1088 < 1576

#### 6.8 RATIO UPINTRON EXON LENGTH

Back to: Overview | ToC

Meaning: median up-stream intron length / exon length

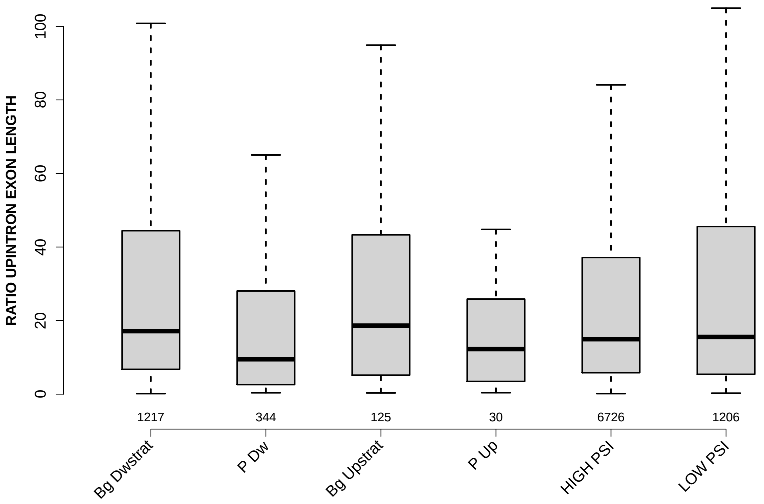

Significant results from Mann-Whitney U test:

- `Bg_Dwstrat` vs `P_Dw` : 5.01453e-11
  mean: 59.9863 > 26.0546 , median: 17.1538 > 9.5056
- `Bg_Dwstrat` vs `P_Up` : 0.0402708
  mean: 59.9863 > 23.9071 , median: 17.1538 > 12.267
- `Bg_Dwstrat` vs `HIGH_PSI` : 0.000442057
  mean: 59.9863 > 48.2685 , median: 17.1538 > 14.9615
- `P_Dw` vs `Bg_Upstrat` : 0.000762116
  mean: 26.0546 < 46.7155 , median: 9.5056 < 18.619
- `P_Dw` vs `HIGH_PSI` : 3.76185e-08
  mean: 26.0546 < 48.2685 , median: 9.5056 < 14.9615
- `P_Dw` vs `LOW_PSI` : 3.33645e-08
  mean: 26.0546 < 60.4703 , median: 9.5056 < 15.5511

#### 6.9 RATIO DOINTRON EXON LENGTH

Back to: Overview | ToC

Meaning: median down-stream intron length / exon length

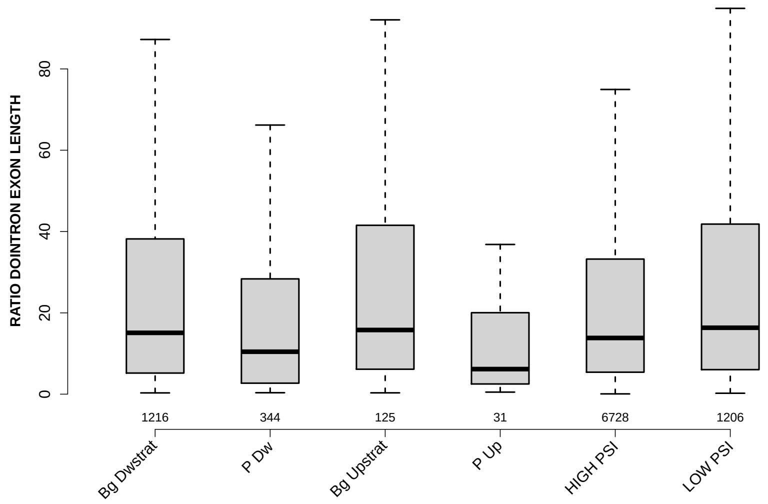

Significant results from Mann-Whitney U test:

- Bg_Dwstrat vs P_Dw : 6.43123e-06
  mean: 43.4873 > 27.0845 , median: 15.0887 > 10.4412
- Bg_Dwstrat vs P_Up : 0.0218955
  mean: 43.4873 > 27.0137 , median: 15.0887 > 6.1667
- Bg_Dwstrat vs LOW_PSI : 0.0416203
  mean: 43.4873 < 66.6507 , median: 15.0887 < 16.3362
- P_Dw vs Bg_Upstrat : 0.00381491
  mean: 27.0845 < 36.8493 , median: 10.4412 < 15.7863
- P_Dw vs HIGH_PSI : 2.25624e-05
  mean: 27.0845 < 39.4243 , median: 10.4412 < 13.8232
- P_Dw vs LOW_PSI : 8.42128e-09
  mean: 27.0845 < 66.6507 , median: 10.4412 < 16.3362
- Bg_Upstrat vs P_Up : 0.0224421
  mean: 36.8493 > 27.0137 , median: 15.7863 > 6.1667
- P_Up vs HIGH_PSI : 0.0275933
  mean: 27.0137 < 39.4243 , median: 6.1667 < 13.8232
- P_Up vs LOW_PSI : 0.00680335
  mean: 27.0137 < 66.6507 , median: 6.1667 < 16.3362
- HIGH_PSI vs LOW_PSI : 2.8367e-05
  mean: 39.4243 < 66.6507 , median: 13.8232 < 16.3362

#### 6.10 EXON GCC

Back to: Overview | ToC

Meaning: GC content of entire exon sequence

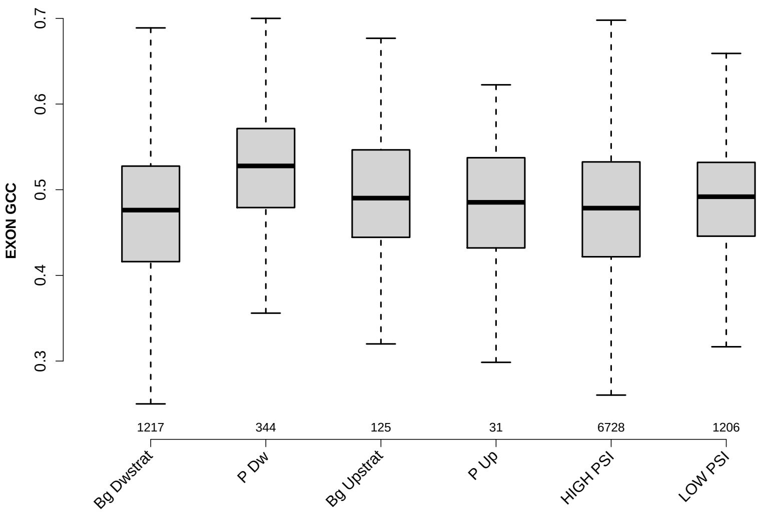

Significant results from Mann-Whitney U test:

- Bg_Dwstrat vs P_Dw : 2.95074e-26
  mean: 0.472243 < 0.524157 , median: 0.47619 < 0.527778
- Bg_Dwstrat vs Bg_Upstrat : 0.0155803
  mean: 0.472243 < 0.489795 , median: 0.47619 < 0.490196
- Bg_Dwstrat vs LOW_PSI : 4.69155e-07
  mean: 0.472243 < 0.486056 , median: 0.47619 < 0.491837
- P_Dw vs Bg_Upstrat : 4.95362e-05
  mean: 0.524157 > 0.489795 , median: 0.527778 > 0.490196
- P_Dw vs P_Up : 0.00610649
  mean: 0.524157 > 0.485194 , median: 0.527778 > 0.485294
- P_Dw vs HIGH_PSI : 9.85827e-28
  mean: 0.524157 > 0.47693 , median: 0.527778 > 0.478511
- P_Dw vs LOW_PSI : 8.79797e-17
  mean: 0.524157 > 0.486056 , median: 0.527778 > 0.491837
- HIGH_PSI vs LOW_PSI : 1.88356e-06
  mean: 0.47693 < 0.486056 , median: 0.478511 < 0.491837

#### 6.11 UPINTRON GCC

Back to: Overview | ToC

Meaning: GC content of entire up-stream intron sequence

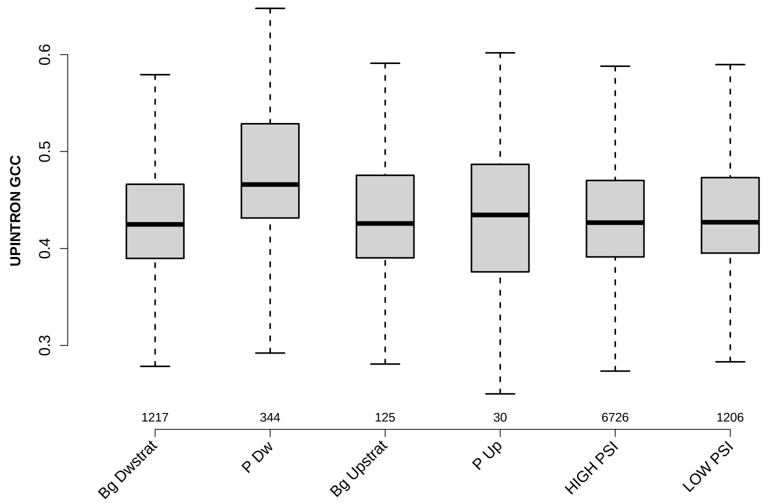

Significant results from Mann-Whitney U test:

- Bg_Dwstrat vs P_Dw : 5.88161e-28
  mean: 0.431295 < 0.479933 , median: 0.42485 < 0.466
- Bg_Dwstrat vs LOW_PSI : 0.0465734
  mean: 0.431295 < 0.437047 , median: 0.42485 < 0.427089
- P_Dw vs Bg_Upstrat : 5.58226e-08
  mean: 0.479933 > 0.440527 , median: 0.466 > 0.425815
- P_Dw vs P_Up : 0.0187773
  mean: 0.479933 > 0.445296 , median: 0.466 > 0.434616
- P_Dw vs HIGH_PSI : 3.13112e-30
  mean: 0.479933 > 0.433915 , median: 0.466 > 0.426699
- P_Dw vs LOW_PSI : 8.14504e-23
  mean: 0.479933 > 0.437047 , median: 0.466 > 0.427089

#### 6.12 UPEXON GCC

Back to: Overview | ToC

Meaning: GC content of entire up-stream exon sequence

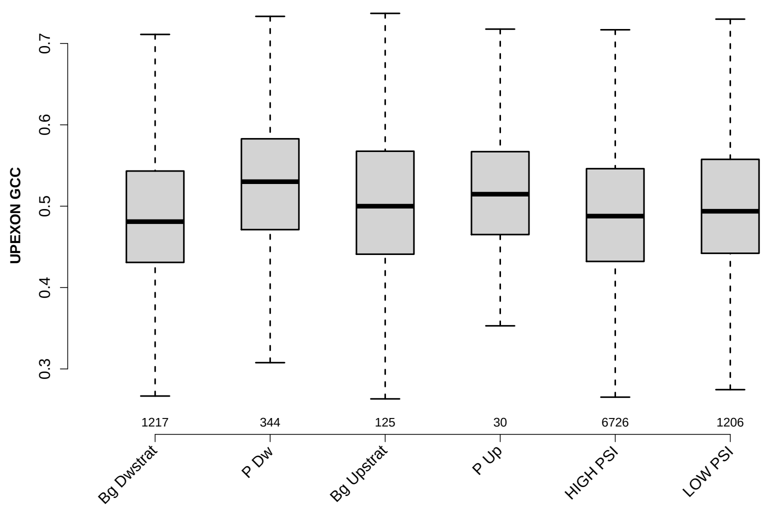

Significant results from Mann-Whitney U test:

- Bg_Dwstrat vs P_Dw : 2.06475e-14
  mean: 0.488876 < 0.526615 , median: 0.481013 < 0.530128
- Bg_Dwstrat vs Bg_Upstrat : 0.0164687
  mean: 0.488876 < 0.510481 , median: 0.481013 < 0.5
- Bg_Dwstrat vs LOW_PSI : 0.00015066
  mean: 0.488876 < 0.502655 , median: 0.481013 < 0.493768
- P_Dw vs Bg_Upstrat : 0.0409606
  mean: 0.526615 > 0.510481 , median: 0.530128 > 0.5
- P_Dw vs HIGH_PSI : 1.39728e-14
  mean: 0.526615 > 0.492204 , median: 0.530128 > 0.487805
- P_Dw vs LOW_PSI : 5.55183e-07
  mean: 0.526615 > 0.502655 , median: 0.530128 > 0.493768
- Bg_Upstrat vs HIGH_PSI : 0.0459568
  mean: 0.510481 > 0.492204 , median: 0.5 > 0.487805
- HIGH_PSI vs LOW_PSI : 0.000590232
  mean: 0.492204 < 0.502655 , median: 0.487805 < 0.493768

#### 6.13 DOINTRON GCC

Back to: Overview | ToC

Meaning: GC content of entire down-stream intron sequence

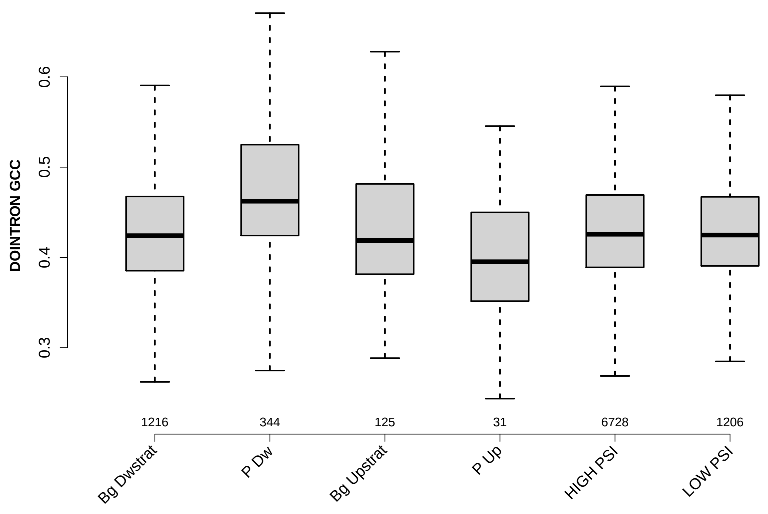

Significant results from Mann-Whitney U test:

- `Bg_Dwstrat` vs `P_Dw` : 1.05831e-23
  mean: 0.429738 < 0.474312 , median: 0.424089 < 0.462267
- `Bg_Dwstrat` vs `P_Up` : 0.0118724
  mean: 0.429738 > 0.394255 , median: 0.424089 > 0.395217
- `P_Dw` vs `Bg_Upstrat` : 4.38856e-08
  mean: 0.474312 > 0.434127 , median: 0.462267 > 0.41882
- `P_Dw` vs `P_Up` : 3.2016e-07
  mean: 0.474312 > 0.394255 , median: 0.462267 > 0.395217
- `P_Dw` vs `HIGH_PSI` : 8.98107e-26
  mean: 0.474312 > 0.431952 , median: 0.462267 > 0.425689
- `P_Dw` vs `LOW_PSI` : 6.6171e-24
  mean: 0.474312 > 0.429031 , median: 0.462267 > 0.424801
- `Bg_Upstrat` vs `P_Up` : 0.0229708
  mean: 0.434127 > 0.394255 , median: 0.41882 > 0.395217
- `P_Up` vs `HIGH_PSI` : 0.00574162
  mean: 0.394255 < 0.431952 , median: 0.395217 < 0.425689
- `P_Up` vs `LOW_PSI` : 0.00674102
  mean: 0.394255 < 0.429031 , median: 0.395217 < 0.424801

#### 6.14 DOEXON GCC

Back to: Overview | ToC

Meaning: GC content of entire down-stream exon sequence

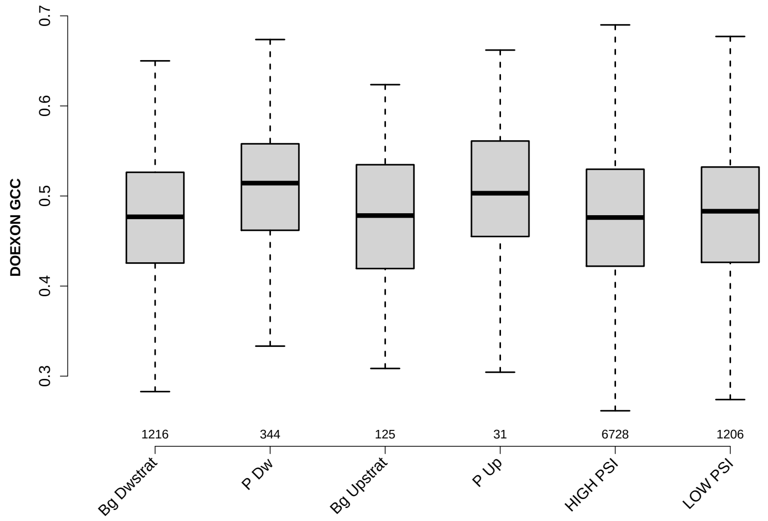

Significant results from Mann-Whitney U test:

- `Bg_Dwstrat` vs `P_Dw` : 2.5695e-12
  mean: 0.475839 < 0.506085 , median: 0.476864 < 0.514286
- `P_Dw` vs `Bg_Upstrat` : 6.89199e-05
  mean: 0.506085 > 0.475174 , median: 0.514286 > 0.478261
- `P_Dw` vs `HIGH_PSI` : 3.62938e-14
  mean: 0.506085 > 0.475597 , median: 0.514286 > 0.476103
- `P_Dw` vs `LOW_PSI` : 1.68564e-09
  mean: 0.506085 > 0.479005 , median: 0.514286 > 0.483051

#### 6.15 RATIO UPEXON EXON GCC

Back to: Overview | ToC

Meaning: UPEXON GCC / EXON GCC

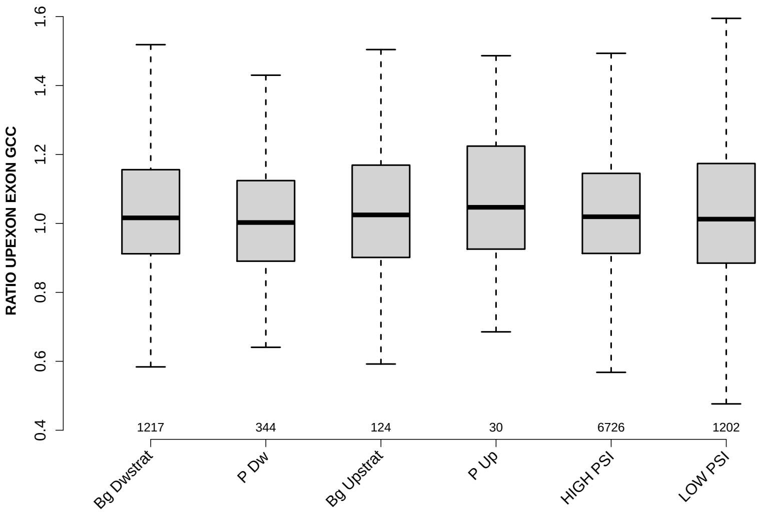

Significant results from Mann-Whitney U test:

- `Bg_Dwstrat` vs `P_Dw` : 0.0404056
  mean: 1.0559 > 1.0174 , median: 1.0161 > 1.0026
- `P_Dw` vs `HIGH_PSI` : 0.0285583
  mean: 1.0174 < 1.0494 , median: 1.0026 < 1.0194

#### 6.16 RATIO UPINTRON EXON GCC

Back to: Overview | ToC

Meaning: UPINTRON GCC / EXON GCC

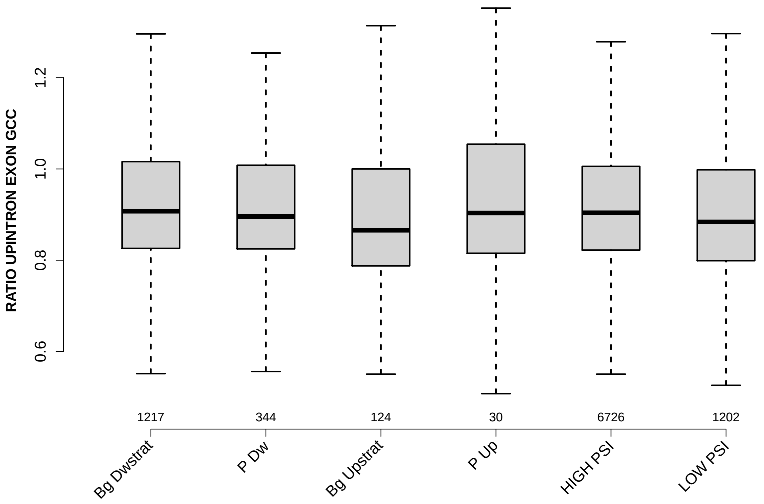

Significant results from Mann-Whitney U test:

- `Bg_Dwstrat` vs `Bg_Upstrat` : 0.0426782
  mean: 0.927892 > 0.908859 , median: 0.907448 > 0.86574
- `Bg_Dwstrat` vs `LOW_PSI` : 0.000172028
  mean: 0.927892 > 0.912411 , median: 0.907448 > 0.883916
- `P_Dw` vs `LOW_PSI` : 0.0284233
  mean: 0.923844 > 0.912411 , median: 0.895865 > 0.883916
- `HIGH_PSI` vs `LOW_PSI` : 9.70677e-05
  mean: 0.922923 > 0.912411 , median: 0.903951 > 0.883916

#### 6.17 RATIO DOINTRON EXON GCC

Back to: Overview | ToC

Meaning: DOINTRON GCC / EXON GCC

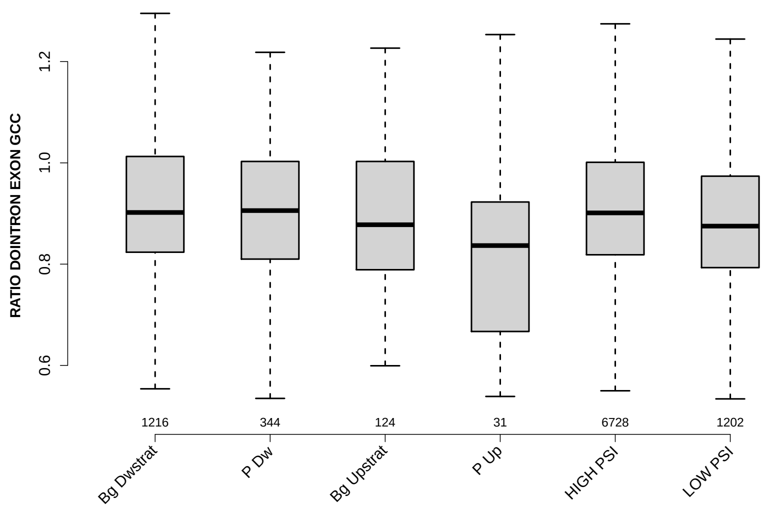

Significant results from Mann-Whitney U test:

- Bg_Dwstrat vs Bg_Upstrat : 0.0244567
  mean: 0.924926 > 0.893267 , median: 0.90201 > 0.877805
- Bg_Dwstrat vs P_Up : 0.00234479
  mean: 0.924926 > 0.828765 , median: 0.90201 > 0.83667
- Bg_Dwstrat vs LOW_PSI : 1.22195e-07
  mean: 0.924926 > 0.895832 , median: 0.90201 > 0.875075
- P_Dw vs P_Up : 0.00602688
  mean: 0.91325 > 0.828765 , median: 0.905755 > 0.83667
- P_Dw vs LOW_PSI : 0.00581276
  mean: 0.91325 > 0.895832 , median: 0.905755 > 0.875075
- P_Up vs HIGH_PSI : 0.00330706
  mean: 0.828765 < 0.918696 , median: 0.83667 < 0.901204
- P_Up vs LOW_PSI : 0.0374358
  mean: 0.828765 < 0.895832 , median: 0.83667 < 0.875075
- HIGH_PSI vs LOW_PSI : 5.6964e-09
  mean: 0.918696 > 0.895832 , median: 0.901204 > 0.875075

#### 6.18 RATIO DOEXON EXON GCC

Back to: Overview | ToC

Meaning: DOEXON GCC / EXON GCC

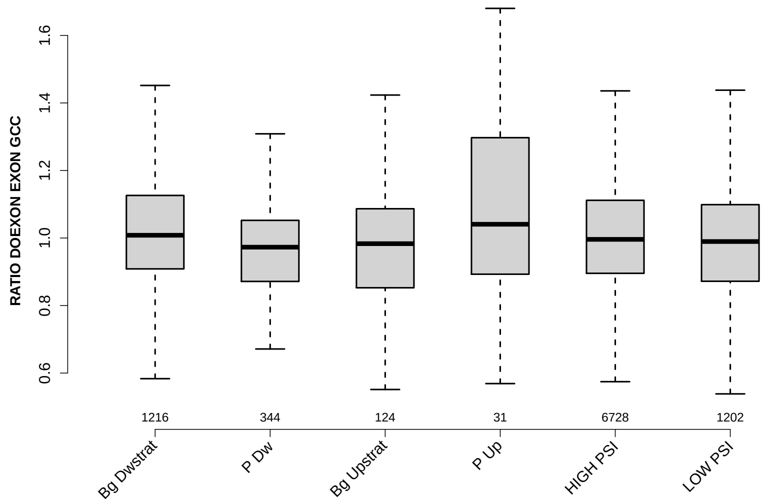

Significant results from Mann-Whitney U test:

- `Bg_Dwstrat` vs `P_Dw` : 5.12807e-06
  mean: 1.0257 > 0.975676 , median: 1.0081 > 0.972869
- `Bg_Dwstrat` vs `Bg_Upstrat` : 0.0120089
  mean: 1.0257 > 0.981398 , median: 1.0081 > 0.983207
- `Bg_Dwstrat` vs `HIGH_PSI` : 0.0215206
  mean: 1.0257 > 1.0125 , median: 1.0081 > 0.995986
- `Bg_Dwstrat` vs `LOW_PSI` : 0.000329476
  mean: 1.0257 > 1.0025 , median: 1.0081 > 0.989604
- `P_Dw` vs `HIGH_PSI` : 0.000195041
  mean: 0.975676 < 1.0125 , median: 0.972869 < 0.995986
- `HIGH_PSI` vs `LOW_PSI` : 0.0113034
  mean: 1.0125 > 1.0025 , median: 0.995986 > 0.989604

#### 6.19 SF1 HIGHESTSCORE 3SS UPINTRON

Back to: Overview | ToC

Meaning: highest score of a SF1 positon weight matrix trained with human data in the last 150 nt 3 prime intron positons of up-stream intron

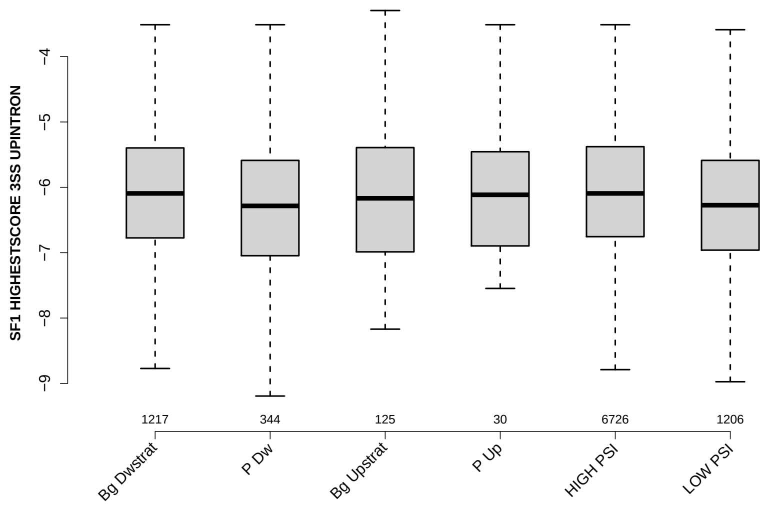

Significant results from Mann-Whitney U test:

- Bg_Dwstrat vs P_Dw : 0.00111265
  mean: -6.08482 > -6.31151 , median: -6.09302 > -6.28413
- Bg_Dwstrat vs LOW_PSI : 0.000127878
  mean: -6.08482 > -6.22325 , median: -6.09302 > -6.27371
- P_Dw vs HIGH_PSI : 7.72003e-05
  mean: -6.31151 < -6.04852 , median: -6.28413 < -6.09302
- HIGH_PSI vs LOW_PSI : 2.94071e-08
  mean: -6.04852 > -6.22325 , median: -6.09302 > -6.27371

#### 6.20 SF1 HIGHESTSCORE 3SS DOINTRON

Back to: Overview | ToC

Meaning: highest score of a SF1 positon weight matrix trained with human data in the last 150 nt 3 prime intron positons of down-stream intron

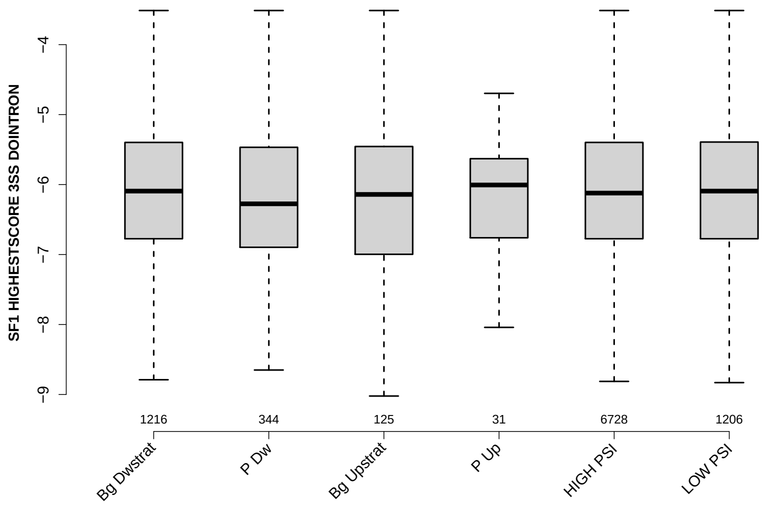

Significant results from Mann-Whitney U test:

- Bg_Dwstrat vs P_Dw : 0.0201748
  mean: -6.06529 > -6.22481 , median: -6.09302 > -6.27484
- P_Dw vs HIGH_PSI : 0.0262783
  mean: -6.22481 < -6.08306 , median: -6.27484 < -6.12133
- P_Dw vs LOW_PSI : 0.0173506
  mean: -6.22481 < -6.06687 , median: -6.27484 < -6.09302

#### 6.21 UP 5SS 20INT10EX GCC

Back to: Overview | ToC

Meaning: GC content of up-stream 5ss sequence (20int+10ex positions)

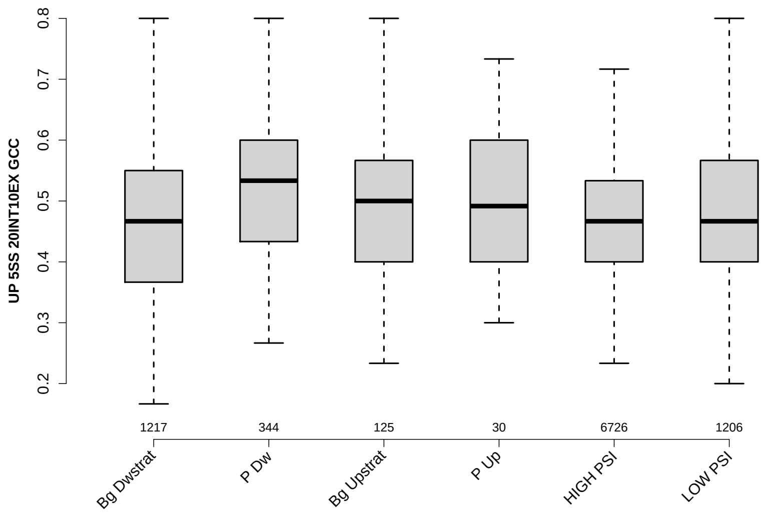

Significant results from Mann-Whitney U test:

- Bg_Dwstrat vs P_Dw : 1.00382e-13
  mean: 0.466119 < 0.518556 , median: 0.466667 < 0.533333
- Bg_Dwstrat vs LOW_PSI : 0.00653329
  mean: 0.466119 < 0.479602 , median: 0.466667 = 0.466667
- P_Dw vs Bg_Upstrat : 0.00934644
  mean: 0.518556 > 0.488 , median: 0.533333 > 0.5
- P_Dw vs HIGH_PSI : 6.97865e-15
  mean: 0.518556 > 0.468756 , median: 0.533333 > 0.466667
- P_Dw vs LOW_PSI : 1.52728e-07
  mean: 0.518556 > 0.479602 , median: 0.533333 > 0.466667
- HIGH_PSI vs LOW_PSI : 0.00612999
  mean: 0.468756 < 0.479602 , median: 0.466667 = 0.466667

#### 6.22 GCC 3SS 20INT10EX

Back to: Overview | ToC

Meaning: GC content of 3ss sequence (20int+10ex positions)

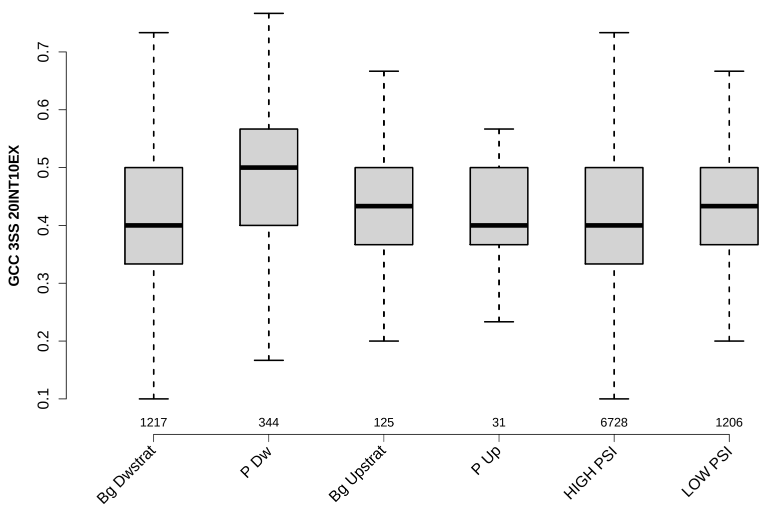

Significant results from Mann-Whitney U test:

- Bg_Dwstrat vs P_Dw : 1.23634e-20
  mean: 0.412216 < 0.478488 , median: 0.4 < 0.5
- Bg_Dwstrat vs Bg_Upstrat : 0.00604086
  mean: 0.412216 < 0.440533 , median: 0.4 < 0.433333
- Bg_Dwstrat vs LOW_PSI : 0.000107276
  mean: 0.412216 < 0.427944 , median: 0.4 < 0.433333
- P_Dw vs Bg_Upstrat : 0.00149522
  mean: 0.478488 > 0.440533 , median: 0.5 > 0.433333
- P_Dw vs P_Up : 0.000694515
  mean: 0.478488 > 0.410753 , median: 0.5 > 0.4
- P_Dw vs HIGH_PSI : 1.70115e-22
  mean: 0.478488 > 0.415111 , median: 0.5 > 0.4
- P_Dw vs LOW_PSI : 2.31887e-13
  mean: 0.478488 > 0.427944 , median: 0.5 > 0.433333
- Bg_Upstrat vs HIGH_PSI : 0.0110892
  mean: 0.440533 > 0.415111 , median: 0.433333 > 0.4
- HIGH_PSI vs LOW_PSI : 5.14955e-05
  mean: 0.415111 < 0.427944 , median: 0.4 < 0.433333

#### 6.23 GCC 5SS 20INT10EX

Back to: Overview | ToC

Meaning: GC content of 5ss sequence (20int+10ex positions)

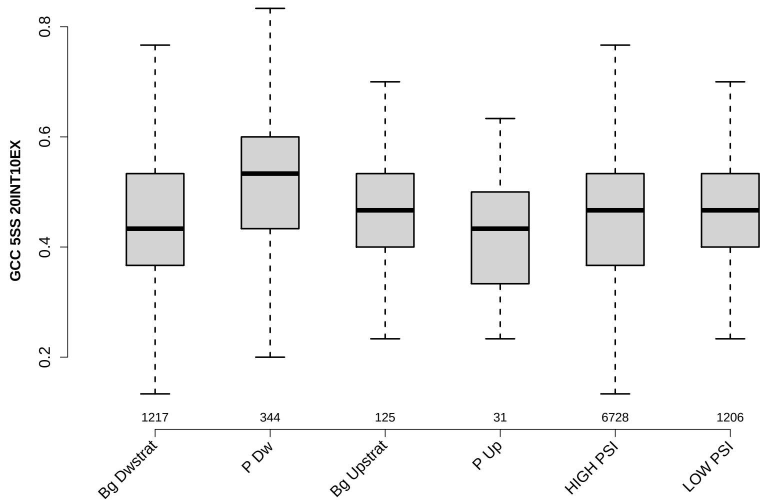

Significant results from Mann-Whitney U test:

- `Bg_Dwstrat` vs `P_Dw` : 2.94825e-15
  mean: 0.447083 < 0.503391 , median: 0.433333 < 0.533333
- `Bg_Dwstrat` vs `HIGH_PSI` : 0.0184691
  mean: 0.447083 < 0.45491 , median: 0.433333 < 0.466667
- `P_Dw` vs `Bg_Upstrat` : 0.00137177
  mean: 0.503391 > 0.466133 , median: 0.533333 > 0.466667
- `P_Dw` vs `P_Up` : 0.00154936
  mean: 0.503391 > 0.430108 , median: 0.533333 > 0.433333
- `P_Dw` vs `HIGH_PSI` : 1.68448e-14
  mean: 0.503391 > 0.45491 , median: 0.533333 > 0.466667
- `P_Dw` vs `LOW_PSI` : 1.23533e-15
  mean: 0.503391 > 0.450415 , median: 0.533333 > 0.466667

#### 6.24 DO 3SS 20INT10EX GCC

Back to: Overview | ToC

Meaning: GC content of down-stream 3ss sequence (20int+10ex positions)

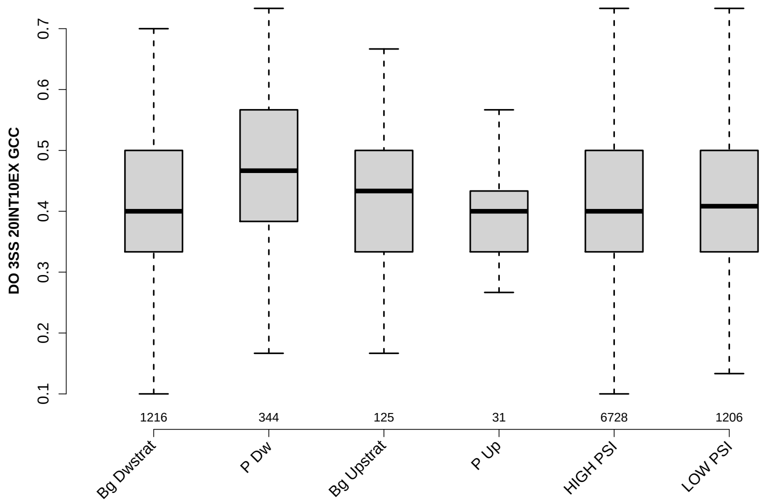

Significant results from Mann-Whitney U test:

- `Bg_Dwstrat` vs `P_Dw` : 1.82169e-11
  mean: 0.414035 < 0.461483 , median: 0.4 < 0.466667
- `P_Dw` vs `Bg_Upstrat` : 2.70478e-05
  mean: 0.461483 > 0.4116 , median: 0.466667 > 0.433333
- `P_Dw` vs `P_Up` : 0.00687894
  mean: 0.461483 > 0.406452 , median: 0.466667 > 0.4
- `P_Dw` vs `HIGH_PSI` : 1.06993e-12
  mean: 0.461483 > 0.416971 , median: 0.466667 > 0.4
- `P_Dw` vs `LOW_PSI` : 8.24464e-11
  mean: 0.461483 > 0.41686 , median: 0.466667 > 0.408333

#### 6.25 MAXENTSCR HSAMODEL UPSTRM 5SS

Back to: Overview | ToC

Meaning: maximum entropy score of 5ss of up-stream exon using a model trained with human splice sites

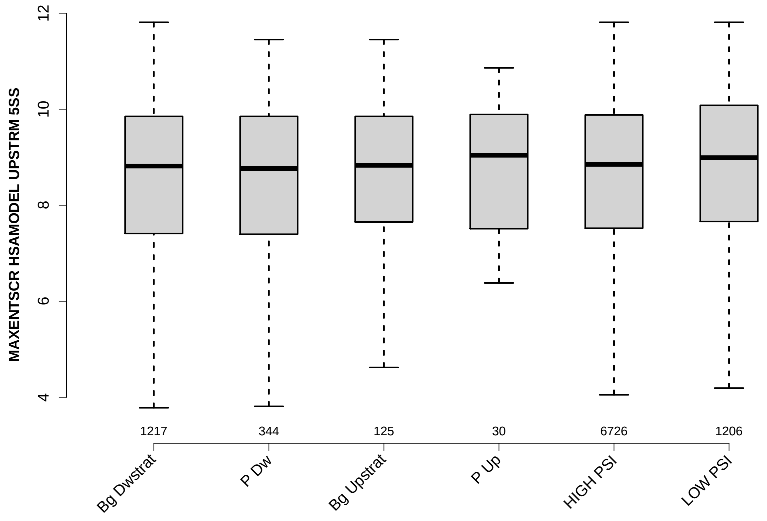

Significant results from Mann-Whitney U test:

- `Bg_Dwstrat` vs `LOW_PSI` : 0.00745361
  mean: 7.9084 < 8.5773 , median: 8.815 < 8.99
- `P_Dw` vs `LOW_PSI` : 0.0450125
  mean: 8.2209 < 8.5773 , median: 8.765 < 8.99
- `HIGH_PSI` vs `LOW_PSI` : 0.015593
  mean: 8.4394 < 8.5773 , median: 8.85 < 8.99

#### 6.26 MAXENTSCR HSAMODEL 3SS

Back to: Overview | ToC

Meaning: maximum entropy score of 3ss using a model trained with human splice sites

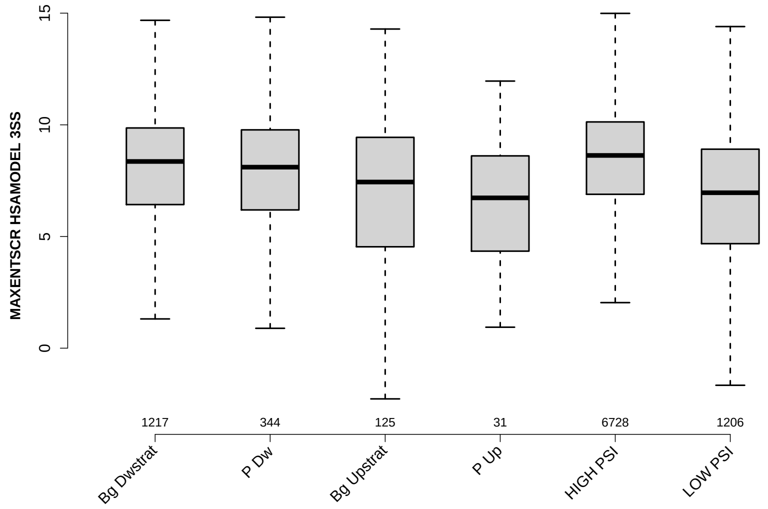

Significant results from Mann-Whitney U test:

- `Bg_Dwstrat` vs `Bg_Upstrat` : 0.00498044
  mean: 7.6581 > 6.5258 , median: 8.36 > 7.44
- `Bg_Dwstrat` vs `P_Up` : 0.0044846
  mean: 7.6581 > 6.589 , median: 8.36 > 6.73
- `Bg_Dwstrat` vs `HIGH_PSI` : 5.31137e-05
  mean: 7.6581 < 8.3973 , median: 8.36 < 8.63
- `Bg_Dwstrat` vs `LOW_PSI` : 7.21984e-27
  mean: 7.6581 > 6.3804 , median: 8.36 > 6.96
- `P_Dw` vs `Bg_Upstrat` : 0.0326791
  mean: 7.8561 > 6.5258 , median: 8.105 > 7.44
- `P_Dw` vs `P_Up` : 0.0109617
  mean: 7.8561 > 6.589 , median: 8.105 > 6.73
- `P_Dw` vs `HIGH_PSI` : 0.000898408
  mean: 7.8561 < 8.3973 , median: 8.105 < 8.63
- `P_Dw` vs `LOW_PSI` : 8.1728e-11
  mean: 7.8561 > 6.3804 , median: 8.105 > 6.96
- `Bg_Upstrat` vs `HIGH_PSI` : 1.8392e-05
  mean: 6.5258 < 8.3973 , median: 7.44 < 8.63
- `P_Up` vs `HIGH_PSI` : 0.000300135
  mean: 6.589 < 8.3973 , median: 6.73 < 8.63

- `HIGH_PSI` vs `LOW_PSI` : 1.89162e-73
  mean: 8.3973 > 6.3804 , median: 8.63 > 6.96

#### 6.27 MAXENTSCR HSAMODEL 5SS

Back to: Overview | ToC

Meaning: maximum entropy score of 5ss using a model trained with human splice sites

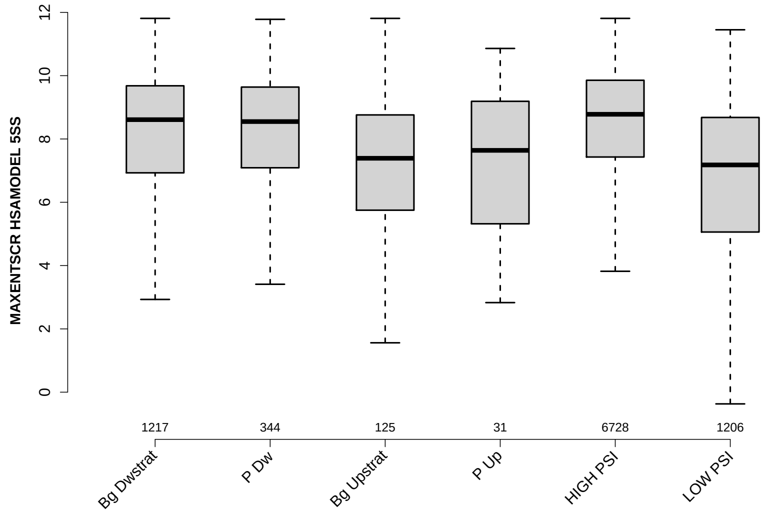

Significant results from Mann-Whitney U test:

- Bg_Dwstrat vs Bg_Upstrat : 1.99477e-07
  mean: 7.6254 > 6.3274 , median: 8.61 > 7.39
- Bg_Dwstrat vs P_Up : 0.0325407
  mean: 7.6254 > 7.1535 , median: 8.61 > 7.64
- Bg_Dwstrat vs HIGH_PSI : 4.77579e-06
  mean: 7.6254 < 8.3829 , median: 8.61 < 8.78
- Bg_Dwstrat vs LOW_PSI : 5.76687e-39
  mean: 7.6254 > 6.4028 , median: 8.61 > 7.18
- P_Dw vs Bg_Upstrat : 3.33725e-06
  mean: 7.9974 > 6.3274 , median: 8.55 > 7.39
- P_Dw vs P_Up : 0.043406
  mean: 7.9974 > 7.1535 , median: 8.55 > 7.64
- P_Dw vs HIGH_PSI : 0.00180104
  mean: 7.9974 < 8.3829 , median: 8.55 < 8.78
- P_Dw vs LOW_PSI : 3.29089e-18
  mean: 7.9974 > 6.4028 , median: 8.55 > 7.18
- Bg_Upstrat vs HIGH_PSI : 1.09006e-12
  mean: 6.3274 < 8.3829 , median: 7.39 < 8.78
- P_Up vs HIGH_PSI : 0.00268599
  mean: 7.1535 < 8.3829 , median: 7.64 < 8.78

- `HIGH_PSI` vs `LOW_PSI` : 1.63811e-104
  mean: 8.3829 > 6.4028 , median: 8.78 > 7.18

#### 6.28 MAXENTSCR HSAMODEL DOWNSTRM 3SS

Back to: Overview | ToC

Meaning: maximum entropy score of 3ss of down-stream exon using a model trained with human splice sites

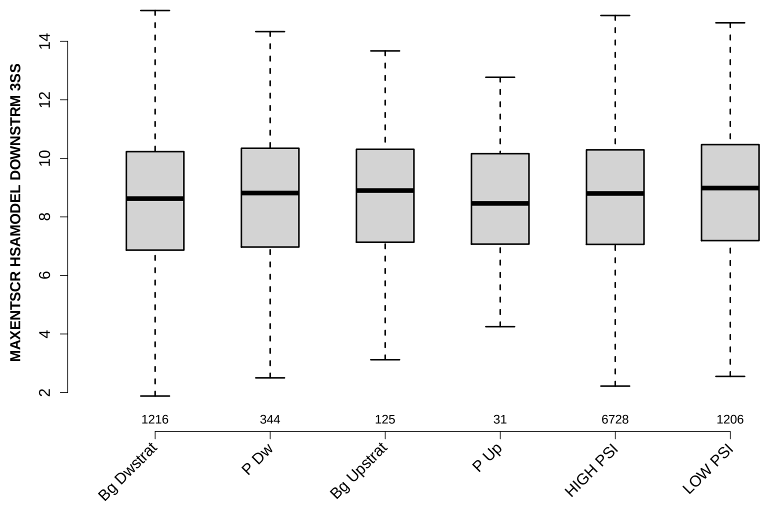

Significant results from Mann-Whitney U test:

- `Bg_Dwstrat` vs `LOW_PSI` : 0.00290708
  mean: 8.0458 < 8.658 , median: 8.625 < 8.985
- `HIGH_PSI` vs `LOW_PSI` : 0.0336775
  mean: 8.511 < 8.658 , median: 8.8 < 8.985

#### 6.29 DIST FROM MAXBP TO 3SS UPINTRON

Back to: Overview | ToC

Meaning: distance to 3ss of best precited BP

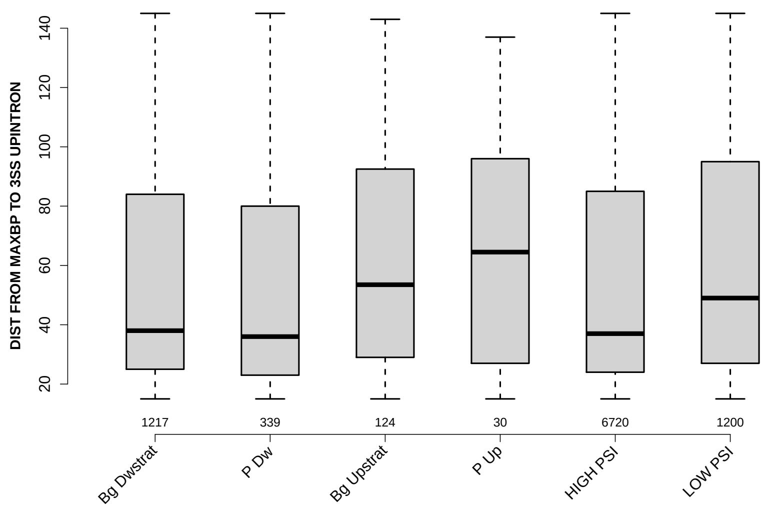

Significant results from Mann-Whitney U test:

- Bg_Dwstrat vs Bg_Upstrat : 0.00685065
  mean: 56.2662 < 64.5161 , median: 38 < 53.5
- Bg_Dwstrat vs LOW_PSI : 0.000131458
  mean: 56.2662 < 62.1158 , median: 38 < 49
- P_Dw vs Bg_Upstrat : 0.00274466
  mean: 54.8319 < 64.5161 , median: 36 < 53.5
- P_Dw vs LOW_PSI : 0.000479244
  mean: 54.8319 < 62.1158 , median: 36 < 49
- Bg_Upstrat vs HIGH_PSI : 0.00163335
  mean: 64.5161 > 55.5583 , median: 53.5 > 37
- HIGH_PSI vs LOW_PSI : 1.9157e-09
  mean: 55.5583 < 62.1158 , median: 37 < 49

#### 6.30 SCORE FOR MAXBP SEQ UPINTRON

Back to: Overview | ToC

Meaning: BP sequence score of best predicted BP

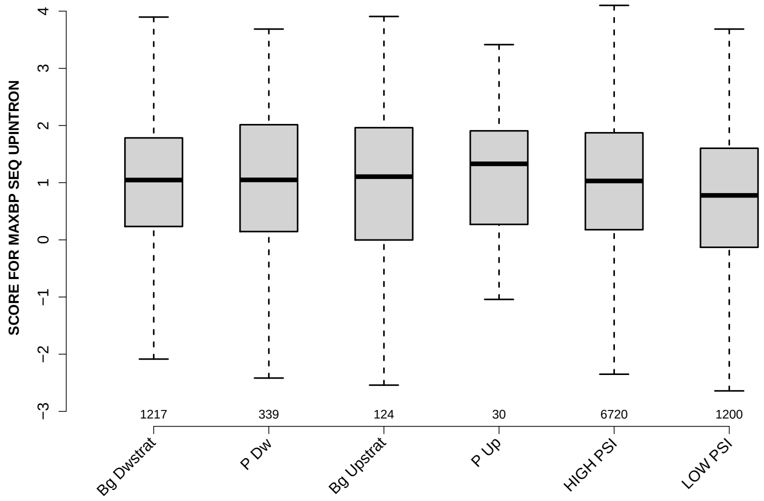

Significant results from Mann-Whitney U test:

- `Bg_Dwstrat` vs `LOW_PSI` : 8.73266e-07
  mean: 1.0007 > 0.747279 , median: 1.0462 > 0.777644
- `P_Dw` vs `LOW_PSI` : 0.00177745
  mean: 0.994349 > 0.747279 , median: 1.0489 > 0.777644
- `HIGH_PSI` vs `LOW_PSI` : 2.09972e-10
  mean: 1.0065 > 0.747279 , median: 1.0303 > 0.777644

#### 6.31 PYRIMIDINECONT MAXBP UPINTRON

Back to: Overview | ToC

Meaning: Pyrimidine content between the BP adenine and the 3 prime splice site for best BP

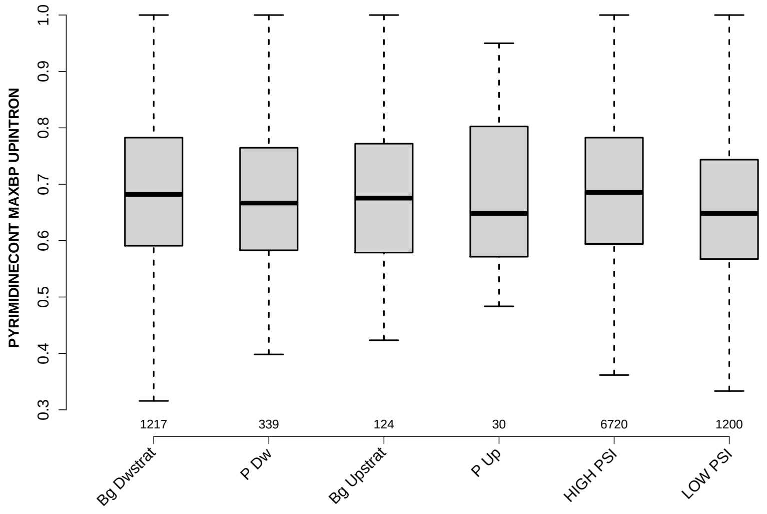

Significant results from Mann-Whitney U test:

- `Bg_Dwstrat` vs `LOW_PSI` : 7.56535e-08
  mean: 0.685626 > 0.658969 , median: 0.681818 > 0.648162
- `P_Dw` vs `HIGH_PSI` : 0.0352363
  mean: 0.675244 < 0.691152 , median: 0.666667 < 0.685335
- `P_Dw` vs `LOW_PSI` : 0.0448129
  mean: 0.675244 > 0.658969 , median: 0.666667 > 0.648162
- `HIGH_PSI` vs `LOW_PSI` : 7.95324e-16
  mean: 0.691152 > 0.658969 , median: 0.685335 > 0.648162

#### 6.32 POLYPYRITRAC OFFSET MAXBP UPINTRON

Back to: Overview | ToC

Meaning: Polypyrimidine track offset relative to the BP adenine for best BP

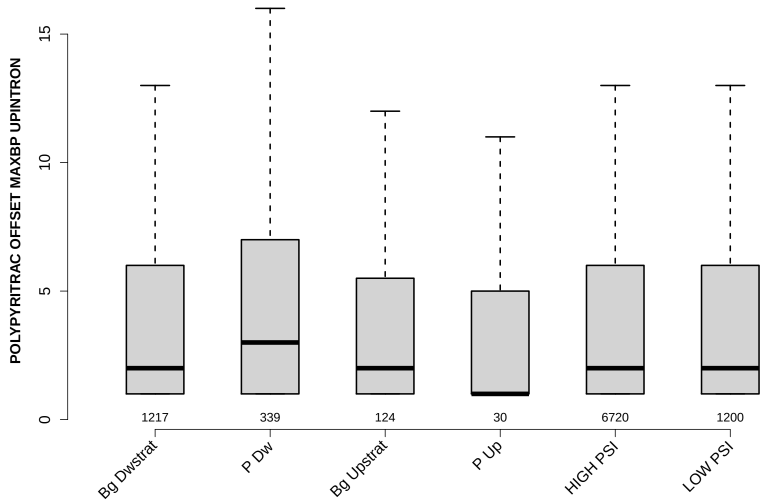

Significant results from Mann-Whitney U test:

- Bg_Dwstrat vs P_Dw : 0.000208113
  mean: 4.0164 < 5.4277 , median: 2 < 3
- Bg_Dwstrat vs LOW_PSI : 0.037406
  mean: 4.0164 < 4.9383 , median: 2 = 2
- P_Dw vs HIGH_PSI : 6.23888e-05
  mean: 5.4277 > 4.0061 , median: 3 > 2
- P_Dw vs LOW_PSI : 0.024821
  mean: 5.4277 > 4.9383 , median: 3 > 2
- HIGH_PSI vs LOW_PSI : 0.0125436
  mean: 4.0061 < 4.9383 , median: 2 = 2

#### 6.33 POLYPYRITRAC LEN MAXBP UPINTRON

Back to: Overview | ToC

Meaning: Polypyrimidine track length for best BP

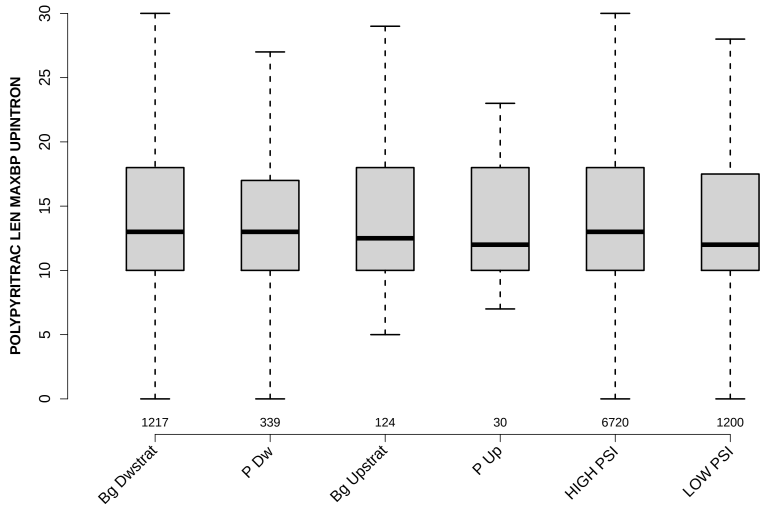

Significant results from Mann-Whitney U test:

- `Bg_Dwstrat` vs `LOW_PSI` : 0.0151575
  mean: 14.7017 > 14.6567 , median: 13 > 12
- `HIGH_PSI` vs `LOW_PSI` : 0.00434324
  mean: 14.7149 > 14.6567 , median: 13 > 12

#### 6.34 POLYPYRITRAC SCORE MAXBP UPINTRON

Back to: Overview | ToC

Meaning: Polypyrimidine track score for best BP

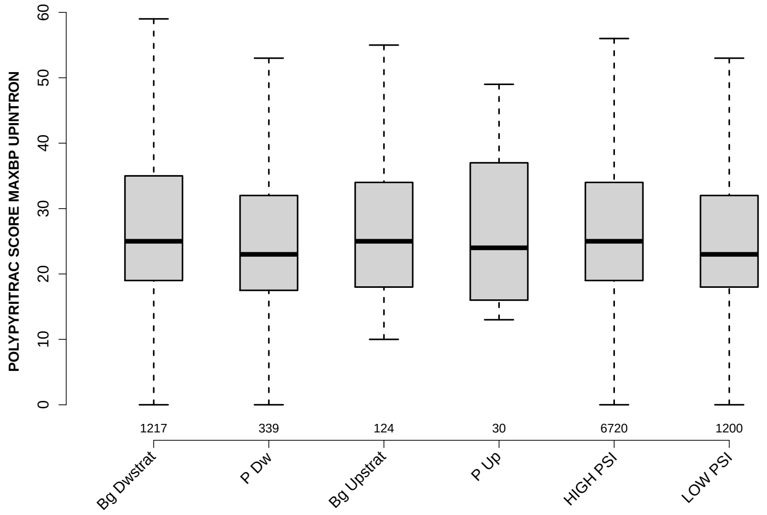

Significant results from Mann-Whitney U test:

- Bg_Dwstrat vs P_Dw : 0.0117101
  mean: 27.9844 > 25.9499 , median: 25 > 23
- Bg_Dwstrat vs LOW_PSI : 0.00231902
  mean: 27.9844 > 27.7225 , median: 25 > 23
- P_Dw vs HIGH_PSI : 0.00963916
  mean: 25.9499 < 28.078 , median: 23 < 25
- HIGH_PSI vs LOW_PSI : 0.000266642
  mean: 28.078 > 27.7225 , median: 25 > 23

#### 6.35 BPSCORE MAXBP UPINTRON

Back to: Overview | ToC

Meaning: SVM classification score of best BP

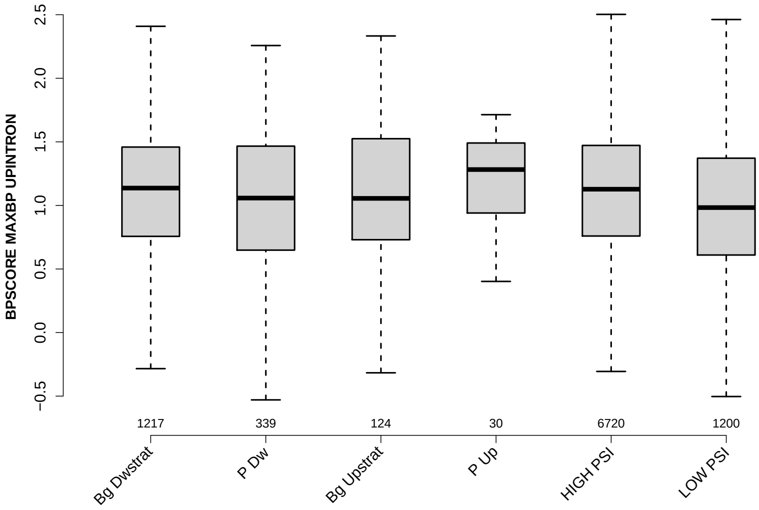

Significant results from Mann-Whitney U test:

- `Bg_Dwstrat` vs `LOW_PSI` : 1.15332e-09
  mean: 1.0892 > 0.914979 , median: 1.1364 > 0.982977
- `P_Dw` vs `HIGH_PSI` : 0.0324404
  mean: 0.975689 < 1.0942 , median: 1.0579 < 1.1275
- `Bg_Upstrat` vs `LOW_PSI` : 0.0202327
  mean: 1.0831 > 0.914979 , median: 1.0554 > 0.982977
- `P_Up` vs `LOW_PSI` : 0.015323
  mean: 1.1844 > 0.914979 , median: 1.2821 > 0.982977
- `HIGH_PSI` vs `LOW_PSI` : 3.4211e-15
  mean: 1.0942 > 0.914979 , median: 1.1275 > 0.982977

#### 6.36 NUM PREDICTED BPS UPINTRON

Back to: Overview | ToC

Meaning: number of all predicted BPs which have a positive BP score

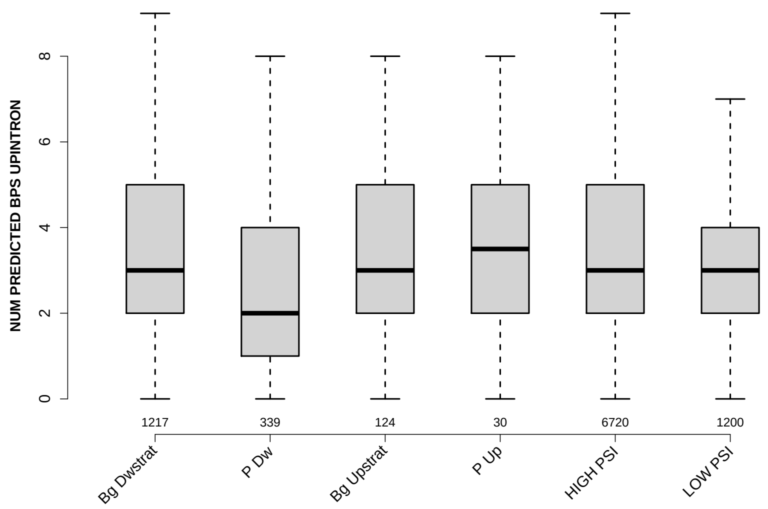

Significant results from Mann-Whitney U test:

- Bg_Dwstrat vs P_Dw : 5.54162e-15
  mean: 3.493 > 2.59 , median: 3 > 2
- Bg_Dwstrat vs LOW_PSI : 1.86835e-05
  mean: 3.493 > 3.1658 , median: 3 = 3
- P_Dw vs Bg_Upstrat : 4.22426e-05
  mean: 2.59 < 3.4113 , median: 2 < 3
- P_Dw vs P_Up : 0.0127691
  mean: 2.59 < 3.4333 , median: 2 < 3.5
- P_Dw vs HIGH_PSI : 1.01262e-16
  mean: 2.59 < 3.4536 , median: 2 < 3
- P_Dw vs LOW_PSI : 4.9877e-06
  mean: 2.59 < 3.1658 , median: 2 < 3
- HIGH_PSI vs LOW_PSI : 4.08254e-07
  mean: 3.4536 > 3.1658 , median: 3 = 3

#### 6.37 MEDIAN DIST FROM BP TO 3SS UPINTRON

Back to: Overview | ToC

Meaning: like DIST FROM MAXBP TO 3SS but median over top-3 predicted BPs

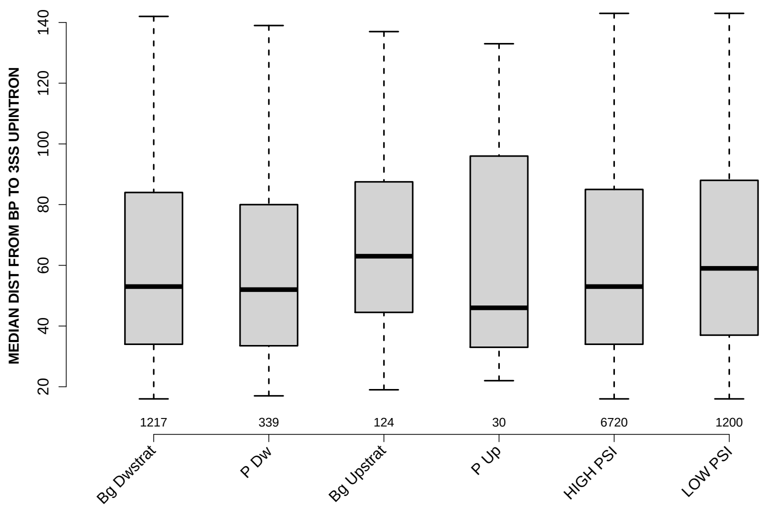

Significant results from Mann-Whitney U test:

- `Bg_Dwstrat` vs `Bg_Upstrat` : 0.0114078
  mean: 61.304 < 66.7056 , median: 53 < 63
- `Bg_Dwstrat` vs `LOW_PSI` : 0.0101538
  mean: 61.304 < 64.1996 , median: 53 < 59
- `P_Dw` vs `Bg_Upstrat` : 0.0113484
  mean: 60.0885 < 66.7056 , median: 52 < 63
- `P_Dw` vs `LOW_PSI` : 0.0365315
  mean: 60.0885 < 64.1996 , median: 52 < 59
- `Bg_Upstrat` vs `HIGH_PSI` : 0.00767258
  mean: 66.7056 > 60.9685 , median: 63 > 53
- `HIGH_PSI` vs `LOW_PSI` : 0.00047277
  mean: 60.9685 < 64.1996 , median: 53 < 59

#### 6.38 MEDIAN SCORE FOR BPSEQ UPINTRON

Back to: Overview | ToC

Meaning: like SCORE FOR MAXBP SEQ but median over top-3 predicted BPs

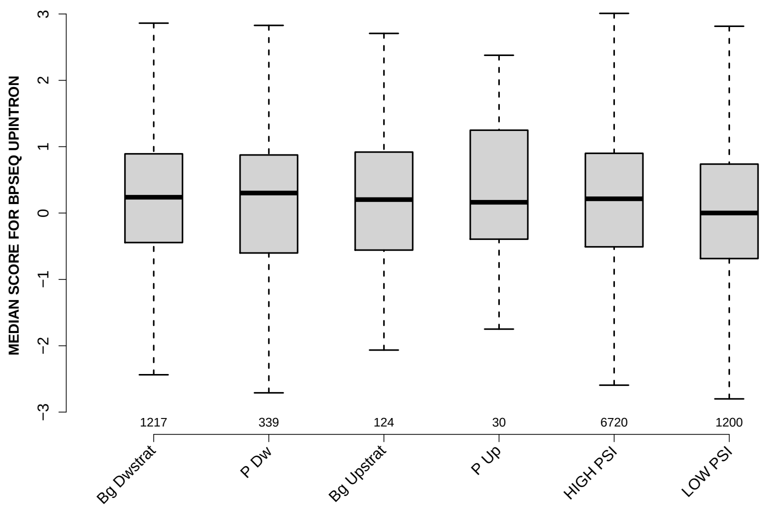

Significant results from Mann-Whitney U test:

- `Bg_Dwstrat` vs `LOW_PSI` : 1.85863e-06
  mean: 0.217693 > 0.011832 , median: 0.238131 > 0.000812688
- `P_Dw` vs `LOW_PSI` : 0.00508848
  mean: 0.183173 > 0.011832 , median: 0.30173 > 0.000812688
- `HIGH_PSI` vs `LOW_PSI` : 1.58089e-08
  mean: 0.201832 > 0.011832 , median: 0.214651 > 0.000812688

#### 6.39 MEDIAN PYRIMIDINECONT UPINTRON

Back to: Overview | ToC

Meaning: like PYRIMIDINECONT MAXBP but median over top-3 predicted BPs

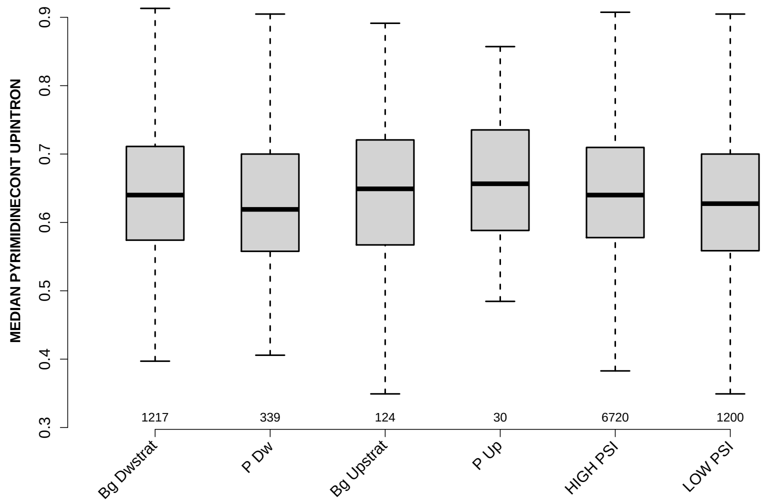

Significant results from Mann-Whitney U test:

- Bg_Dwstrat vs P_Dw : 0.00808645
  mean: 0.643386 > 0.628771 , median: 0.64 > 0.619048
- Bg_Dwstrat vs LOW_PSI : 0.000951656
  mean: 0.643386 > 0.630386 , median: 0.64 > 0.627451
- P_Dw vs HIGH_PSI : 0.000992192
  mean: 0.628771 < 0.64712 , median: 0.619048 < 0.64
- HIGH_PSI vs LOW_PSI : 7.26269e-07
  mean: 0.64712 > 0.630386 , median: 0.64 > 0.627451

#### 6.40 MEDIAN POLYPYRITRAC OFFSET UPINTRON

Back to: Overview | ToC

Meaning: like POLYPYRITRAC OFFSET MAXBP but median over top-3 predicted BPs

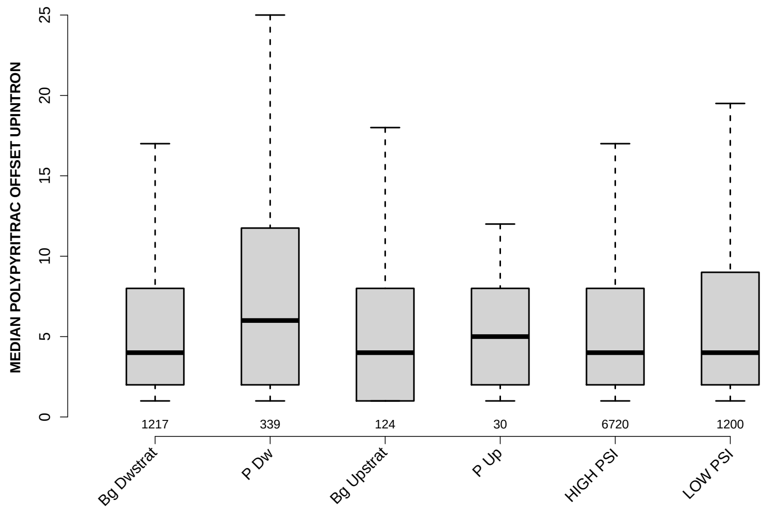

Significant results from Mann-Whitney U test:

- `Bg_Dwstrat` vs `P_Dw` : 2.94382e-09
  mean: 6.0049 < 9.4307 , median: 4 < 6
- `Bg_Dwstrat` vs `LOW_PSI` : 0.00130046
  mean: 6.0049 < 7.3642 , median: 4 = 4
- `P_Dw` vs `Bg_Upstrat` : 0.00142133
  mean: 9.4307 > 6.8629 , median: 6 > 4
- `P_Dw` vs `HIGH_PSI` : 8.5879e-11
  mean: 9.4307 > 6.0569 , median: 6 > 4
- `P_Dw` vs `LOW_PSI` : 0.000171125
  mean: 9.4307 > 7.3642 , median: 6 > 4
- `HIGH_PSI` vs `LOW_PSI` : 6.1334e-05
  mean: 6.0569 < 7.3642 , median: 4 = 4

#### 6.41 MEDIAN POLYPYRITRAC LEN UPINTRON

Back to: Overview | ToC

Meaning: like POLYPYRITRAC LEN MAXBP but median over top-3 predicted BPs

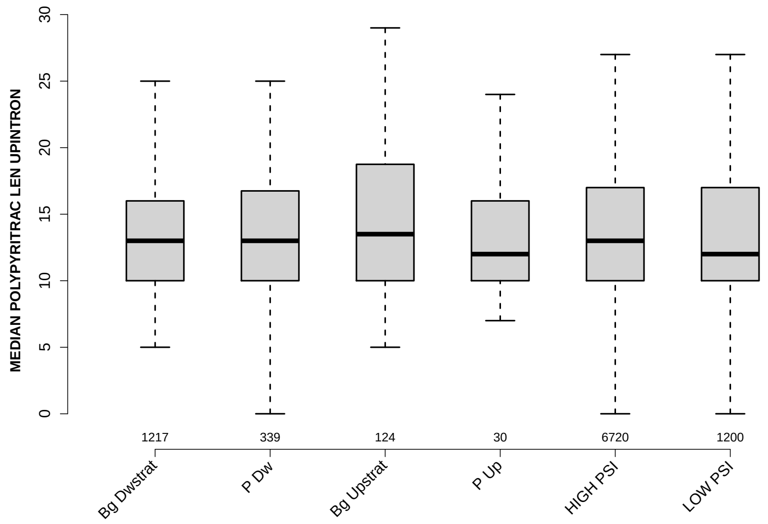

Significant results from Mann-Whitney U test:

- none

#### 6.42 MEDIAN POLYPYRITRAC SCORE UPINTRON

Back to: Overview | ToC

Meaning: like POLYPYRITRAC SCORE MAXBP but median over top-3 predicted BPs

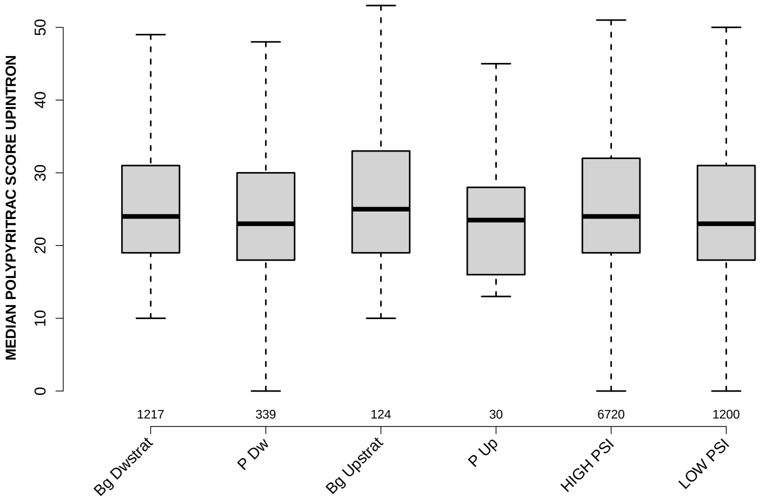

Significant results from Mann-Whitney U test:

- `P_Dw` vs `Bg_Upstrat` : 0.0244785
  mean: 25.1327 < 31.4919 , median: 23 < 25
- `P_Dw` vs `HIGH_PSI` : 0.028502
  mean: 25.1327 < 27.0268 , median: 23 < 24
- `HIGH_PSI` vs `LOW_PSI` : 0.0412494
  mean: 27.0268 < 27.2179 , median: 24 > 23

#### 6.43 MEDIAN BPSCORE UPINTRON

Back to: Overview | ToC

Meaning: like BPSCORE MAXBP but median over top-3 predicted BPs

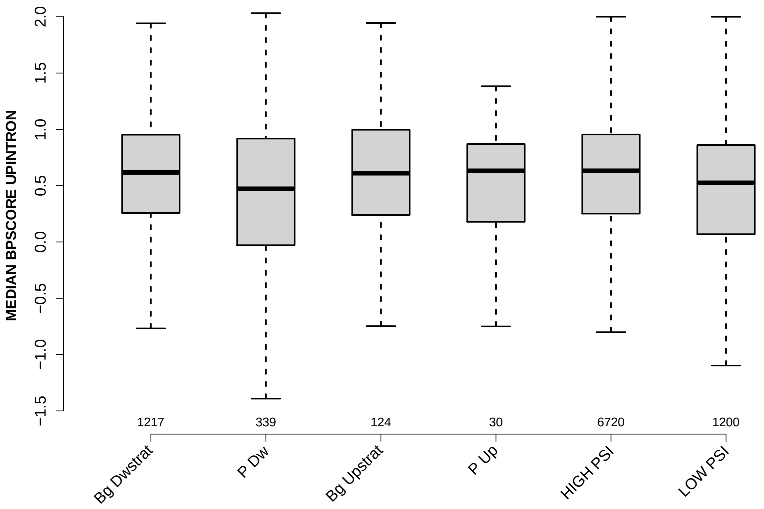

Significant results from Mann-Whitney U test:

- `Bg_Dwstrat` vs `P_Dw` : 3.46684e-05
  mean: 0.523907 > 0.261526 , median: 0.617673 > 0.472187
- `Bg_Dwstrat` vs `LOW_PSI` : 4.81716e-07
  mean: 0.523907 > 0.3577 , median: 0.617673 > 0.525109
- `P_Dw` vs `Bg_Upstrat` : 0.0119535
  mean: 0.261526 < 0.470837 , median: 0.472187 < 0.61077
- `P_Dw` vs `HIGH_PSI` : 3.45178e-06
  mean: 0.261526 < 0.527849 , median: 0.472187 < 0.632114
- `Bg_Upstrat` vs `LOW_PSI` : 0.0292465
  mean: 0.470837 > 0.3577 , median: 0.61077 > 0.525109
- `HIGH_PSI` vs `LOW_PSI` : 1.87209e-11
  mean: 0.527849 > 0.3577 , median: 0.632114 > 0.525109

#### 6.44 DIST FROM MAXBP TO 3SS DOINTRON

Back to: Overview | ToC

Meaning: distance to 3ss of best precited BP

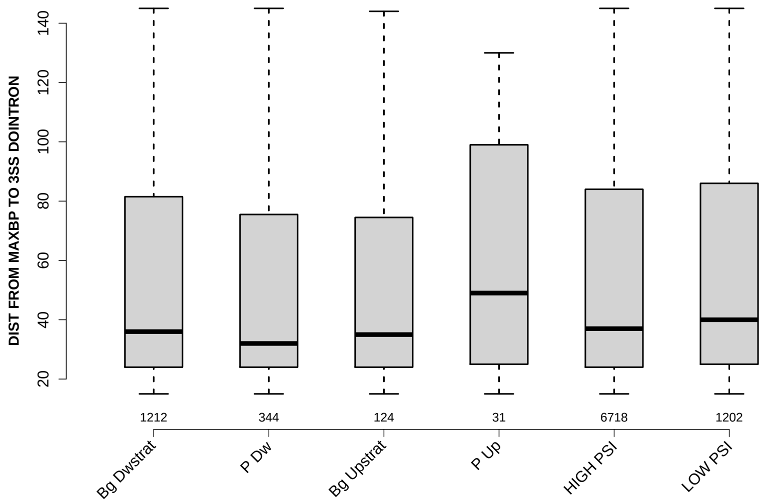

Significant results from Mann-Whitney U test:

- `P_Dw` vs `LOW_PSI` : 0.0338565
  mean: 51.9012 $<$ 56.9842 , median: 32 $<$ 40

#### 6.45 SCORE FOR MAXBP SEQ DOINTRON

Back to: Overview | ToC

Meaning: BP sequence score of best predicted BP

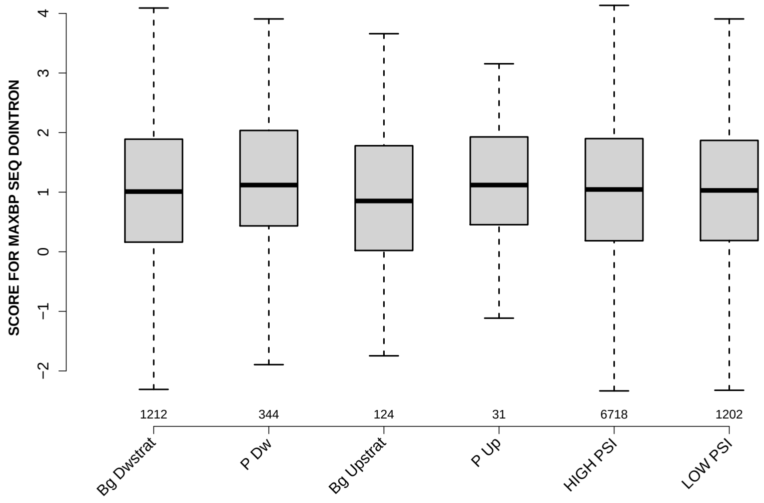

Significant results from Mann-Whitney U test:

- none

#### 6.46 PYRIMIDINECONT MAXBP DOINTRON

Back to: Overview | ToC

Meaning: Pyrimidine content between the BP adenine and the 3 prime splice site for best BP

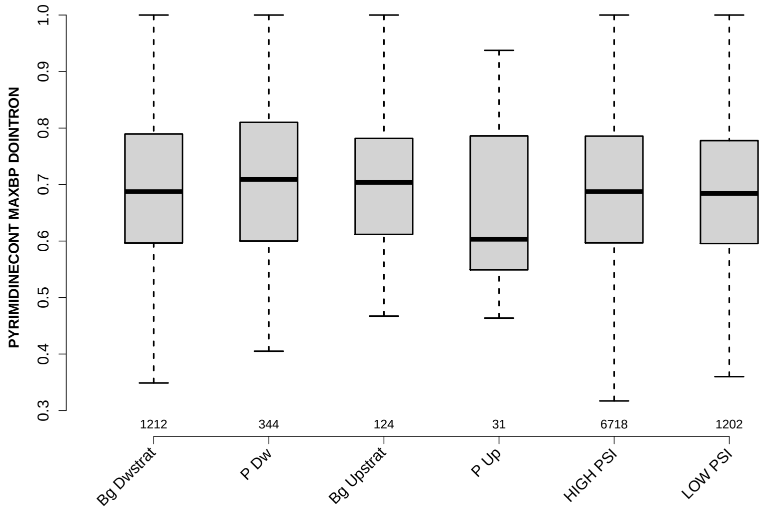

Significant results from Mann-Whitney U test:

- `P_Dw` vs `LOW_PSI` : 0.0285899
  mean: 0.705353 > 0.689647 , median: 0.709005 > 0.684211

#### 6.47 POLYPYRITRAC OFFSET MAXBP DOINTRON

Back to: Overview | ToC

Meaning: Polypyrimidine track offset relative to the BP adenine for best BP

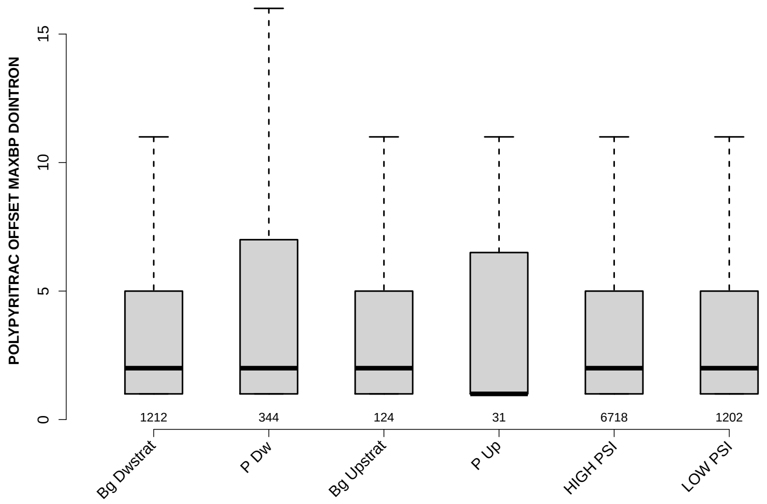

Significant results from Mann-Whitney U test:

- `Bg_Dwstrat` vs `P_Dw` : 0.00212106
  mean: 3.6691 < 4.6657 , median: 2 = 2
- `P_Dw` vs `HIGH_PSI` : 0.0069995
  mean: 4.6657 > 3.8876 , median: 2 = 2
- `P_Dw` vs `LOW_PSI` : 0.0146626
  mean: 4.6657 > 3.728 , median: 2 = 2

#### 6.48 POLYPYRITRAC LEN MAXBP DOINTRON

Back to: Overview | ToC

Meaning: Polypyrimidine track length for best BP

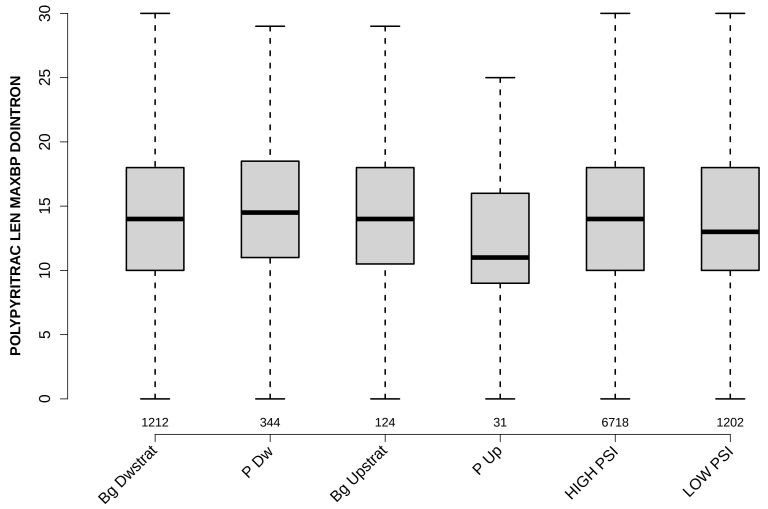

Significant results from Mann-Whitney U test:

- `P_Dw` vs `P_Up` : 0.0173514
  mean: 15.532 > 12.7419 , median: 14.5 > 11
- `P_Dw` vs `HIGH_PSI` : 0.0447688
  mean: 15.532 > 15.0168 , median: 14.5 > 14

#### 6.49 POLYPYRITRAC SCORE MAXBP DOINTRON

Back to: Overview | ToC

Meaning: Polypyrimidine track score for best BP

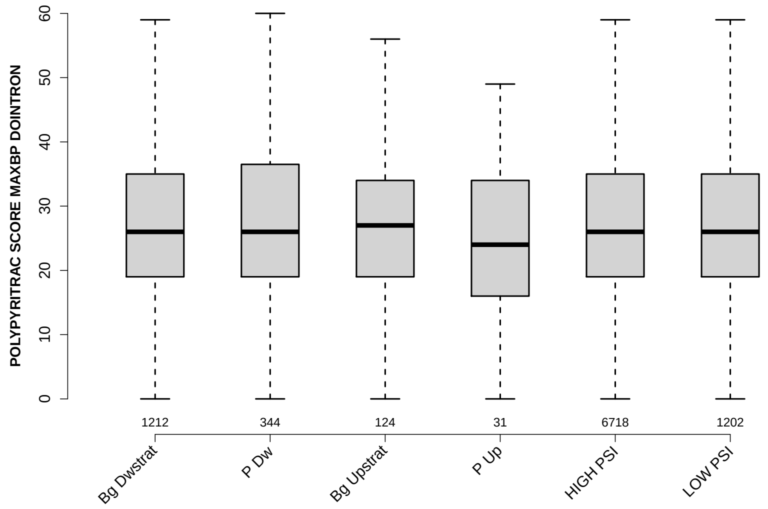

Significant results from Mann-Whitney U test:

- none

#### 6.50 BPSCORE MAXBP DOINTRON

Back to: Overview | ToC

Meaning: SVM classification score of best BP

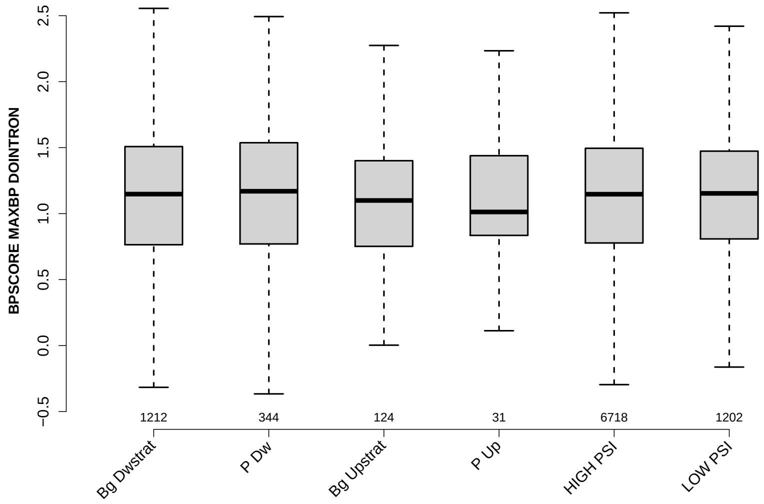

Significant results from Mann-Whitney U test:

- none

#### 6.51 NUM PREDICTED BPS DOINTRON

Back to: Overview | ToC

Meaning: number of all predicted BPs which have a positive BP score

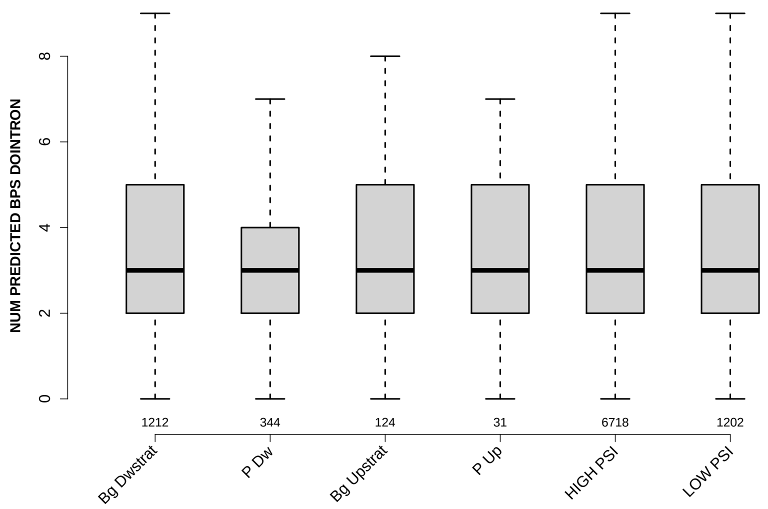

Significant results from Mann-Whitney U test:

- Bg_Dwstrat vs P_Dw : 2.8745e-09
  mean: 3.5578 > 2.8081 , median: 3 = 3
- P_Dw vs Bg_Upstrat : 0.0154964
  mean: 2.8081 < 3.3468 , median: 3 = 3
- P_Dw vs HIGH_PSI : 1.60903e-09
  mean: 2.8081 < 3.4802 , median: 3 = 3
- P_Dw vs LOW_PSI : 2.41625e-09
  mean: 2.8081 < 3.5657 , median: 3 = 3

#### 6.52 MEDIAN DIST FROM BP TO 3SS DOINTRON

Back to: Overview | ToC

Meaning: like DIST FROM MAXBP TO 3SS but median over top-3 predicted BPs

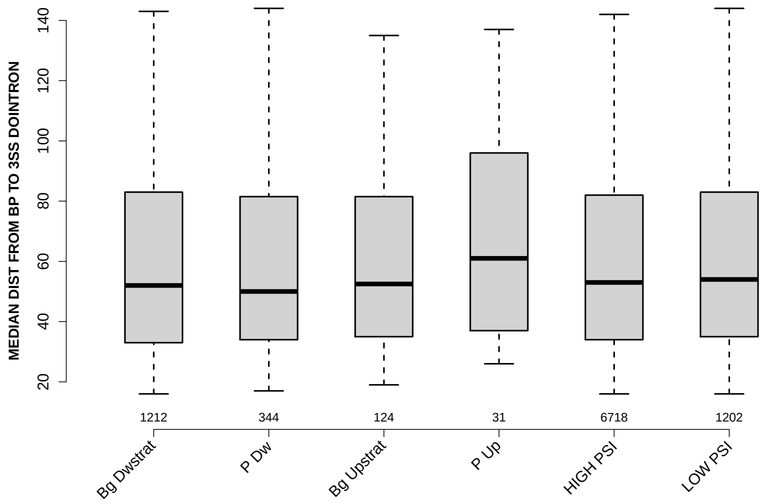

Significant results from Mann-Whitney U test:

- none

#### 6.53 MEDIAN SCORE FOR BPSEQ DOINTRON

Back to: Overview | ToC

Meaning: like SCORE FOR MAXBP SEQ but median over top-3 predicted BPs

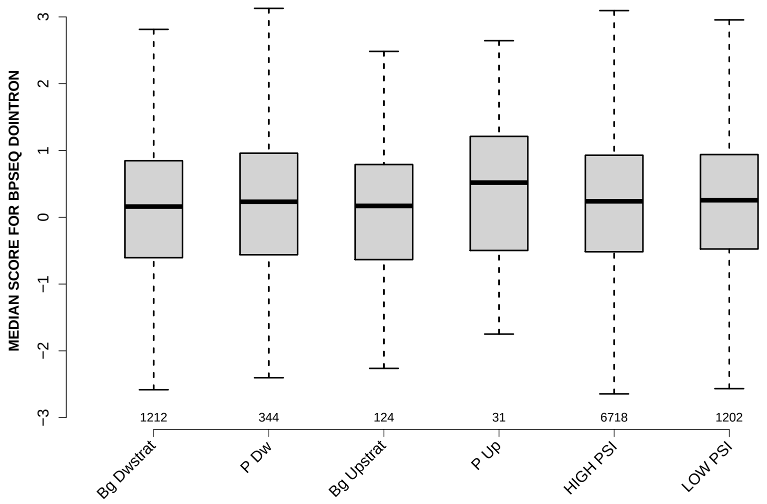

Significant results from Mann-Whitney U test:

- `Bg_Dwstrat` vs `LOW_PSI` : 0.0353051
  mean: 0.149369 < 0.227168 , median: 0.160506 < 0.2552

#### 6.54 MEDIAN PYRIMIDINECONT DOINTRON

Back to: Overview | ToC

Meaning: like PYRIMIDINECONT MAXBP but median over top-3 predicted BPs

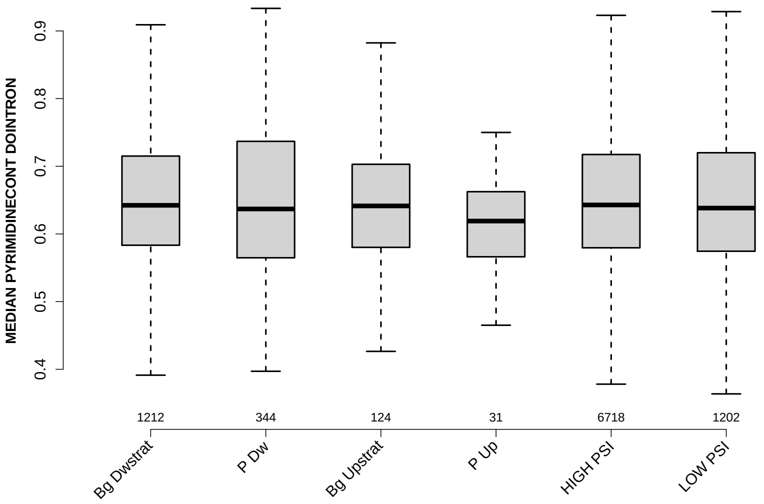

Significant results from Mann-Whitney U test:

- none

#### 6.55 MEDIAN POLYPYRITRAC OFFSET DOINTRON

Back to: Overview | ToC

Meaning: like POLYPYRITRAC OFFSET MAXBP but median over top-3 predicted BPs

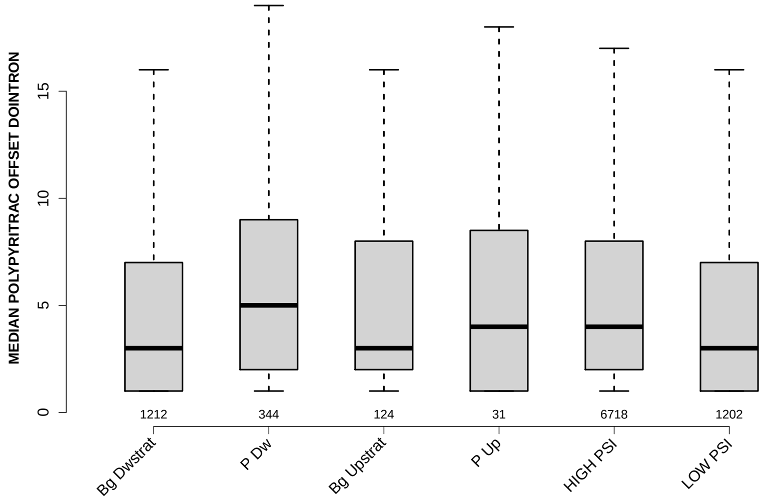

Significant results from Mann-Whitney U test:

- `Bg_Dwstrat` vs `P_Dw` : 1.06282e-06
  mean: 5.5173 < 7.7878 , median: 3 < 5
- `Bg_Dwstrat` vs `HIGH_PSI` : 0.00231446
  mean: 5.5173 < 6.0331 , median: 3 < 4
- `P_Dw` vs `HIGH_PSI` : 0.000208966
  mean: 7.7878 > 6.0331 , median: 5 > 4
- `P_Dw` vs `LOW_PSI` : 0.00015016
  mean: 7.7878 > 5.8087 , median: 5 > 3

#### 6.56 MEDIAN POLYPYRITRAC LEN DOINTRON

Back to: Overview | ToC

Meaning: like POLYPYRITRAC LEN MAXBP but median over top-3 predicted BPs

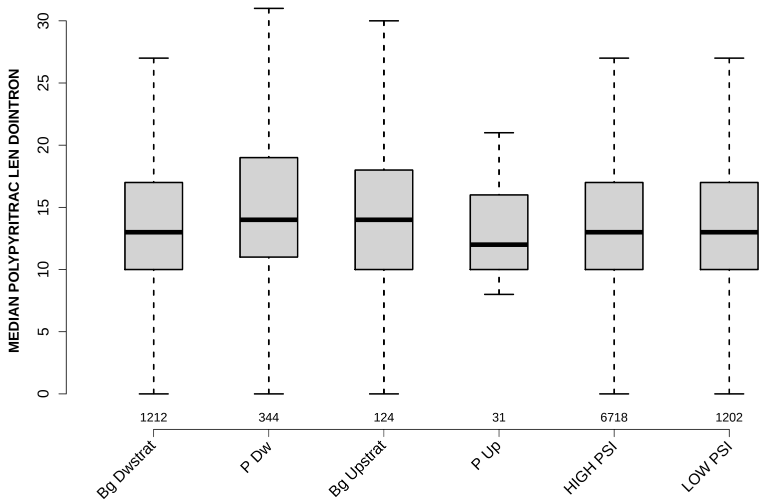

Significant results from Mann-Whitney U test:

- `Bg_Dwstrat` vs `P_Dw` : 0.0200431
  mean: 14.5421 < 15.5189 , median: 13 < 14
- `P_Dw` vs `P_Up` : 0.0457393
  mean: 15.5189 > 14 , median: 14 > 12
- `P_Dw` vs `HIGH_PSI` : 0.00453105
  mean: 15.5189 > 14.5534 , median: 14 > 13
- `P_Dw` vs `LOW_PSI` : 0.00927176
  mean: 15.5189 > 14.4859 , median: 14 > 13

#### 6.57 MEDIAN POLYPYRITRAC SCORE DOINTRON

Back to: Overview | ToC

Meaning: like POLYPYRITRAC SCORE MAXBP but median over top-3 predicted BPs

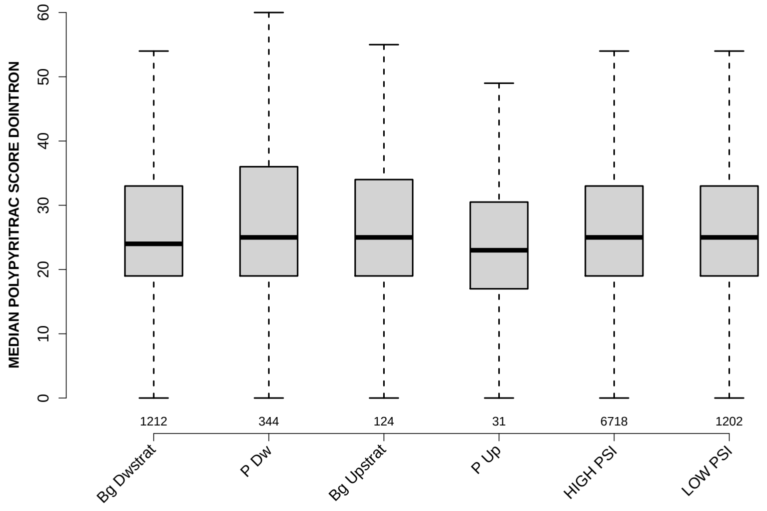

Significant results from Mann-Whitney U test:
- none

#### 6.58 MEDIAN BPSCORE DOINTRON

Back to: Overview | ToC

Meaning: like BPSCORE MAXBP but median over top-3 predicted BPs

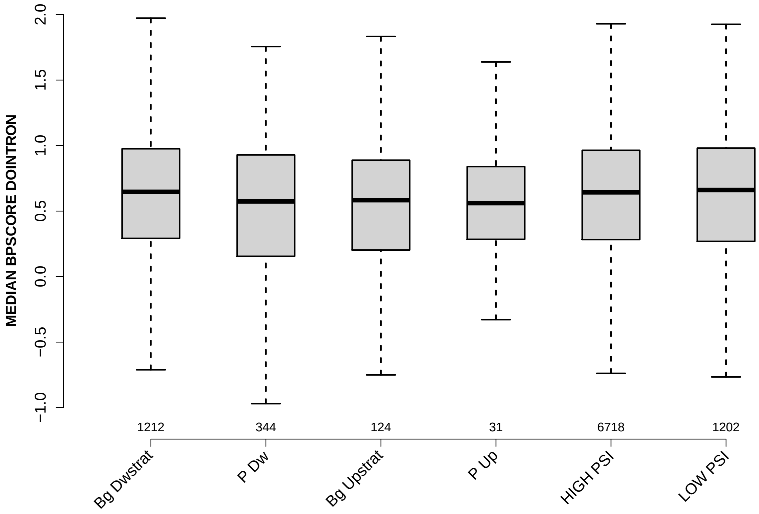

Significant results from Mann-Whitney U test:

- Bg_Dwstrat vs P_Dw : 0.0229115
  mean: 0.555026 > 0.417852 , median: 0.64671 > 0.574329
- P_Dw vs HIGH_PSI : 0.0298713
  mean: 0.417852 < 0.539659 , median: 0.574329 < 0.643701
- P_Dw vs LOW_PSI : 0.029582
  mean: 0.417852 < 0.553182 , median: 0.574329 < 0.661029

#### 6.59 MEDIAN TR LENGTH

Back to: Overview | ToC

Meaning: median length of transcripts the exon occurs in

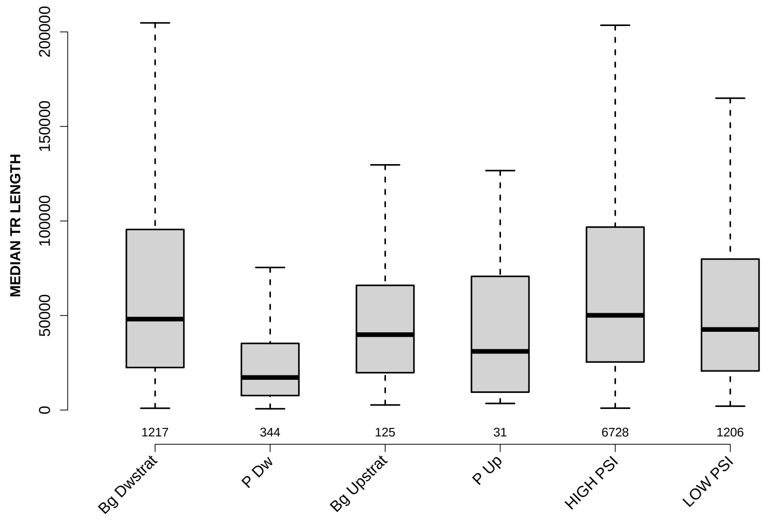

Significant results from Mann-Whitney U test:

- Bg_Dwstrat vs P_Dw : 2.237e-40
  mean: 76598.8488 > 30452.0988 , median: 48106 > 17229.5
- Bg_Dwstrat vs Bg_Upstrat : 0.0197804
  mean: 76598.8488 > 53563.076 , median: 48106 > 39881
- Bg_Dwstrat vs P_Up : 0.0381893
  mean: 76598.8488 > 55806.7581 , median: 48106 > 31039
- Bg_Dwstrat vs LOW_PSI : 0.0108861
  mean: 76598.8488 > 63093.0896 , median: 48106 > 42619
- P_Dw vs Bg_Upstrat : 2.02972e-10
  mean: 30452.0988 < 53563.076 , median: 17229.5 < 39881
- P_Dw vs HIGH_PSI : 8.48917e-57
  mean: 30452.0988 < 76543.0673 , median: 17229.5 < 50110
- P_Dw vs LOW_PSI : 6.11649e-36
  mean: 30452.0988 < 63093.0896 , median: 17229.5 < 42619
- Bg_Upstrat vs HIGH_PSI : 0.00156512
  mean: 53563.076 < 76543.0673 , median: 39881 < 50110
- P_Up vs HIGH_PSI : 0.0153908
  mean: 55806.7581 < 76543.0673 , median: 31039 < 50110
- HIGH_PSI vs LOW_PSI : 1.00574e-07
  mean: 76543.0673 > 63093.0896 , median: 50110 > 42619

#### 6.60 MEDIAN EXON NUMBER

Back to: Overview | ToC

Meaning: ... of transcripts where exon was found in

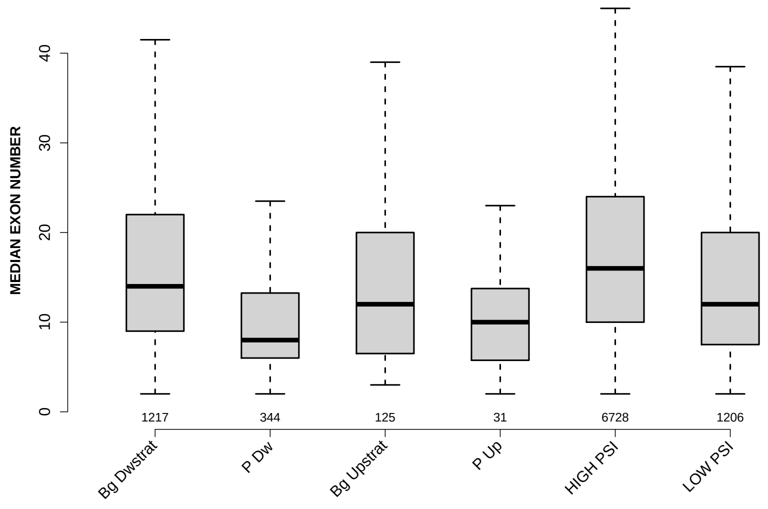

Significant results from Mann-Whitney U test:

- Bg_Dwstrat vs P_Dw : 4.59242e-26
  mean: 17.6561 > 10.7398 , median: 14 > 8
- Bg_Dwstrat vs Bg_Upstrat : 0.00714264
  mean: 17.6561 > 14.924 , median: 14 > 12
- Bg_Dwstrat vs P_Up : 0.00407682
  mean: 17.6561 > 12.1935 , median: 14 > 10
- Bg_Dwstrat vs HIGH_PSI : 1.45932e-06
  mean: 17.6561 < 19.2398 , median: 14 < 16
- Bg_Dwstrat vs LOW_PSI : 2.74226e-06
  mean: 17.6561 > 15.2952 , median: 14 > 12
- P_Dw vs Bg_Upstrat : 0.00111024
  mean: 10.7398 < 14.924 , median: 8 < 12
- P_Dw vs HIGH_PSI : 3.88067e-47
  mean: 10.7398 < 19.2398 , median: 8 < 16
- P_Dw vs LOW_PSI : 8.20999e-14
  mean: 10.7398 < 15.2952 , median: 8 < 12
- Bg_Upstrat vs HIGH_PSI : 1.30379e-05
  mean: 14.924 < 19.2398 , median: 12 < 16
- P_Up vs HIGH_PSI : 0.00023009
  mean: 12.1935 < 19.2398 , median: 10 < 16

- HIGH_PSI vs LOW_PSI : 6.63277e-28
  mean: 19.2398 > 15.2952 , median: 16 > 12

#### 6.61 EXON MEDIANRELATIVERANK

Back to: Overview | ToC

Meaning: relative rank = rank / number of all exons in transcript, is between 0 and 1

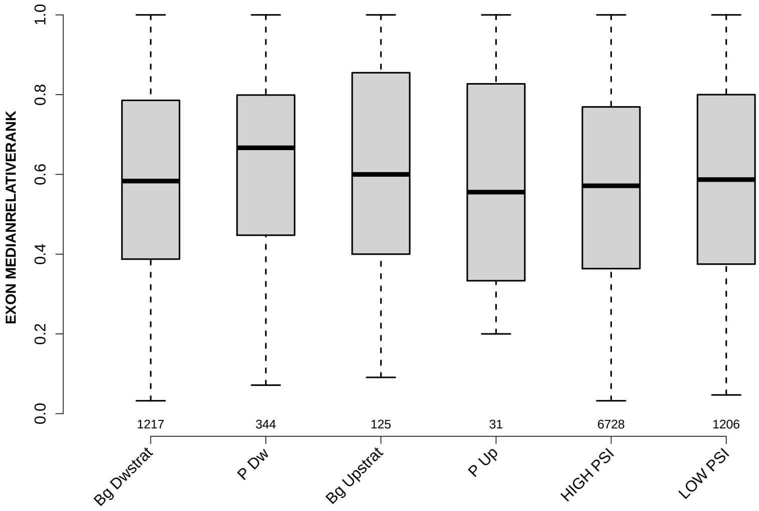

Significant results from Mann-Whitney U test:

- `Bg_Dwstrat` vs `P_Dw` : 0.00314392
  mean: 0.573359 < 0.619939 , median: 0.583333 < 0.666667
- `P_Dw` vs `HIGH_PSI` : 1.97356e-05
  mean: 0.619939 > 0.560021 , median: 0.666667 > 0.571429
- `P_Dw` vs `LOW_PSI` : 0.02373
  mean: 0.619939 > 0.583309 , median: 0.666667 > 0.586975
- `Bg_Upstrat` vs `HIGH_PSI` : 0.0353678
  mean: 0.609473 > 0.560021 , median: 0.6 > 0.571429
- `HIGH_PSI` vs `LOW_PSI` : 0.00515261
  mean: 0.560021 < 0.583309 , median: 0.571429 < 0.586975

#### 6.62 EXON MEDIANRELATIVERANK 3BINS

Back to: Overview | ToC

Meaning: median bin into which EXON MEDIANRELATIVERANK falls when binning 0-1 into 3 bins

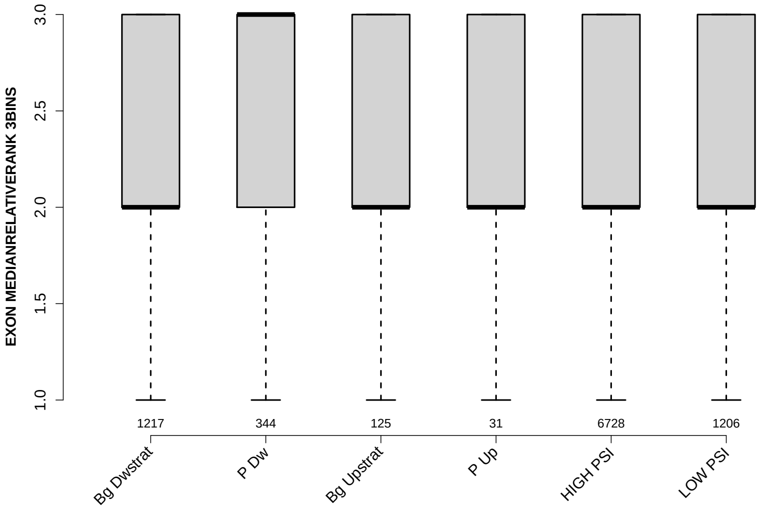

Significant results from Mann-Whitney U test:

- Bg_Dwstrat vs P_Dw : 3.46809e-05
  mean: 2.2219 < 2.4128 , median: 2 < 3
- P_Dw vs HIGH_PSI : 9.59386e-08
  mean: 2.4128 > 2.1885 , median: 3 > 2
- P_Dw vs LOW_PSI : 0.000116147
  mean: 2.4128 > 2.2322 , median: 3 > 2

#### 6.63 EXON MEDIANRELATIVERANK 5BINS

Back to: Overview | ToC

Meaning: similar to EXON MEDIANRELATIVERANK 3BINS with 5 bins

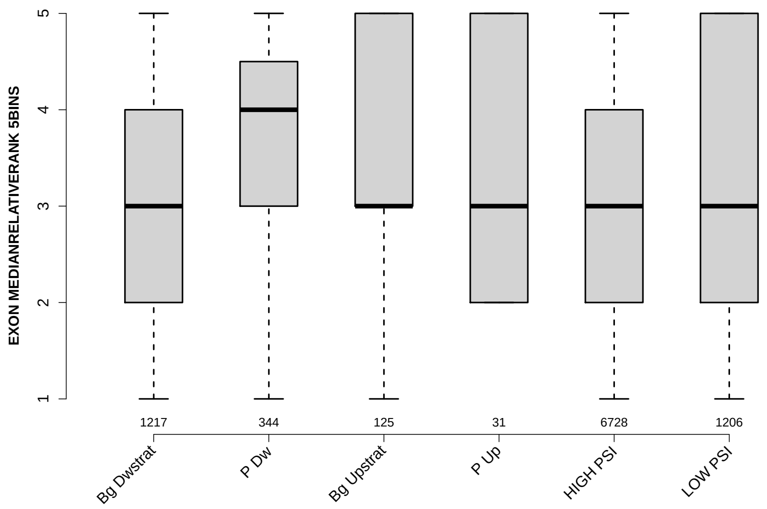

Significant results from Mann-Whitney U test:

- `Bg_Dwstrat` vs `P_Dw` : 0.00494934
  mean: 3.3739 < 3.5959 , median: 3 < 4
- `P_Dw` vs `HIGH_PSI` : 2.85755e-05
  mean: 3.5959 > 3.3017 , median: 4 > 3
- `P_Dw` vs `LOW_PSI` : 0.0145859
  mean: 3.5959 > 3.398 , median: 4 > 3
- `HIGH_PSI` vs `LOW_PSI` : 0.0160538
  mean: 3.3017 < 3.398 , median: 3 = 3

#### 6.64 EXON MEDIANRELATIVERANK 10BINS

Back to: Overview | ToC

Meaning: similar to EXON MEDIANRELATIVERANK 3BINS with 10 bins

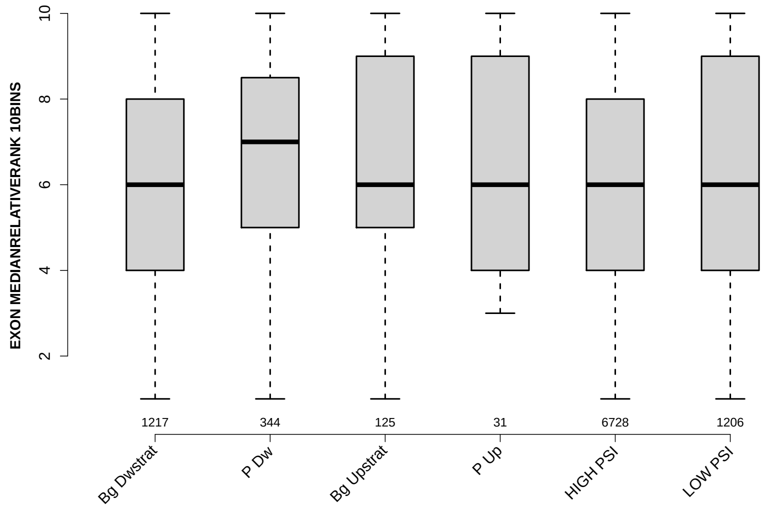

Significant results from Mann-Whitney U test:

- `Bg_Dwstrat` vs `P_Dw` : 0.00382489
  mean: 6.2818 < 6.7355 , median: 6 < 7
- `P_Dw` vs `HIGH_PSI` : 1.48598e-05
  mean: 6.7355 > 6.1315 , median: 7 > 6
- `P_Dw` vs `LOW_PSI` : 0.0152356
  mean: 6.7355 > 6.3358 , median: 7 > 6
- `HIGH_PSI` vs `LOW_PSI` : 0.00975755
  mean: 6.1315 < 6.3358 , median: 6 = 6

#### 6.65 NTRS ALL FOR GENE

Back to: Overview | ToC

Meaning: number of transcripts of gene where the exon was found in

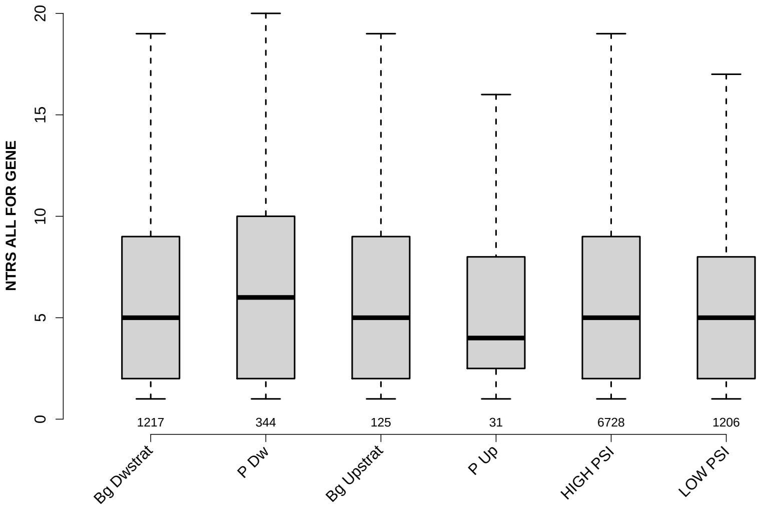

Significant results from Mann-Whitney U test:

- `Bg_Dwstrat` vs `P_Dw` : 0.0115767
  mean: 6.1044 < 6.9157 , median: 5 < 6
- `P_Dw` vs `HIGH_PSI` : 0.00954971
  mean: 6.9157 > 6.0902 , median: 6 > 5
- `P_Dw` vs `LOW_PSI` : 0.000212392
  mean: 6.9157 > 5.6385 , median: 6 > 5
- `HIGH_PSI` vs `LOW_PSI` : 0.00904806
  mean: 6.0902 > 5.6385 , median: 5 = 5

#### 6.66 PROP FIRST EXON

Back to: Overview | ToC

Meaning: NTRS WITH EXON AS FIRST EXON / NTRS WITH EXON

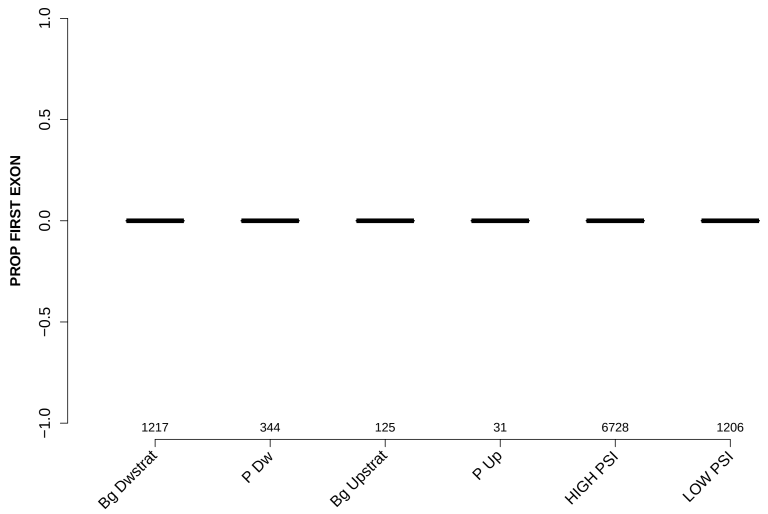

Significant results from Mann-Whitney U test:

- `Bg_Dwstrat` vs `P_Dw` : 0.00513249
  mean: 0.0209927 < 0.0330749 , median: 0 = 0
- `Bg_Dwstrat` vs `LOW_PSI` : 7.77165e-07
  mean: 0.0209927 > 0.009062 , median: 0 = 0
- `P_Dw` vs `Bg_Upstrat` : 0.0168868
  mean: 0.0330749 > 0.0130984 , median: 0 = 0
- `P_Dw` vs `HIGH_PSI` : 0.00804627
  mean: 0.0330749 > 0.0235694 , median: 0 = 0
- `P_Dw` vs `LOW_PSI` : 9.76062e-12
  mean: 0.0330749 > 0.009062 , median: 0 = 0
- `P_Up` vs `LOW_PSI` : 0.00170866
  mean: 0.054315 > 0.009062 , median: 0 = 0
- `HIGH_PSI` vs `LOW_PSI` : 1.39953e-10
  mean: 0.0235694 > 0.009062 , median: 0 = 0

#### 6.67 PROP LAST EXON

Back to: Overview | ToC

Meaning: NTRS WITH EXON AS LAST EXON / NTRS WITH EXON

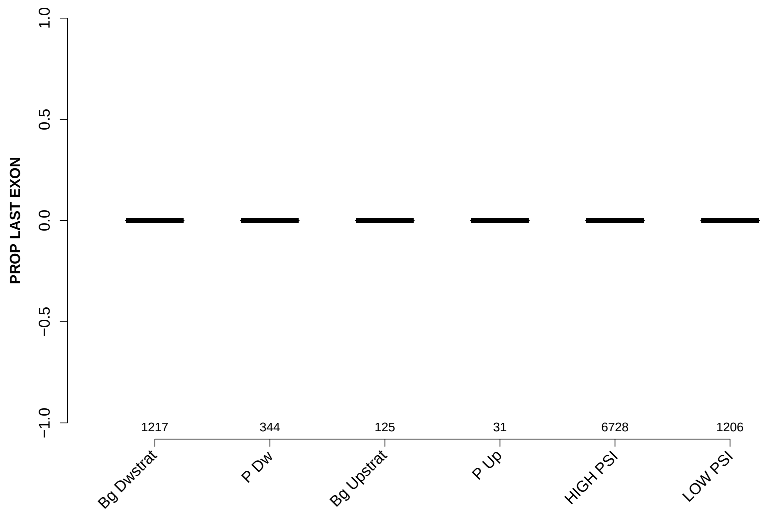

Significant results from Mann-Whitney U test:

- Bg_Dwstrat vs P_Dw : 0.000447823
  mean: 0.0423357 < 0.0558746 , median: 0 = 0
- Bg_Dwstrat vs LOW_PSI : 2.34331e-16
  mean: 0.0423357 > 0.0128661 , median: 0 = 0
- P_Dw vs Bg_Upstrat : 0.00211932
  mean: 0.0558746 > 0.023063 , median: 0 = 0
- P_Dw vs HIGH_PSI : 1.30666e-06
  mean: 0.0558746 > 0.0360322 , median: 0 = 0
- P_Dw vs LOW_PSI : 8.56608e-26
  mean: 0.0558746 > 0.0128661 , median: 0 = 0
- Bg_Upstrat vs LOW_PSI : 0.0074979
  mean: 0.023063 > 0.0128661 , median: 0 = 0
- HIGH_PSI vs LOW_PSI : 1.05396e-17
  mean: 0.0360322 > 0.0128661 , median: 0 = 0

#### 6.68 PROP INTERNAL EXON

Back to: Overview | ToC

Meaning: NTRS WITH EXON AS INTERNAL EXON / NTRS WITH EXON

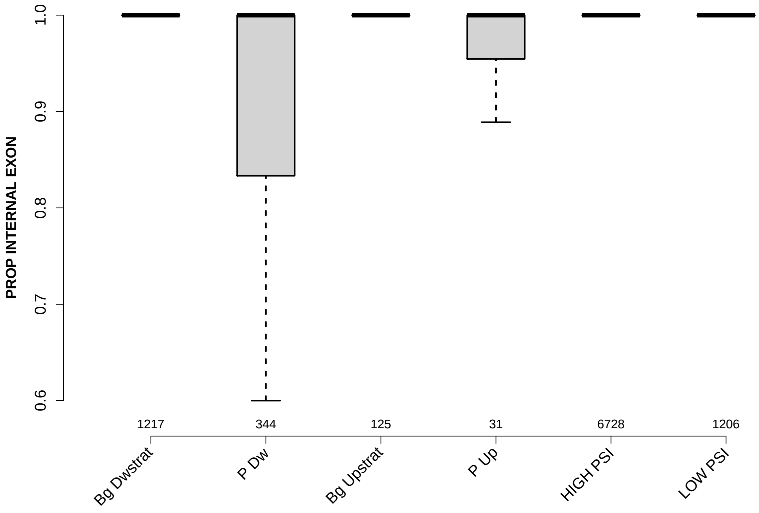

Significant results from Mann-Whitney U test:

- Bg_Dwstrat vs P_Dw : 4.97956e-05
  mean: 0.936857 > 0.91105 , median: 1 = 1
- Bg_Dwstrat vs LOW_PSI : 6.86973e-20
  mean: 0.936857 < 0.978072 , median: 1 = 1
- P_Dw vs Bg_Upstrat : 0.000244097
  mean: 0.91105 < 0.963839 , median: 1 = 1
- P_Dw vs HIGH_PSI : 8.73928e-07
  mean: 0.91105 < 0.940398 , median: 1 = 1
- P_Dw vs LOW_PSI : 1.15832e-30
  mean: 0.91105 < 0.978072 , median: 1 = 1
- Bg_Upstrat vs LOW_PSI : 0.0053824
  mean: 0.963839 < 0.978072 , median: 1 = 1
- P_Up vs LOW_PSI : 0.00128928
  mean: 0.912723 < 0.978072 , median: 1 = 1
- HIGH_PSI vs LOW_PSI : 4.63095e-25
  mean: 0.940398 < 0.978072 , median: 1 = 1

#### 6.69 PROP EXON IN UTR

Back to: Overview | ToC

Meaning: NTRS WITH EXON IN UTR / NTRS WITH EXON

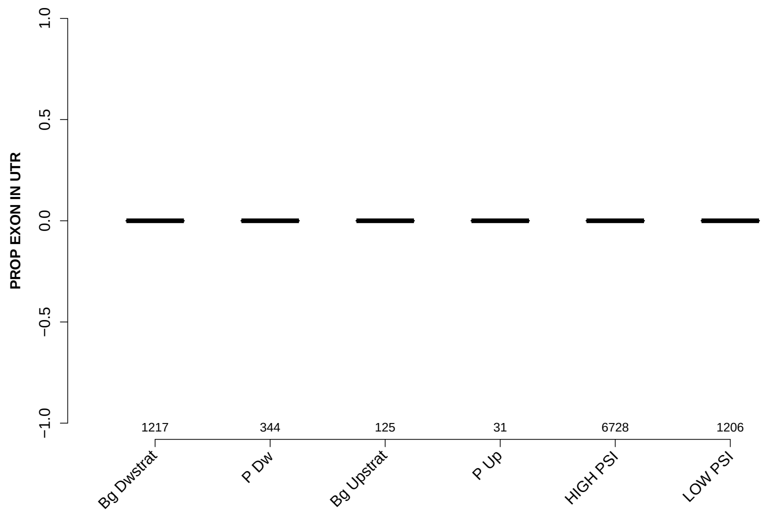

Significant results from Mann-Whitney U test:

- Bg_Dwstrat vs P_Dw : p value = NA
- Bg_Dwstrat vs Bg_Upstrat : p value = NA
- Bg_Dwstrat vs P_Up : p value = NA
- Bg_Dwstrat vs HIGH_PSI : p value = NA
- Bg_Dwstrat vs LOW_PSI : p value = NA
- P_Dw vs Bg_Upstrat : p value = NA
- P_Dw vs P_Up : p value = NA
- P_Dw vs HIGH_PSI : p value = NA
- P_Dw vs LOW_PSI : p value = NA
- Bg_Upstrat vs P_Up : p value = NA
- Bg_Upstrat vs HIGH_PSI : p value = NA
- Bg_Upstrat vs LOW_PSI : p value = NA
- P_Up vs HIGH_PSI : p value = NA
- P_Up vs LOW_PSI : p value = NA
- HIGH_PSI vs LOW_PSI : p value = NA

### Comparison of exons grouped into: Bg-Dwstrat, P-Dw, Bg-Upstrat, P-Up, HIGH-PSI, LOW-PSI

September 16, 2020
Matt version 1.3.0

#### Contents

**1 Infos** 4

**2 Warning: Please read this note carefully** 4

**3 Notes for publishing results** 4

**4 Data sets** 5

**5 Overview: Features with statistically significant differences (p-val $\leq 0.05$)** 6

**6 Details: Box plots and statistical assessments for all features** 20

### 1 Infos

Visualizations of exon features for different groups of exons. Each exon occurs in exactly one gene, but might occur in several transcripts of that gene. Hence, for some features like the exon length, there is exactly one value for each exon. For other features, e.g., length of the up-stream exon(s), which could be different in different transcripts, there might be several values for each exon. Consequently, in the latter cases, the median of these value gets reported.

### 2 Warning: Please read this note carefully

Please keep in mind that some features might affect other features. Especially: all branch-point features get extracted from sub-sequences of introns, by standard the last 150 nt at the 3' end of each intron (if you haven't changed this) always neglecting the first 20 nt at their 5' end. If introns of one set are especially short, i.e., many are shorter then these 150 nt, then the shorter intron length might affect branch-point features. For example, there might be less branch points found in shorter introns or their distance to the 3' intron ends might be generally shorter simply because of their shorter intron length.

### 3 Notes for publishing results

The Matt paper: *Matt: Unix tools for alternative splicing analysis, A. Gohr, M. Irimia, Bioinformatics, 2018, bty606, DOI: 10.1093/bioinformatics/bty606*
When publishing results wrt. splice site strengths which you determined for your data using matt, please cite: *Maximum entropy modeling of short sequence motifs with applications to RNA splicing signals, Yeo et al., 2003, DOI: 10.1089/1066527041410418*
When publishing results wrt. branch point features which you determined for your data with matt, please cite: *Genome-wide association between branch point properties and alternative splicing, Corvelo et al., 2010, DOI: 10.1371/journal.pcbi.1001016*
When publishing results with respect to the binding strength of the human Sf1 splicing factor, you might refert to where the Sf1 binding motif comes from: *Analysis of in situ pre-mRNA targets of human splicing factor SF1 reveals a function in alternative splicing, Margherita Corioni, Nicolas Antih, Goranka Tanackovic, Mihaela Zavolan, and Angela Kramer, 2011, DOI: 10.1093/nar/gkq1042*
The Sf1 binding motif is described in supplement, page 13, table S2: Weight matrix of the binding specificity of SF1.

#### 4 Data sets

Input file:
../bta_final.tab
Selection criteria for defining exons groups:
Bg_Dwstrat : having value Bg_Dwstrat in column GROUP
P_Dw : having value P_Dw in column GROUP
Bg_Upstrat : having value Bg_Upstrat in column GROUP
P_Up : having value P_Up in column GROUP
HIGH_PSI : having value HIGH_PSI in column GROUP
LOW_PSI : having value LOW_PSI in column GROUP
Exon duplicates removal: yes
Numbers of exons per group before / after neglecting exons which were not found in GTF file (gene annotation). For the comparisons only exons which were found in the gene annotation are used. These numbers might change slightly for each feature if NAs occur.
Bg_Dwstrat: 2584 / 2554
P_Dw: 646 / 640
Bg_Upstrat: 604 / 590
P_Up: 152 / 148
HIGH_PSI: 5108 / 5078
LOW_PSI: 955 / 950

### 5 Overview: Features with statistically significant differences (p-val $\leq 0.05$)

MAXENTSCR HSAMODEL 3SS

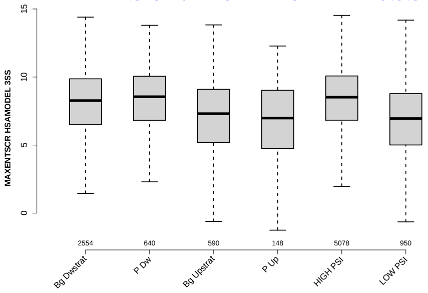

MAXENTSCR HSAMODEL 5SS

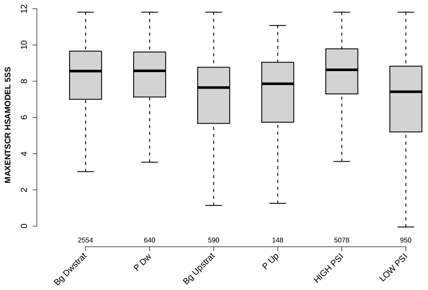

DOINTRON MEDIANLENGTH

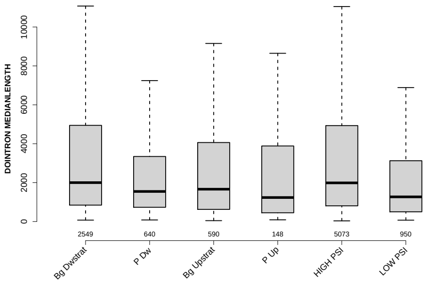

PYRIMIDINECONT MAXBP UPINTRON

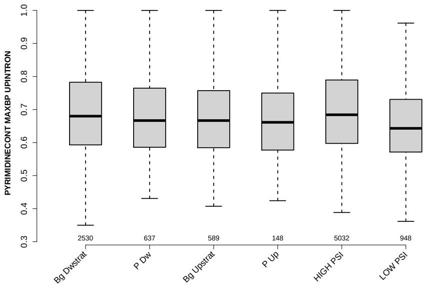

#### UPINTRON MEDIANLENGTH

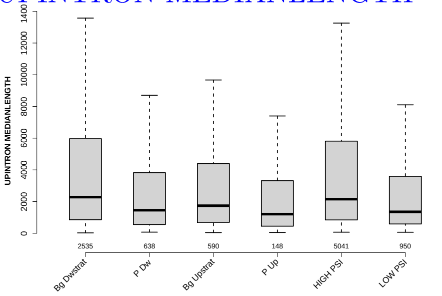

#### MEDIAN TR LENGTH

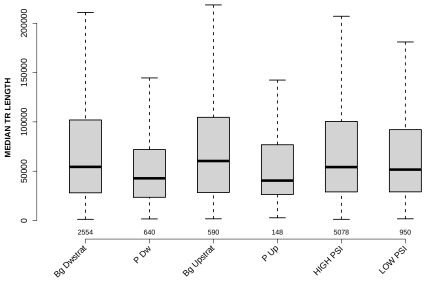

#### RATIO DOEXON EXON LENGTH

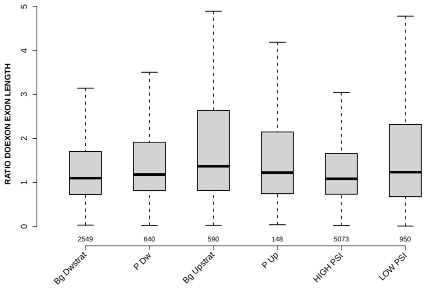

#### EXON LENGTH

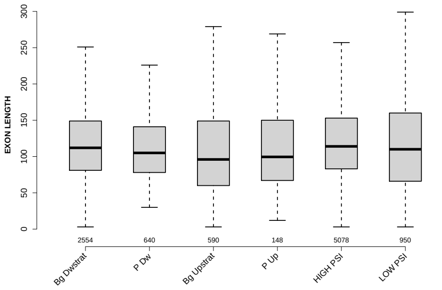

#### PROP EXON IN UTR

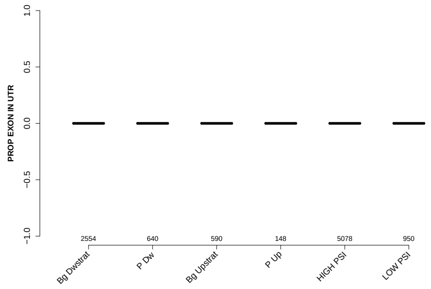

#### RATIO DOINTRON EXON LENGTH

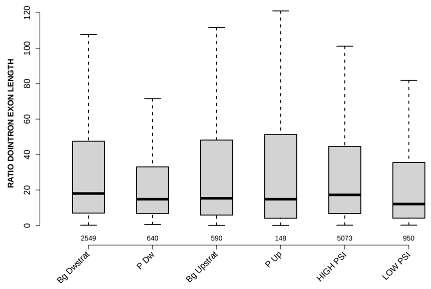

#### RATIO UPINTRON EXON LENGTH

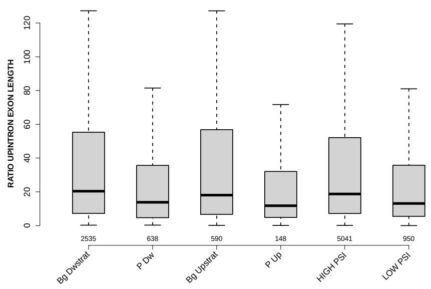

#### DIST FROM MAXBP TO 3SS UPINTRON

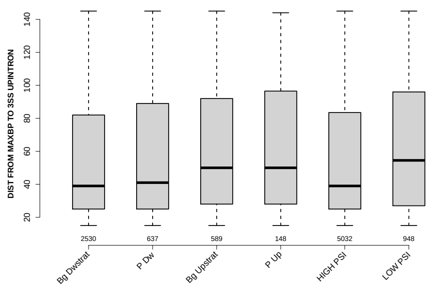

### RATIO UPEXON EXON LENGTH

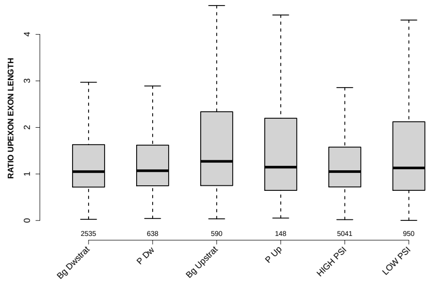

### MEDIAN PYRIMIDINECONT UPINTRON

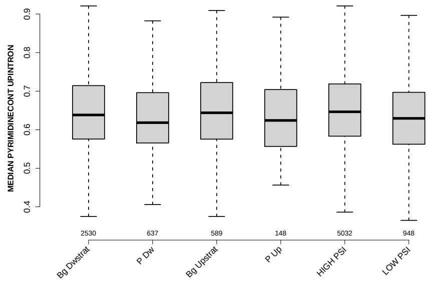

### UPEXON GCC

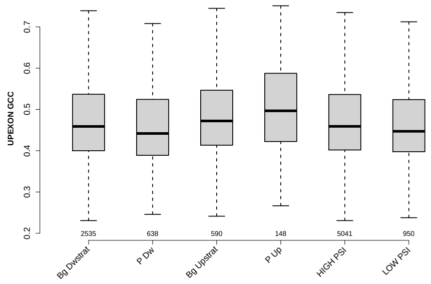

### GCC 3SS 20INT10EX

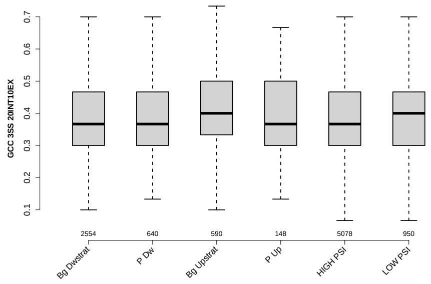

#### UP 5SS 20INT10EX GCC

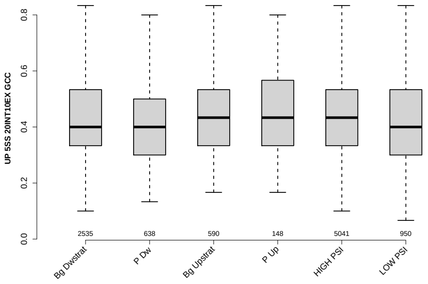

#### POLYPYRITRAC SCORE MAXBP UPINTRON

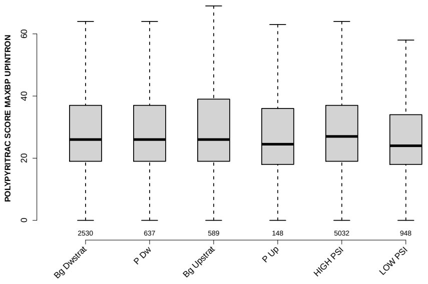

#### EXON MEDIANRELATIVERANK

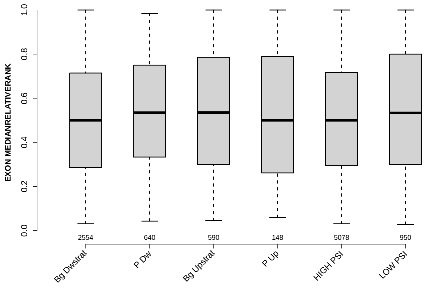

#### MEDIAN EXON NUMBER

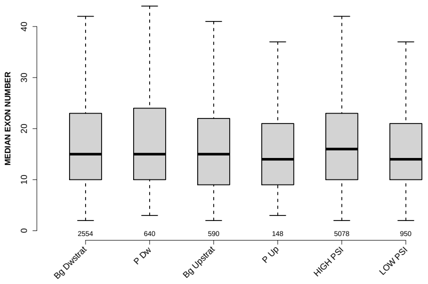

#### EXON MEDIANRELATIVERANK 10BINS

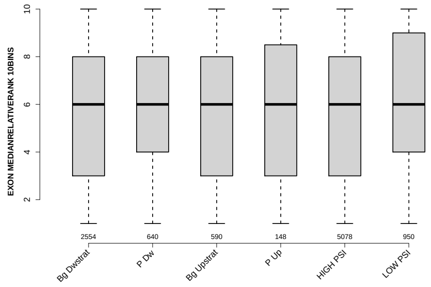

#### EXON MEDIANRELATIVERANK 5BINS

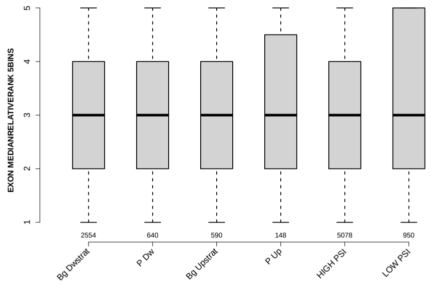

#### NUM PREDICTED BPS UPINTRON

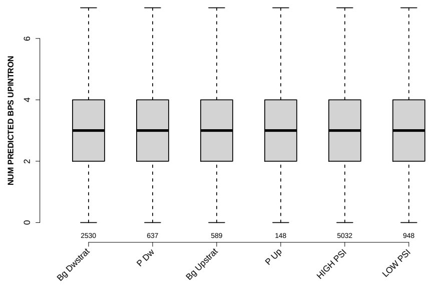

#### MEDIAN BPSCORE UPINTRON

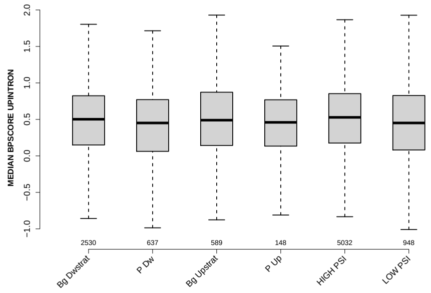

### DOEXON GCC

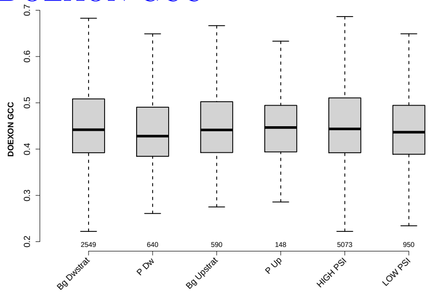

### DOINTRON GCC

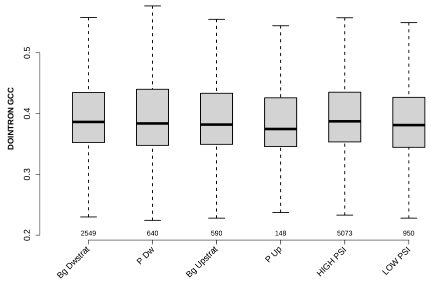

### MEDIAN DIST FROM BP TO 3SS UPINTRON

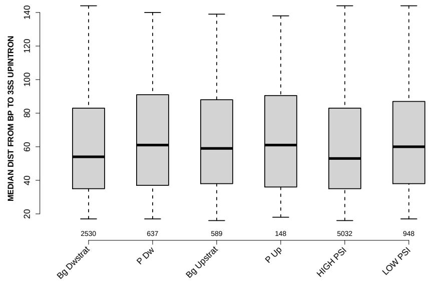

### MEDIAN POLYPYRITRAC OFFSET UPINTRON

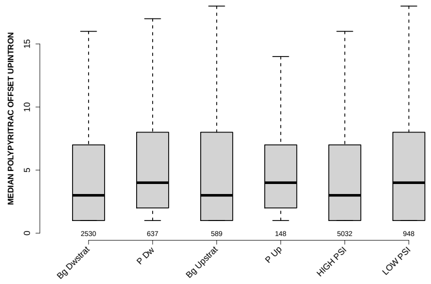

### EXON MEDIANRELATIVERANK 3BINS

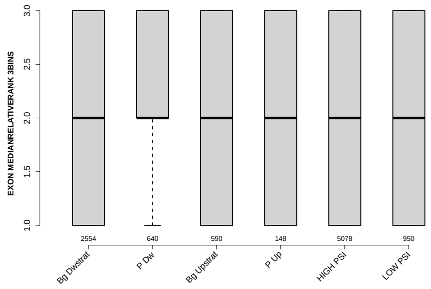

### MEDIAN POLYPYRITRAC LEN UPINTRON

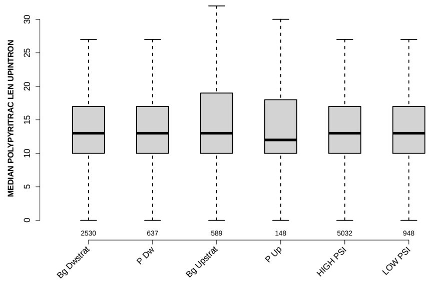

### PROP FIRST EXON

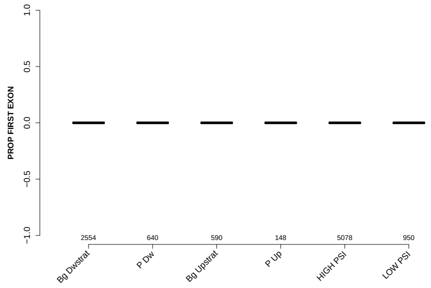

### RATIO UPEXON EXON GCC

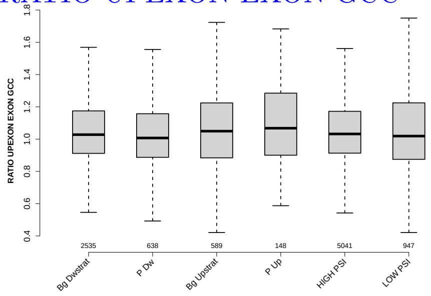

### NTRS ALL FOR GENE

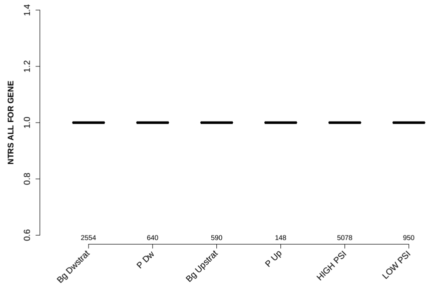

### POLYPYRITRAC LEN MAXBP DOINTRON

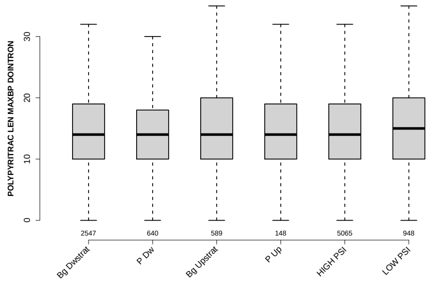

### PROP INTERNAL EXON

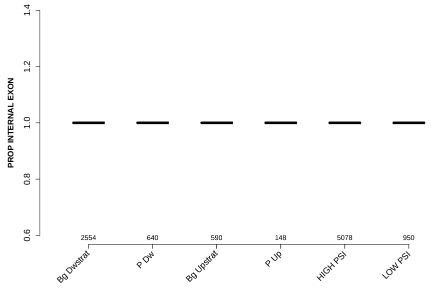

### PYRIMIDINECONT MAXBP DOINTRON

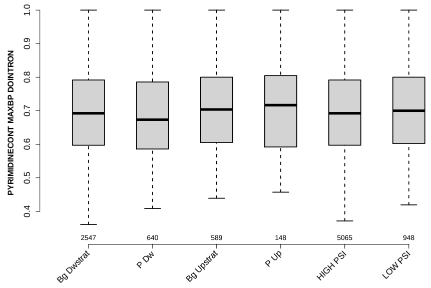

#### BPSCORE MAXBP UPINTRON

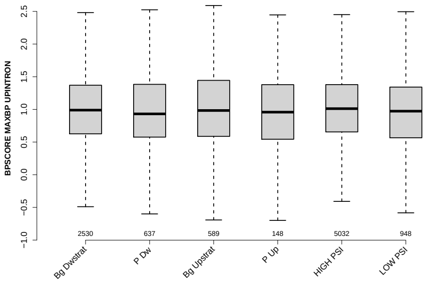

#### MAXENTSCR HSAMODEL UPSTRM 5SS

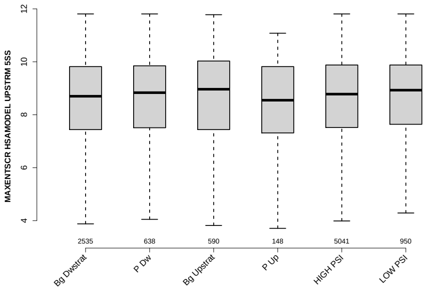

#### POLYPYRITRAC LEN MAXBP UPINTRON

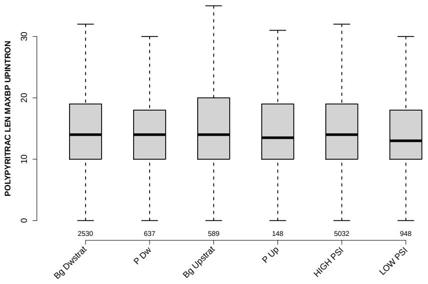

#### GCC 5SS 20INT10EX

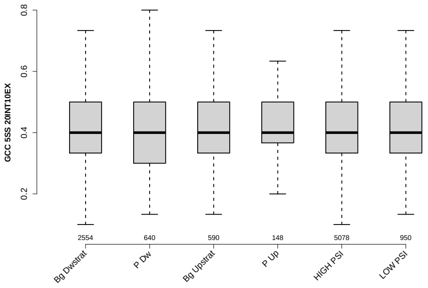

### MEDIAN PYRIMIDINECONT DOINTRON

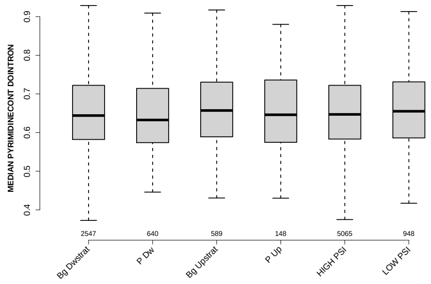

### POLYPYRITRAC SCORE MAXBP DOINTRON

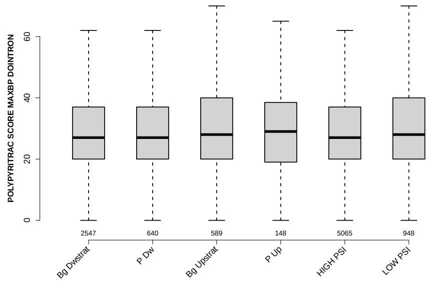

### UPINTRON GCC

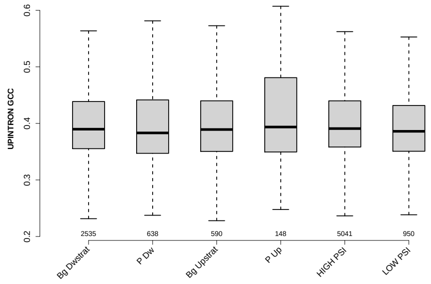

### DOEXON MEDIANLENGTH

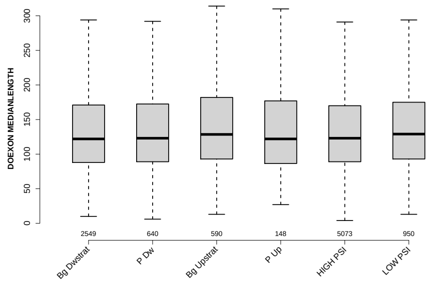

### EXON GCC

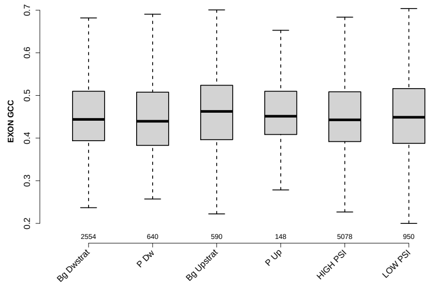

### MEDIAN SCORE FOR BPSEQ DOINTRON

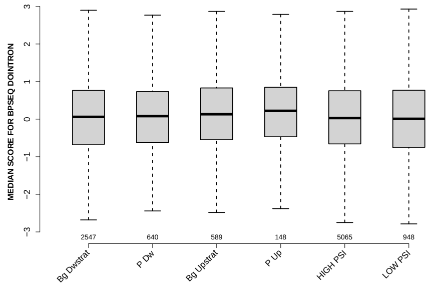

### RATIO DOINTRON EXON GCC

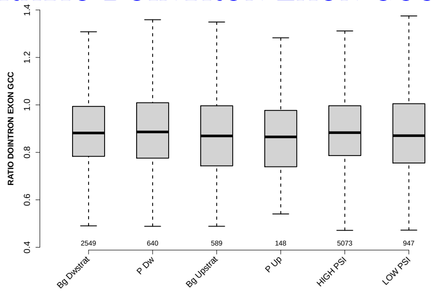

### POLYPYRITRAC OFFSET MAXBP UPINTRON

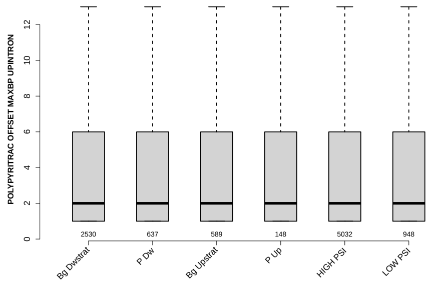

#### MEDIAN POLYPYRITRAC SCORE UPINTRON

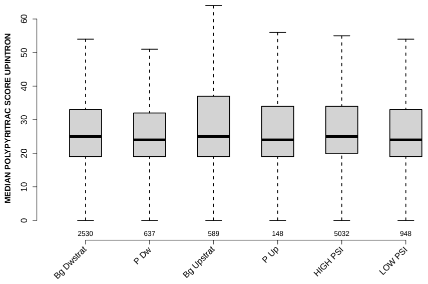

#### RATIO DOEXON EXON GCC

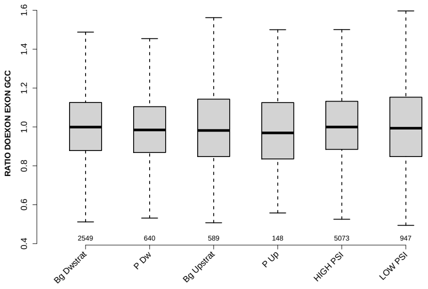

#### MEDIAN POLYPYRITRAC LEN DOINTRON

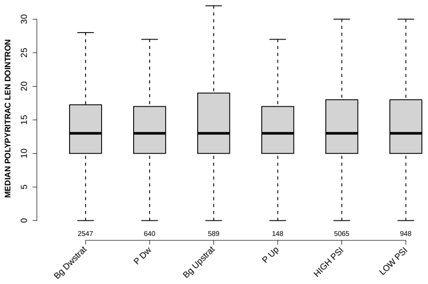

#### BPSCORE MAXBP DOINTRON

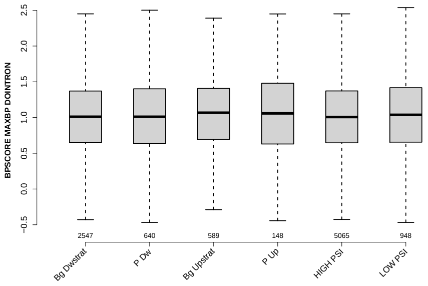

### MEDIAN POLYPYRITRAC OFFSET DOINTRON

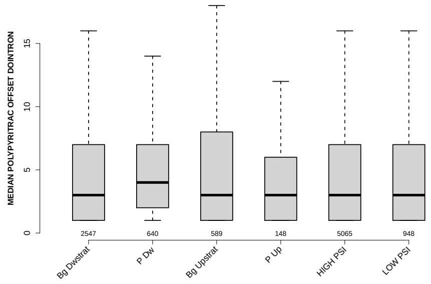

### PROP LAST EXON

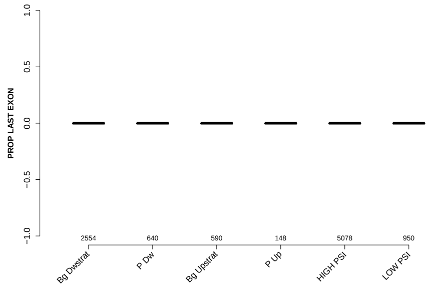

### MEDIAN POLYPYRITRAC SCORE DOINTRON

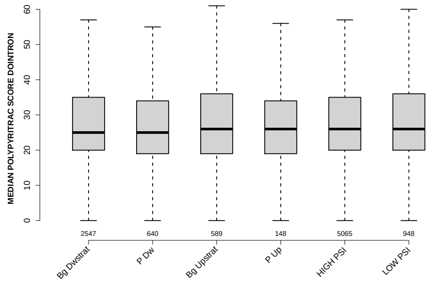

### 6 Details: Box plots and statistical assessments for all features

#### 6.1 EXON LENGTH

Back to: Overview | ToC

Meaning:

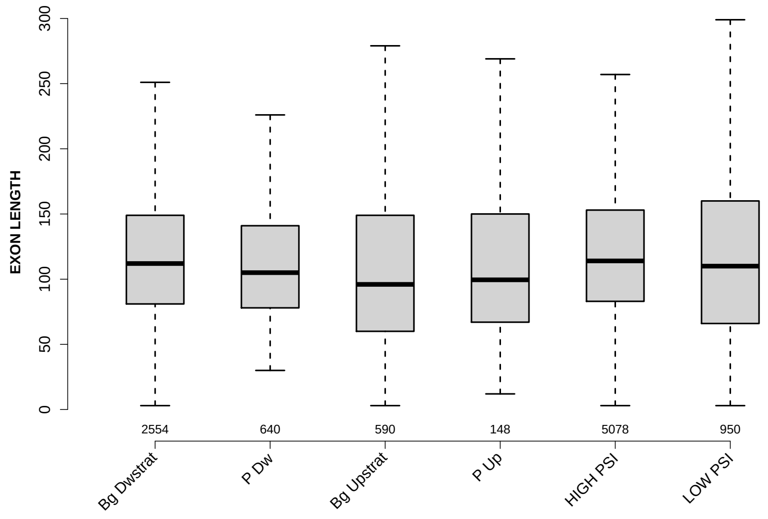

Significant results from Mann-Whitney U test:

- `Bg_Dwstrat` vs `P_Dw` : 0.0103854
  mean: 134.0732 > 129.5328 , median: 112 > 105
- `Bg_Dwstrat` vs `Bg_Upstrat` : 6.14556e-08
  mean: 134.0732 < 134.6254 , median: 112 > 96
- `Bg_Dwstrat` vs `P_Up` : 0.0321314
  mean: 134.0732 < 147.1486 , median: 112 > 99.5
- `Bg_Dwstrat` vs `LOW_PSI` : 0.0199155
  mean: 134.0732 < 143.5189 , median: 112 > 110
- `P_Dw` vs `Bg_Upstrat` : 0.00554072
  mean: 129.5328 < 134.6254 , median: 105 > 96
- `P_Dw` vs `HIGH_PSI` : 0.000233189
  mean: 129.5328 < 131.7564 , median: 105 < 114
- `Bg_Upstrat` vs `HIGH_PSI` : 1.25961e-10
  mean: 134.6254 > 131.7564 , median: 96 < 114
- `Bg_Upstrat` vs `LOW_PSI` : 0.0173428
  mean: 134.6254 < 143.5189 , median: 96 < 110

- `P_Up` vs `HIGH_PSI` : 0.00915702
  mean: $147.1486 > 131.7564$ , median: $99.5 < 114$
- `HIGH_PSI` vs `LOW_PSI` : 0.000655211
  mean: $131.7564 < 143.5189$ , median: $114 > 110$

#### 6.2 UPEXON MEDIANLENGTH

Back to: Overview | ToC

Meaning: median length of up-stream exon

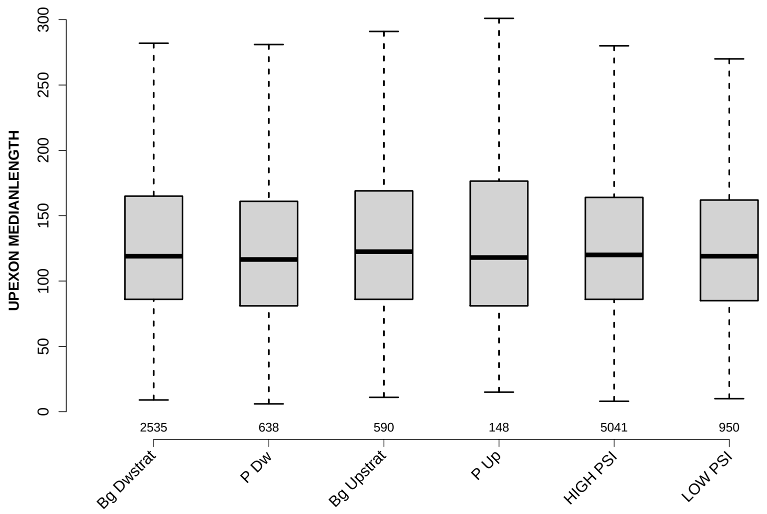

Significant results from Mann-Whitney U test:

- none

#### 6.3 DOEXON MEDIANLENGTH

Back to: Overview | ToC

Meaning: median length of down-stream exon

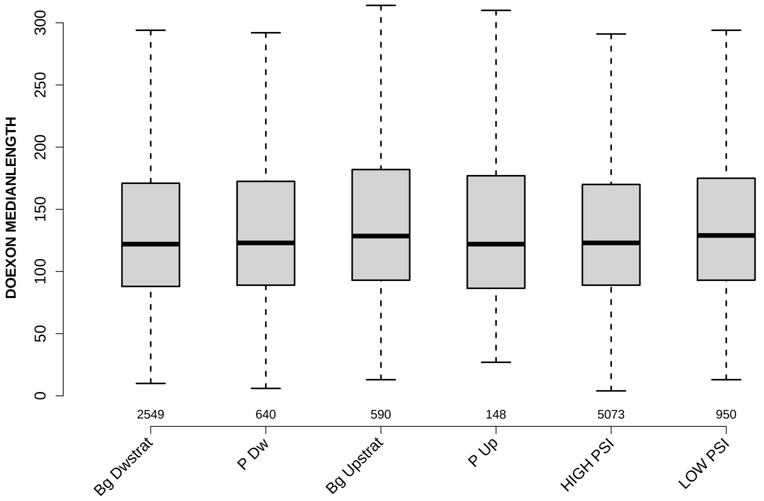

Significant results from Mann-Whitney U test:

- `Bg_Dwstrat` vs `Bg_Upstrat` : 0.0134166
  mean: 191.4563 < 233.1237 , median: 122 < 128.5
- `Bg_Dwstrat` vs `LOW_PSI` : 0.0164616
  mean: 191.4563 < 210.4563 , median: 122 < 129
- `Bg_Upstrat` vs `HIGH_PSI` : 0.0211888
  mean: 233.1237 > 189.6441 , median: 128.5 > 123
- `HIGH_PSI` vs `LOW_PSI` : 0.0273599
  mean: 189.6441 < 210.4563 , median: 123 < 129

#### 6.4 RATIO UPEXON EXON LENGTH

Back to: Overview | ToC

Meaning: median up-stream exon length / exon length

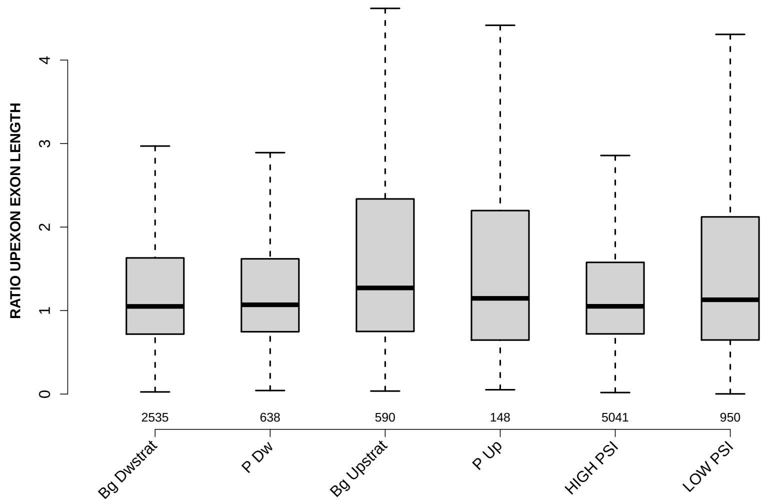

Significant results from Mann-Whitney U test:

- `Bg_Dwstrat` vs `Bg_Upstrat` : 1.4242e-07
  mean: 1.5235 < 2.7066 , median: 1.05 < 1.2712
- `Bg_Dwstrat` vs `LOW_PSI` : 0.0303775
  mean: 1.5235 < 2.8544 , median: 1.05 < 1.1295
- `P_Dw` vs `Bg_Upstrat` : 8.95169e-05
  mean: 1.5662 < 2.7066 , median: 1.0694 < 1.2712
- `Bg_Upstrat` vs `HIGH_PSI` : 4.23331e-09
  mean: 2.7066 > 1.4765 , median: 1.2712 > 1.051
- `Bg_Upstrat` vs `LOW_PSI` : 0.0129771
  mean: 2.7066 < 2.8544 , median: 1.2712 > 1.1295
- `HIGH_PSI` vs `LOW_PSI` : 0.00820343
  mean: 1.4765 < 2.8544 , median: 1.051 < 1.1295

#### 6.5 RATIO DOEXON EXON LENGTH

Back to: Overview | ToC

Meaning: median down-stream exon length / exon length

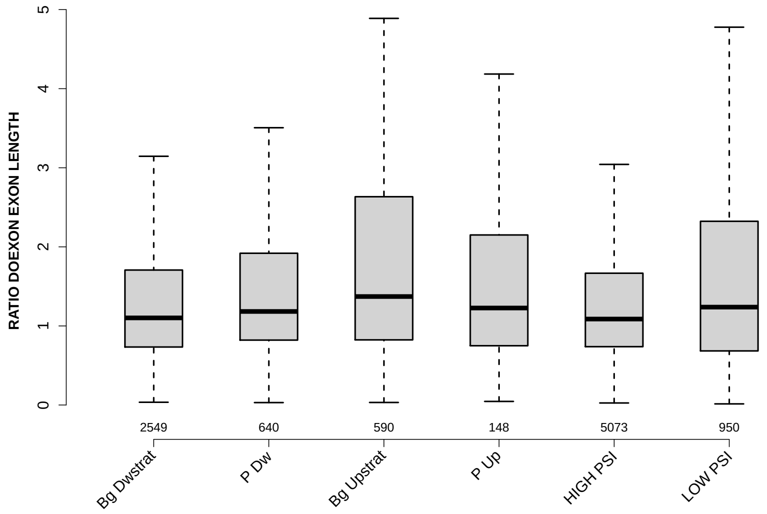

Significant results from Mann-Whitney U test:

- Bg_Dwstrat vs P_Dw : 0.00221219
  mean: 1.9285 > 1.8717 , median: 1.1014 < 1.1828
- Bg_Dwstrat vs Bg_Upstrat : 2.3746e-09
  mean: 1.9285 < 3.1582 , median: 1.1014 < 1.3717
- Bg_Dwstrat vs LOW_PSI : 0.000917111
  mean: 1.9285 < 4.0907 , median: 1.1014 < 1.2378
- P_Dw vs Bg_Upstrat : 0.00343489
  mean: 1.8717 < 3.1582 , median: 1.1828 < 1.3717
- P_Dw vs HIGH_PSI : 0.00017202
  mean: 1.8717 > 1.8533 , median: 1.1828 > 1.087
- Bg_Upstrat vs HIGH_PSI : 1.59815e-11
  mean: 3.1582 > 1.8533 , median: 1.3717 > 1.087
- Bg_Upstrat vs LOW_PSI : 0.0197477
  mean: 3.1582 < 4.0907 , median: 1.3717 > 1.2378
- HIGH_PSI vs LOW_PSI : 5.36648e-05
  mean: 1.8533 < 4.0907 , median: 1.087 < 1.2378

#### 6.6 UPINTRON MEDIANLENGTH

Back to: Overview | ToC

Meaning: median length of up-stream introns

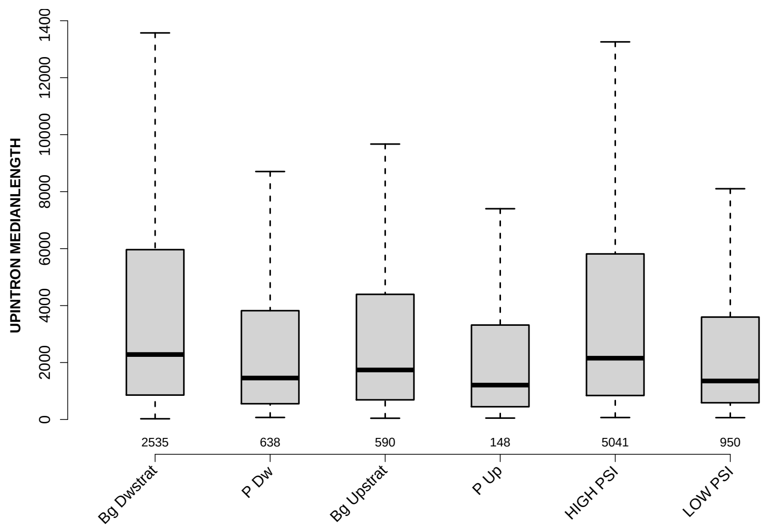

Significant results from Mann-Whitney U test:

- Bg_Dwstrat vs P_Dw : 5.09211e-11
  mean: 6883.3548 > 3642.1787 , median: 2282 > 1456.5
- Bg_Dwstrat vs Bg_Upstrat : 0.000683109
  mean: 6883.3548 > 4819.4322 , median: 2282 > 1740
- Bg_Dwstrat vs P_Up : 1.33438e-05
  mean: 6883.3548 > 3708.0642 , median: 2282 > 1208.5
- Bg_Dwstrat vs LOW_PSI : 1.09709e-15
  mean: 6883.3548 > 3833.9379 , median: 2282 > 1353.5
- P_Dw vs Bg_Upstrat : 0.0123601
  mean: 3642.1787 < 4819.4322 , median: 1456.5 < 1740
- P_Dw vs HIGH_PSI : 1.18249e-10
  mean: 3642.1787 < 6656.6092 , median: 1456.5 < 2153
- Bg_Upstrat vs P_Up : 0.0126377
  mean: 4819.4322 > 3708.0642 , median: 1740 > 1208.5
- Bg_Upstrat vs HIGH_PSI : 0.00195441
  mean: 4819.4322 < 6656.6092 , median: 1740 < 2153
- Bg_Upstrat vs LOW_PSI : 0.00356997
  mean: 4819.4322 > 3833.9379 , median: 1740 > 1353.5
- P_Up vs HIGH_PSI : 2.8064e-05
  mean: 3708.0642 < 6656.6092 , median: 1208.5 < 2153

- `HIGH_PSI` vs `LOW_PSI` : 8.01651e-16
  mean: 6656.6092 > 3833.9379 , median: 2153 > 1353.5

#### 6.7 DOINTRON MEDIANLENGTH

Back to: Overview | ToC

Meaning: median length of down-stream introns

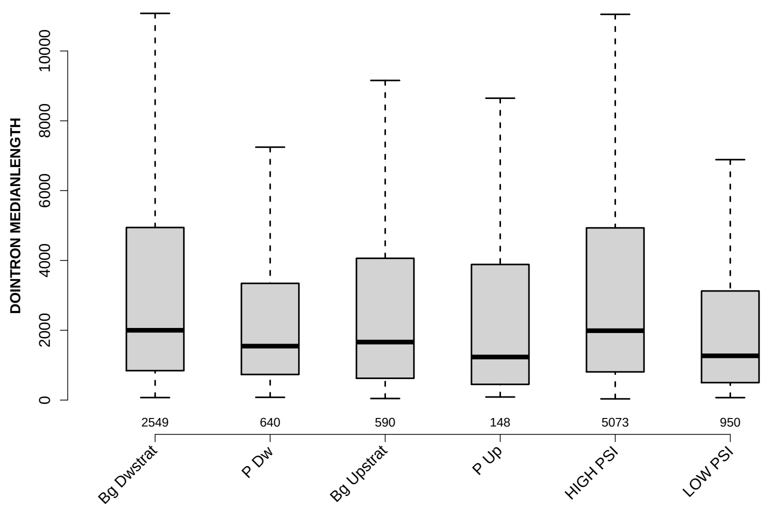

Significant results from Mann-Whitney U test:

- `Bg_Dwstrat` vs `P_Dw` : 2.44193e-05
  mean: 5476.1883 > 3246.4219 , median: 2000 > 1546.5
- `Bg_Dwstrat` vs `Bg_Upstrat` : 0.000809048
  mean: 5476.1883 > 4812.6347 , median: 2000 > 1662.5
- `Bg_Dwstrat` vs `P_Up` : 0.00323931
  mean: 5476.1883 > 4114.7365 , median: 2000 > 1234.5
- `Bg_Dwstrat` vs `LOW_PSI` : 2.19475e-16
  mean: 5476.1883 > 3619.9511 , median: 2000 > 1268.5
- `P_Dw` vs `HIGH_PSI` : 2.16457e-05
  mean: 3246.4219 < 5569.5115 , median: 1546.5 < 1988
- `P_Dw` vs `LOW_PSI` : 0.00325781
  mean: 3246.4219 < 3619.9511 , median: 1546.5 > 1268.5
- `Bg_Upstrat` vs `HIGH_PSI` : 0.000812048
  mean: 4812.6347 < 5569.5115 , median: 1662.5 < 1988
- `Bg_Upstrat` vs `LOW_PSI` : 0.00315919
  mean: 4812.6347 > 3619.9511 , median: 1662.5 > 1268.5
- `P_Up` vs `HIGH_PSI` : 0.00359788
  mean: 4114.7365 < 5569.5115 , median: 1234.5 < 1988
- `HIGH_PSI` vs `LOW_PSI` : 8.15661e-18
  mean: 5569.5115 > 3619.9511 , median: 1988 > 1268.5

#### 6.8 RATIO UPINTRON EXON LENGTH

Back to: Overview | ToC

Meaning: median up-stream intron length / exon length

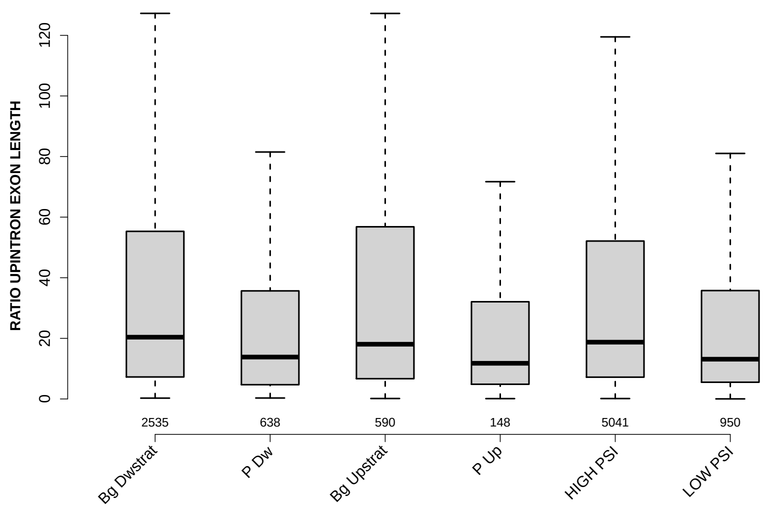

Significant results from Mann-Whitney U test:

- Bg_Dwstrat vs P_Dw : 1.07327e-08
  mean: 70.7248 > 37.8451 , median: 20.3814 > 13.8218
- Bg_Dwstrat vs P_Up : 0.000345709
  mean: 70.7248 > 40.503 , median: 20.3814 > 11.7679
- Bg_Dwstrat vs LOW_PSI : 2.31433e-09
  mean: 70.7248 < 139.1717 , median: 20.3814 > 13.1172
- P_Dw vs Bg_Upstrat : 0.000207575
  mean: 37.8451 < 69.5559 , median: 13.8218 < 18.0598
- P_Dw vs HIGH_PSI : 1.10212e-07
  mean: 37.8451 < 65.6412 , median: 13.8218 < 18.7391
- Bg_Upstrat vs P_Up : 0.00396605
  mean: 69.5559 > 40.503 , median: 18.0598 > 11.7679
- Bg_Upstrat vs LOW_PSI : 0.000414148
  mean: 69.5559 < 139.1717 , median: 18.0598 > 13.1172
- P_Up vs HIGH_PSI : 0.000954051
  mean: 40.503 < 65.6412 , median: 11.7679 < 18.7391
- HIGH_PSI vs LOW_PSI : 2.06874e-08
  mean: 65.6412 < 139.1717 , median: 18.7391 > 13.1172

#### 6.9 RATIO DOINTRON EXON LENGTH

Back to: Overview | ToC

Meaning: median down-stream intron length / exon length

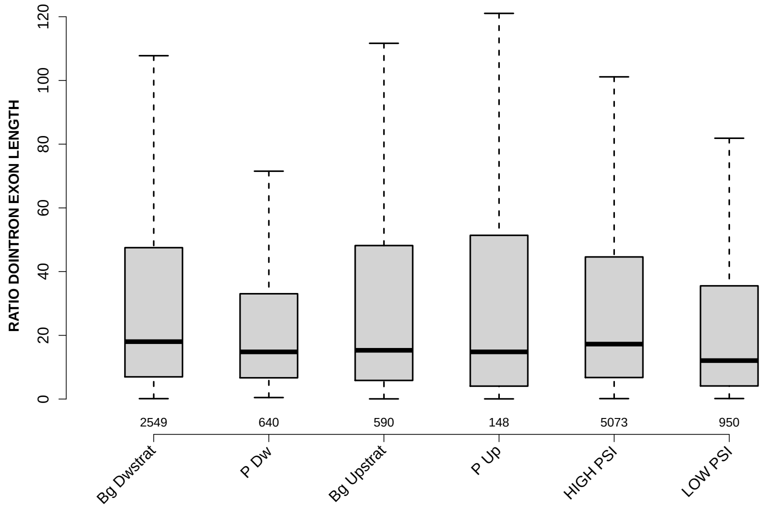

Significant results from Mann-Whitney U test:

- `Bg_Dwstrat` vs `P_Dw` : 0.000899113
  mean: 54.0274 > 31.9573 , median: 18.0185 > 14.8063
- `Bg_Dwstrat` vs `LOW_PSI` : 1.21068e-09
  mean: 54.0274 < 81.8413 , median: 18.0185 > 12.071
- `P_Dw` vs `HIGH_PSI` : 0.00304456
  mean: 31.9573 < 53.7682 , median: 14.8063 < 17.2475
- `P_Dw` vs `LOW_PSI` : 0.0246602
  mean: 31.9573 < 81.8413 , median: 14.8063 > 12.071
- `Bg_Upstrat` vs `LOW_PSI` : 0.00100828
  mean: 79.5031 < 81.8413 , median: 15.3022 > 12.071
- `HIGH_PSI` vs `LOW_PSI` : 1.47445e-09
  mean: 53.7682 < 81.8413 , median: 17.2475 > 12.071

#### 6.10 EXON GCC

Back to: Overview | ToC

Meaning: GC content of entire exon sequence

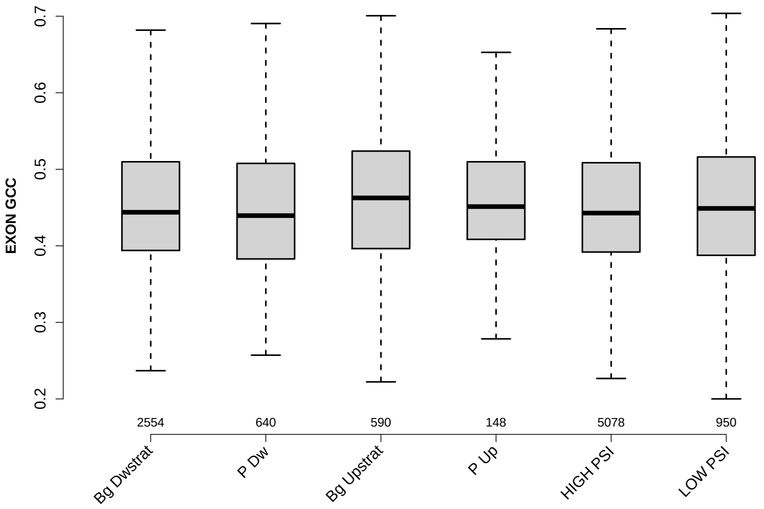

Significant results from Mann-Whitney U test:

- `Bg_Dwstrat` vs `Bg_Upstrat` : 0.0371666
  mean: 0.454684 < 0.460744 , median: 0.443835 < 0.462574
- `P_Dw` vs `Bg_Upstrat` : 0.0148211
  mean: 0.451739 < 0.460744 , median: 0.439477 < 0.462574
- `Bg_Upstrat` vs `HIGH_PSI` : 0.0171208
  mean: 0.460744 > 0.454001 , median: 0.462574 > 0.442803
- `Bg_Upstrat` vs `LOW_PSI` : 0.0435904
  mean: 0.460744 > 0.449449 , median: 0.462574 > 0.44888

#### 6.11 UPINTRON GCC

Back to: Overview | ToC

Meaning: GC content of entire up-stream intron sequence

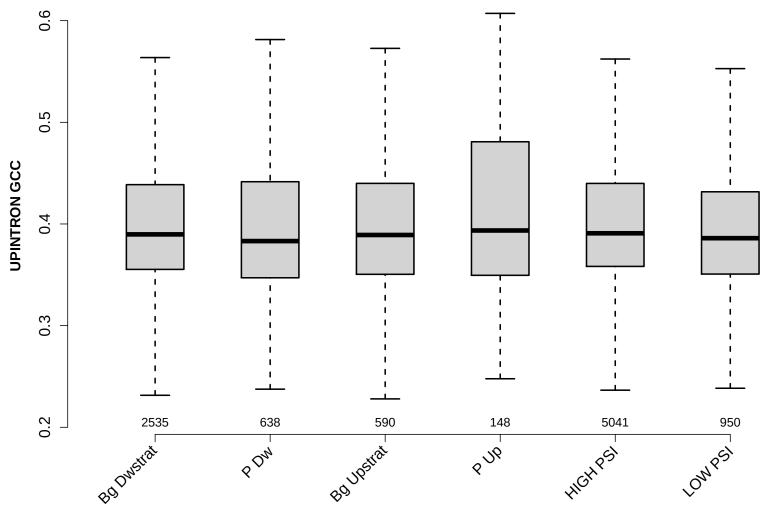

Significant results from Mann-Whitney U test:

- P_Dw vs HIGH_PSI : 0.0134029
  mean: 0.40391 < 0.408145 , median: 0.383158 < 0.390841
- HIGH_PSI vs LOW_PSI : 0.0133334
  mean: 0.408145 > 0.40223 , median: 0.390841 > 0.386017

#### 6.12 UPEXON GCC

Back to: Overview | ToC

Meaning: GC content of entire up-stream exon sequence

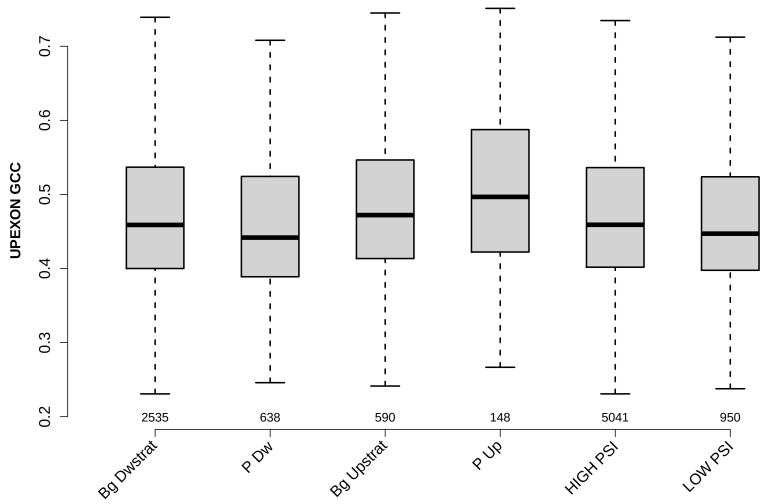

Significant results from Mann-Whitney U test:

- Bg_Dwstrat vs P_Dw : 0.000336181
  mean: 0.475889 > 0.460018 , median: 0.458824 > 0.441764
- Bg_Dwstrat vs Bg_Upstrat : 0.00640045
  mean: 0.475889 < 0.489178 , median: 0.458824 < 0.472111
- Bg_Dwstrat vs P_Up : 0.000692306
  mean: 0.475889 < 0.504396 , median: 0.458824 < 0.496667
- Bg_Dwstrat vs LOW_PSI : 0.0404072
  mean: 0.475889 > 0.46833 , median: 0.458824 > 0.447059
- P_Dw vs Bg_Upstrat : 6.54167e-07
  mean: 0.460018 < 0.489178 , median: 0.441764 < 0.472111
- P_Dw vs P_Up : 1.50362e-06
  mean: 0.460018 < 0.504396 , median: 0.441764 < 0.496667
- P_Dw vs HIGH_PSI : 0.000151456
  mean: 0.460018 < 0.474982 , median: 0.441764 < 0.459016
- Bg_Upstrat vs HIGH_PSI : 0.00375196
  mean: 0.489178 > 0.474982 , median: 0.472111 > 0.459016
- Bg_Upstrat vs LOW_PSI : 0.000101732
  mean: 0.489178 > 0.46833 , median: 0.472111 > 0.447059
- P_Up vs HIGH_PSI : 0.000465384
  mean: 0.504396 > 0.474982 , median: 0.496667 > 0.459016

- P_Up vs LOW_PSI : 5.0156e-05
  mean: 0.504396 > 0.46833 , median: 0.496667 > 0.447059
- HIGH_PSI vs LOW_PSI : 0.0287194
  mean: 0.474982 > 0.46833 , median: 0.459016 > 0.447059

#### 6.13 DOINTRON GCC

Back to: Overview | ToC

Meaning: GC content of entire down-stream intron sequence

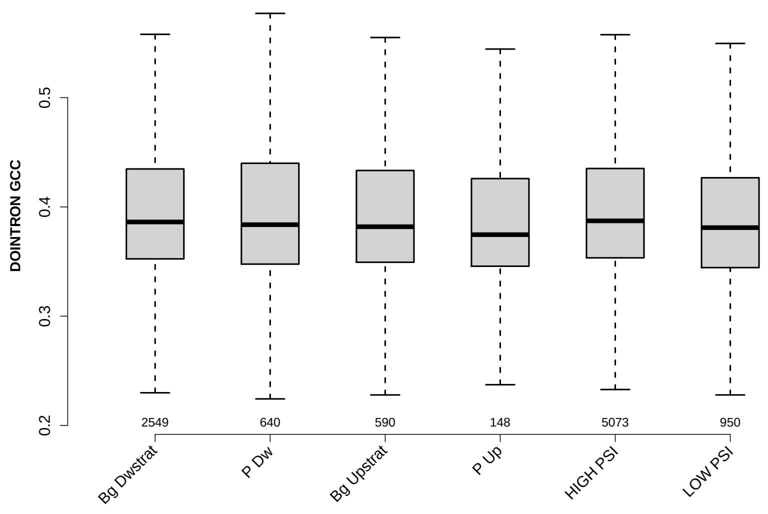

Significant results from Mann-Whitney U test:

- `Bg_Dwstrat` vs `P_Up` : 0.0461719
  mean: 0.40243 > 0.393467 , median: 0.386207 > 0.374597
- `Bg_Dwstrat` vs `LOW_PSI` : 0.00325909
  mean: 0.40243 > 0.392939 , median: 0.386207 > 0.381036
- `P_Up` vs `HIGH_PSI` : 0.0285159
  mean: 0.393467 < 0.403579 , median: 0.374597 < 0.387224
- `HIGH_PSI` vs `LOW_PSI` : 0.000367439
  mean: 0.403579 > 0.392939 , median: 0.387224 > 0.381036

#### 6.14 DOEXON GCC

Back to: Overview | ToC

Meaning: GC content of entire down-stream exon sequence

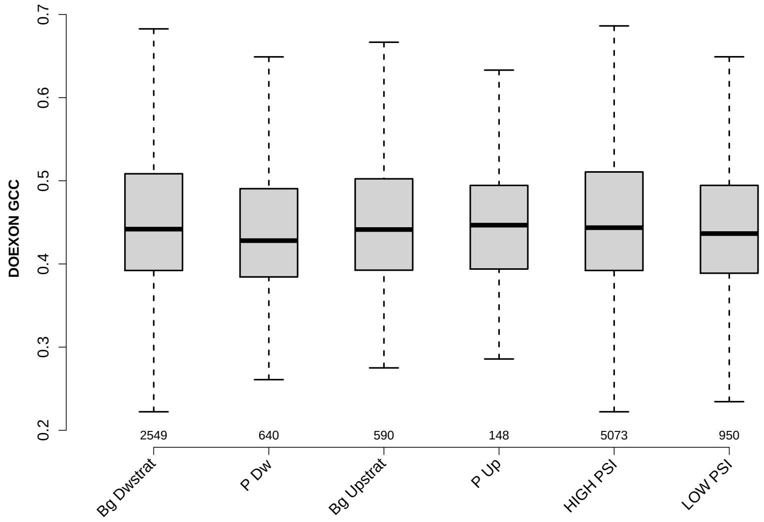

Significant results from Mann-Whitney U test:

- `Bg_Dwstrat` vs `P_Dw` : 0.00195889
  mean: 0.452686 > 0.442211 , median: 0.44186 > 0.428026
- `P_Dw` vs `Bg_Upstrat` : 0.0130229
  mean: 0.442211 < 0.452997 , median: 0.428026 < 0.4414
- `P_Dw` vs `HIGH_PSI` : 0.000269361
  mean: 0.442211 < 0.454135 , median: 0.428026 < 0.443609
- `HIGH_PSI` vs `LOW_PSI` : 0.0451769
  mean: 0.454135 > 0.447948 , median: 0.443609 > 0.436528

#### 6.15 RATIO UPEXON EXON GCC

Back to: Overview | ToC

Meaning: UPEXON GCC / EXON GCC

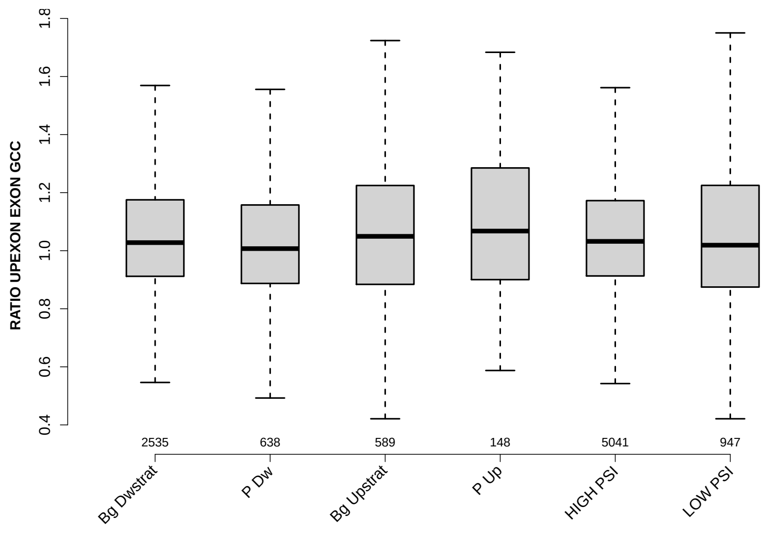

Significant results from Mann-Whitney U test:

- Bg_Dwstrat vs P_Dw : 0.0207975
  mean: 1.0671 > 1.0424 , median: 1.0278 > 1.0071
- Bg_Dwstrat vs P_Up : 0.0466445
  mean: 1.0671 < 1.1343 , median: 1.0278 < 1.0677
- P_Dw vs Bg_Upstrat : 0.0203087
  mean: 1.0424 < 1.0929 , median: 1.0071 < 1.0494
- P_Dw vs P_Up : 0.00439257
  mean: 1.0424 < 1.1343 , median: 1.0071 < 1.0677
- P_Dw vs HIGH_PSI : 0.00990314
  mean: 1.0424 < 1.0667 , median: 1.0071 < 1.0321
- P_Up vs LOW_PSI : 0.0346838
  mean: 1.1343 > 1.0727 , median: 1.0677 > 1.019

#### 6.16 RATIO UPINTRON EXON GCC

Back to: Overview | ToC

Meaning: UPINTRON GCC / EXON GCC

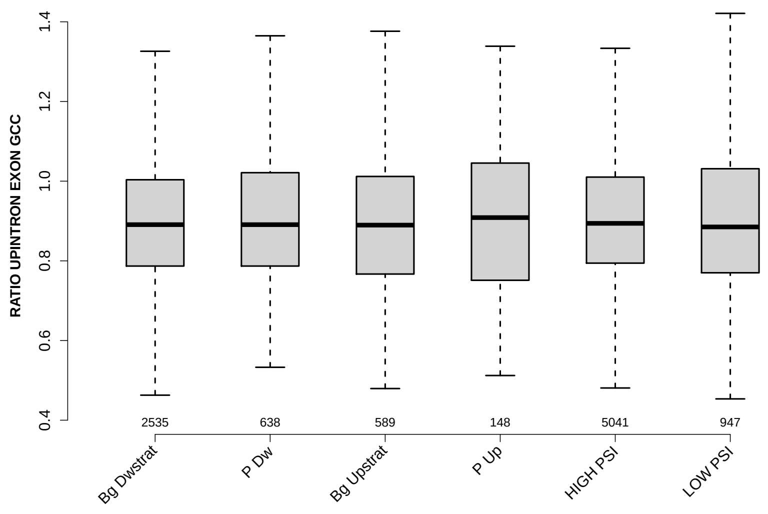

Significant results from Mann-Whitney U test:

- none

#### 6.17 RATIO DOINTRON EXON GCC

Back to: Overview | ToC

Meaning: DOINTRON GCC / EXON GCC

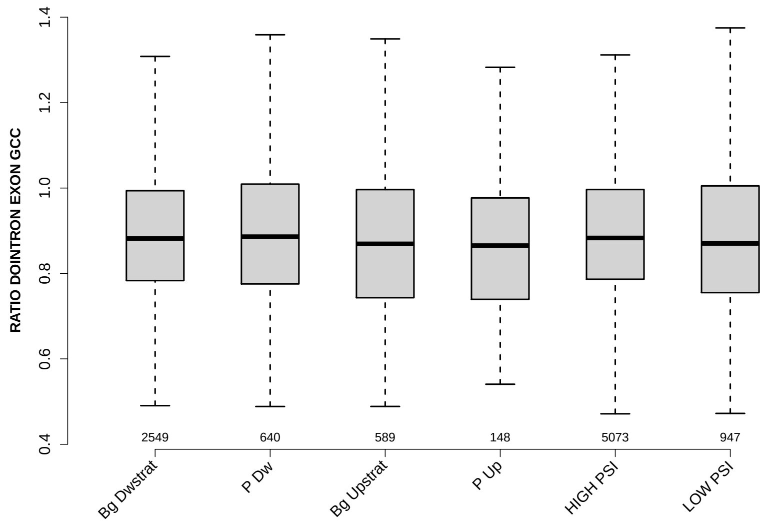

Significant results from Mann-Whitney U test:

- P_Dw vs Bg_Upstrat : 0.0332519
  mean: 0.90812 > 0.888145 , median: 0.886095 > 0.869301
- P_Dw vs P_Up : 0.0463563
  mean: 0.90812 > 0.873287 , median: 0.886095 > 0.865197
- Bg_Upstrat vs HIGH_PSI : 0.0188597
  mean: 0.888145 < 0.902299 , median: 0.869301 < 0.88311
- P_Up vs HIGH_PSI : 0.0489958
  mean: 0.873287 < 0.902299 , median: 0.865197 < 0.88311
- HIGH_PSI vs LOW_PSI : 0.026562
  mean: 0.902299 > 0.89282 , median: 0.88311 > 0.870407

#### 6.18 RATIO DOEXON EXON GCC

Back to: Overview | ToC

Meaning: DOEXON GCC / EXON GCC

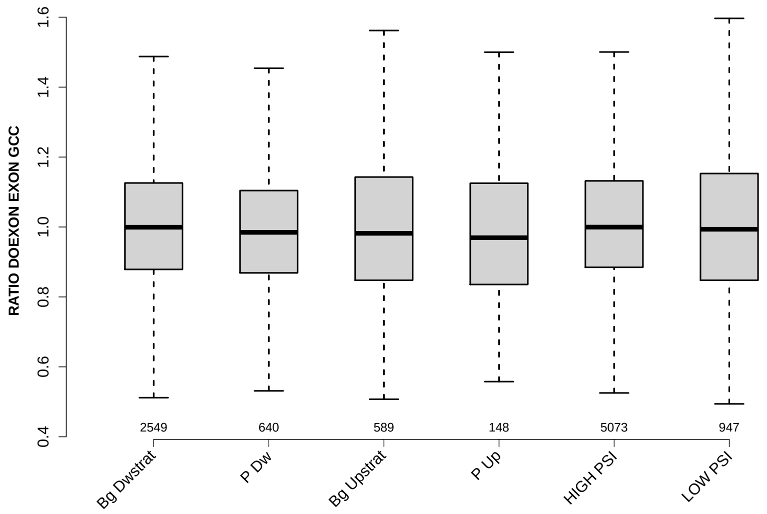

Significant results from Mann-Whitney U test:

- P_Dw vs HIGH_PSI : 0.0229069
  mean: 1.0017 < 1.0189 , median: 0.984591 < 0.999621

#### 6.19 SF1 HIGHESTSCORE 3SS UPINTRON

Back to: Overview | ToC

Meaning: highest score of a SF1 positon weight matrix trained with human data in the last 150 nt 3 prime intron positons of up-stream intron

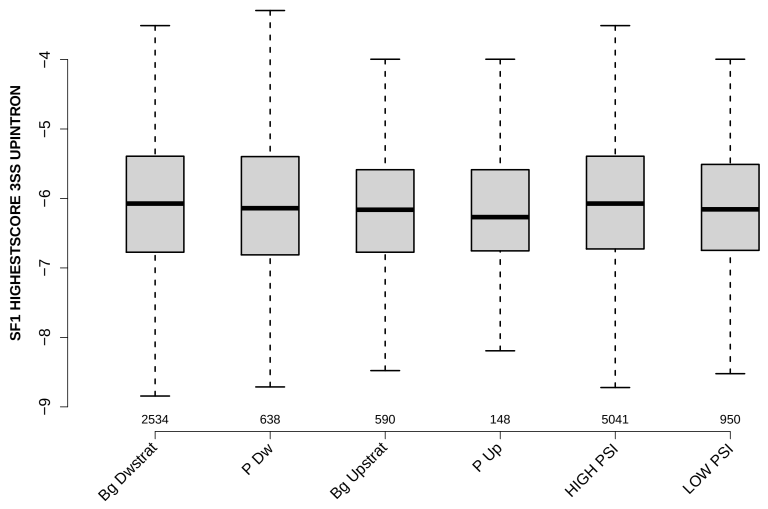

Significant results from Mann-Whitney U test:

- none

#### 6.20 SF1 HIGHESTSCORE 3SS DOINTRON

Back to: Overview | ToC

Meaning: highest score of a SF1 positon weight matrix trained with human data in the last 150 nt 3 prime intron positons of down-stream intron

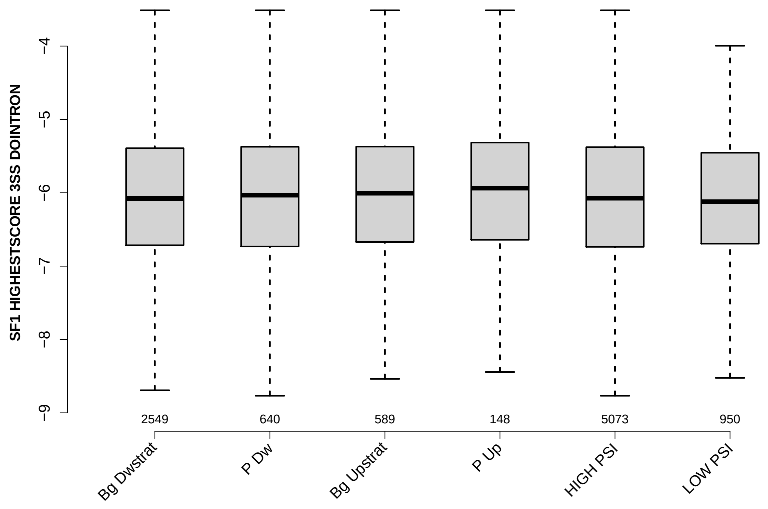

Significant results from Mann-Whitney U test:

- none

#### 6.21 UP 5SS 20INT10EX GCC

Back to: Overview | ToC

Meaning: GC content of up-stream 5ss sequence (20int+10ex positions)

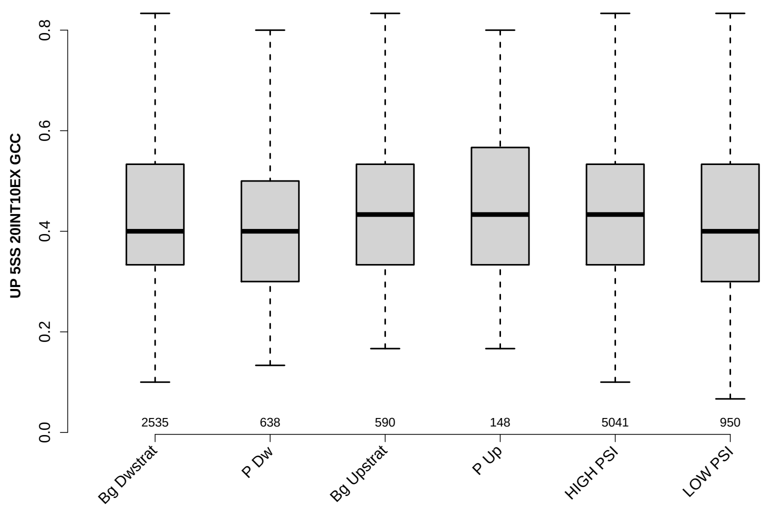

Significant results from Mann-Whitney U test:

- `Bg_Dwstrat` vs `P_Dw` : 0.00165872
  mean: 0.437035 > 0.418056 , median: 0.4 = 0.4
- `Bg_Dwstrat` vs `Bg_Upstrat` : 0.0147213
  mean: 0.437035 < 0.452938 , median: 0.4 < 0.433333
- `Bg_Dwstrat` vs `LOW_PSI` : 0.0234828
  mean: 0.437035 > 0.425789 , median: 0.4 = 0.4
- `P_Dw` vs `Bg_Upstrat` : 1.21228e-05
  mean: 0.418056 < 0.452938 , median: 0.4 < 0.433333
- `P_Dw` vs `P_Up` : 0.0218448
  mean: 0.418056 < 0.448761 , median: 0.4 < 0.433333
- `P_Dw` vs `HIGH_PSI` : 0.000648664
  mean: 0.418056 < 0.437274 , median: 0.4 < 0.433333
- `Bg_Upstrat` vs `HIGH_PSI` : 0.0145008
  mean: 0.452938 > 0.437274 , median: 0.433333 = 0.433333
- `Bg_Upstrat` vs `LOW_PSI` : 0.000146469
  mean: 0.452938 > 0.425789 , median: 0.433333 > 0.4
- `HIGH_PSI` vs `LOW_PSI` : 0.010535
  mean: 0.437274 > 0.425789 , median: 0.433333 > 0.4

#### 6.22 GCC 3SS 20INT10EX

Back to: Overview | ToC

Meaning: GC content of 3ss sequence (20int+10ex positions)

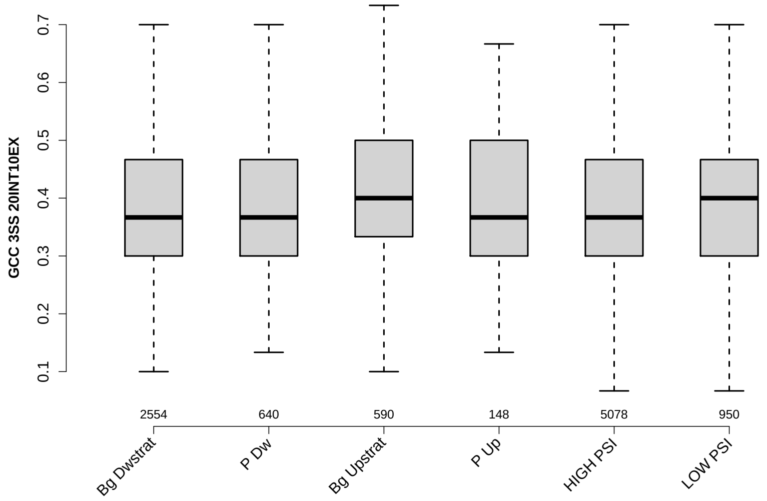

Significant results from Mann-Whitney U test:

- `Bg_Dwstrat` vs `Bg_Upstrat` : 9.50683e-05
  mean: 0.389702 < 0.406271 , median: 0.366667 < 0.4
- `Bg_Dwstrat` vs `LOW_PSI` : 0.0101804
  mean: 0.389702 < 0.397228 , median: 0.366667 < 0.4
- `P_Dw` vs `Bg_Upstrat` : 0.0056037
  mean: 0.392604 < 0.406271 , median: 0.366667 < 0.4
- `Bg_Upstrat` vs `HIGH_PSI` : 5.43674e-06
  mean: 0.406271 > 0.387679 , median: 0.4 > 0.366667
- `HIGH_PSI` vs `LOW_PSI` : 0.000892212
  mean: 0.387679 < 0.397228 , median: 0.366667 < 0.4

#### 6.23 GCC 5SS 20INT10EX

Back to: Overview | ToC

Meaning: GC content of 5ss sequence (20int+10ex positions)

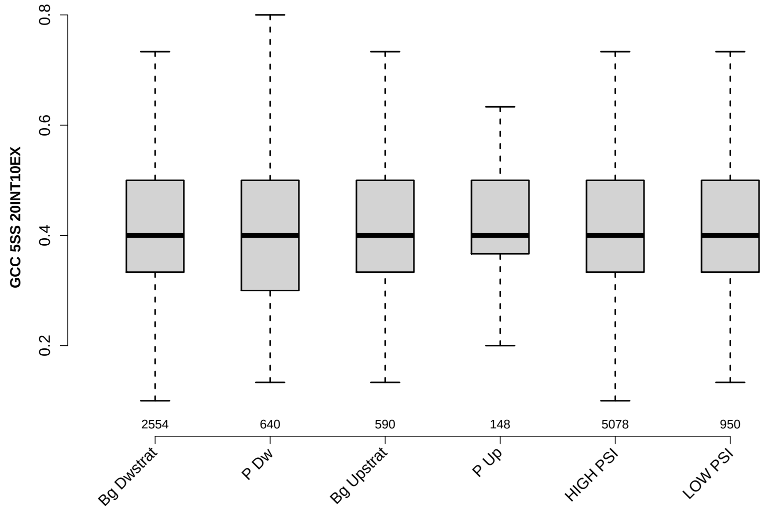

Significant results from Mann-Whitney U test:

- `Bg_Dwstrat` vs `Bg_Upstrat` : 0.0095765
  mean: 0.412555 < 0.421977 , median: 0.4 = 0.4
- `Bg_Upstrat` vs `HIGH_PSI` : 0.0215096
  mean: 0.421977 > 0.414999 , median: 0.4 = 0.4

#### 6.24 DO 3SS 20INT10EX GCC

Back to: Overview | ToC

Meaning: GC content of down-stream 3ss sequence (20int+10ex positions)

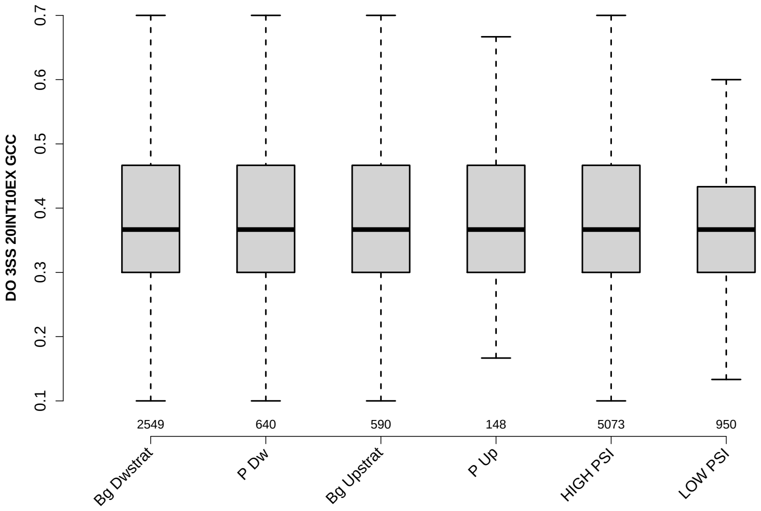

Significant results from Mann-Whitney U test:

- none

#### 6.25 MAXENTSCR HSAMODEL UPSTRM 5SS

Back to: Overview | ToC

Meaning: maximum entropy score of 5ss of up-stream exon using a model trained with human splice sites

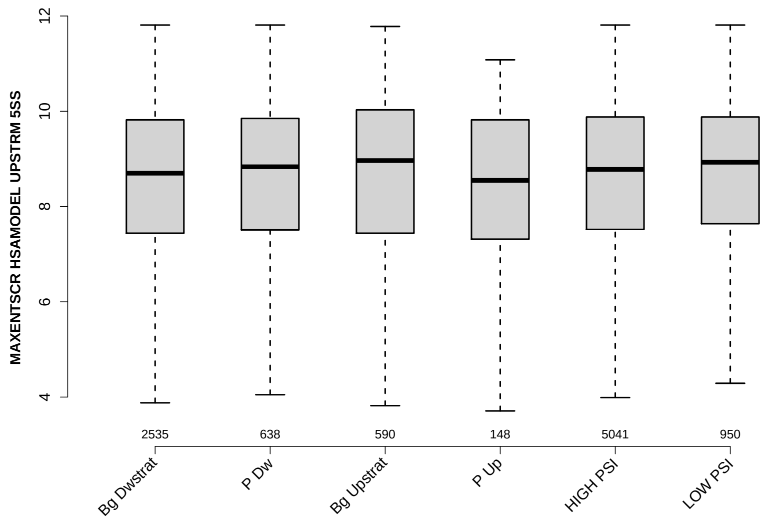

Significant results from Mann-Whitney U test:

- `Bg_Dwstrat` vs `Bg_Upstrat` : 0.0411867
  mean: 8.1136 < 8.1611 , median: 8.7 < 8.965
- `Bg_Dwstrat` vs `LOW_PSI` : 0.00765417
  mean: 8.1136 < 8.4687 , median: 8.7 < 8.93

#### 6.26 MAXENTSCR HSAMODEL 3SS

Back to: Overview | ToC

Meaning: maximum entropy score of 3ss using a model trained with human splice sites

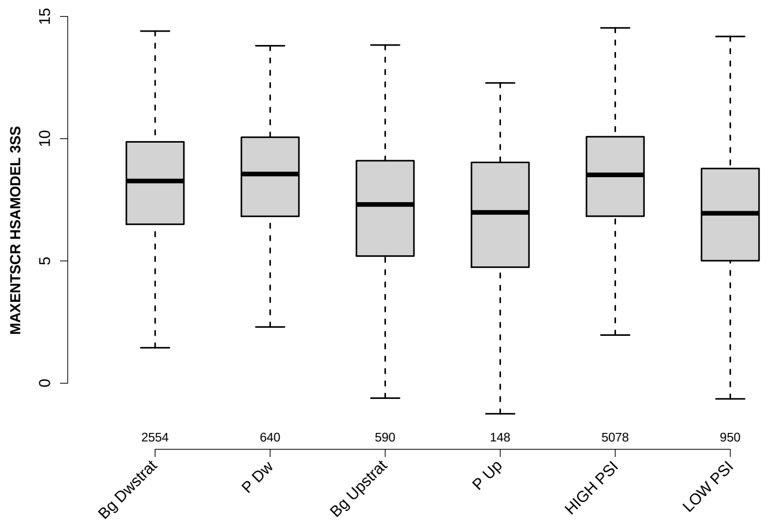

Significant results from Mann-Whitney U test:

- Bg_Dwstrat vs P_Dw : 0.00803932
  mean: 7.8393 < 8.3662 , median: 8.27 < 8.555
- Bg_Dwstrat vs Bg_Upstrat : 2.21637e-14
  mean: 7.8393 > 6.7478 , median: 8.27 > 7.31
- Bg_Dwstrat vs P_Up : 4.52768e-07
  mean: 7.8393 > 6.6478 , median: 8.27 > 6.985
- Bg_Dwstrat vs HIGH_PSI : 1.81573e-05
  mean: 7.8393 < 8.3058 , median: 8.27 < 8.52
- Bg_Dwstrat vs LOW_PSI : 1.30309e-32
  mean: 7.8393 > 6.5217 , median: 8.27 > 6.95
- P_Dw vs Bg_Upstrat : 1.28059e-16
  mean: 8.3662 > 6.7478 , median: 8.555 > 7.31
- P_Dw vs P_Up : 1.31479e-09
  mean: 8.3662 > 6.6478 , median: 8.555 > 6.985
- P_Dw vs LOW_PSI : 1.04432e-29
  mean: 8.3662 > 6.5217 , median: 8.555 > 6.95
- Bg_Upstrat vs HIGH_PSI : 1.50677e-25
  mean: 6.7478 < 8.3058 , median: 7.31 < 8.52
- P_Up vs HIGH_PSI : 1.76056e-10
  mean: 6.6478 < 8.3058 , median: 6.985 < 8.52

- `HIGH_PSI` vs `LOW_PSI` : 2.02947e-56
  mean: 8.3058 > 6.5217 , median: 8.52 > 6.95

#### 6.27 MAXENTSCR HSAMODEL 5SS

Back to: Overview | ToC

Meaning: maximum entropy score of 5ss using a model trained with human splice sites

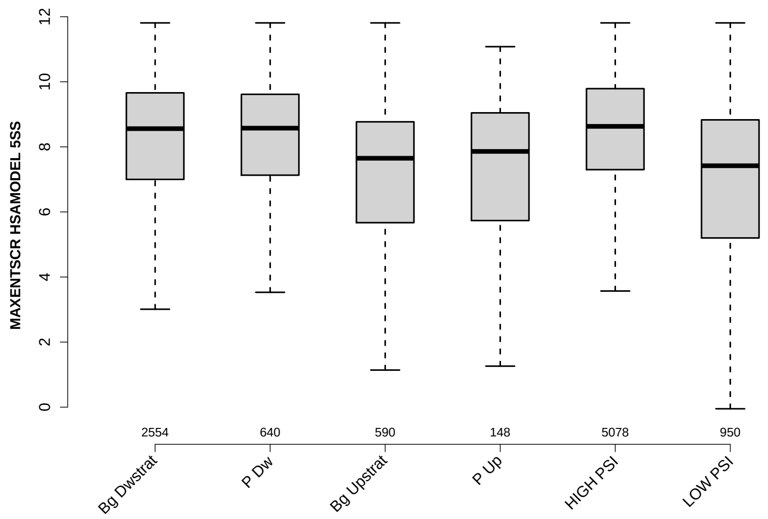

Significant results from Mann-Whitney U test:

- `Bg_Dwstrat` vs `Bg_Upstrat` : 2.42146e-20
  mean: 7.8832 > 6.7779 , median: 8.56 > 7.65
- `Bg_Dwstrat` vs `P_Up` : 5.37872e-05
  mean: 7.8832 > 6.9855 , median: 8.56 > 7.86
- `Bg_Dwstrat` vs `HIGH_PSI` : 0.00322219
  mean: 7.8832 < 8.2026 , median: 8.56 < 8.63
- `Bg_Dwstrat` vs `LOW_PSI` : 3.08431e-36
  mean: 7.8832 > 6.6397 , median: 8.56 > 7.42
- `P_Dw` vs `Bg_Upstrat` : 1.40832e-14
  mean: 8.0527 > 6.7779 , median: 8.575 > 7.65
- `P_Dw` vs `P_Up` : 9.39846e-05
  mean: 8.0527 > 6.9855 , median: 8.575 > 7.86
- `P_Dw` vs `LOW_PSI` : 4.52859e-22
  mean: 8.0527 > 6.6397 , median: 8.575 > 7.42
- `Bg_Upstrat` vs `HIGH_PSI` : 8.28014e-30
  mean: 6.7779 < 8.2026 , median: 7.65 < 8.63
- `P_Up` vs `HIGH_PSI` : 8.18013e-07
  mean: 6.9855 < 8.2026 , median: 7.86 < 8.63
- `HIGH_PSI` vs `LOW_PSI` : 4.10684e-54
  mean: 8.2026 > 6.6397 , median: 8.63 > 7.42

#### 6.28 MAXENTSCR HSAMODEL DOWNSTRM 3SS

Back to: Overview | ToC

Meaning: maximum entropy score of 3ss of down-stream exon using a model trained with human splice sites

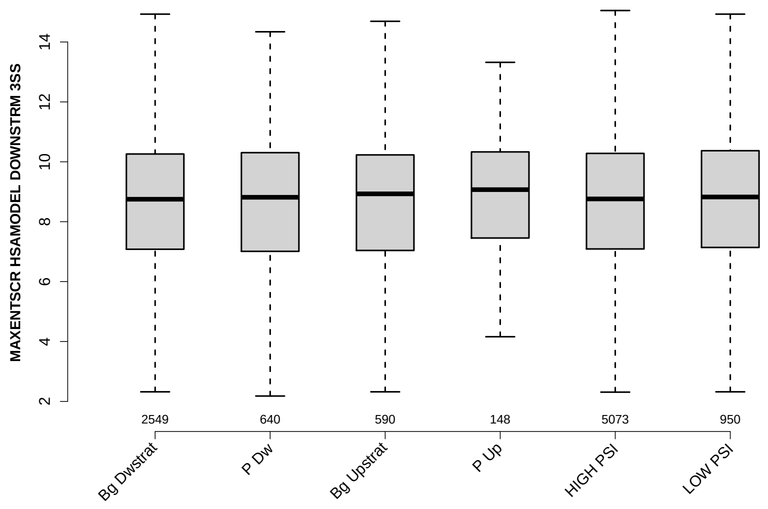

Significant results from Mann-Whitney U test:

- none

#### 6.29 DIST FROM MAXBP TO 3SS UPINTRON

Back to: Overview | ToC

Meaning: distance to 3ss of best precited BP

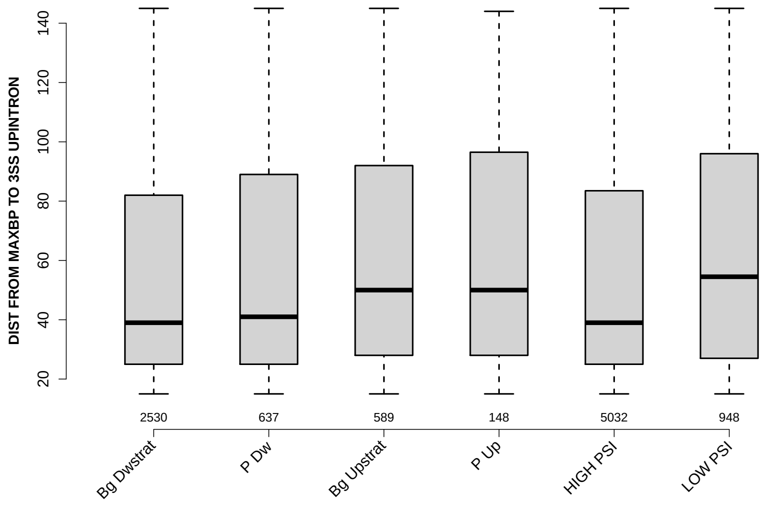

Significant results from Mann-Whitney U test:

- Bg_Dwstrat vs Bg_Upstrat : 2.57432e-05
  mean: 55.5656 < 62.3107 , median: 39 < 50
- Bg_Dwstrat vs P_Up : 0.0331555
  mean: 55.5656 < 62.5338 , median: 39 < 50
- Bg_Dwstrat vs LOW_PSI : 9.70276e-09
  mean: 55.5656 < 63.5401 , median: 39 < 54.5
- P_Dw vs Bg_Upstrat : 0.0232593
  mean: 58.314 < 62.3107 , median: 41 < 50
- P_Dw vs LOW_PSI : 0.00253991
  mean: 58.314 < 63.5401 , median: 41 < 54.5
- Bg_Upstrat vs HIGH_PSI : 2.1642e-05
  mean: 62.3107 > 55.997 , median: 50 > 39
- P_Up vs HIGH_PSI : 0.0381726
  mean: 62.5338 > 55.997 , median: 50 > 39
- HIGH_PSI vs LOW_PSI : 3.07079e-09
  mean: 55.997 < 63.5401 , median: 39 < 54.5

#### 6.30 SCORE FOR MAXBP SEQ UPINTRON

Back to: Overview | ToC

Meaning: BP sequence score of best predicted BP

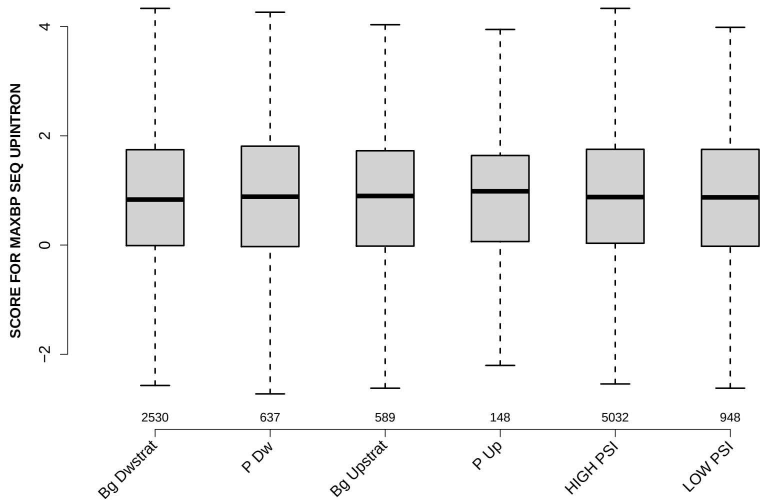

Significant results from Mann-Whitney U test:

- none

#### 6.31 PYRIMIDINECONT MAXBP UPINTRON

Back to: Overview | ToC

Meaning: Pyrimidine content between the BP adenine and the 3 prime splice site for best BP

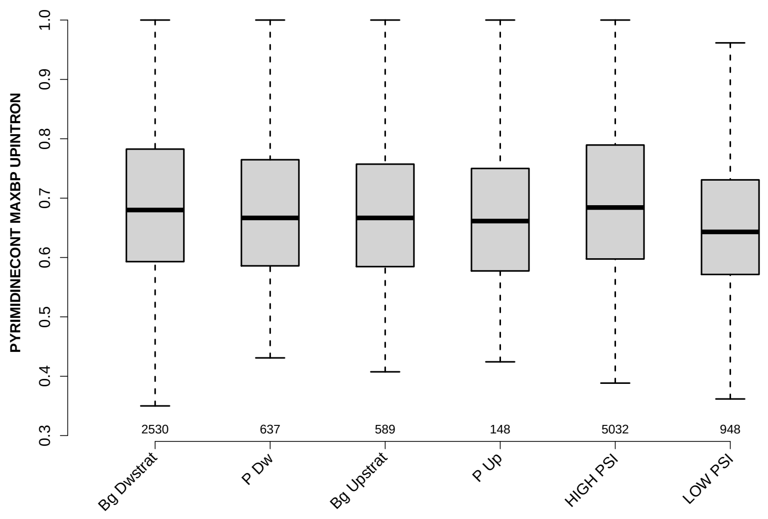

Significant results from Mann-Whitney U test:

- Bg_Dwstrat vs Bg_Upstrat : 0.0324241
  mean: 0.689039 > 0.676106 , median: 0.68 > 0.666667
- Bg_Dwstrat vs LOW_PSI : 2.46828e-11
  mean: 0.689039 > 0.657024 , median: 0.68 > 0.643134
- P_Dw vs HIGH_PSI : 0.00404084
  mean: 0.679227 < 0.694354 , median: 0.666667 < 0.684211
- P_Dw vs LOW_PSI : 0.000928699
  mean: 0.679227 > 0.657024 , median: 0.666667 > 0.643134
- Bg_Upstrat vs HIGH_PSI : 0.0015489
  mean: 0.676106 < 0.694354 , median: 0.666667 < 0.684211
- Bg_Upstrat vs LOW_PSI : 0.00222503
  mean: 0.676106 > 0.657024 , median: 0.666667 > 0.643134
- P_Up vs HIGH_PSI : 0.0180473
  mean: 0.669583 < 0.694354 , median: 0.661391 < 0.684211
- HIGH_PSI vs LOW_PSI : 6.92471e-17
  mean: 0.694354 > 0.657024 , median: 0.684211 > 0.643134

#### 6.32 POLYPYRITRAC OFFSET MAXBP UPINTRON

Back to: Overview | ToC

Meaning: Polypyrimidine track offset relative to the BP adenine for best BP

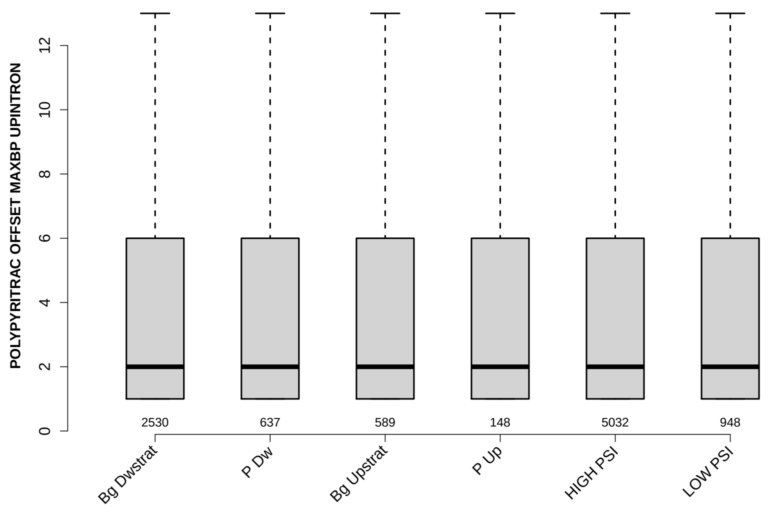

Significant results from Mann-Whitney U test:

- Bg_Dwstrat vs P_Dw : 0.0251185
  mean: 4.0941 < 4.7049 , median: 2 = 2
- P_Dw vs HIGH_PSI : 0.0200271
  mean: 4.7049 > 4.0803 , median: 2 = 2

#### 6.33 POLYPYRITRAC LEN MAXBP UPINTRON

Back to: Overview | ToC

Meaning: Polypyrimidine track length for best BP

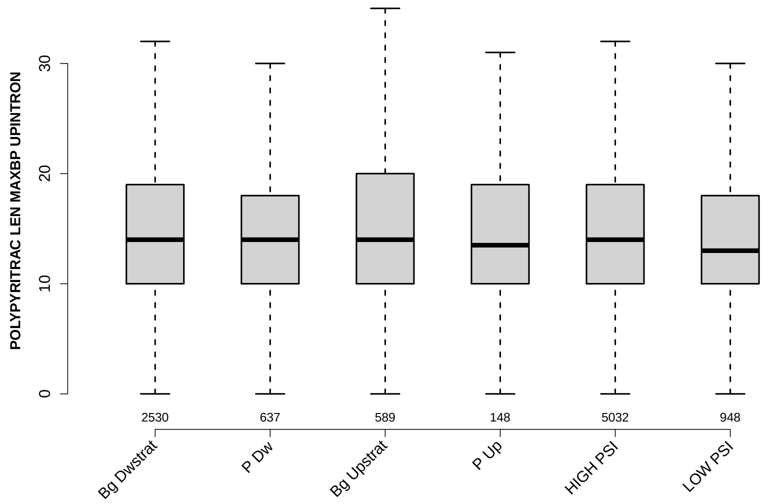

Significant results from Mann-Whitney U test:

- `Bg_Dwstrat` vs `LOW_PSI` : 0.0272079
  mean: 15.3976 > 15.1477 , median: 14 > 13
- `Bg_Upstrat` vs `LOW_PSI` : 0.00806701
  mean: 16.6027 > 15.1477 , median: 14 > 13
- `HIGH_PSI` vs `LOW_PSI` : 0.0194969
  mean: 15.4614 > 15.1477 , median: 14 > 13

#### 6.34 POLYPYRITRAC SCORE MAXBP UPINTRON

Back to: Overview | ToC

Meaning: Polypyrimidine track score for best BP

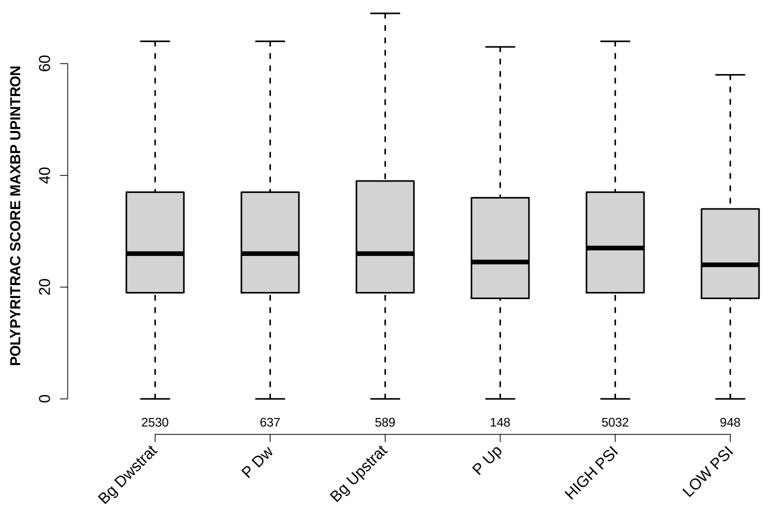

Significant results from Mann-Whitney U test:

- `Bg_Dwstrat` vs `LOW_PSI` : 0.000167989
  mean: 30.0573 > 28.827 , median: 26 > 24
- `P_Dw` vs `LOW_PSI` : 0.00502874
  mean: 30.2747 > 28.827 , median: 26 > 24
- `Bg_Upstrat` vs `LOW_PSI` : 0.00511046
  mean: 32.0323 > 28.827 , median: 26 > 24
- `HIGH_PSI` vs `LOW_PSI` : 1.3514e-05
  mean: 30.4086 > 28.827 , median: 27 > 24

#### 6.35 BPSCORE MAXBP UPINTRON

Back to: Overview | ToC

Meaning: SVM classification score of best BP

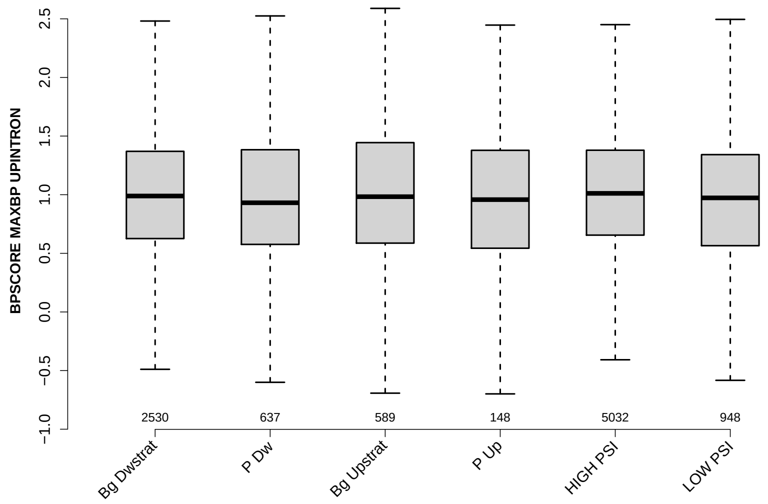

Significant results from Mann-Whitney U test:

- `P_Dw` vs `HIGH_PSI` : 0.0209274
  mean: 0.950837 < 1.0093 , median: 0.931285 < 1.0114
- `HIGH_PSI` vs `LOW_PSI` : 0.00708267
  mean: 1.0093 > 0.943046 , median: 1.0114 > 0.972861

#### 6.36 NUM PREDICTED BPS UPINTRON

Back to: Overview | ToC

Meaning: number of all predicted BPs which have a positive BP score

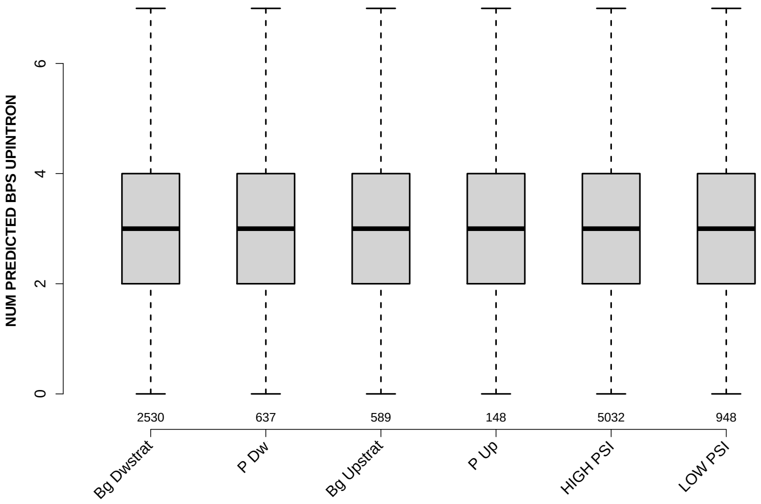

Significant results from Mann-Whitney U test:

- Bg_Dwstrat vs P_Dw : 0.00512798
  mean: 3.2549 > 3.0283 , median: 3 = 3
- Bg_Dwstrat vs LOW_PSI : 0.04085
  mean: 3.2549 > 3.116 , median: 3 = 3
- P_Dw vs HIGH_PSI : 0.000156047
  mean: 3.0283 < 3.3144 , median: 3 = 3
- HIGH_PSI vs LOW_PSI : 0.00151383
  mean: 3.3144 > 3.116 , median: 3 = 3

#### 6.37 MEDIAN DIST FROM BP TO 3SS UPINTRON

Back to: Overview | ToC

Meaning: like DIST FROM MAXBP TO 3SS but median over top-3 predicted BPs

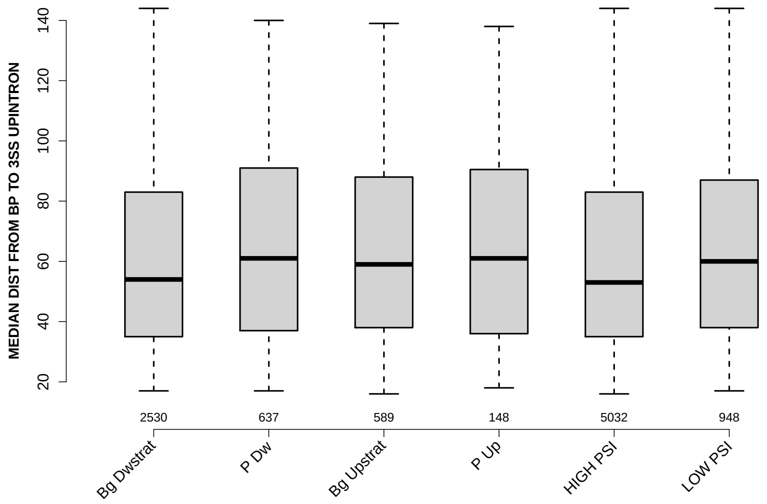

Significant results from Mann-Whitney U test:

- `Bg_Dwstrat` vs `P_Dw` : 0.00496023
  mean: 61.3996 < 65.4254 , median: 54 < 61
- `Bg_Dwstrat` vs `Bg_Upstrat` : 0.0181536
  mean: 61.3996 < 64.1961 , median: 54 < 59
- `Bg_Dwstrat` vs `LOW_PSI` : 0.00347409
  mean: 61.3996 < 64.2104 , median: 54 < 60
- `P_Dw` vs `HIGH_PSI` : 0.00204241
  mean: 65.4254 > 61.0949 , median: 61 > 53
- `Bg_Upstrat` vs `HIGH_PSI` : 0.00846463
  mean: 64.1961 > 61.0949 , median: 59 > 53
- `HIGH_PSI` vs `LOW_PSI` : 0.00089913
  mean: 61.0949 < 64.2104 , median: 53 < 60

#### 6.38 MEDIAN SCORE FOR BPSEQ UPINTRON

Back to: Overview | ToC

Meaning: like SCORE FOR MAXBP SEQ but median over top-3 predicted BPs

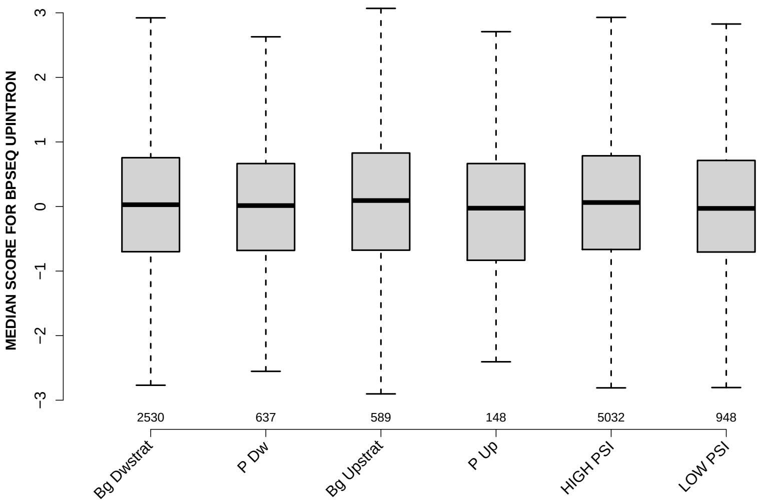

Significant results from Mann-Whitney U test:

- none

#### 6.39 MEDIAN PYRIMIDINECONT UPINTRON

Back to: Overview | ToC

Meaning: like PYRIMIDINECONT MAXBP but median over top-3 predicted BPs

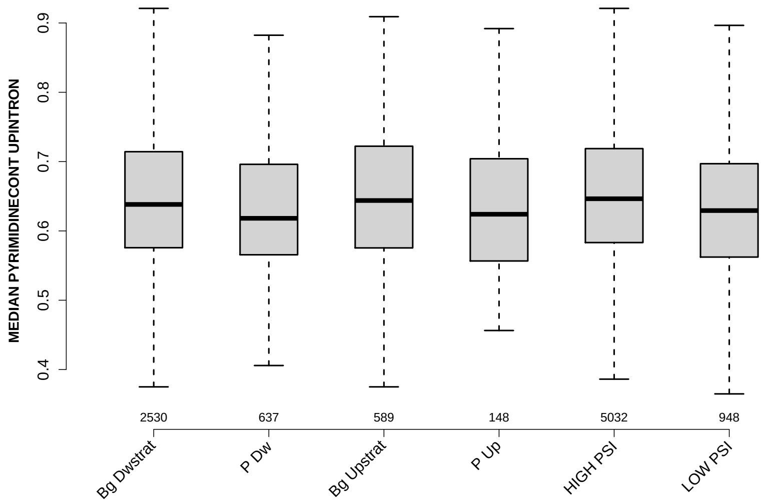

Significant results from Mann-Whitney U test:

- Bg_Dwstrat vs P_Dw : 0.000573053
  mean: 0.646592 > 0.633613 , median: 0.638197 > 0.618182
- Bg_Dwstrat vs HIGH_PSI : 0.00841354
  mean: 0.646592 < 0.65279 , median: 0.638197 < 0.646403
- Bg_Dwstrat vs LOW_PSI : 0.000746273
  mean: 0.646592 > 0.633121 , median: 0.638197 > 0.629331
- P_Dw vs Bg_Upstrat : 0.00270699
  mean: 0.633613 < 0.648496 , median: 0.618182 < 0.643836
- P_Dw vs HIGH_PSI : 1.69455e-07
  mean: 0.633613 < 0.65279 , median: 0.618182 < 0.646403
- Bg_Upstrat vs LOW_PSI : 0.00384368
  mean: 0.648496 > 0.633121 , median: 0.643836 > 0.629331
- P_Up vs HIGH_PSI : 0.0407719
  mean: 0.639053 < 0.65279 , median: 0.624038 < 0.646403
- HIGH_PSI vs LOW_PSI : 5.09315e-08
  mean: 0.65279 > 0.633121 , median: 0.646403 > 0.629331

#### 6.40 MEDIAN POLYPYRITRAC OFFSET UPINTRON

Back to: Overview | ToC

Meaning: like POLYPYRITRAC OFFSET MAXBP but median over top-3 predicted BPs

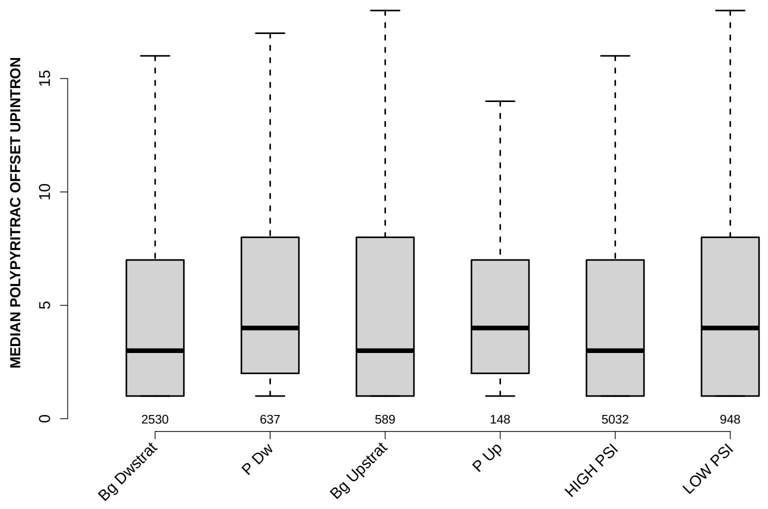

Significant results from Mann-Whitney U test:

- `Bg_Dwstrat` vs `P_Dw` : 0.0267496
  mean: 5.9581 < 6.7308 , median: 3 < 4
- `P_Dw` vs `HIGH_PSI` : 0.00136726
  mean: 6.7308 > 5.648 , median: 4 > 3

#### 6.41 MEDIAN POLYPYRITRAC LEN UPINTRON

Back to: Overview | ToC

Meaning: like POLYPYRITRAC LEN MAXBP but median over top-3 predicted BPs

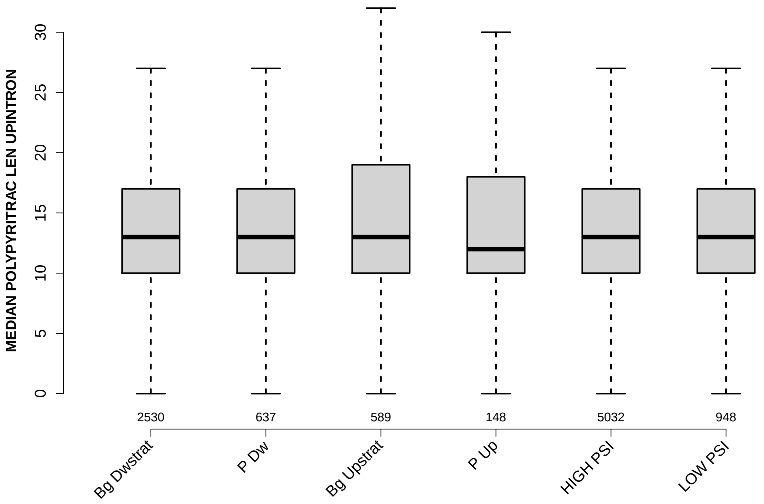

Significant results from Mann-Whitney U test:

- `Bg_Dwstrat` vs `Bg_Upstrat` : 0.0466671
  mean: 14.7225 < 16.0297 , median: 13 = 13
- `P_Dw` vs `Bg_Upstrat` : 0.00316932
  mean: 14.2865 < 16.0297 , median: 13 = 13
- `P_Dw` vs `HIGH_PSI` : 0.0278828
  mean: 14.2865 < 14.8023 , median: 13 = 13
- `Bg_Upstrat` vs `LOW_PSI` : 0.0429602
  mean: 16.0297 > 14.7558 , median: 13 = 13

#### 6.42 MEDIAN POLYPYRITRAC SCORE UPINTRON

Back to: Overview | ToC

Meaning: like POLYPYRITRAC SCORE MAXBP but median over top-3 predicted BPs

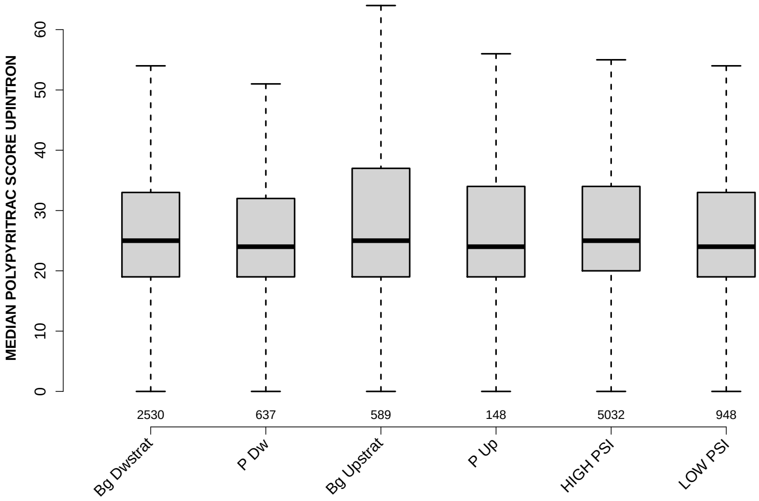

Significant results from Mann-Whitney U test:

- `P_Dw` vs `Bg_Upstrat` : 0.0440457
  mean: 28.033 < 31.0195 , median: 24 < 25
- `P_Dw` vs `HIGH_PSI` : 0.0395985
  mean: 28.033 < 28.9109 , median: 24 < 25
- `Bg_Upstrat` vs `LOW_PSI` : 0.0339269
  mean: 31.0195 > 28.2046 , median: 25 > 24
- `HIGH_PSI` vs `LOW_PSI` : 0.0209139
  mean: 28.9109 > 28.2046 , median: 25 > 24

#### 6.43 MEDIAN BPSCORE UPINTRON

Back to: Overview | ToC

Meaning: like BPSCORE MAXBP but median over top-3 predicted BPs

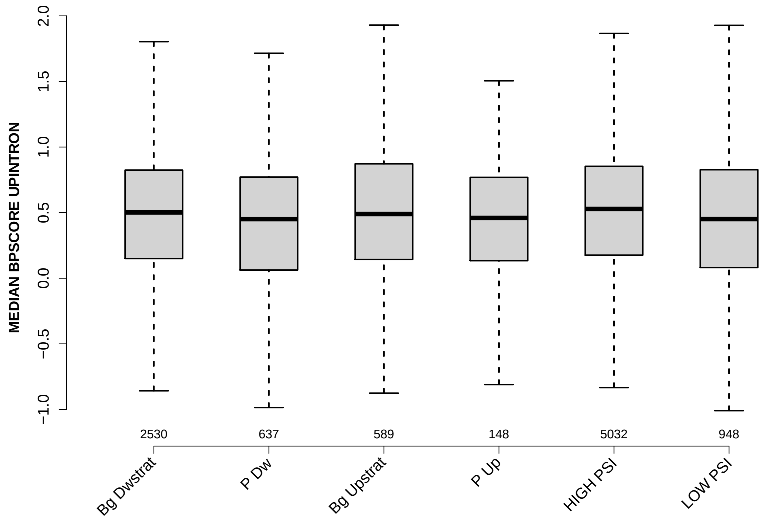

Significant results from Mann-Whitney U test:

- `Bg_Dwstrat` vs `P_Dw` : 0.0241788
  mean: 0.43016 > 0.365007 , median: 0.502223 > 0.451114
- `Bg_Dwstrat` vs `HIGH_PSI` : 0.0222001
  mean: 0.43016 < 0.467506 , median: 0.502223 < 0.52799
- `P_Dw` vs `Bg_Upstrat` : 0.0487337
  mean: 0.365007 < 0.457 , median: 0.451114 < 0.48957
- `P_Dw` vs `HIGH_PSI` : 0.000226891
  mean: 0.365007 < 0.467506 , median: 0.451114 < 0.52799
- `HIGH_PSI` vs `LOW_PSI` : 0.000273983
  mean: 0.467506 > 0.388494 , median: 0.52799 > 0.45125

#### 6.44 DIST FROM MAXBP TO 3SS DOINTRON

Back to: Overview | ToC

Meaning: distance to 3ss of best precited BP

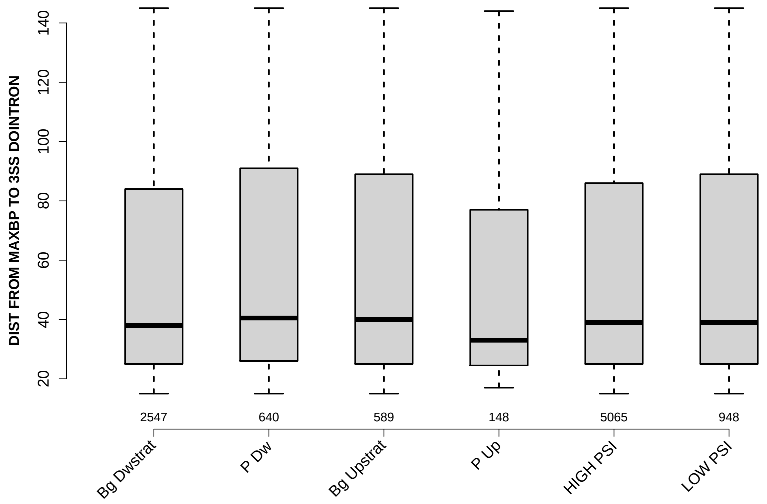

Significant results from Mann-Whitney U test:

- none

#### 6.45 SCORE FOR MAXBP SEQ DOINTRON

Back to: Overview | ToC

Meaning: BP sequence score of best predicted BP

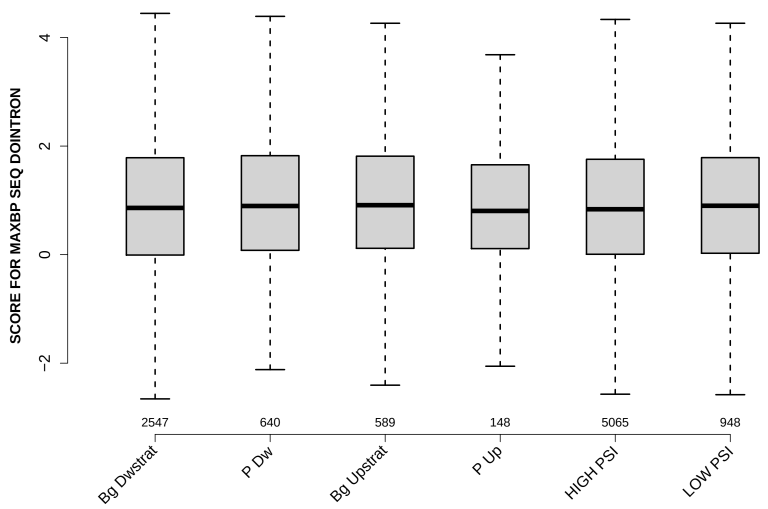

Significant results from Mann-Whitney U test:

- none

#### 6.46 PYRIMIDINECONT MAXBP DOINTRON

Back to: Overview | ToC

Meaning: Pyrimidine content between the BP adenine and the 3 prime splice site for best BP

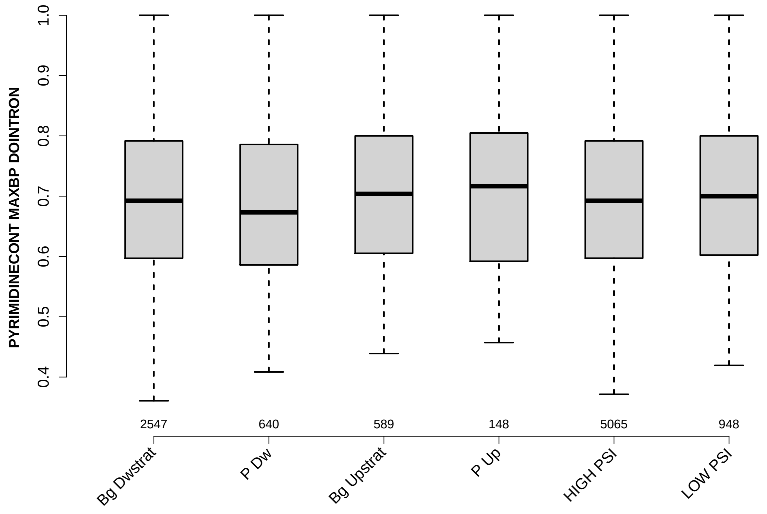

Significant results from Mann-Whitney U test:

- `Bg_Dwstrat` vs `P_Dw` : 0.0407524
  mean: 0.697584 > 0.687164 , median: 0.692308 > 0.673286
- `P_Dw` vs `Bg_Upstrat` : 0.00685557
  mean: 0.687164 < 0.706337 , median: 0.673286 < 0.703704
- `P_Dw` vs `HIGH_PSI` : 0.0312103
  mean: 0.687164 < 0.697834 , median: 0.673286 < 0.692308
- `P_Dw` vs `LOW_PSI` : 0.0125907
  mean: 0.687164 < 0.702356 , median: 0.673286 < 0.7

#### 6.47 POLYPYRITRAC OFFSET MAXBP DOINTRON

Back to: Overview | ToC

Meaning: Polypyrimidine track offset relative to the BP adenine for best BP

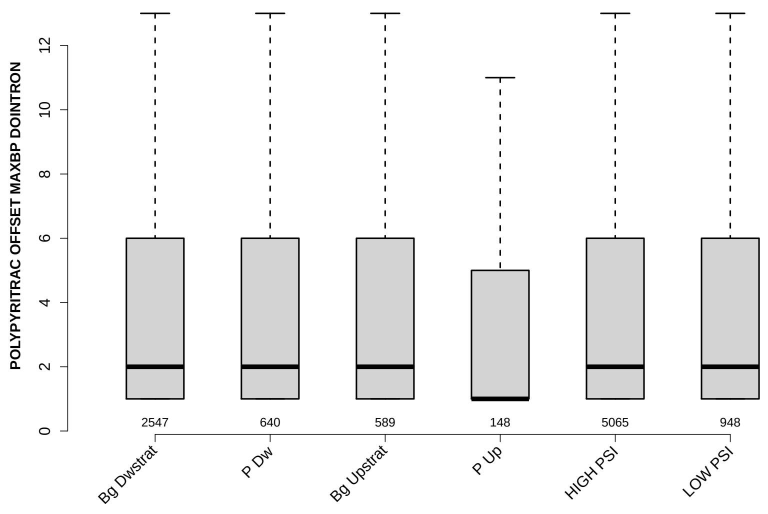

Significant results from Mann-Whitney U test:

- none

#### 6.48 POLYPYRITRAC LEN MAXBP DOINTRON

Back to: Overview | ToC

Meaning: Polypyrimidine track length for best BP

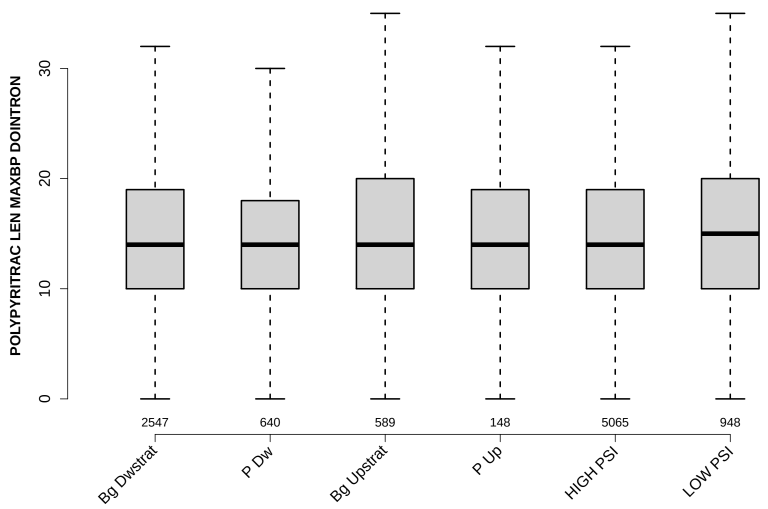

Significant results from Mann-Whitney U test:

- `Bg_Dwstrat` vs `LOW_PSI` : 0.00564143
  mean: 15.3581 < 16.3228 , median: 14 < 15
- `P_Dw` vs `Bg_Upstrat` : 0.0307758
  mean: 15.0047 < 16.3379 , median: 14 = 14
- `P_Dw` vs `LOW_PSI` : 0.00556986
  mean: 15.0047 < 16.3228 , median: 14 < 15
- `HIGH_PSI` vs `LOW_PSI` : 0.0125303
  mean: 15.4693 < 16.3228 , median: 14 < 15

#### 6.49 POLYPYRITRAC SCORE MAXBP DOINTRON

Back to: Overview | ToC

Meaning: Polypyrimidine track score for best BP

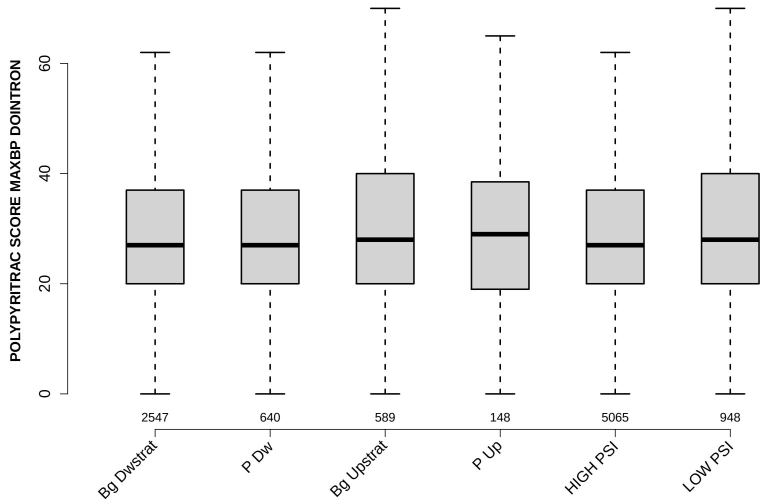

Significant results from Mann-Whitney U test:

- Bg_Dwstrat vs Bg_Upstrat : 0.0303814
  mean: 30.1786 < 32.601 , median: 27 < 28
- Bg_Dwstrat vs LOW_PSI : 0.0128559
  mean: 30.1786 < 32.2954 , median: 27 < 28
- P_Dw vs Bg_Upstrat : 0.0454645
  mean: 29.7922 < 32.601 , median: 27 < 28
- P_Dw vs LOW_PSI : 0.0297366
  mean: 29.7922 < 32.2954 , median: 27 < 28
- Bg_Upstrat vs HIGH_PSI : 0.0429166
  mean: 32.601 > 30.3889 , median: 28 > 27
- HIGH_PSI vs LOW_PSI : 0.0173619
  mean: 30.3889 < 32.2954 , median: 27 < 28

#### 6.50 BPSCORE MAXBP DOINTRON

Back to: Overview | ToC

Meaning: SVM classification score of best BP

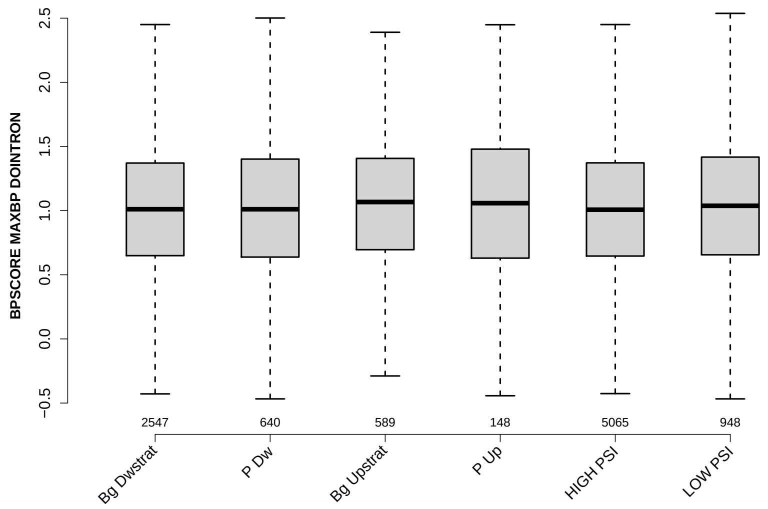

Significant results from Mann-Whitney U test:

- `Bg_Upstrat` vs `HIGH_PSI` : 0.033014
  mean: 1.067 > 1.0045 , median: 1.0671 > 1.0072

#### 6.51 NUM PREDICTED BPS DOINTRON

Back to: Overview | ToC

Meaning: number of all predicted BPs which have a positive BP score

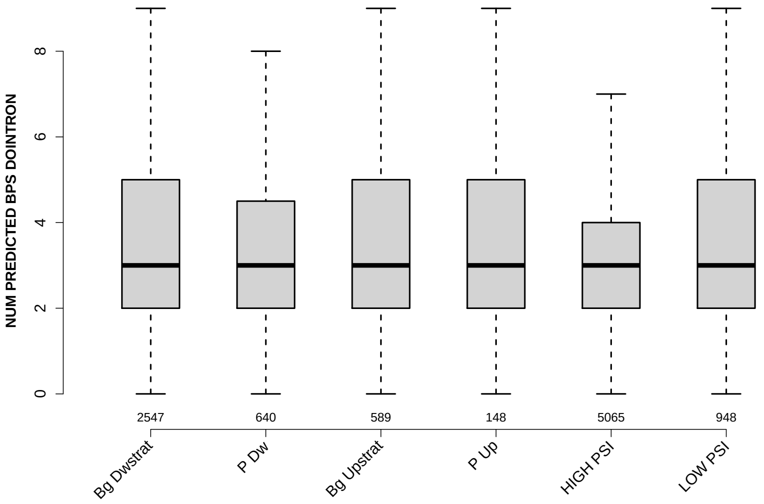

Significant results from Mann-Whitney U test:

- none

#### 6.52 MEDIAN DIST FROM BP TO 3SS DOINTRON

Back to: Overview | ToC

Meaning: like DIST FROM MAXBP TO 3SS but median over top-3 predicted BPs

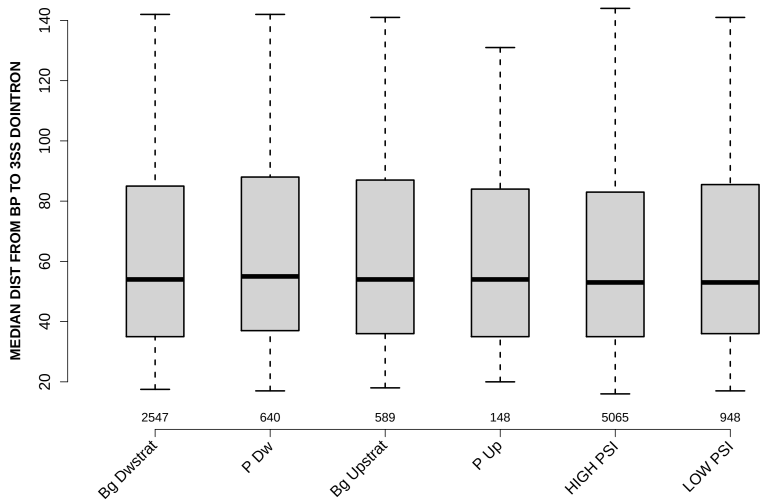

Significant results from Mann-Whitney U test:

- none

#### 6.53 MEDIAN SCORE FOR BPSEQ DOINTRON

Back to: Overview | ToC

Meaning: like SCORE FOR MAXBP SEQ but median over top-3 predicted BPs

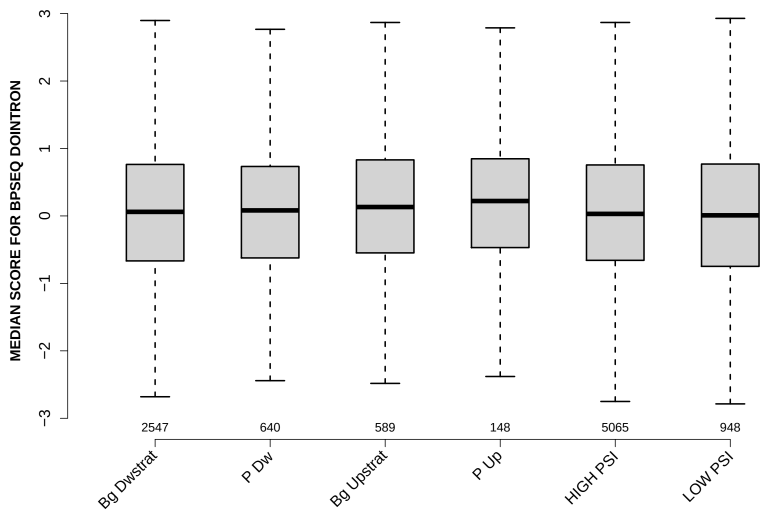

Significant results from Mann-Whitney U test:

- `Bg_Upstrat` vs `HIGH_PSI` : 0.0284286
  mean: 0.171096 > 0.0682715 , median: 0.132013 > 0.0303217
- `Bg_Upstrat` vs `LOW_PSI` : 0.0156659
  mean: 0.171096 > 0.0385431 , median: 0.132013 > 0.00879501

#### 6.54 MEDIAN PYRIMIDINECONT DOINTRON

Back to: Overview | ToC

Meaning: like PYRIMIDINECONT MAXBP but median over top-3 predicted BPs

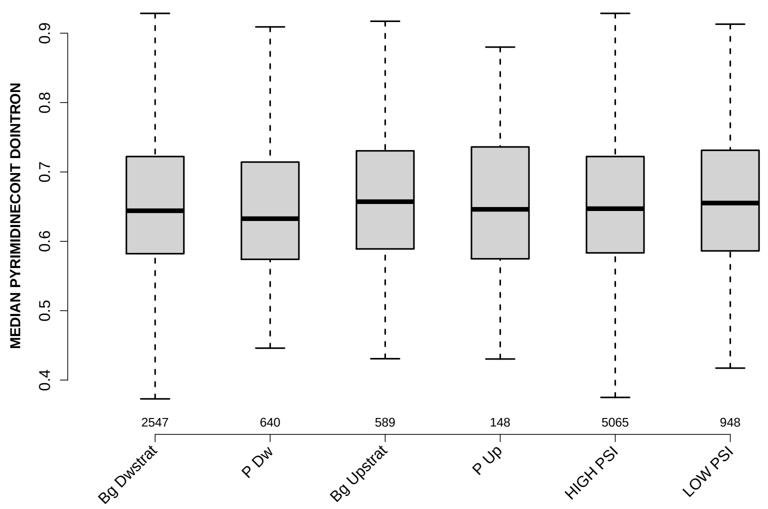

Significant results from Mann-Whitney U test:

- P_Dw vs Bg_Upstrat : 0.0102178
  mean: 0.648239 < 0.661454 , median: 0.632653 < 0.657143
- P_Dw vs HIGH_PSI : 0.049135
  mean: 0.648239 < 0.654977 , median: 0.632653 < 0.647059
- P_Dw vs LOW_PSI : 0.00982767
  mean: 0.648239 < 0.660625 , median: 0.632653 < 0.655172

#### 6.55 MEDIAN POLYPYRITRAC OFFSET DOINTRON

Back to: Overview | ToC

Meaning: like POLYPYRITRAC OFFSET MAXBP but median over top-3 predicted BPs

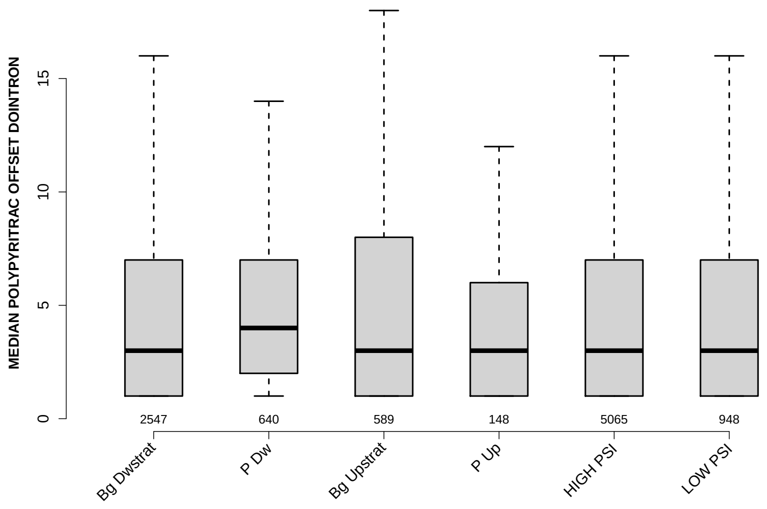

Significant results from Mann-Whitney U test:

- `P_Dw` vs `LOW_PSI` : 0.0364511
  mean: 5.6344 > 5.2627 , median: 4 > 3

#### 6.56 MEDIAN POLYPYRITRAC LEN DOINTRON

Back to: Overview | ToC

Meaning: like POLYPYRITRAC LEN MAXBP but median over top-3 predicted BPs

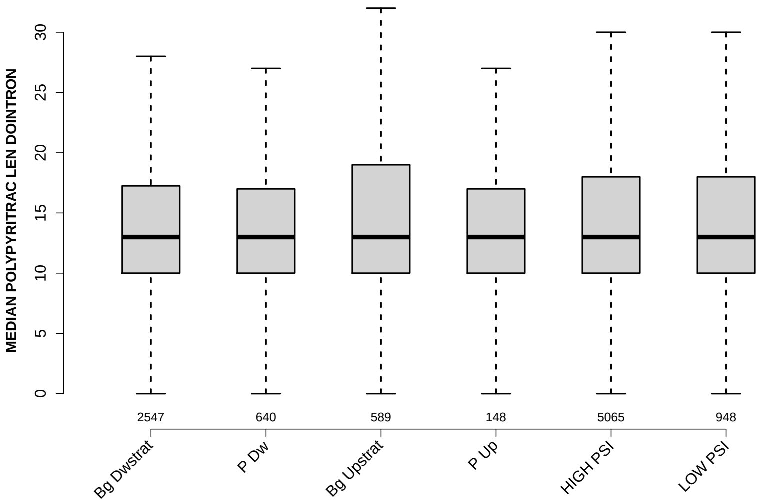

Significant results from Mann-Whitney U test:

- `P_Dw` vs `LOW_PSI` : 0.0243995
  mean: $14.3352 < 15.4014$ , median: $13 = 13$

#### 6.57 MEDIAN POLYPYRITRAC SCORE DOINTRON

Back to: Overview | ToC

Meaning: like POLYPYRITRAC SCORE MAXBP but median over top-3 predicted BPs

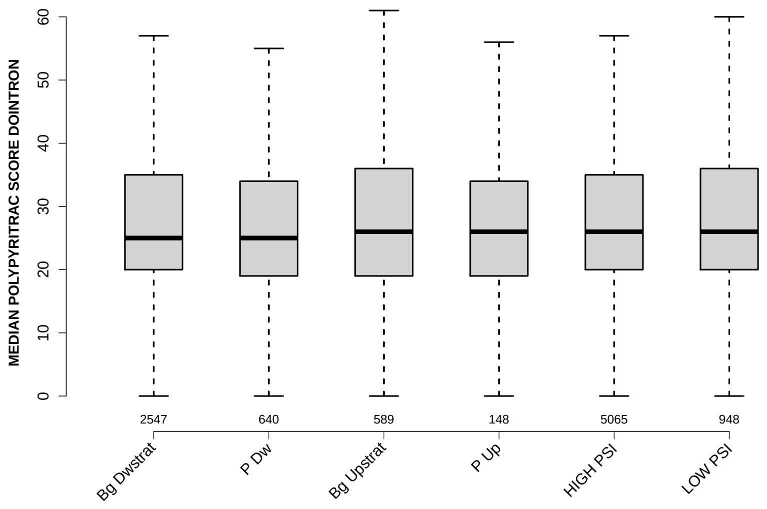

Significant results from Mann-Whitney U test:

- `P_Dw` vs `LOW_PSI` : 0.0452168
  mean: 28.1891 < 30.4188 , median: 25 < 26

#### 6.58 MEDIAN BPSCORE DOINTRON

Back to: Overview | ToC

Meaning: like BPSCORE MAXBP but median over top-3 predicted BPs

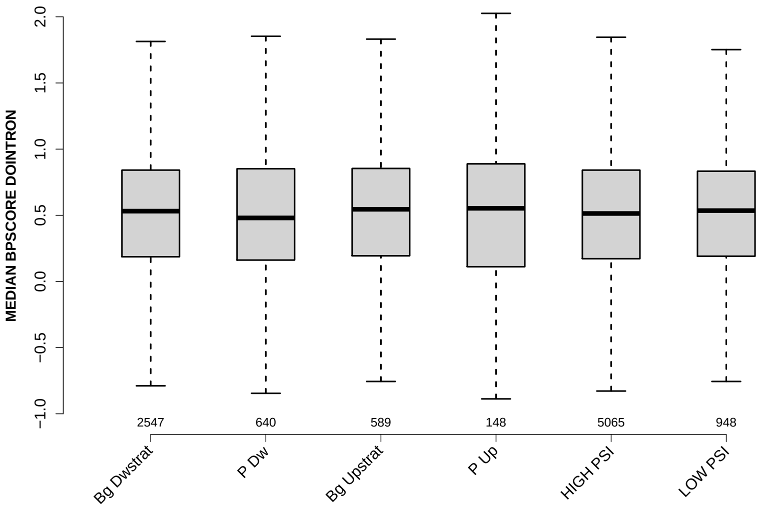

Significant results from Mann-Whitney U test:

- none

#### 6.59 MEDIAN TR LENGTH

Back to: Overview | ToC

Meaning: median length of transcripts the exon occurs in

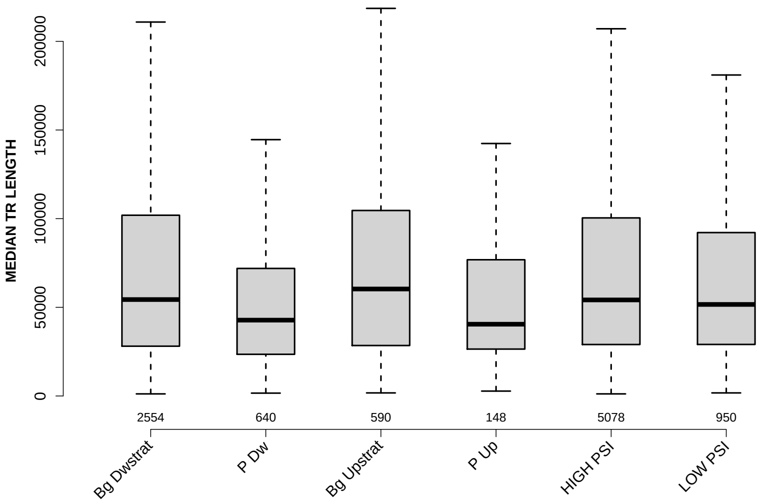

Significant results from Mann-Whitney U test:

- Bg_Dwstrat vs P_Dw : 2.17195e-10
  mean: 82663.7128 > 57174.4734 , median: 54372.5 > 42765.5
- Bg_Dwstrat vs P_Up : 0.0102901
  mean: 82663.7128 > 61367.6757 , median: 54372.5 > 40468.5
- P_Dw vs Bg_Upstrat : 9.22794e-09
  mean: 57174.4734 < 82523.728 , median: 42765.5 < 60329.5
- P_Dw vs HIGH_PSI : 3.12535e-12
  mean: 57174.4734 < 83525.7209 , median: 42765.5 < 54167
- P_Dw vs LOW_PSI : 7.65318e-07
  mean: 57174.4734 < 73517.1747 , median: 42765.5 < 51617
- Bg_Upstrat vs P_Up : 0.003726
  mean: 82523.728 > 61367.6757 , median: 60329.5 > 40468.5
- P_Up vs HIGH_PSI : 0.00630021
  mean: 61367.6757 < 83525.7209 , median: 40468.5 < 54167
- P_Up vs LOW_PSI : 0.0332589
  mean: 61367.6757 < 73517.1747 , median: 40468.5 < 51617

#### 6.60 MEDIAN EXON NUMBER

Back to: Overview | ToC

Meaning: ... of transcripts where exon was found in

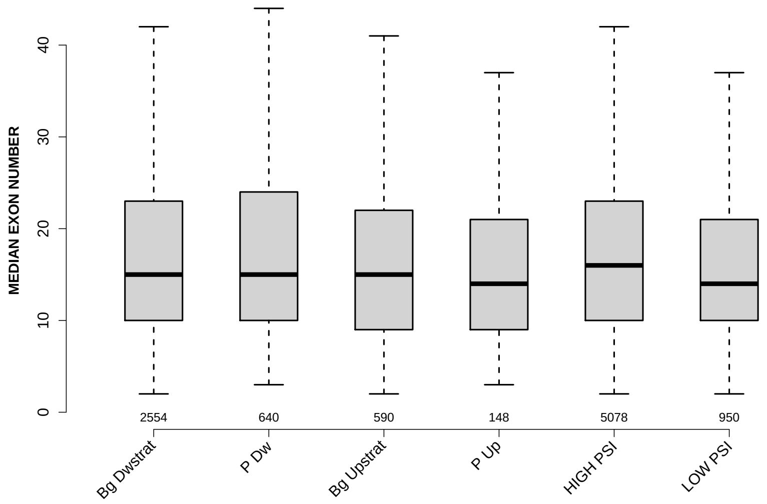

Significant results from Mann-Whitney U test:

- Bg_Dwstrat vs LOW_PSI : 0.00481114
  mean: 18.211 > 16.3842 , median: 15 > 14
- Bg_Upstrat vs HIGH_PSI : 0.00652098
  mean: 17.1237 < 18.1003 , median: 15 < 16
- P_Up vs HIGH_PSI : 0.0212916
  mean: 16.0169 < 18.1003 , median: 14 < 16
- HIGH_PSI vs LOW_PSI : 3.74643e-05
  mean: 18.1003 > 16.3842 , median: 16 > 14

#### 6.61 EXON MEDIANRELATIVERANK

Back to: Overview | ToC

Meaning: relative rank = rank / number of all exons in transcript, is between 0 and 1

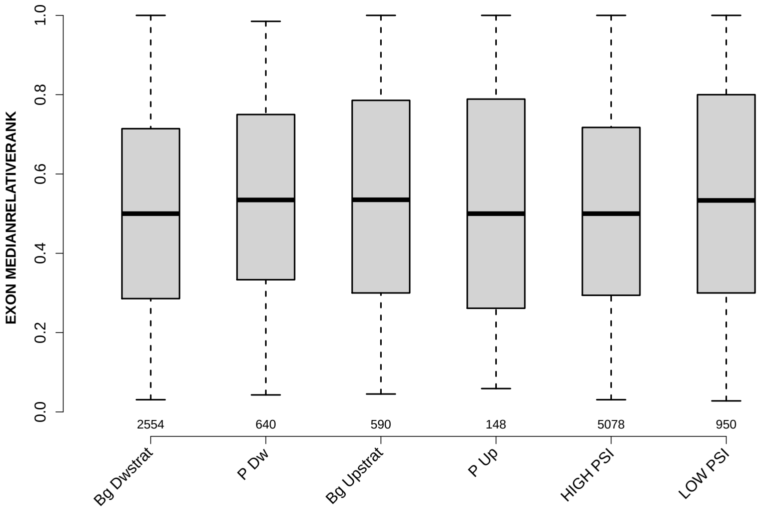

Significant results from Mann-Whitney U test:

- `Bg_Dwstrat` vs `P_Dw` : 0.00843201
  mean: 0.503215 < 0.532238 , median: 0.5 < 0.534524
- `Bg_Dwstrat` vs `Bg_Upstrat` : 0.00106206
  mean: 0.503215 < 0.546861 , median: 0.5 < 0.534959
- `Bg_Dwstrat` vs `LOW_PSI` : 2.44172e-05
  mean: 0.503215 < 0.54969 , median: 0.5 < 0.533333
- `P_Dw` vs `HIGH_PSI` : 0.0236867
  mean: 0.532238 > 0.508495 , median: 0.534524 > 0.5
- `Bg_Upstrat` vs `HIGH_PSI` : 0.00278233
  mean: 0.546861 > 0.508495 , median: 0.534959 > 0.5
- `HIGH_PSI` vs `LOW_PSI` : 6.71162e-05
  mean: 0.508495 < 0.54969 , median: 0.5 < 0.533333

#### 6.62 EXON MEDIANRELATIVERANK 3BINS

Back to: Overview | ToC

Meaning: median bin into which EXON MEDIANRELATIVERANK falls when binning 0-1 into 3 bins

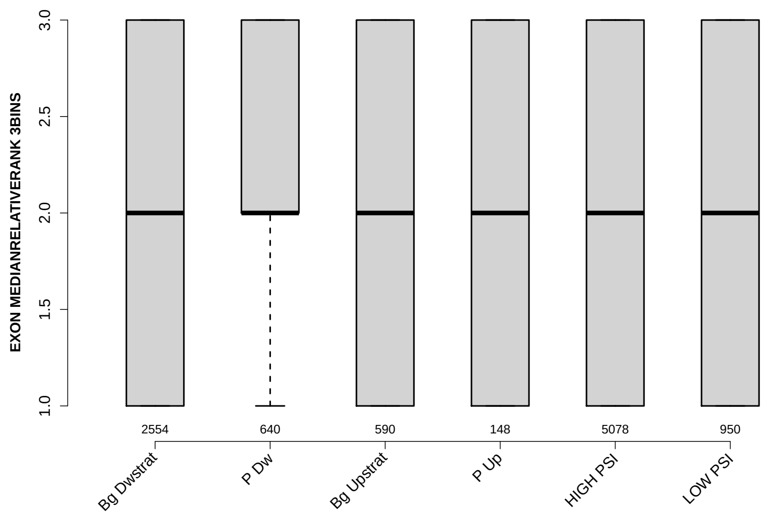

Significant results from Mann-Whitney U test:

- Bg_Dwstrat vs P_Dw : 0.00786406
  mean: 2.0294 < 2.1203 , median: 2 = 2
- Bg_Dwstrat vs Bg_Upstrat : 0.0139429
  mean: 2.0294 < 2.1153 , median: 2 = 2
- Bg_Dwstrat vs LOW_PSI : 0.00240031
  mean: 2.0294 < 2.1179 , median: 2 = 2
- P_Dw vs HIGH_PSI : 0.0226653
  mean: 2.1203 > 2.0461 , median: 2 = 2
- Bg_Upstrat vs HIGH_PSI : 0.0357473
  mean: 2.1153 > 2.0461 , median: 2 = 2
- HIGH_PSI vs LOW_PSI : 0.00772393
  mean: 2.0461 < 2.1179 , median: 2 = 2

#### 6.63 EXON MEDIANRELATIVERANK 5BINS

Back to: Overview | ToC

Meaning: similar to EXON MEDIANRELATIVERANK 3BINS with 5 bins

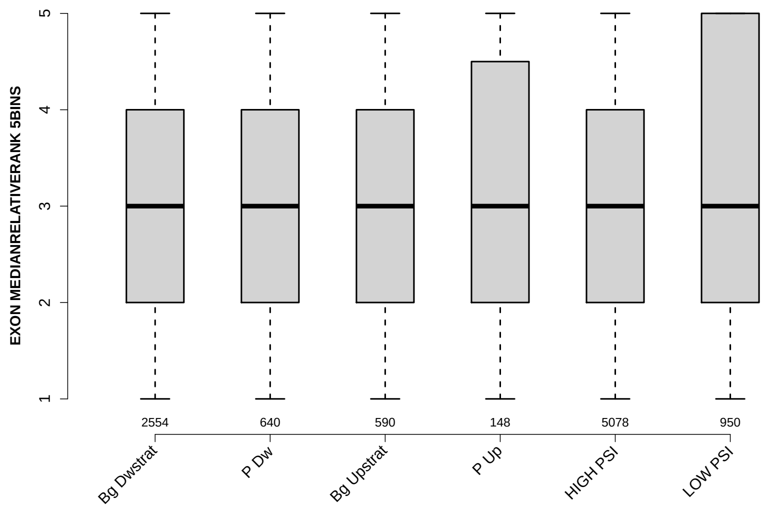

Significant results from Mann-Whitney U test:

- Bg_Dwstrat vs P_Dw : 0.00693677
  mean: 3.0129 < 3.1703 , median: 3 = 3
- Bg_Dwstrat vs Bg_Upstrat : 0.00305482
  mean: 3.0129 < 3.1949 , median: 3 = 3
- Bg_Dwstrat vs LOW_PSI : 7.41081e-05
  mean: 3.0129 < 3.2179 , median: 3 = 3
- P_Dw vs HIGH_PSI : 0.0382326
  mean: 3.1703 > 3.0565 , median: 3 = 3
- Bg_Upstrat vs HIGH_PSI : 0.0174283
  mean: 3.1949 > 3.0565 , median: 3 = 3
- HIGH_PSI vs LOW_PSI : 0.000745985
  mean: 3.0565 < 3.2179 , median: 3 = 3

#### 6.64 EXON MEDIANRELATIVERANK 10BINS

Back to: Overview | ToC

Meaning: similar to EXON MEDIANRELATIVERANK 3BINS with 10 bins

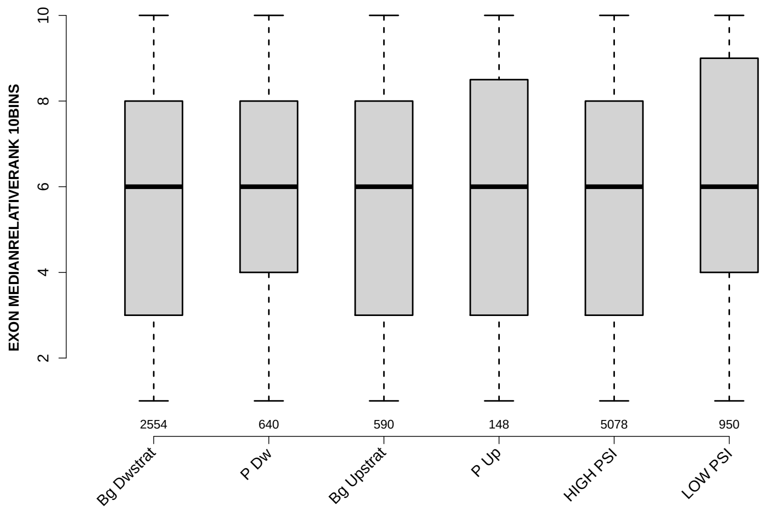

Significant results from Mann-Whitney U test:

- Bg_Dwstrat vs P_Dw : 0.00854558
  mean: 5.5619 < 5.8531 , median: 6 = 6
- Bg_Dwstrat vs Bg_Upstrat : 0.00140939
  mean: 5.5619 < 5.9525 , median: 6 = 6
- Bg_Dwstrat vs LOW_PSI : 3.90084e-05
  mean: 5.5619 < 5.9905 , median: 6 = 6
- P_Dw vs HIGH_PSI : 0.0286517
  mean: 5.8531 > 5.6219 , median: 6 = 6
- Bg_Upstrat vs HIGH_PSI : 0.00436713
  mean: 5.9525 > 5.6219 , median: 6 = 6
- HIGH_PSI vs LOW_PSI : 0.000137933
  mean: 5.6219 < 5.9905 , median: 6 = 6

#### 6.65 NTRS ALL FOR GENE

Back to: Overview | ToC

Meaning: number of transcripts of gene where the exon was found in

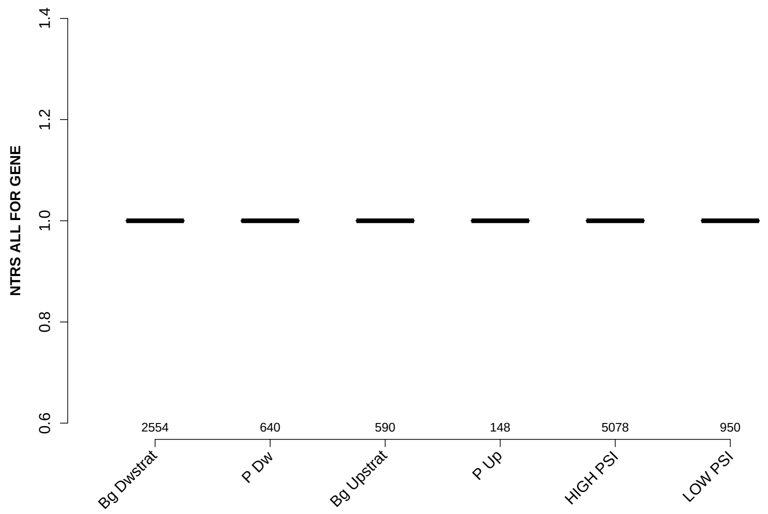

Significant results from Mann-Whitney U test:

- Bg_Dwstrat vs Bg_Upstrat : 0.0124415
  mean: 1.1222 < 1.1627 , median: 1 = 1
- P_Dw vs Bg_Upstrat : 0.00478528
  mean: 1.1047 < 1.1627 , median: 1 = 1
- Bg_Upstrat vs HIGH_PSI : 0.00856443
  mean: 1.1627 > 1.1217 , median: 1 = 1
- Bg_Upstrat vs LOW_PSI : 0.0215088
  mean: 1.1627 > 1.1116 , median: 1 = 1

#### 6.66 PROP FIRST EXON

Back to: Overview | ToC

Meaning: NTRS WITH EXON AS FIRST EXON / NTRS WITH EXON

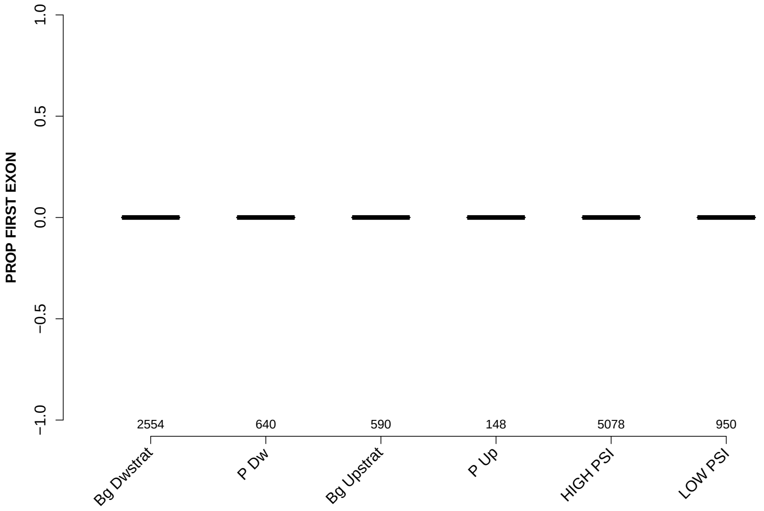

Significant results from Mann-Whitney U test:

- `Bg_Dwstrat` vs `Bg_Upstrat` : 0.020724
  mean: 0.0082224 > 0 , median: 0 = 0
- `Bg_Dwstrat` vs `LOW_PSI` : 0.0033461
  mean: 0.0082224 > 0 , median: 0 = 0
- `Bg_Upstrat` vs `P_Up` : p value = NA
- `Bg_Upstrat` vs `HIGH_PSI` : 0.0285072
  mean: 0 < 0.00771301 , median: 0 = 0
- `Bg_Upstrat` vs `LOW_PSI` : p value = NA
- `P_Up` vs `LOW_PSI` : p value = NA
- `HIGH_PSI` vs `LOW_PSI` : 0.00545773
  mean: 0.00771301 > 0 , median: 0 = 0

#### 6.67 PROP LAST EXON

Back to: Overview | ToC

Meaning: NTRS WITH EXON AS LAST EXON / NTRS WITH EXON

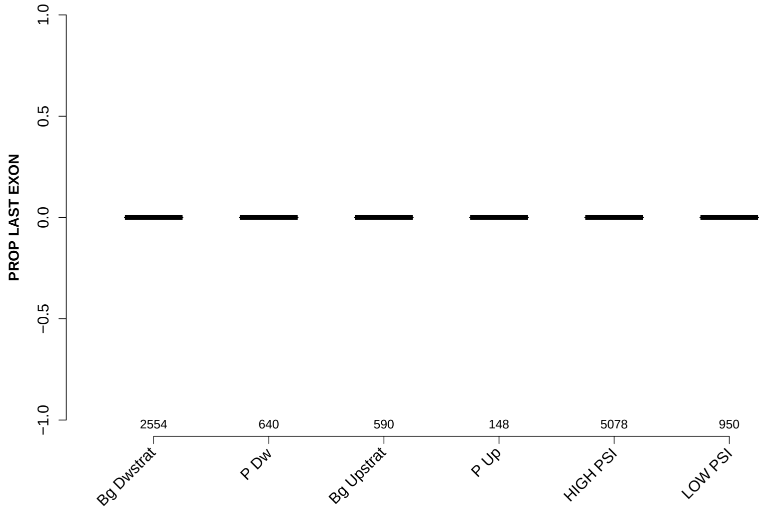

Significant results from Mann-Whitney U test:

- P_Dw vs P_Up : 0.0378707
  mean: $0 < 0.00337838$ , median: $0 = 0$

#### 6.68 PROP INTERNAL EXON

Back to: Overview | ToC

Meaning: NTRS WITH EXON AS INTERNAL EXON / NTRS WITH EXON

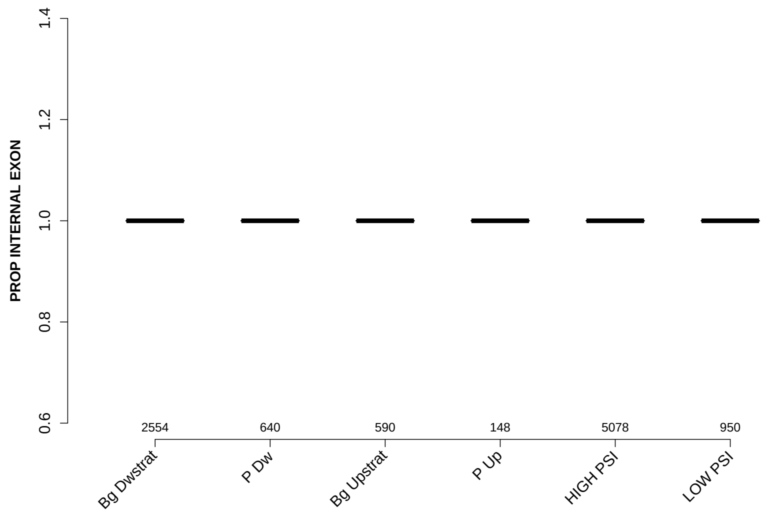

Significant results from Mann-Whitney U test:

- `Bg_Dwstrat` vs `P_Dw` : 0.0440165
  mean: 0.989233 < 0.996875 , median: 1 = 1
- `Bg_Dwstrat` vs `LOW_PSI` : 0.00617901
  mean: 0.989233 < 0.998947 , median: 1 = 1
- `HIGH_PSI` vs `LOW_PSI` : 0.0176329
  mean: 0.990908 < 0.998947 , median: 1 = 1

#### 6.69 PROP EXON IN UTR

Back to: Overview | ToC

Meaning: NTRS WITH EXON IN UTR / NTRS WITH EXON

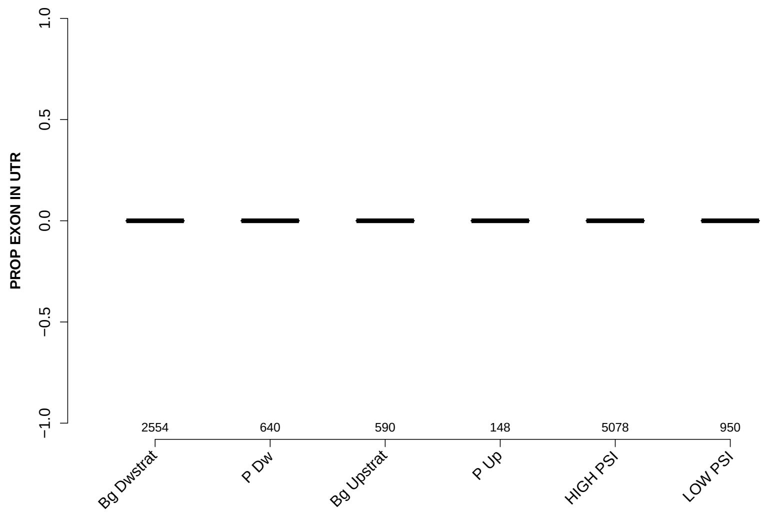

Significant results from Mann-Whitney U test:

- Bg_Dwstrat vs Bg_Upstrat : 6.07913e-06
  mean: 0.0429392 > 0.0059322 , median: 0 = 0
- Bg_Dwstrat vs LOW_PSI : 9.7125e-10
  mean: 0.0429392 > 0.00210526 , median: 0 = 0
- P_Dw vs Bg_Upstrat : 0.00305237
  mean: 0.0265625 > 0.0059322 , median: 0 = 0
- P_Dw vs LOW_PSI : 2.82656e-05
  mean: 0.0265625 > 0.00210526 , median: 0 = 0
- Bg_Upstrat vs HIGH_PSI : 1.60438e-05
  mean: 0.0059322 < 0.0393528 , median: 0 = 0
- P_Up vs LOW_PSI : 0.0222935
  mean: 0.0135135 > 0.00210526 , median: 0 = 0
- HIGH_PSI vs LOW_PSI : 4.33877e-09
  mean: 0.0393528 > 0.00210526 , median: 0 = 0
